# navigate: an open-source platform for smart light-sheet microscopy

**DOI:** 10.1101/2024.02.09.579083

**Authors:** Zach Marin, Xiaoding Wang, Dax W. Collison, Conor McFadden, Jinlong Lin, Hazel Borges, Bingying Chen, Dushyant Mehra, Qionghua Shen, Seweryn Galecki, Stephan Daetwyler, Reto Fiolka, Kevin M. Dean

## Abstract

***navigate*** is a turnkey, open-source software solution designed to enhance light-sheet fluorescence microscopy (LSFM) by integrating smart microscopy techniques into a user-friendly framework. It enables automated, intelligent imaging with a Python-based control system that supports GUI-reconfigurable acquisition routines and the integration of diverse hardware sets. As a comprehensive package, navigate democratizes access to advanced LSFM capabilities, facilitating the development and implementation of smart microscopy workflows without requiring deep programming knowledge or specialized expertise in light-sheet microscopy.

## Introduction

LSFM enables fast, gentle, and large volume imaging, making it an increasingly important tool for cell biological discovery *in vitro* and *in vivo*. In both live cells and fixed tissues, many important cellular interactions and microenvironmental niches are exceedingly rare. Automatic identification of such events is computationally expensive, especially if such analyses are performed for many biological replicates or through time. In living specimens, such as developing embryos, cellular dynamics often require an experimenter to not only detect the initial event, but manually update the microscope’s configuration to track movement or risk losing the sample. To combat such challenges, researchers have started to apply smart microscopy techniques to LSFM^1-4^. Despite smart imaging approaches existing for over a decade^5^, most LSFMs operate in a stubbornly classical operation paradigm. In part, this is because each LSFM is optimized for specific biological contexts with distinct hardware and control requirements, which may not be supported by existing smart routines. Compounding the problem, most LSFM control software has been built in a piece-meal fashion by combining closed-source LabView or Java code with separate Java or Python routines that must be installed and/or compiled separately (Table S1), making it difficult for users without advanced programming capabilities to adapt and reuse available smart routines. To address these challenges and democratize smart LSFM, we developed ***navigate***, an open-source, Python-based LSFM control software with easily reconfigured hardware controls and GUI-reconfigurable smart acquisition routines (Fig. 1A & B).

**Figure 1:**
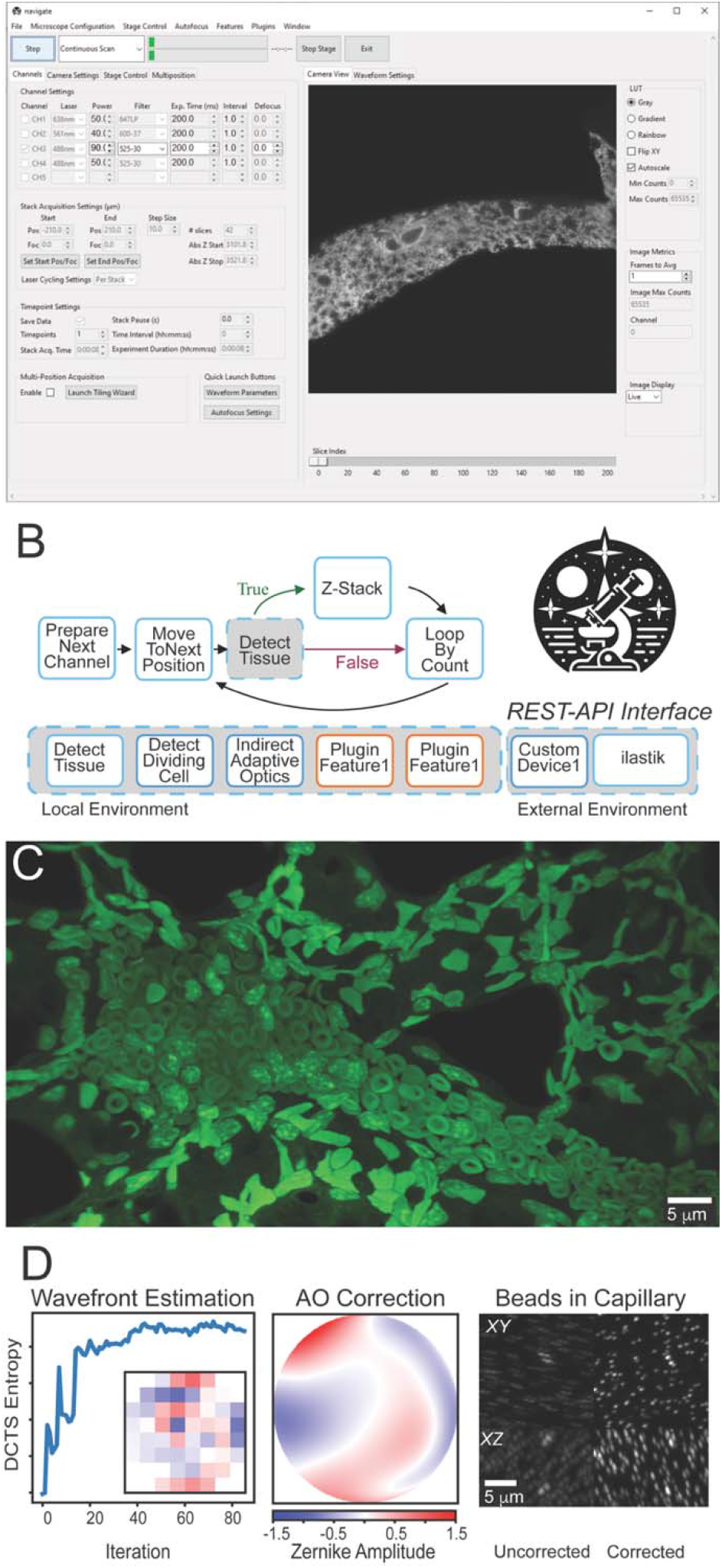
navigate provides a GUI with access to reconfigurable smart acquisition pipelines and runs on multiple types of light-sheet configurations. (A) A screenshot of the navigate GUI running on a mesoSPIM^11^ with the Channels and Camera View tabs selected (see the user guide in Supplementary Information for a full description). (B) An example of a reconfigurable, decision-based pipeline that takes a Z-Stack if there is tissue found at a particular stage position. The user guide in the Supplementary Note features a tutorial on how to build this pipeline. Individual features can be swapped for other features that ship with the software or with custom-built plugin features. (C) Example image of an expanded mouse lung, captured with CT-ASLM using navigate. (D) navigate can integrate advanced optical techniques, such as adaptive optics. (Left) An iterative sensorless adaptive optics scheme uses the Discrete Cosine Transform Shannon (DCTS) Entropy to estimate aberrations. Inset shows the final deformable mirror positions. (Middle) Final estimated adaptive optics wavefront. (Right) Fluorescent beads within a capillary, before and after adaptive optics, in both XY and XZ directions.

## Results

***navigate*** enables biologists and technology developers alike to establish and re-use smart microscopy pipelines on diverse sets of hardware from within a single framework. While generalizable, Python-based frameworks for smart microscopy have been built, they are designed for stimulated emission depletion^6^ or single-molecule localization microscopy^7^ and do not yet address LSFM’s specific acquisition challenges, including decoupled illumination and detection optomechanics and a lack of an optical substrate for focus maintenance. While another framework with a GUI-editable image post-processing pipeline exists^7^, to the authors’ knowledge, ***navigate*** is the only acquisition software in which decision-based acquisition routines based on image processing results can be modified from within a GUI (see Table S1). For image-based feedback, custom analysis routines can be implemented in a plugin to work within ***navigate’s*** environment. ***navigate*** supports the addition of REST-API interfaces for two-way communication with image analysis programs running outside of Python, or in different Python environments, such as Ilastik^8^, enabling developers to make calls to state-of-the-art software while avoiding dependency conflicts. The plugin architecture of ***navigate*** also facilitates the addition of new hardware, empowering users to integrate otherwise unsupported devices. We believe this flexibility fosters the software’s natural evolution and its capacity to seamlessly accommodate the diverse modalities and acquisition formats inherent to LSFM technology.

***navigate*** can readily be installed and operated without knowledge of software development or an expertise in light-sheet microscopy. Out of the box, it is equipped with an intuitive and rigorously tested graphical user interface (GUI) and is capable of capturing rich biological data (Fig. 1C) from axially swept, digitally scanned, oblique plane, field synthesis, projection and confocal projection light-sheet imaging modalities in both laser– and sample-scanning formats. ***navigate*** virtualizes microscope control, allowing multiple microscopes to be operated on the fly using a single set of hardware. For optically complex specimens, ***navigate*** provides indirect optimization routines for sensorless adaptive optics (Fig. 1***D***). It writes to next-generation, pyramidal BigDataViewer H5/N5 data formats^9,10^, enabling easy movement from acquisition into established image registration and analysis pipelines and fast visual feedback in ImageJ. Data generated adheres to Open Microscopy Environment standards and imports readily into OMERO image data repositories. To ensure academic reproducibility, consistency, and reliability, ***navigate*** is well documented, and each release of ***navigate*** is versioned, archived, and given a citable digital object identifier. We believe ***navigate’s*** extensibility, user-friendliness even for non-experts, and reproducibility positions ***navigate*** as the turn-key solution for smart LSFM.

The source code for ***navigate*** is available at https://github.com/TheDeanLab/navigate and queries and feedback can be provided by creating an issue on GitHub. Documentation on how to extend ***navigate*** is provided in the Supplementary Note (and is kept up to date at https://thedeanlab.github.io/navigate/) and a plugin template is available at https://github.com/TheDeanLab/navigate-plugin-template. While initially designed for LSFM, navigate possesses a versatile architecture that makes it suitable for a broad range of camera-based imaging modalities. Thus, we hope ***navigate*** empowers users from all scientific backgrounds to construct and implement novel intelligent imaging workflows, greatly improving our ability to unravel the cellular complexities of life.

## Acknowledgments

This work was funded by the NIH National Institute of General Medical Science (RM1GM145399 and R35GM133522) and National Cancer Institute (U54CA268072). The authors wish to thank Drs. Gaudenz Danuser, and Felix Zhou (UT Southwestern Medical Center) for helpful discussions about software design, Drs. Andrew G. York and Nathaniel H. Thayer (Calico Life Sciences LLC) for their open-source concurrency toolkit, and Evan Wylie, Sampath Rapuri, Renil Gupta, and Samir Mamtani for their contributions to the graphical user interface.

## Author Contributions

ZM, XW, DC and KMD designed the software. ZM, XW, DC, BC, CF, DM, SD and KMD wrote the code. JL, BC, CM and HB extensively tested the software on multiple microscopes. QS, HB, and SG provided biological samples for testing. CM and RF implemented the adaptive optics routines. KD supervised the design and development of software, microscopes, and tissue preparation methods. ZM, XW and KMD wrote the manuscript. All authors discussed the project and edited the manuscript.

## Supplementary Material

Compiled online documentation.

## Supplementary Tables

**Table S1:**
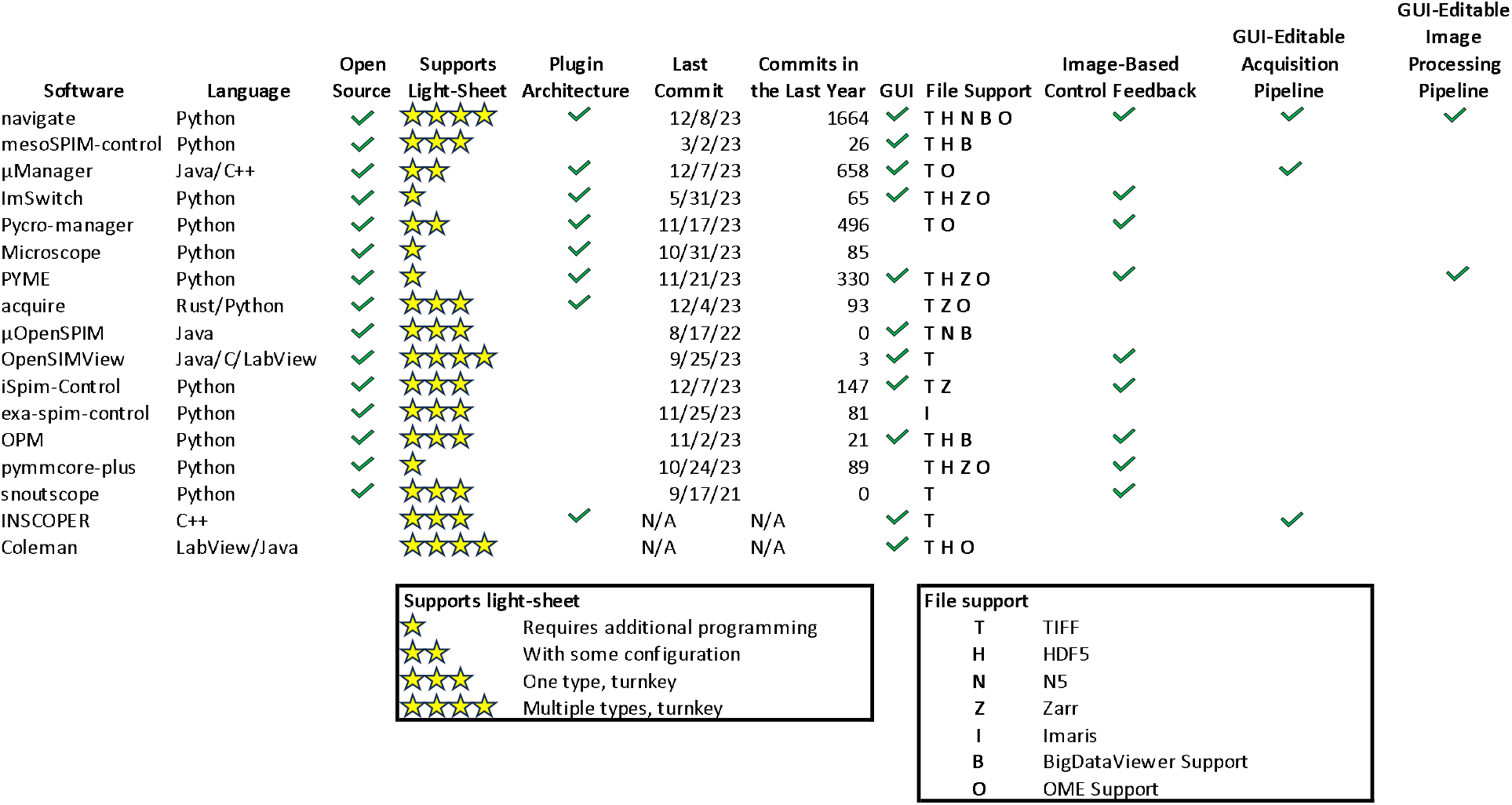
Comparative Analysis of Software Packages. A comparison with commonly used microscopy acquisition software is summarized. We believe ***navigate*** has many features that make it a desirable choice for smart, light-sheet acquisition for biologists, microscopists, and software developers. It offers:

- Turnkey support for standard hardware and multiple types of light sheets.
- A GUI-editable smart acquisition pipeline.
- A plugin architecture that lets developers easily enable new hardware controls and image-based feedback routines.
- Support for large-volume, next-generation file formats.
- An open-source license.
- Active development.

## Notes

https://github.com/TheDeanLab/navigate

https://thedeanlab.github.io/navigate/

### Competing Interest Statement

K.M.D. and R.F. have a patent covering ASLM (US10989661) and consultancy agreements with 3i, Inc (Denver, CO, USA). K.M.D. has an ownership interest in Discovery Imaging Systems, LLC.

