## Supplementary Material for "navigate: an open-source platform for smart light-sheet microscopy"

### navigate Documentation

*Release 0.0.3*

Dean Lab, UT Southwestern Medical Center

Feb 06, 2024

### GETTING STARTED

**1 Quick Start Guide** **1**

CHAPTER

ONE

### QUICK START GUIDE

This quick start guide covers how to install **navigate**, launch it in synthetic hardware mode to confirm it is working, and save an image to disk.

#### 1.1 Installation

1. *Install Miniconda.* Download and install Miniconda from the official website.
2. *Create and Activate a Conda Environment.* Launch a Miniconda Prompt (or a Terminal on MacOS) and enter the following.

```
(base) conda create -n navigate python=3.9.7
(base) conda activate navigate
```

3. *Install* **navigate**.

```
(navigate) pip install navigate-micro
```

4. *Launch* **navigate** *in synthetic hardware mode.* This will allow you to test its functionality without actual hardware.

```
(navigate) navigate -sh
```

#### 1.2 Saving a Z-Stack to Disk

To save an image to disk, follow these steps:

- Launch **navigate** in synthetic hardware mode as described above.
- Next to the the *Acquire* button on the upper left, make sure that the acquisition mode in the dropdown is set to "Continuous Scan".
- Press *Acquire* (`ctrl + enter`) and confirm that a synthetic noise image is displayed in the *Camera View* window.

**Note:** At least one channel must be selected (checkbox marked) in the *Channel Settings* window, and all of the parameters (e.g., *Power*) in that row must be populated.

- Select the *Channels* tab. In the *Stack Acquisition Settings* window in this tab, press the *Set Start Pos/Foc* button. This specifies the starting `Z` and `F` (e.g., Focus) positions for the stack acquisition.

- Select the *Stage Control* tab (`ctrl + 3`), move the `Z` stage to the desired position (e.g., `100` $\mu$m), go back to the *Channels* tab (`ctrl + 1`), and press the *Set End Pos/Foc* button. This specifies the ending `Z` and `F` positions for the stack acquisition.
- In the *Stack Acquisition Settings* frame, you can now adjust the step size, which determines the number of slices in a z-stack.
- In the *Timepoint Settings* window, select *Save Data* by marking the checkbox. If the number of timepoints is set to `1`, only a single stack will be acquired.
- Next to the the *Acquire* button on the upper left, change the acquisition mode in the dropdown to "Z-Stack", and press *Acquire* (`ctrl + enter`).

•**A *File Saving Dialog* popup window will appear.**

  - With the exception of *Notes*, all fields must be populated. Any spaces in the fields will be replaced with an underscore.
  - *Notes* is saved with the metadata, and can be useful for describing the experiment.
  - *Solvent* is useful for tissue clearing experiments.
  - *File Type* can be set to *.TIFF*, *OME-TIFF*, *H5*, or *N5*. The latter two options are pyramidal file formats that are best used for large datasets and are immediately compatible with BigDataViewer and BigStitcher.
  - Press *Acquire* to begin the acquisition.
  - Once complete, the data can be opened using standard image processing software such as Fiji.

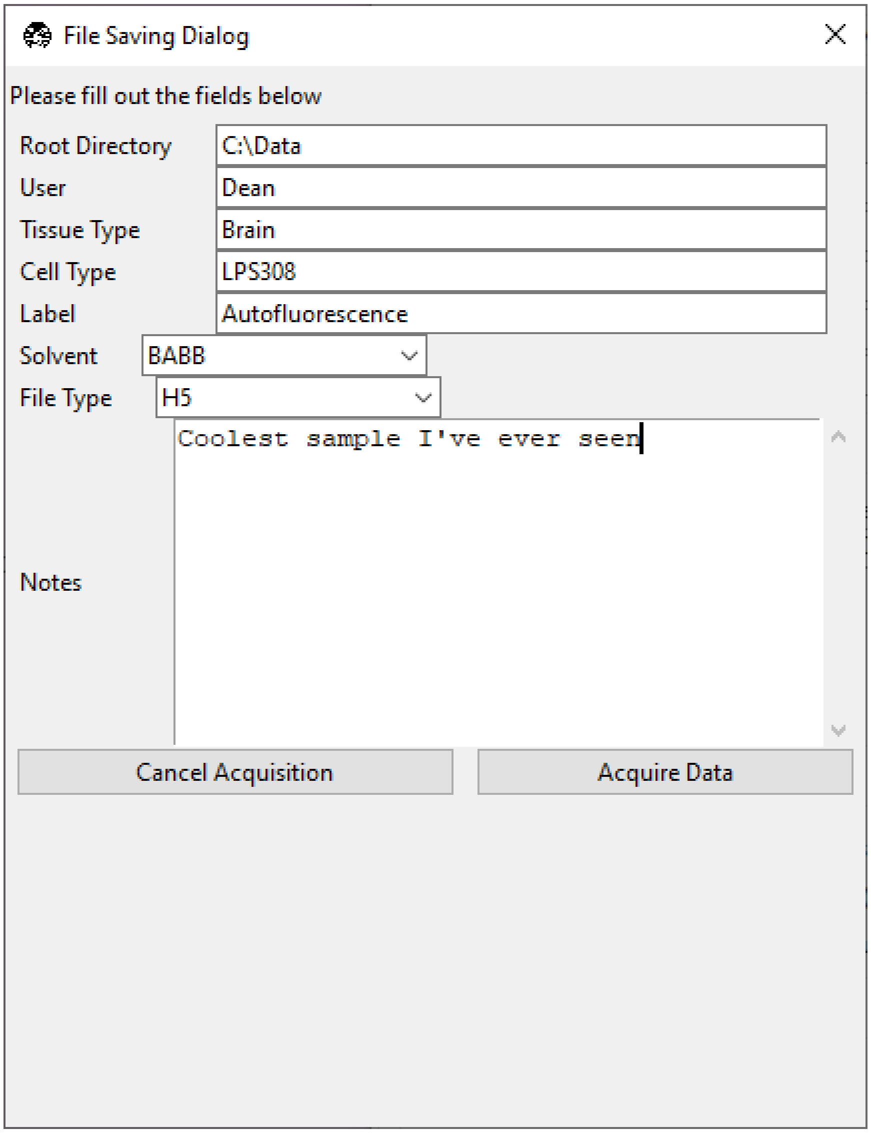
File Saving Dialog
Please fill out the fields below
Root Directory
C:\Data
User
Dean
Tissue Type
Brain
Cell Type
LPS308
Label
Autofluorescence
Solvent
BABB
File Type
H5
Notes
Coolest sample I've ever seen
Cancel Acquisition
Acquire Data

CHAPTER

TWO

### SOFTWARE INSTALLATION

#### 2.1 Computer Specifications

Below are the recommended specifications for **navigate**.

##### 2.1.1 Operating System Compatibility

**Important:** **navigate** is developed for use on Windows-based systems. This is due to the compatibility of device drivers for various microscope hardware components, such as cameras, stages, and data acquisition cards, which are predominantly designed for the Windows environment.

While it is possible to launch the software on a Mac using synthetic hardware mode, users should be aware of known issues with the Tkinter interface. These issues include improper positioning of widgets and problems with resizing the GUI window. As such, the use of **navigate** on MacOS is not recommended.

The software is untested on Linux systems.Users considering the use of **navigate** software on Linux should proceed with caution and be prepared for potential compatibility issues, especially with respect to device drivers.

**Note:** For optimal performance and compatibility, it is strongly recommended to run **navigate** on a Windows machine.

##### 2.1.2 Hardware Considerations

**navigate** will run on a mid-range laptop with at least 8 GB of RAM and a processor with two cores. Most of its operations are undemanding. Saving data at a reasonable rate, however, will require an SSD. The hardware configuration for an example microscope control machine is shown below.

**Important:** Scientific cameras are capable of rapidly generating large amounts of high-resolution data. As such, the read/write speed of the data storage device is a critical for smooth operation of the software. For example, for a standard Hamamatsu camera with a 2048 x 2048 sensor, operating at 16-bit depth and 20 frames per second, the data save rate is approximately ~167 MB/s. While such capabilities are well within the capabilities of modern SSDs, they are beyond the capabilities of most HDDs. Therefore, it is recommended to use a fast SSD data saving operations.

- ***Base Platform***
  - **Product Name**: Colfax SX6300 Workstation
  - **Colfax Part #**: CX-116263

- ***Primary and Secondary CPU***
  - **CPU Model**: Intel Xeon Silver 4215R
  - **Configuration**: 8 Cores / 16 Threads
  - **Frequency**: 3.2 GHz
  - **Cache**: 11 MB
  - **TDP**: 130W
  - **Memory Support**: 2400 MHz
- ***Memory***
  - **Type**: Registered ECC DDR4
  - **Speed**: 3200 MHz
  - **Configuration**: 16 GB per socket, 8 sockets per CPU
  - **Total RAM**: >64 GB (recommended)
- ***Operating System Drive*:**
  - **Type**: M.2 NVMe SSD
  - **Model**: Micron 7450 Max
  - **Capacity**: 800 GB
  - **Endurance**: 3 DWPD
- ***Primary Data Drive*:**
  - **Type**: NVMe SSD
  - **Model**: Samsung PM9A3
  - **Capacity**: 7.68 TB
  - **Interface**: U.2 Gen4
- ***Secondary Data Drive*:**
  - **Type**: SATA HDD
  - **Model**: Seagate Exos X20
  - **Capacity**: 20 TB
  - **Speed**: 7200 RPM
  - **Cache**: 256 MB
  - **Interface**: SATA 6.0 Gb/s
- ***Video Card***
  - **Model**: PNY nVidia T1000
  - **Memory**: 4 GB
  - **Interface**: PCI Express
- ***Network Interface***
  - **Model**: Intel X710-T2L RJ45 Copper
  - **Type**: Dual Port 10GbE

- **Interface**: PCI-E x 8

**Note:** The specifications listed are based on an example system configuration and can be adjusted based on specific needs and availability.

#### 2.2 Quick install

**Setup your Python Environment**

Head over to the miniconda website and install the appropriate version based on your operating system.

**Tip:** It is also handy to have the conda cheatsheet open when first using miniconda to get accustomed to the commands available.

- Windows: Use the Windows taskbar search to find `Anaconda Prompt (Miniconda3)`. Given how frequently you will use this, we recommend pinning it to your taskbar.
- Linux/Mac: Open a Terminal.

**Create a Python environment called navigate that uses Python version 3.9.7**

```
(base) MyComputer ~ $ conda create -n navigate python=3.9.7
```

**Activate the navigate environment**

```
(base) MyComputer ~ $ conda activate navigate
```

The active environment is shown in parentheses on the far-left. Originally, we were in the miniconda `(base)` environment. After activating the navigate environment, it should now show `(navigate)`.

**Install navigate via pip**

To install the latest stable release of **navigate**, run the following command:

```
(navigate) MyComputer ~ $ pip install navigate-micro
```

To install the bleeding edge version of **navigate**, run the following command:

```
(navigate) MyComputer ~ $ pip install git+https://github.com/TheDeanLab/navigate.git
```

**Run navigate software**

```
(navigate) MyComputer ~ $ navigate
```

**Note:** If you are running the software on a computer that is not connected to microscope hardware, you can add the flag `-sh` (`--synthetic-hardware`) to launch the program:

```
navigate -sh
```

#### 2.3 Launching navigate

Open an `Anaconda Prompt (Miniconda3)` and enter the following.

```
(base) conda activate navigate
(navigate) navigate
```

**Note:** If you are running Windows, you can create a desktop shortcut to **navigate** by right-clicking the Desktop, navigating to New and then Shortcut and entering `%windir%\system32\cmd.exe "/c" C:\path\to\miniconda\Scripts\activate.bat navigate && navigate` into the location text box.

#### 2.4 Developer install

**Download Git**

If you do not have Git already installed, you will need to do so before downloading the repo. We also recommend installing GitHub Desktop for a more user-friendly experience.

**Create a directory where the repository will be cloned**

We recommend a path/location that is easy to find and access such as the your Desktop or Documents. Once the folder is created, we will want to change that to our working directory (e.g., `cd`)

- Windows

```
(navigate) C:\Users\Username> cd Desktop
(navigate) C:\Users\Username\Desktop> mkdir Code
(navigate) C:\users\Username\Desktop> cd Code
```

- Linux/Mac

```
(navigate) MyComputer ~ $ mkdir ~/Desktop/Code
(navigate) MyComputer ~ $ cd ~/Desktop/Code
```

**Clone the GitHub repository**

```
(navigate) C:\Users\Username\Code> $ git clone https://github.com/TheDeanLab/navigate.git
```

**Install the Navigate repository**

The last step requires you to change into the navigate directory and the install the repo as an editable package locally on your machine.

```
(navigate) C:\Users\Username\Code> cd navigate
(navigate) C:\Users\Username\Code\navigate> pip install -e .[dev]
```

**Note:** If working in a `zsh` shell, e.g. on a modern MacOS, add single quotes around the call: `pip install -e '.[dev]'`.

#### 2.5 Troubleshooting

If the software is run at an institution with a proxy, you may need to update your proxy settings to allow `pip` and `conda` to install the proper packages.

- This can be done by going to Environment Variables for Windows, or another OS equivalent.
- **Create the following new System Variables (please see that**

    they are both http, this is purposeful and not a typo):

    - Variable = HTTP_PROXY; Value = http://proxy.your_university.edu:1234
    - Variable = HTTPS_PROXY; Value = http://proxy.your_university.edu:1234
- If you continue to have issues then change the value of Variable HTTPS_PROXY to https://proxy.your_university.edu:1234
- If you still have issues then you will need to create/update both configuration files for conda and pip to include proxy settings, if they are not in the paths below you will need to create them. This assumes a Windows perspective. Mac/Linux users will have different paths, they can be found online.
    - The `conda` configuration file can be found at C:\Users\UserProfile\.condarc
    - The `pip` configuration file can be found at C:\Users\UserProfile\pip\pip.ini
- You can also try to set the proxy from within the Anaconda Prompt:
- `set https_proxy=http://username:password@proxy.example.com:8080`
- `set http_proxy=http://username:password@proxy.example.com:8080`

CHAPTER

THREE

### I WANT TO. . .

This section contains how-to documents organized into three levels of difficulty: beginner, intermediate, and advanced.

- The beginner how-to is intended for users who have little to no experience with computer programming and simply wish to acquire an image.
- The intermediate how-to is intended for users who want to learn how to use the graphical user interface to create their own smart acquisition workflows.
- The advanced how-to is intended for users who have experience with computer programming and wish to extend the functionality of **navigate** by adding new devices or writing their own acquisition features.

Through this tiered approach, we hope to provide a gentle introduction to the software while also reaching the maximum number of users.

#### 3.1 Acquire an Image (Beginner)

This guide will describe how to acquire a single image and a z-stack using the **navigate** software package.

##### 3.1.1 Launching the Software Package

###### Open Anaconda Prompt

To start, you need to open the Anaconda Prompt. Follow these steps:

1. On Windows, click on the Start menu.
2. Type `Anaconda Prompt` into the search bar.
3. Click on the Anaconda Prompt application to open it.

---

**Note:** Ensure that Anaconda and **navigate** are already installed on your system. If not, please refer to our *Quick Start Guide* for more information.

---

##### Activate Conda Environment

Once the Anaconda Prompt is open, activate the desired conda environment. By default, the command prompt will open the base environment (as shown in parentheses). To activate **navigate** environment, type the following command into the Anaconda command window and press `Enter`

```
(base) conda activate navigate
```

##### Launch the Software Package

After activating the environment, **navigate** should now be shown in parentheses. After you have already *configured* **navigate**, you can launch it by typing the following command into the Anaconda command window:

```
(navigate) navigate
```

The **navigate** software package will launch and the main window will appear.

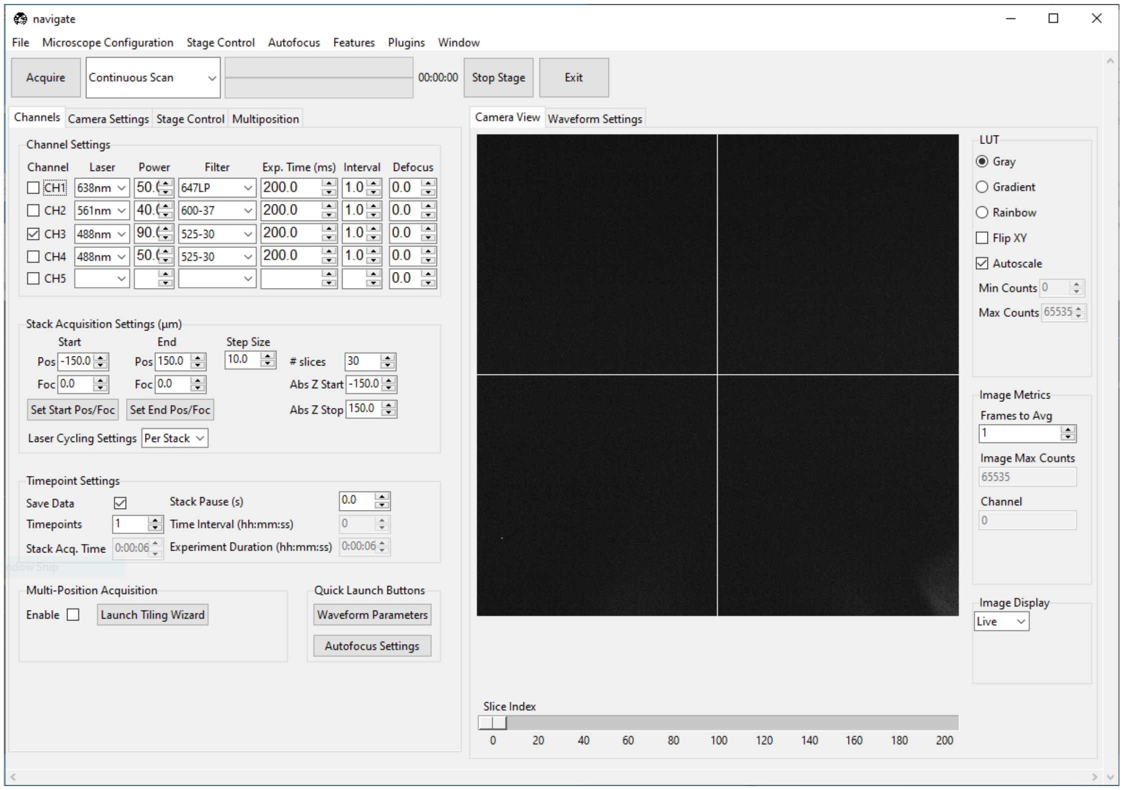

##### 3.1.2 Configure the Channel Settings

- Select the *Channels* tab, which is located on the upper left of the main window.
- Under the *Channel Settings* section, select the number of channels needed for imaging. For each channel selected, you will need to configure the acquisition settings:

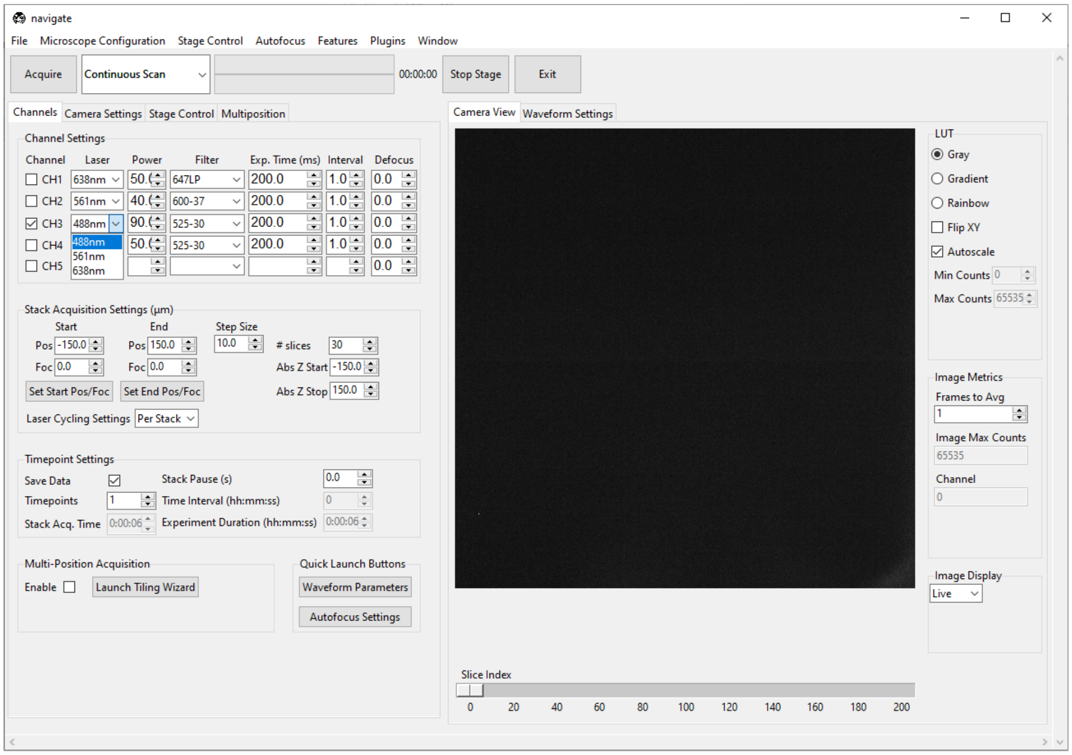

- Select the appropriate *Laser* from the dropdown menu.
- Select the appropriate *Power* for the laser.
- Select the appropriate emission *Filter* from the dropdown menu.

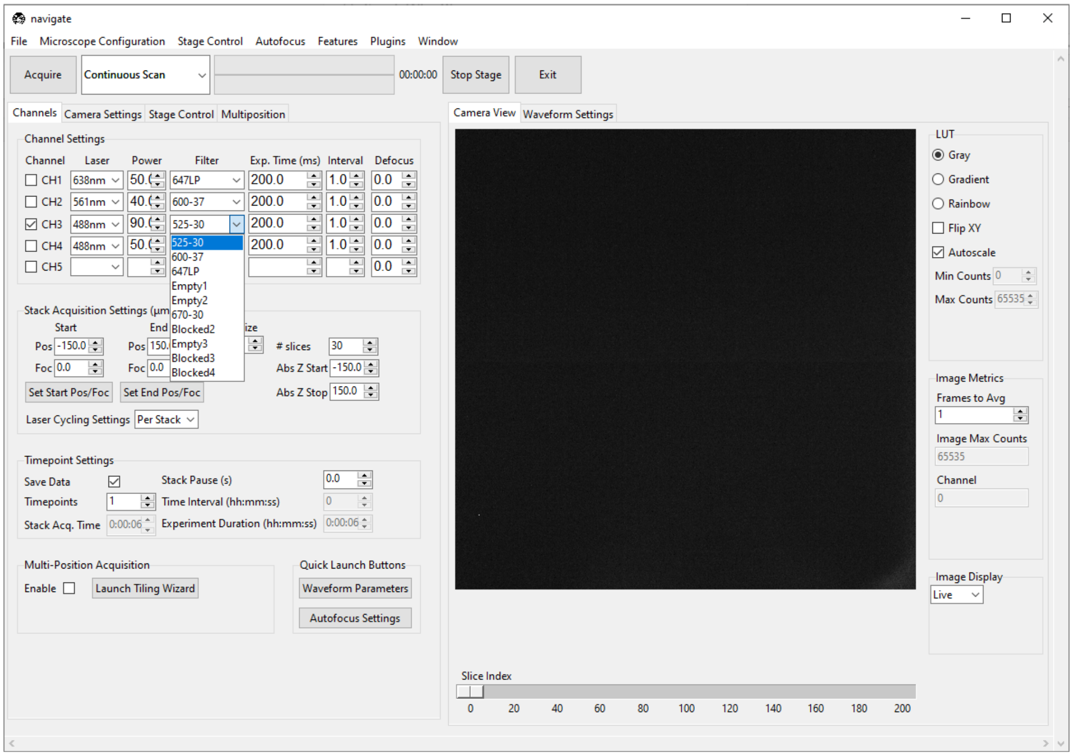

- Specify the camera *Exp. Time (ms)*. A good default value is `100` or `200` ms.
- Specify the *Interval* to be `1.0`. While this feature is not currently implemented, future releases will allow users to image different channels at different time intervals.
- Specify the *Defocus* to be `0`. This feature allows you to adjust for chromatic aberrations that result in focal shifts between each imaging channel.

---

##### 3.1.3 Configure the Camera Settings

- Select the *Camera Settings* tab.
- For standard imaging applications, select *Normal* in the *Sensor Modes* dropdown menu within the *Camera Modes* section.
- If you are using the rolling shutter, select *Light-Sheet* and specify its *Readout Direction* and *Number of Pixels*.

**Note:** For more information on how to configure the rolling shutter for ASLM operation, please refer to *ASLM*.

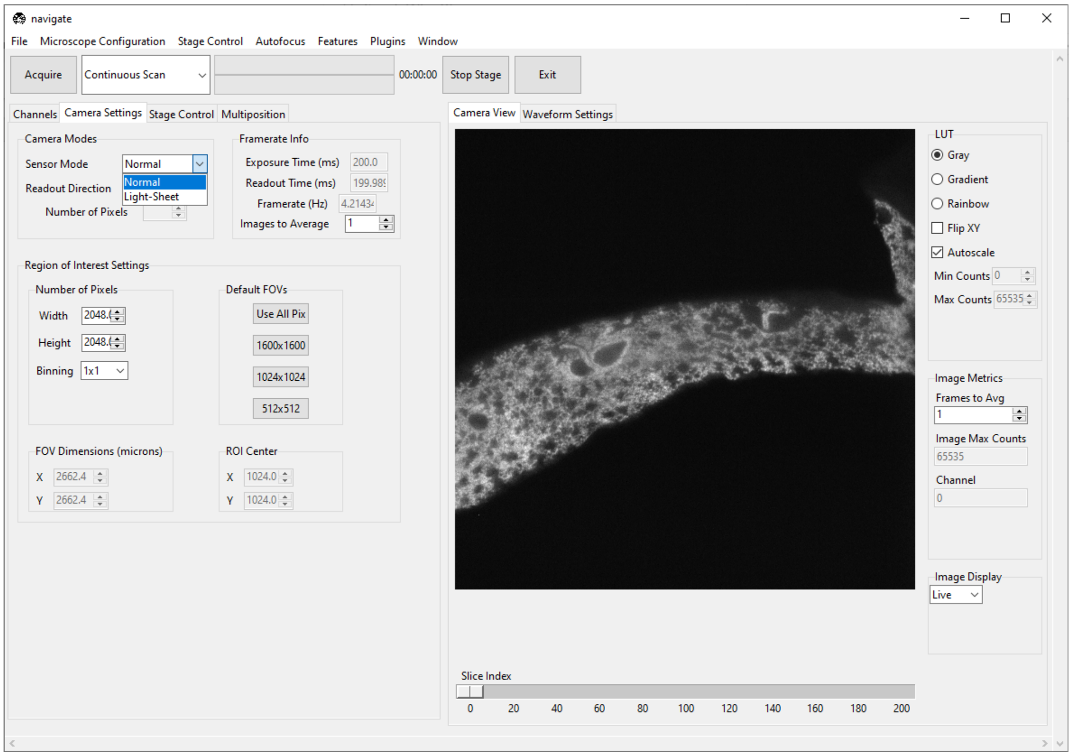

- **Choose the size of your camera's field of view.**
  - Specify the *Region of Interest Settings* by entering the appropriate *Number of Pixels* for both the *Width* and *Height* values. Alternatively, one can select from one of several default values in the *Default FOVs* section.

---

**Note:** The *FOV Dimensions (microns)* is automatically calculated based on the *Number of Pixels* and the *pixel_size* as specified in the *zoom* section of your your `configuration.yaml` file.

```yaml
zoom:
  pixel_size:
    20x: 0.325 # magnification, and pixel size in microns
```

---

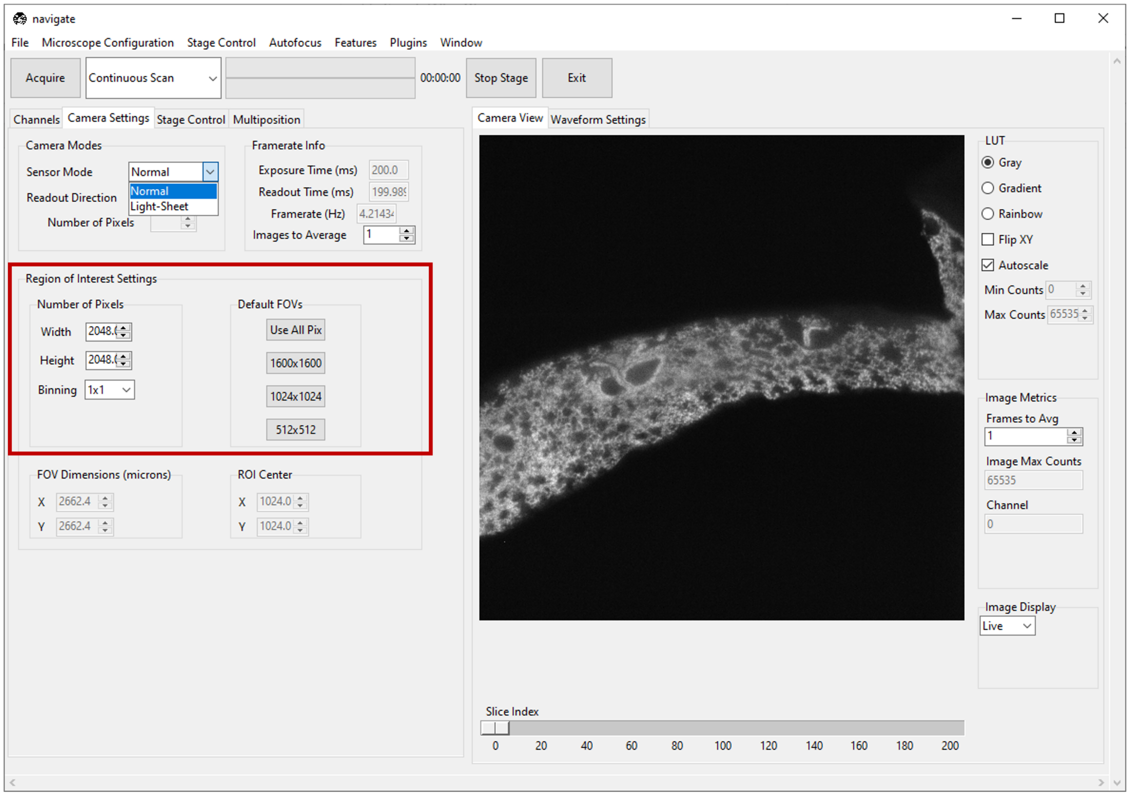

**Note:** If multiple channels are selected, each channel will be acquired with the same camera *Sensor Mode*, *Readout Direction*, and *Region of Interest Settings*.

##### 3.1.4 Acquire in a Continuous Scan Mode

- Select “Continuous Scan” in the dropdown next to the *Acquire* button in the *acquire bar*.

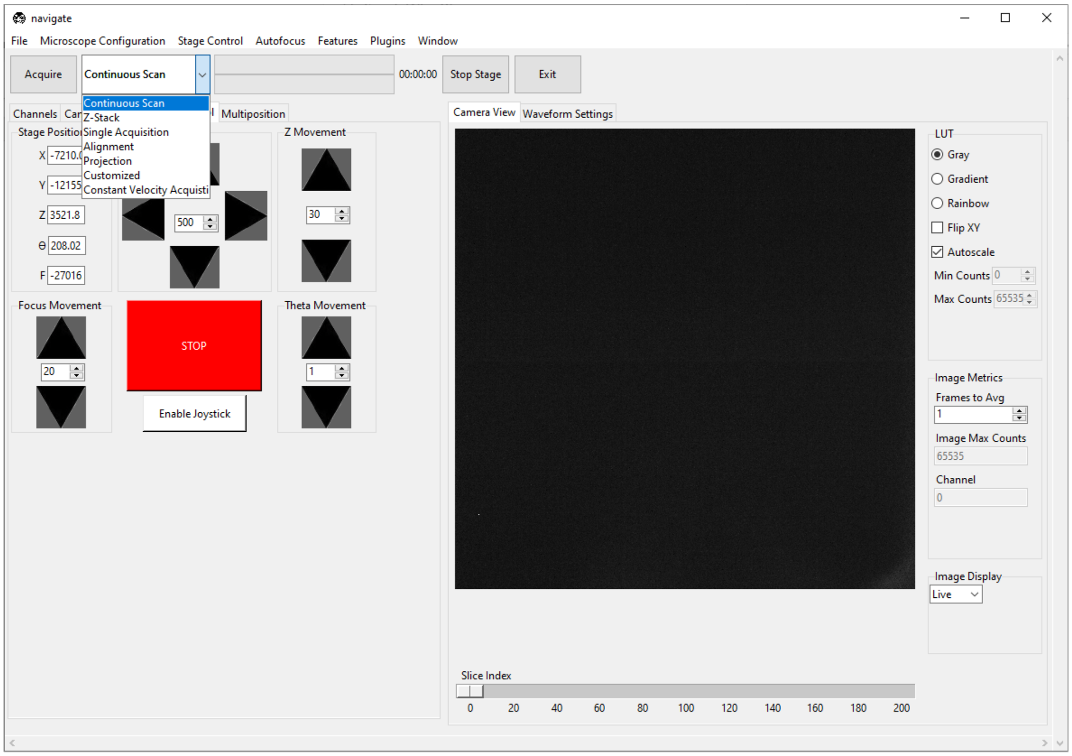

- Press *Acquire*. This will launch a live acquisition mode.

---

**Note:** If multiple channels are selected, each channel will be imaged sequentially. The order of imaging is determined by the order of the channels in the *Channel Settings* section of the *Channels* tab, and will proceed from the top to the bottom of this channel list.

---

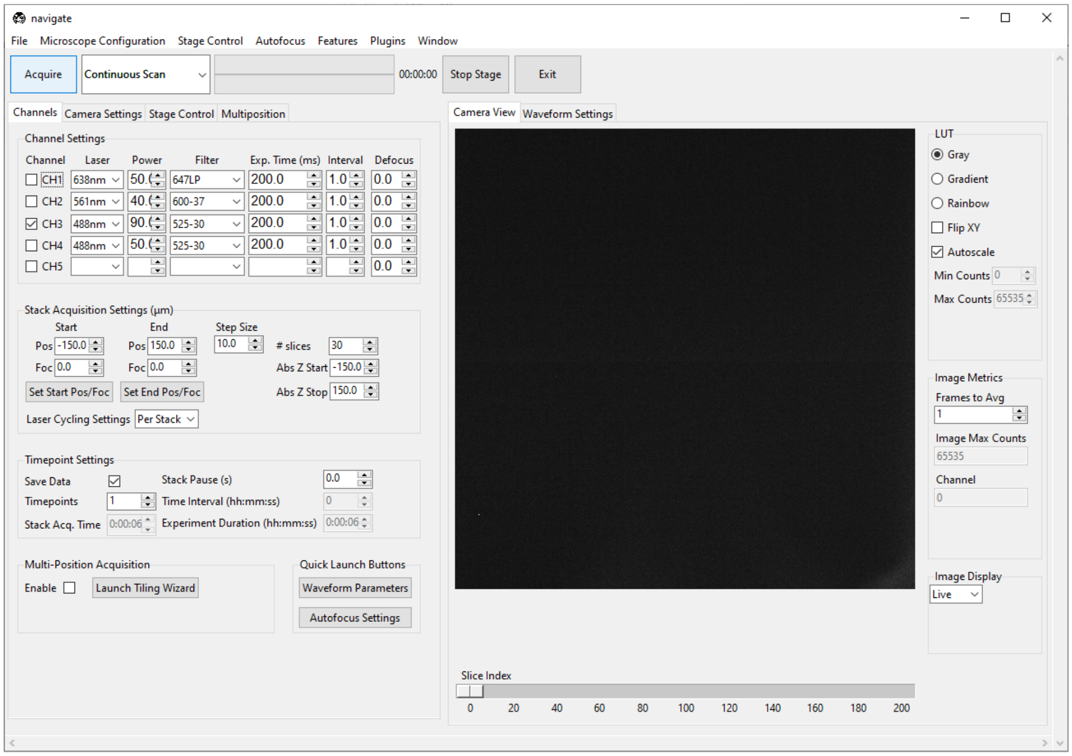

- **Move the stage to identify the location of the sample.**
  - Select the *Stage Control* tab, and use the graphical user interface to move the stage. This includes buttons for moving the stage in `X`, `Y`, `Z` , `F`, and `Theta` directions. * The step size for each axis can be adjusted with the spinbox next to each button. * For stages loaded in a synthetic mode, buttons will be disabled. * Absolute positions can be entered in the text boxes next to each button. * Check *configuration settings* for more information.
  - Alternatively, if available, use the manufacturer-provided joystick to position the sample.

**Note:** The axes for a light-sheet microscope vary in the literature. Here, we define the `Y` axis as the direction of the light-sheet propagation, the `Z` axis as the direction of the detection objective, and the `X` axis as the direction perpendicular to the light-sheet and detection objective axes.

The `F` axis typically controls the position of the detection objective along the detection axis.

The `Theta` axis typically controls the rotation of the sample.

**Warning:** One should always be careful when moving the stage.

If the stage is moved too quickly, the sample and/or microscope may be damaged.

We strongly recommend that you implement stage limits in your configuration file. Please refer to the *configuration settings* for more information.

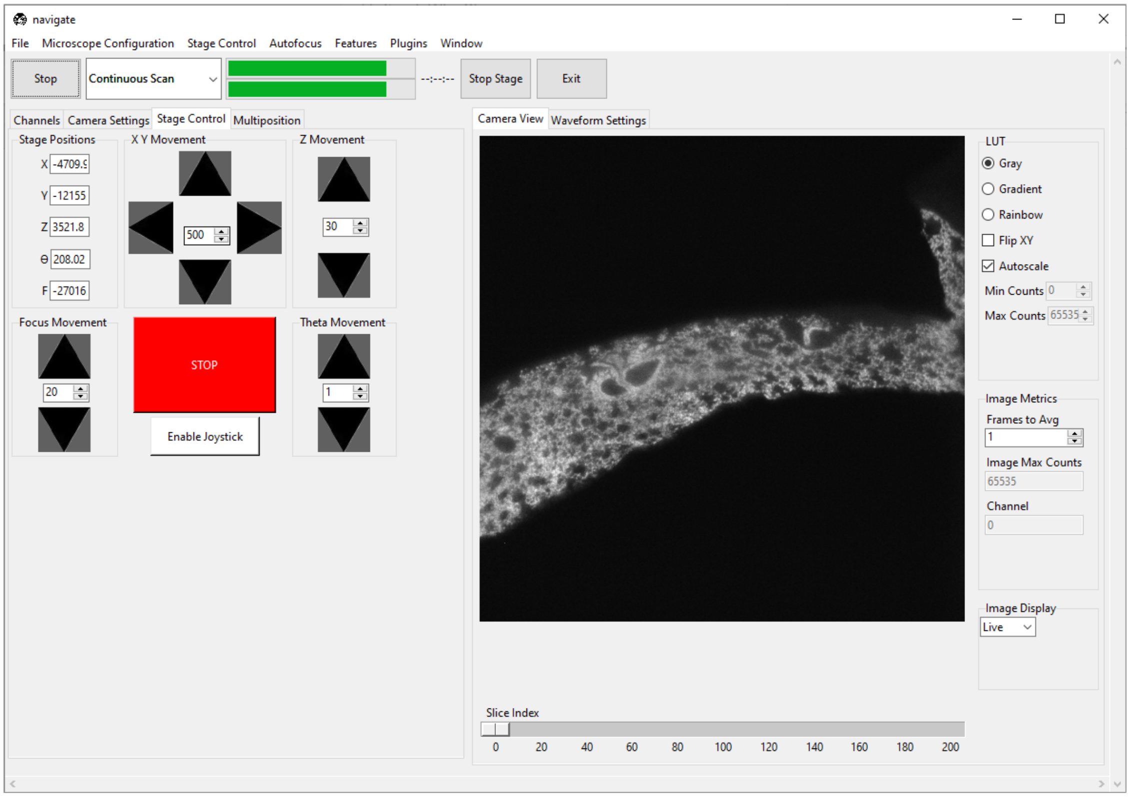

- Press the *Stop* button in the acquisition bar to stop acquisition.

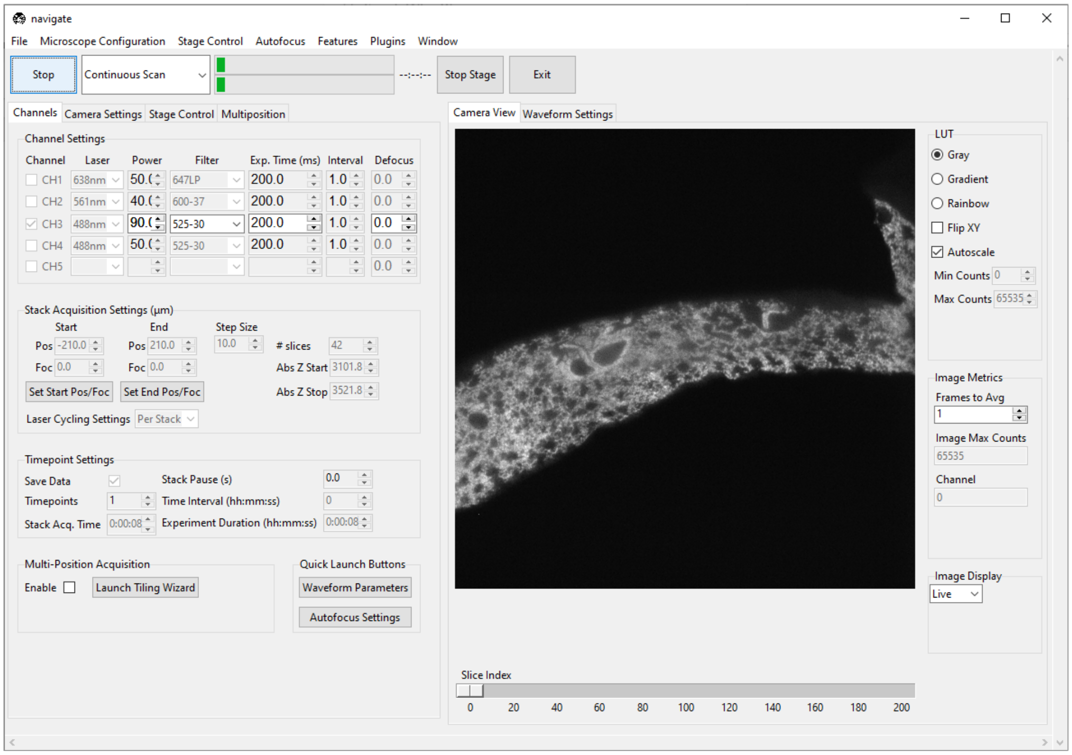

##### 3.1.5 Acquiring a Single Image

- Check the *Save Data* box in the *Timepoint Settings* section under the *Channels* tab to save the acquired images. Check this box before acquiring data.

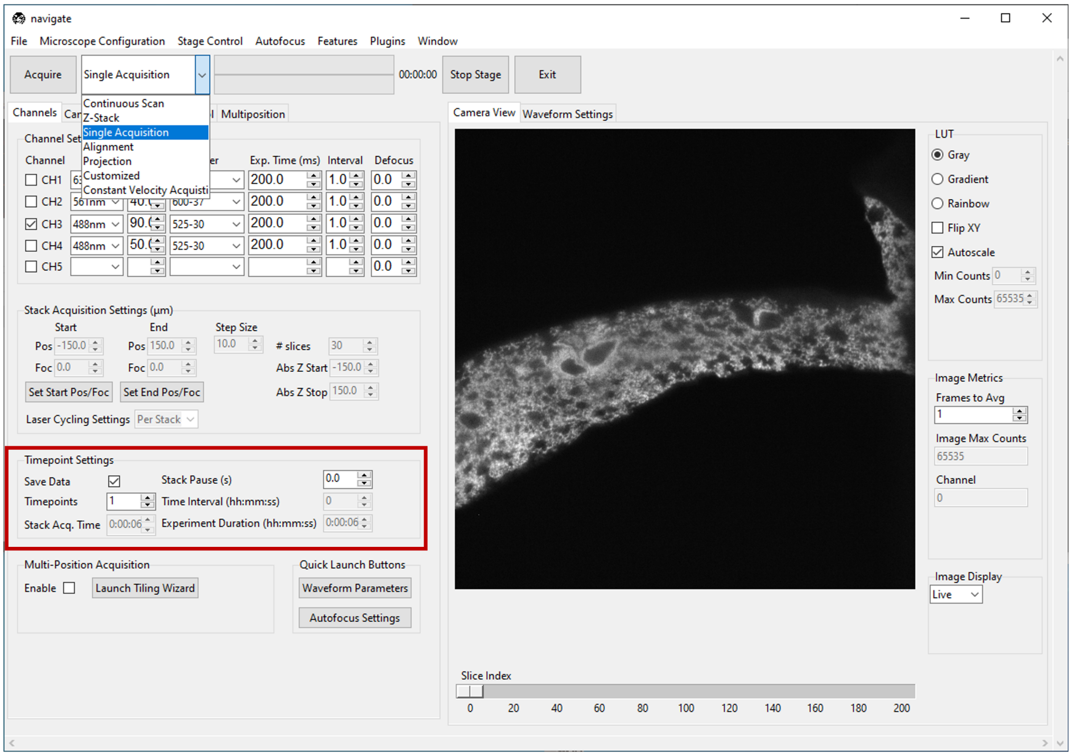

- Select *Single Acquisition* from the dropdown next to the *Acquire* button.

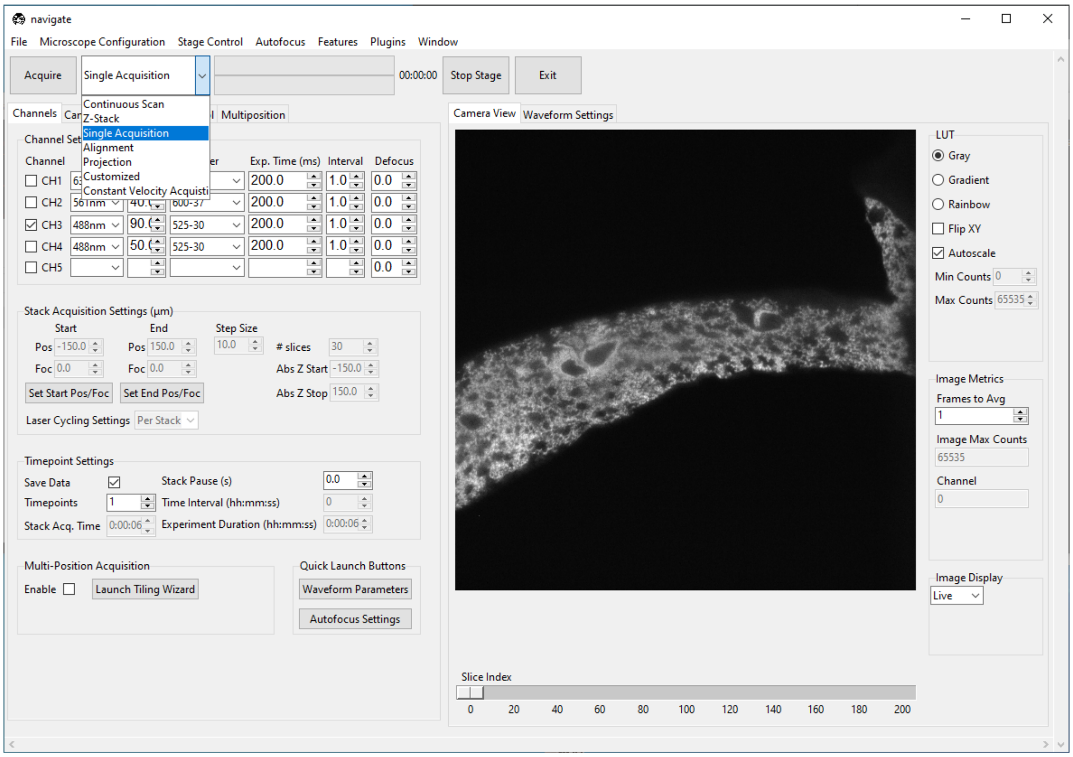

- Press *Acquire* to open the *File Saving Dialog* interface. Enter the sample parameters, notes, location to save file, and filetype in the *File Saving Dialog* that pops up.

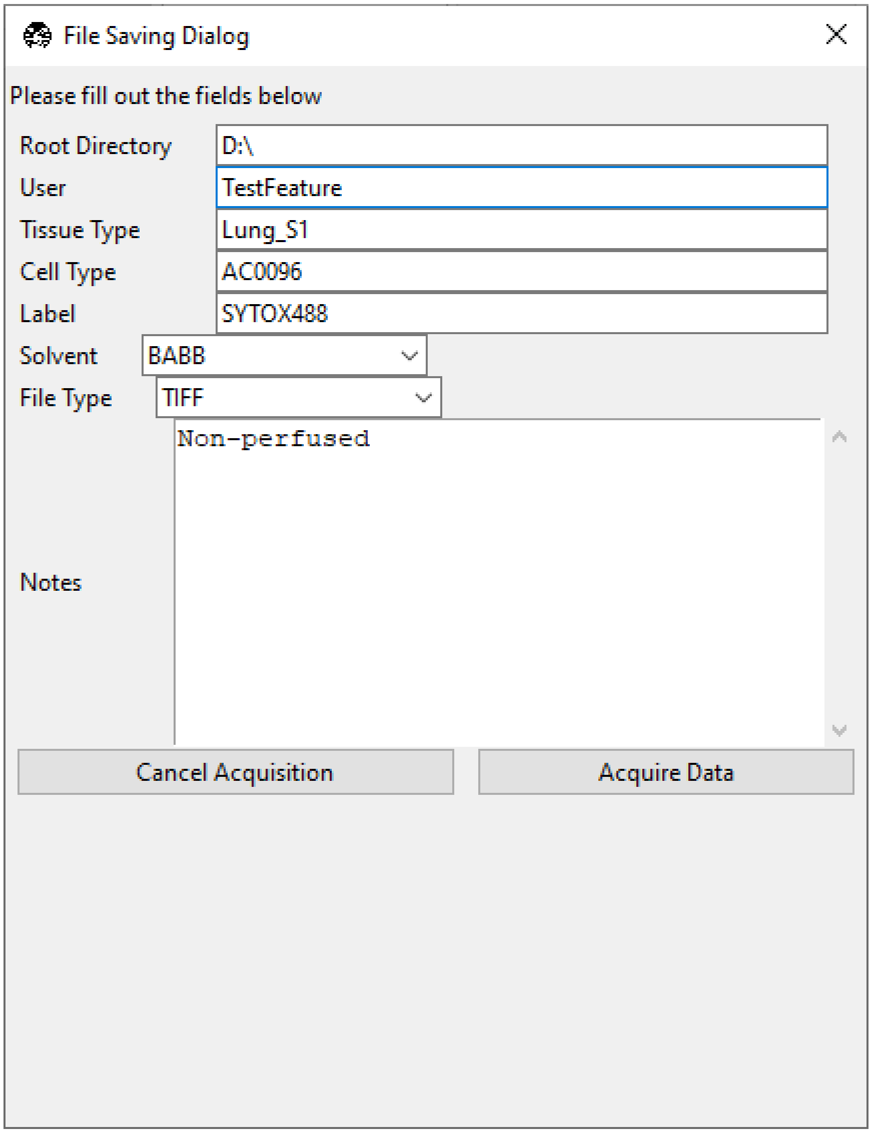

- Press *Acquire Data* to initiate acquisition. Acquisition will automatically stop once the image is acquired.

**Note:** Each acquisition will be saved in a separate folder (e.g., `Cell01`, `Cell02`, …) within the directory specified in the *File Saving Dialog* interface.

Data will not be overwritten between acquisitions.

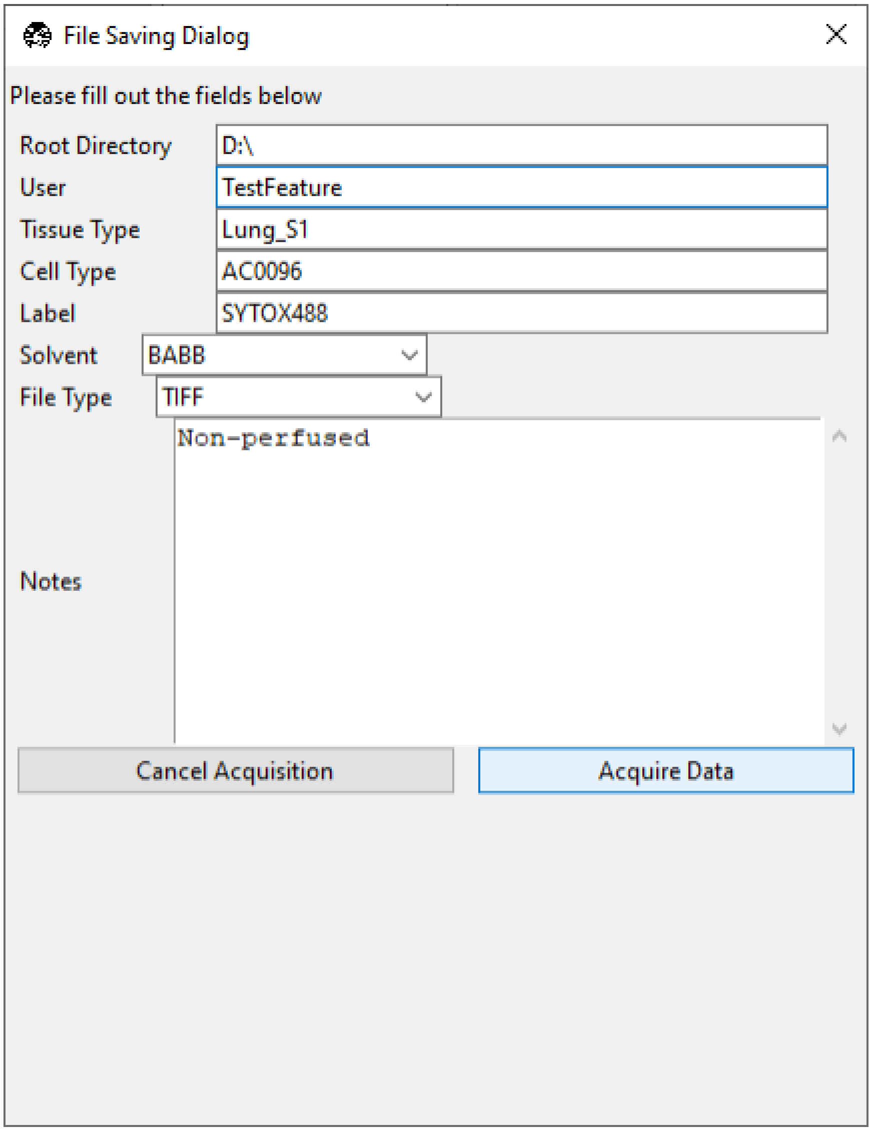

##### 3.1.6 Acquiring a Z-Stack

- Using the *Stage Control*, go to the desired z-position in the sample. Make sure that the sample is in focus. To use the autofocus feature, please refer to the *Autofocus Settings*.

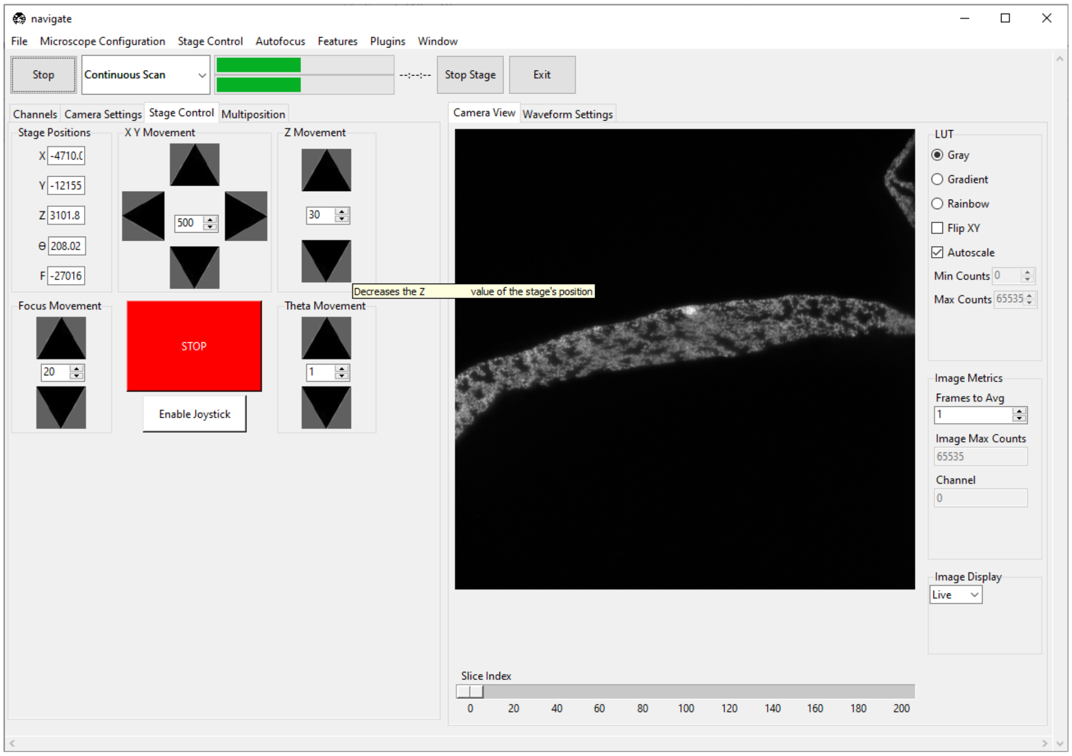

- Under the *Channels* tab, in *Stack Acquisition Settings (μm)* press *Set Start Pos*.

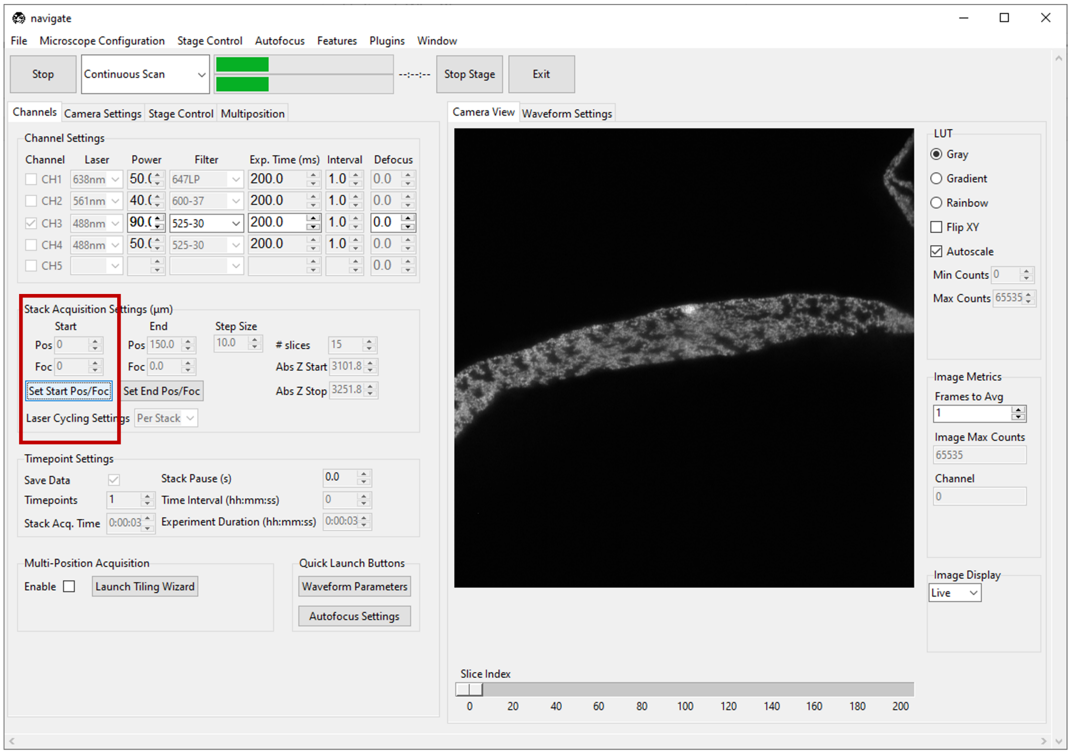

- Using the *Stage Control*, go to a different z-position within the sample. Again, make sure that the sample is in focus.

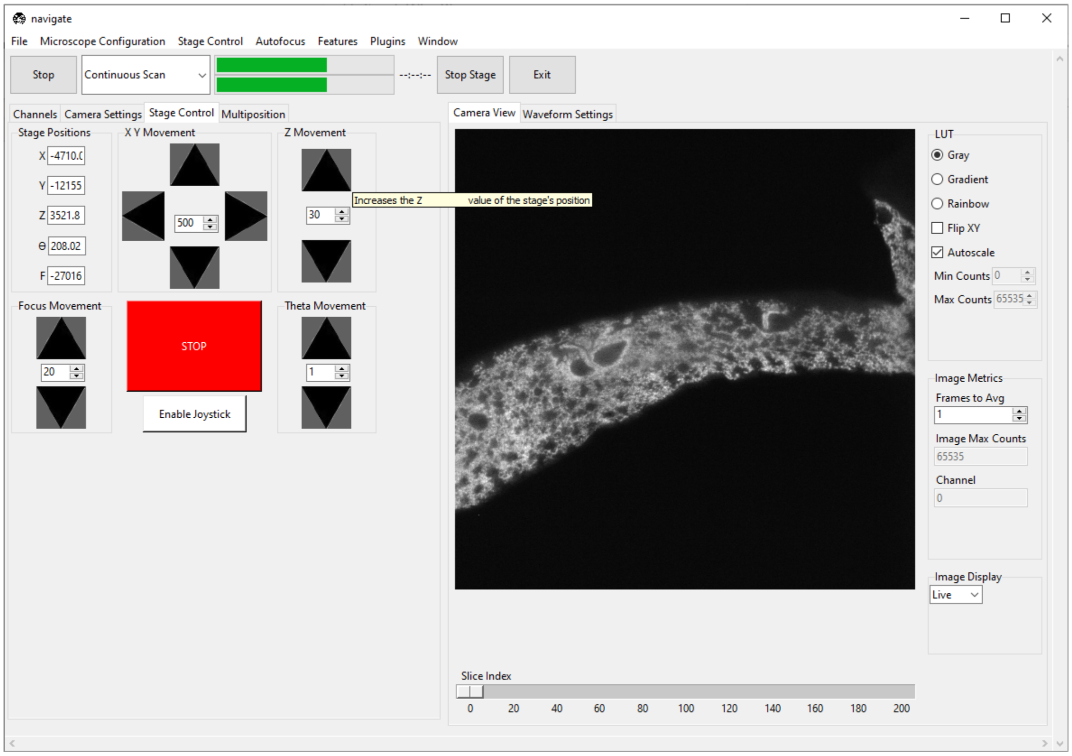

- Under the *Channels* tab, in *Stack Acquisition Settings (µm)* press *Set End Pos*.

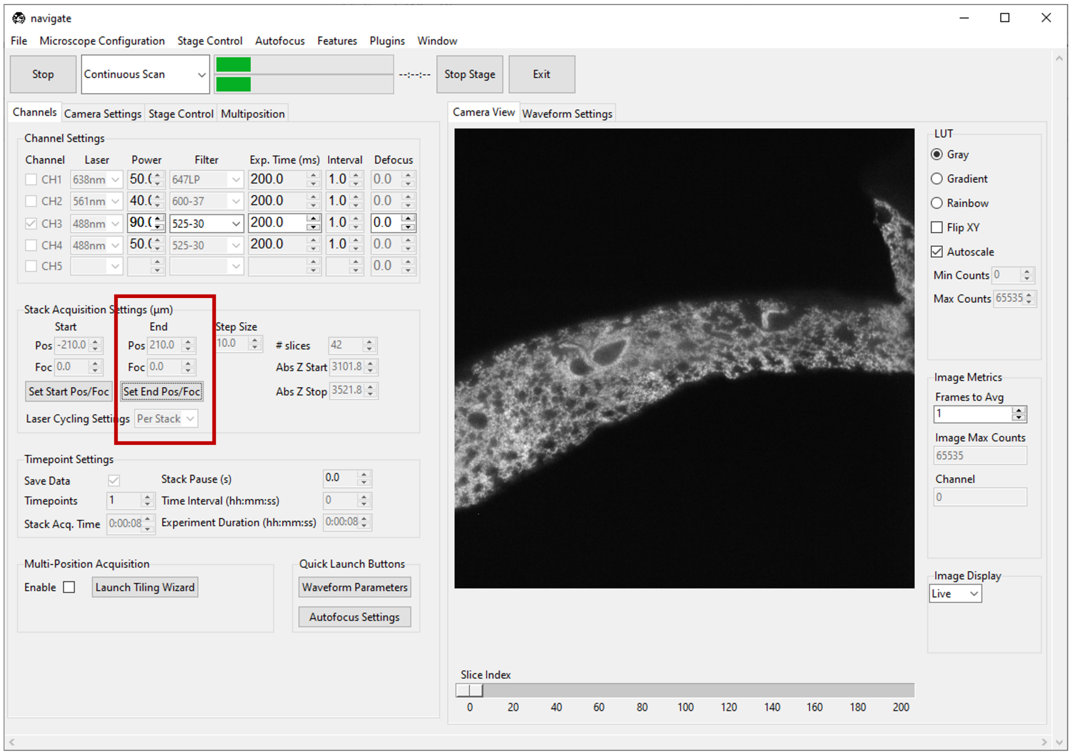

**Note:** If there is a shift in F between the start and stop positions, the F axis will be ramped synchronously with Z to maintain focus.

Check *configuration settings* for more information to determine if focus is enabled in hardware.

Refer to *Imaging on a mesoSPIM BT* section for an example of how to acquire a z-stack with a focus ramp.

- Type the desired step size in microns in the *Step Size* dialog box in *Stack Acquisition Settings (µm)*.

**Note:** The minimum step size, and increment between steps, are graphical user interface defaults that are specified in the `configuration.yaml` file. More information can *configuration settings*

```yaml
gui:
  stack_acquisition:
    step_size:
      min: 0.100
      max: 1000
      step: 0.1
```

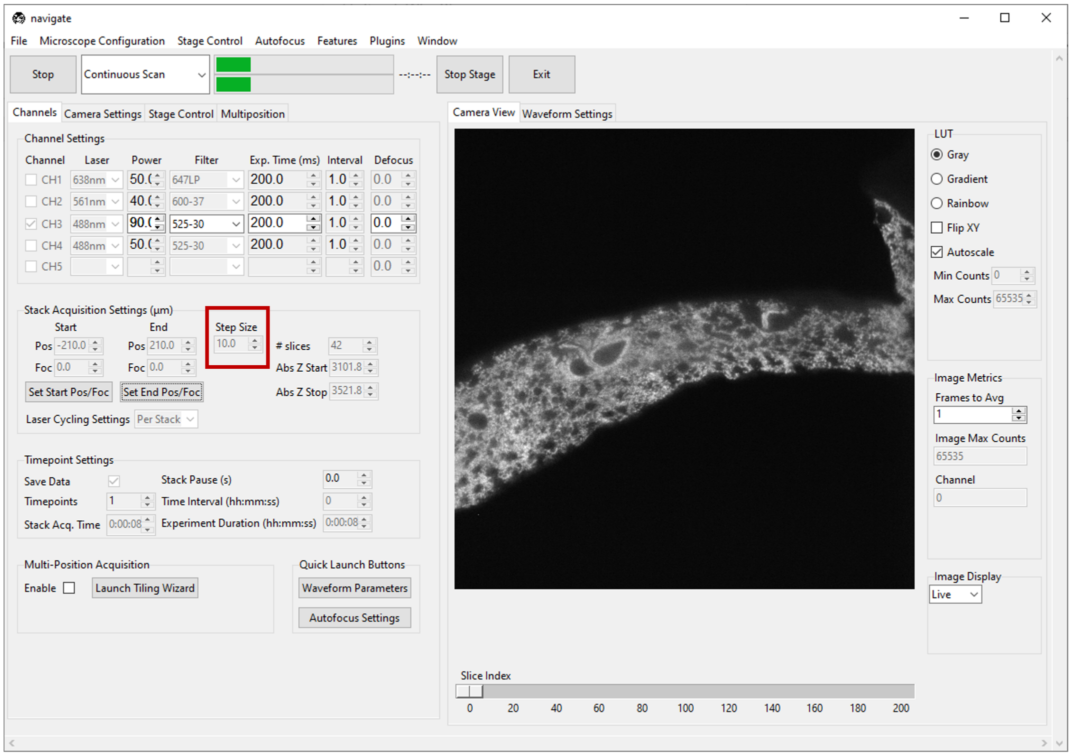

- If using multiple channels for imaging, select either *Per Z* or *Per Stack* under *Laser Cycling Settings* in the *Stack Acquisition Settings (µm)* section under the *Channels* tab.
  - *Per Z* acquires all channels before moving the stage to a new position.
  - *Per Stack* acquires all images in a stack acquisition for a single channel before moving the stage back to the start position and restarting acquisition for the subsequent channel until all channels are imaged.

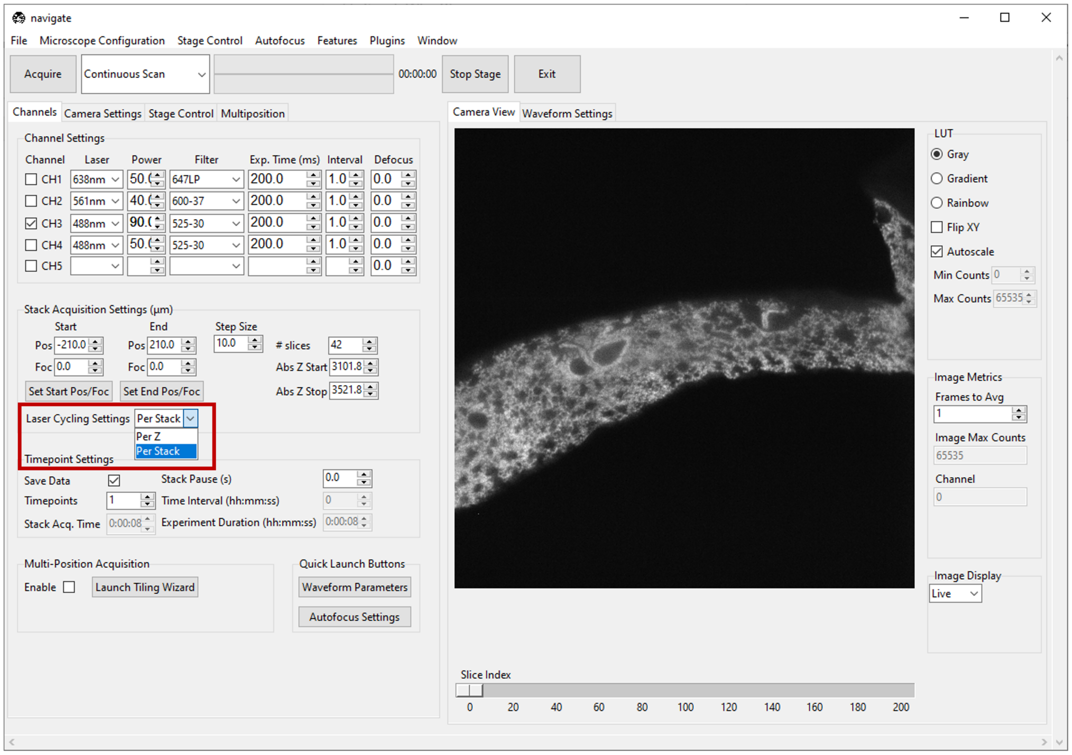

- Select *Z-Stack* from the dropdown next to the *Acquire* button. Press *Acquire*.

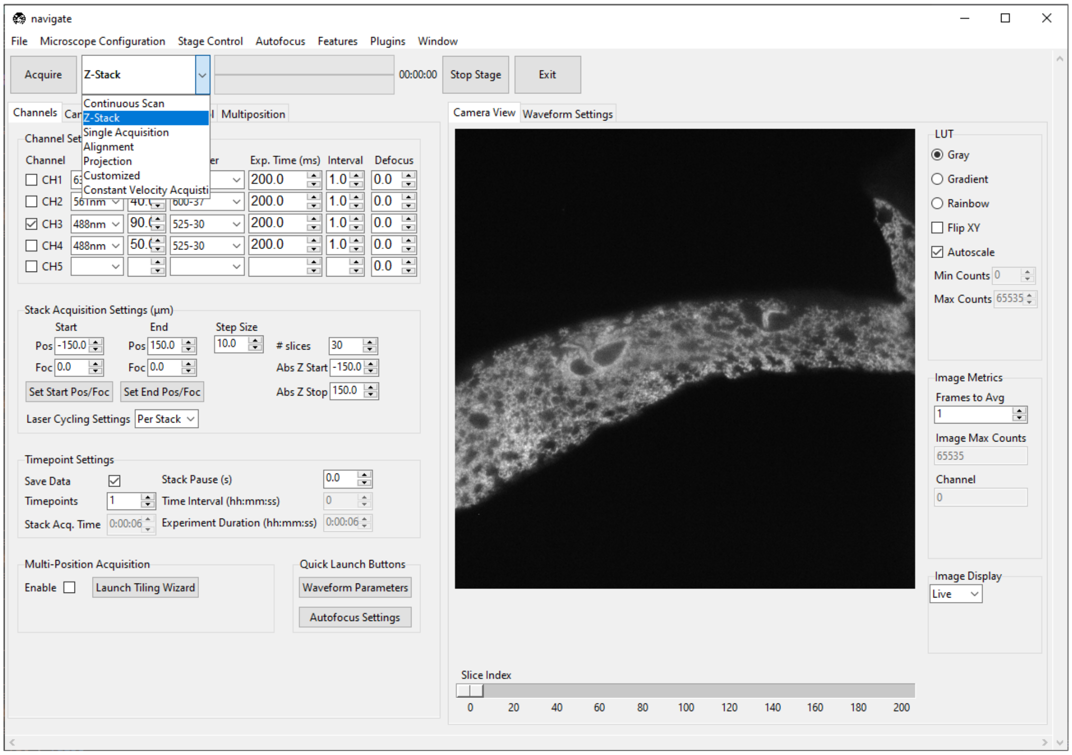

- Enter the sample parameters, notes, location to save file, and filetype in the *File Saving Dialog* that pops up.
- Press *Acquire Data* to initiate acquisition. Acquisition will automatically stop once the image series is acquired.

#### 3.2 Write A Smart Acquisition Routine (Intermediate)

**navigate**'s *feature container* enables us to write acquisition routines on the fly by chaining existing *features* into lists. Please see Currently Implemented Features ffor a complete list of features. Users can build additionals feature within *plugins*.

In this guide, we will use existing features to write a routine that scans through an imaging chamber and takes z-stacks only where it finds the sample.

Suppose there are two positions listed in the *multiposition table*, one containing tissue and one empty, as shown below.

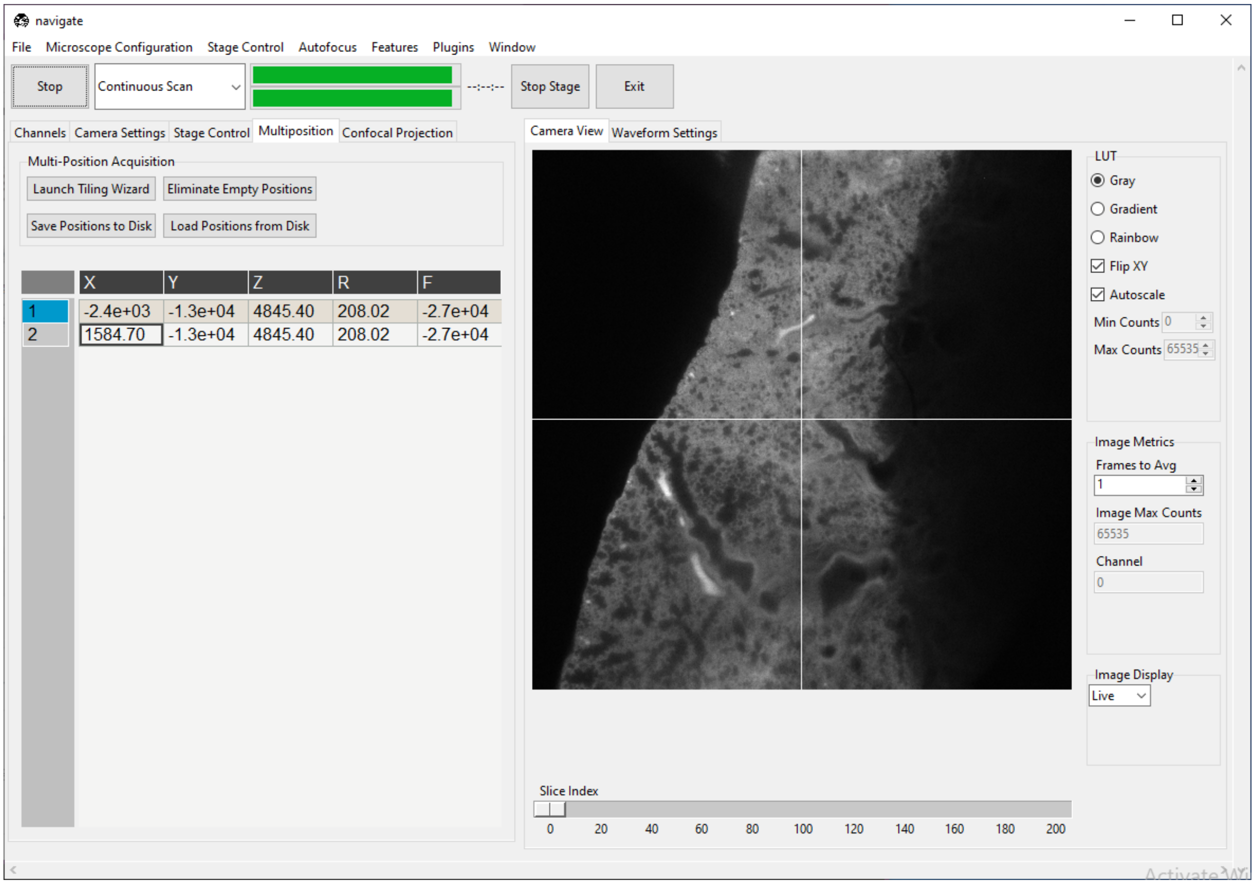

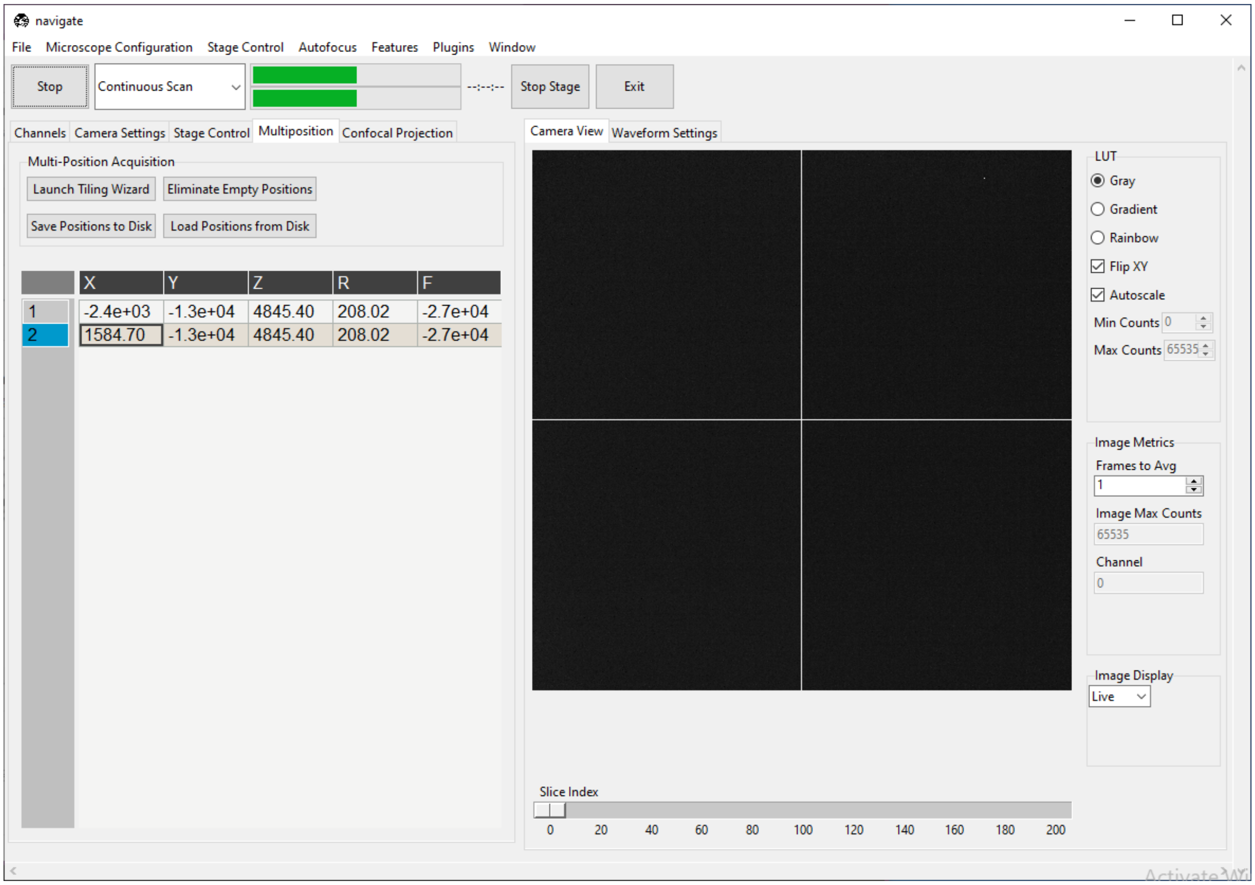

We will build a feature that scans both positions, but only takes a z-stack at the one containing tissue. To access the GUI feature list editor, navigate to *Features* → *Add Customized Feature List*. A window titled "Add New Feature List" will pop up. Enter `TestFeature` in the text box at the top of the popup. Enter

```
[{"name": PrepareNextChannel}]
```

in the text box at the bottom of this popup and press *Preview*. The window should appear as below.

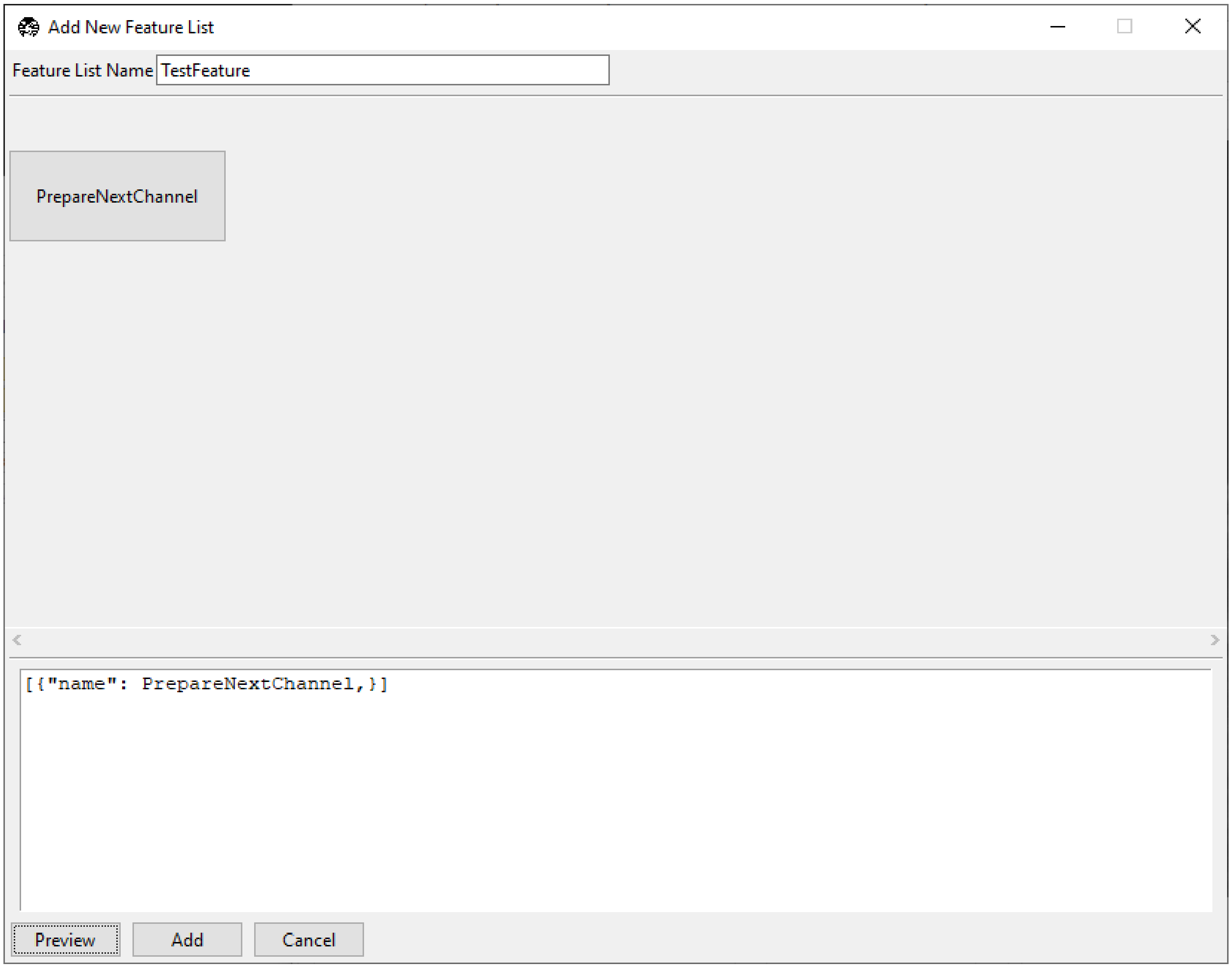

The square brackets `[]` create a sequence of events to run in the feature container. The `{}` braces contain features. In this case, we have a single feature, `PrepareNextChannel`, which will set up the next color channel for acquisition. The complete sequence (as it stands) will take an image in the first *selected color channel*.

We can build much of the rest of our desired acquisition in the GUI. Right-click on the `PrepareNextChannel` tile to reveal the editing menu.

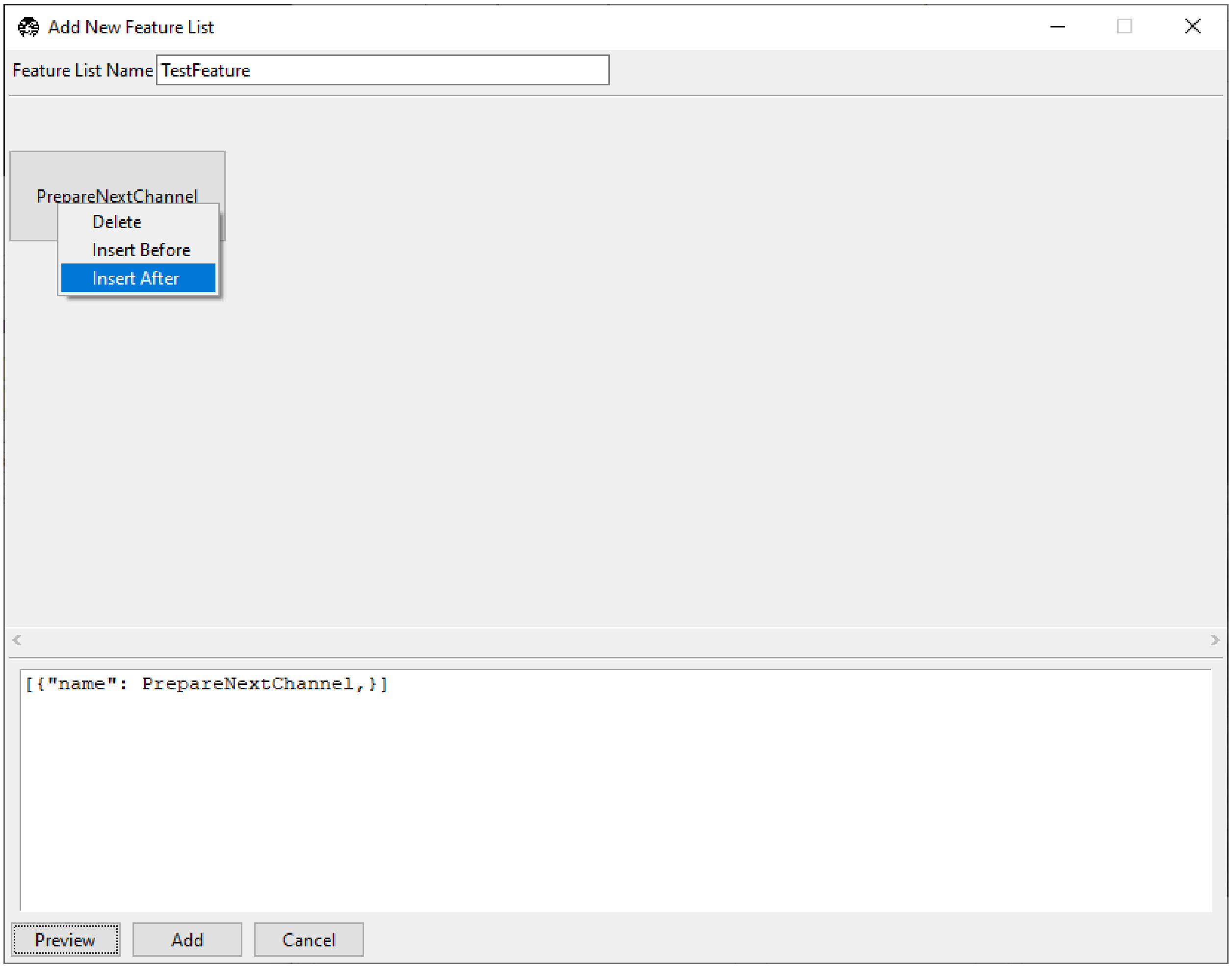

Select *Insert After*. A second copy of `PrepareNextChannel` will appear in the feature list.

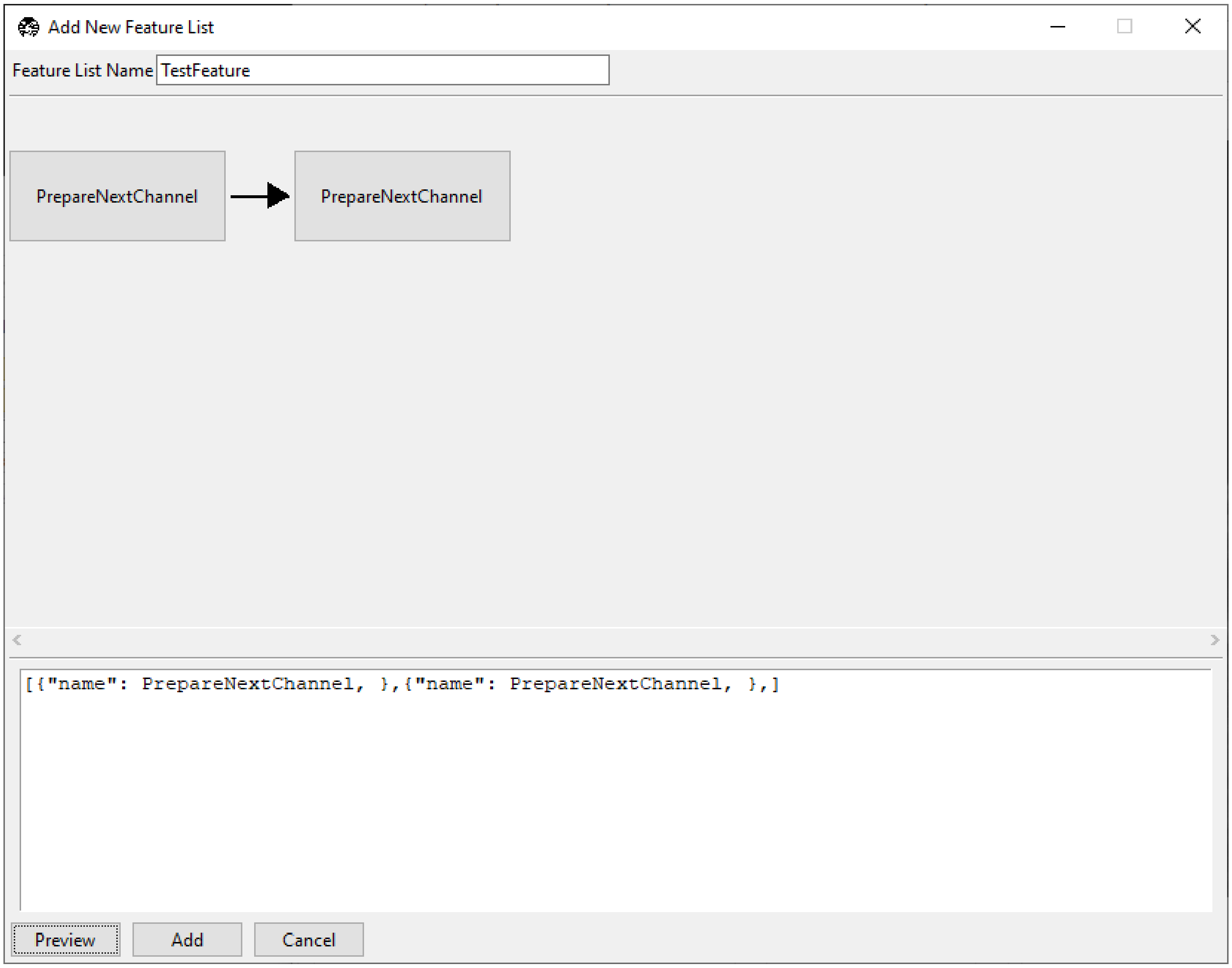

Left-click on this second tile to reveal a popup that allows us to select which feature we want to use in this tile.

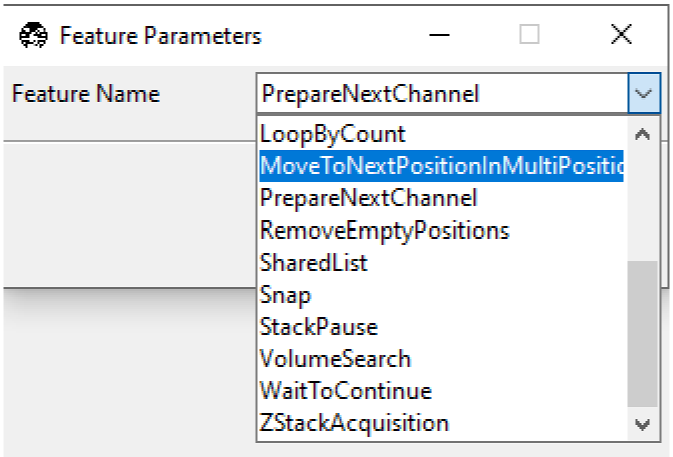

Select `MoveToNextPositionInMultiPositionTable` and then close the popup. Your feature list editing window should now appear as below.

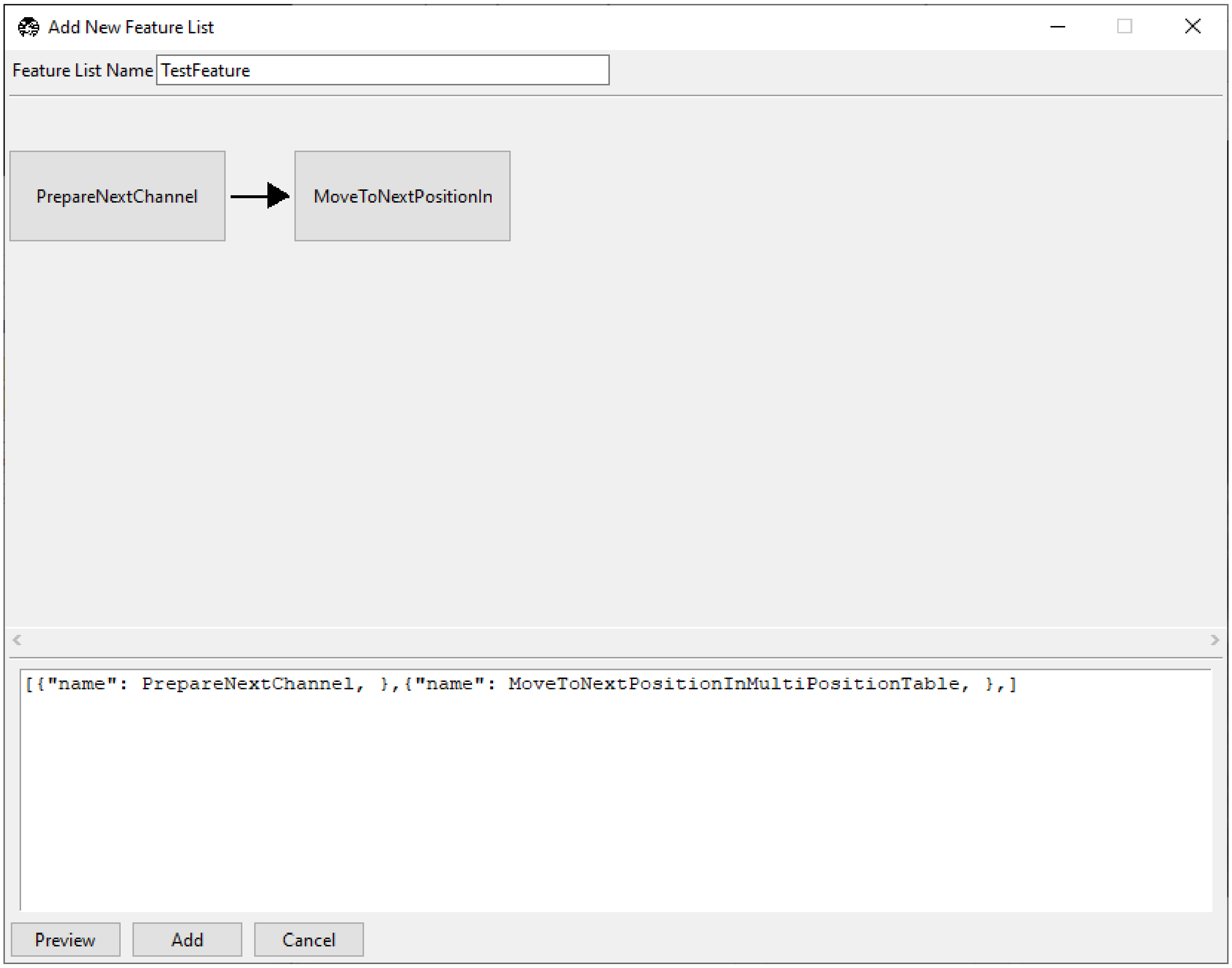

Our feature now takes one image of the first selected color channel at each position in the multiposition table.

Now, right-click `MoveToNextPositionInMultiPositionTable` and press *Insert After*. Click the new tile and change it to `DetectTissueInStackAndReturn`. There are three options associated with this feature.

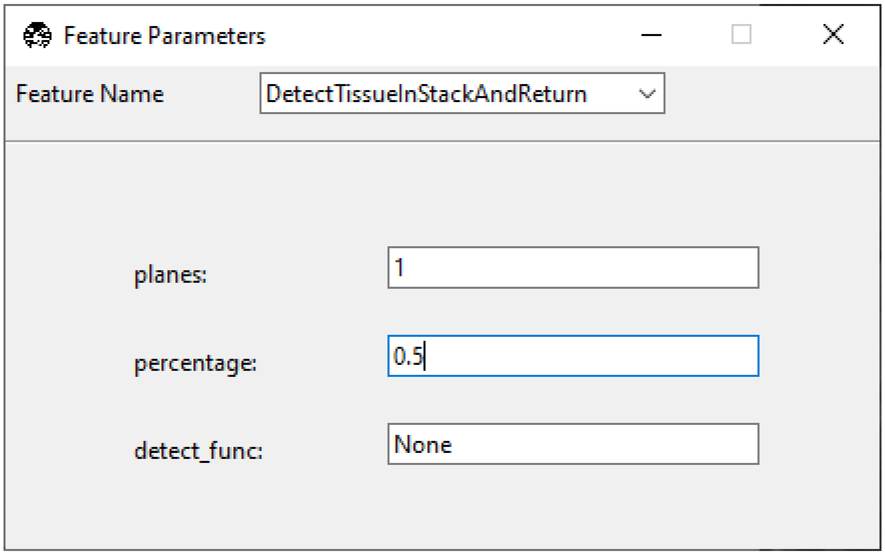

- *planes* indicates how many planes of the z-stack this feature should check for tissue.

- *percentage* indicates what percent of the total image must contain tissue for this feature to return `true` that tissue was detected. 1 indicates that the entire image contains tissue, 0.5 indicates that half of the image contains tissue, and so on.
- *detect_func* is one of the tissue detection functions in remove_empty_tiles. If this is set to `None`, it defaults to `detect_tissue()`, which states that tissue is present if signal is above the Otsu threshold of the stack of images acquired.

In this example, if any plane meets the desired threshold, the feature will return `true` and it will be acquired. If no plane meets the desired threshold, the feature will return `false`

Now, right-click `DetectTissueInStackAndReturn` and press *Insert After*. Click the new tile and change it to `LoopByCount`.

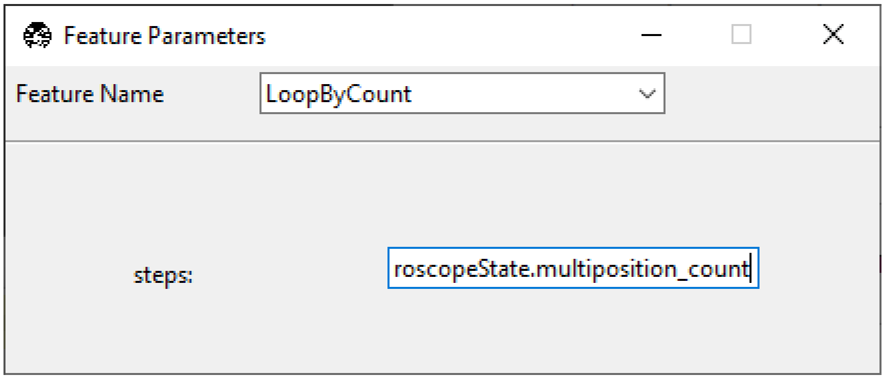

We want to iterate over all of the positions in the multi-position table, so we will set `steps` to `experiment.MicroscopeState.multiposition_count`.

Notice that the acquisition protocol does not appear to loop, but rather still moves in a sequence. This is because all of the tiles are still in the sequence brackets `[]`. We can now enclose the section of the protocol we want to loop in parentheses `()` and press *Preview* to see the update.

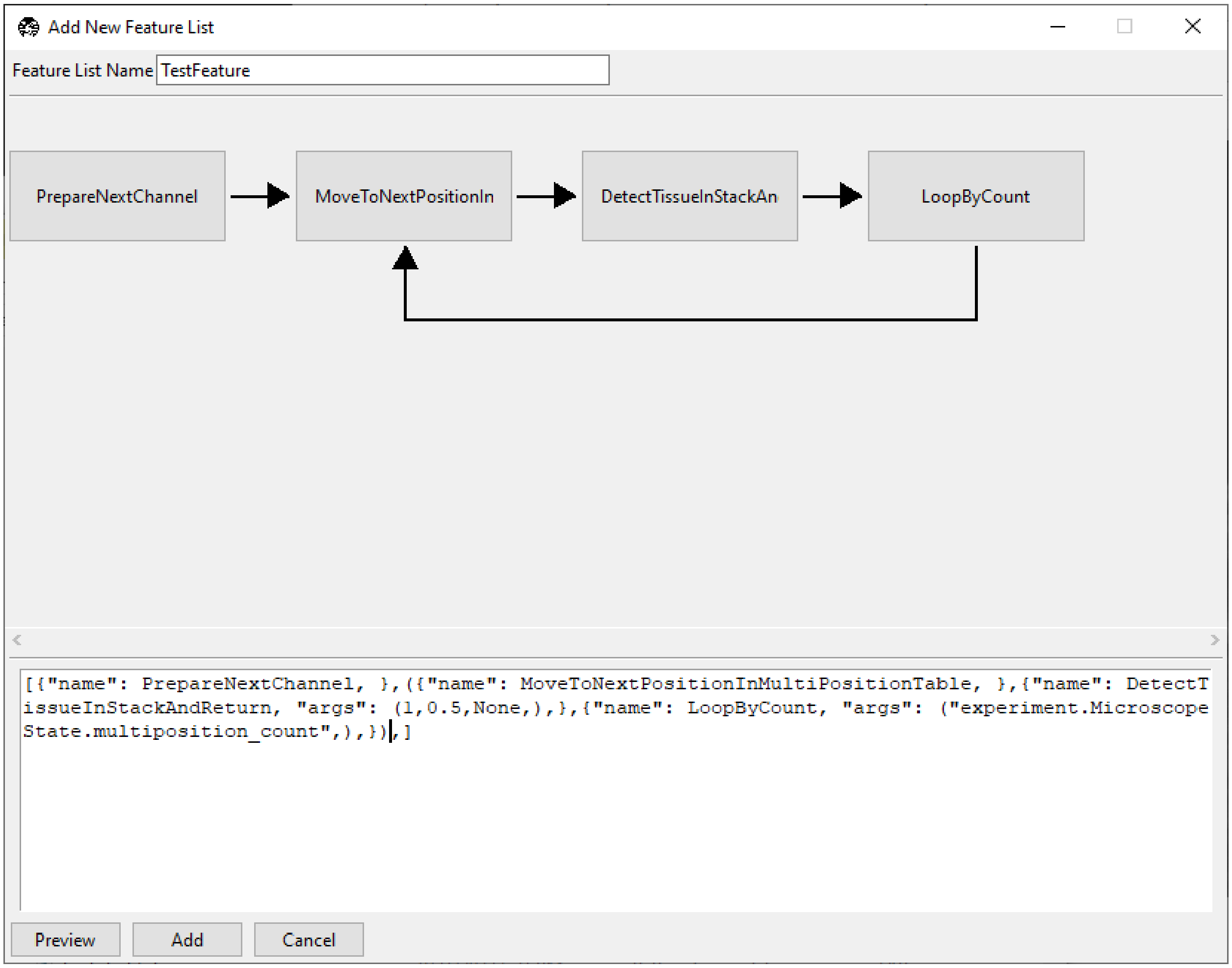

Now, we set up one color channel to image (`PrepareNextChannel`), and then within this channel visit every position in the multiposition table, and detect if there is tissue. However, we do no yet make any decisions of what to do if tissue is found. To do this, we will convert `DetectTissueInStackAndReturn` into a decision node.

To do this, we add `true` and `false` options within the feature braces:

```
{"name": DetectTissueInStackAndReturn,
 "args": (1, 0.5, None),
 "true": [{"name": ZStackAcquisition,"args": (False,False,"z-stack",),}],
 "false": "continue",}
```

Our `true` argument tells the software what to do if tissue is detected. In this case, we take a z-stack at the positions where tissue is found. The `false` argument tells the software how to proceed if no tissue is found. In this case, the `continue` option tells the software to keep moving through the loop to the next position in the multi-position table. Press *Preview* to see the update.

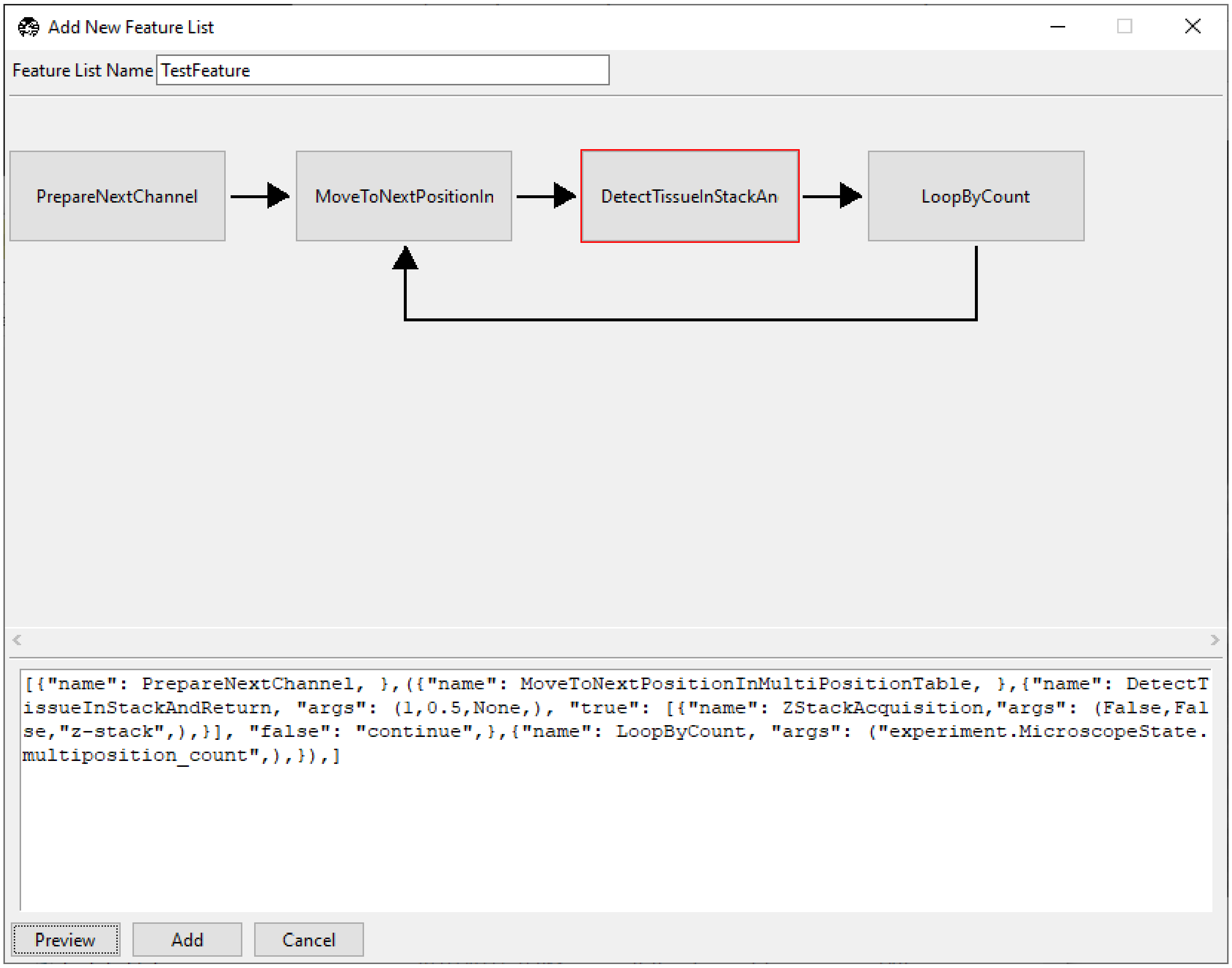

`DetectTissueInStackAndReturn` now has a red border, indicating it is a decision node. Click on it to access the decision node GUI.

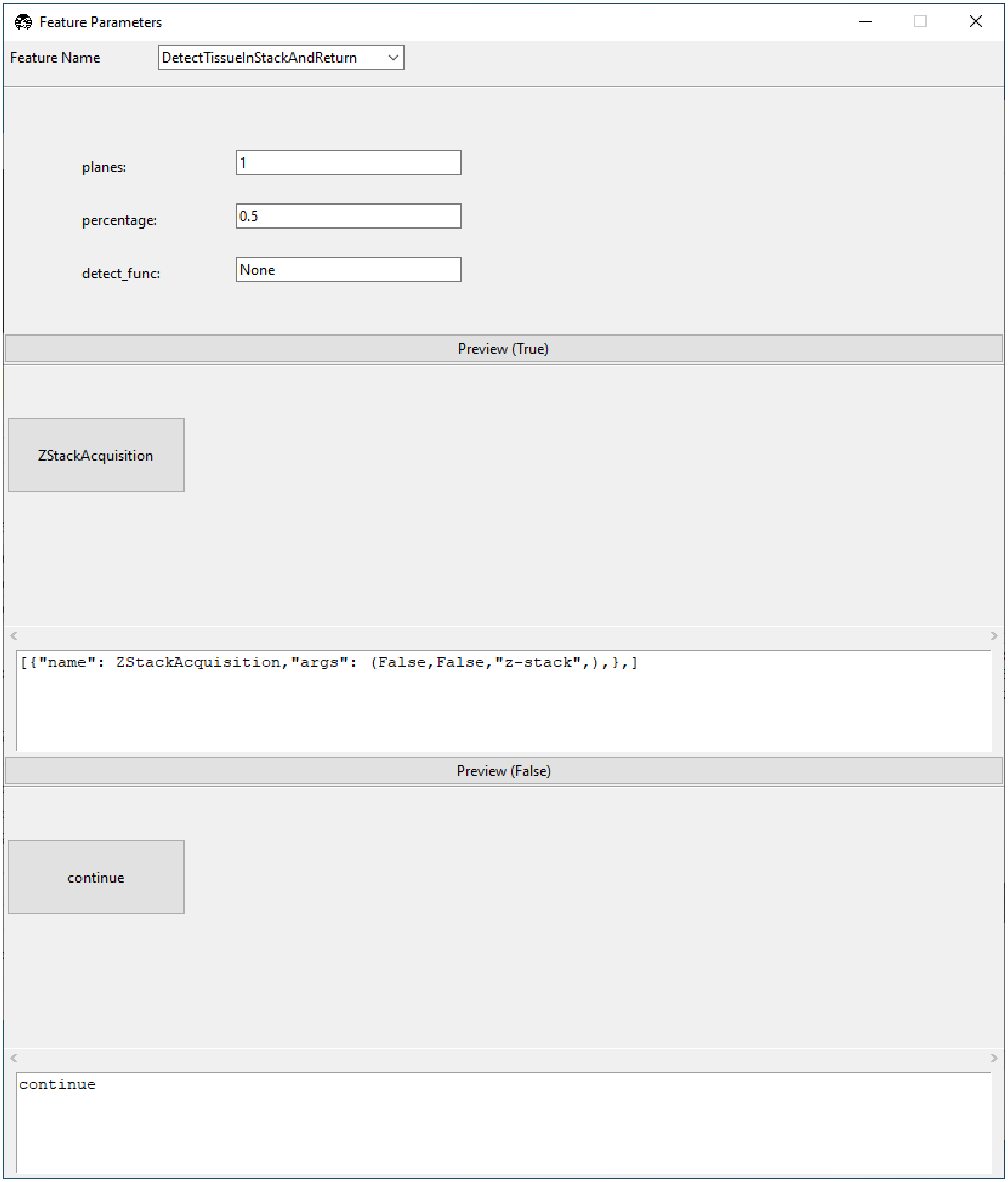

This contains the same settings for `DetectTissueInStackAndReturn` we saw before, but now also features GUI editing windows for the results of `true` and `false` decisions arising from this node.

Close the node window and press *Add* in the "Add New Feature List" window. This feature is now available under *Features* → *TestFeature* and can be run in "Customized" *acquisition mode*.

Select "Customized" acquisition mode, select *Features* → *TestFeature*, and press *Acquire*. For the positions shown at the start of this guide, the software will go to the first position in the multi-position table, decide there is tissue present, and take a z-stack. It will then go to the second position in the multi-position table, find there is no tissue, and decide not to take a z-stack. It will then exit the loop as no more positions are available in the multi-position table.

Now you can use this feature or build another smart acquisition routine suited to your microscope's needs.

#### 3.3 Write a Custom Plugin (Expert)

**navigate**'s *plugin system* enables users to easily incorporate new devices and integrate new features and acquisition modes. In this guide, we will add a new device, titled `CustomDevice`, and a dedicated GUI window to control it. This hypothetical `CustomDevice` is capable of moving a certain distance, rotating a specified number of degrees, and applying a force to halt its movement.

**navigate** plugins are implemented using a Model-View-Controller architecture. The model contains the device-specific code, the view contains the GUI code, and the controller contains the code that communicates between the model and the view.

---

##### 3.3.1 Initial Steps

To ease the addition of a new plugin, we have created a template plugin that can be used as a starting point.

- Go to navigate-plugin-template.
- In the upper right, click "Use this template" and then "Create a new repository".
- In this repository, rename the `plugin_device` folder to `custom_device`.
- Rename the file `plugin_device.py` to `custom_device.py`.

---

##### 3.3.2 Model Code

Create a new custom device using the following code.

```
class CustomDevice:
    """ A Custom Device Class """
    def __init__(self, device_connection, *args):
        """ Initialize the Custom Device

        Parameters
        ----------
        device_connection : object
            The device connection object
        args : list
            The arguments for the device
        """
        self.device_connection = device_connection

    def move(self, step_size=1.0):
        """ Move the Custom Device

        Parameters
```

(continues on next page)

(continued from previous page)

```
        ----------
        step_size : float
            The step size of the movement. Default is 1.0 micron.
        """
        print("*** Custom Device is moving by", step_size)

    def stop(self):
        """ Stop the Custom Device """
        print("*** Stopping the Custom Device!")

    def turn(self, angle=0.1):
        """ Turn the Custom Device

        Parameters
        ----------
        angle : float
            The angle of the rotation. Default is 0.1 degree.
        """
        print("*** Custom Device is turning by", angle)

    @property
    def commands(self):
        """ Return the commands for the Custom Device

        Returns
        -------
        dict
            The commands for the Custom Device
        """
        return {
            "move_custom_device": lambda *args: self.move(args[0]),
            "stop_custom_device": lambda *args: self.stop(),
            "rotate_custom_device": lambda *args: self.rotate(args[0]),
        }
```

All devices should be accompanied by synthetic versions, which enables the software to run without the hardware connected. Thus, in a manner that is similar to the `CustomDevice` class, we edit the code in `synthetic_device.py`, albeit without any calls to the device itself.

```
class SyntheticCustomDevice:
    """ A Synthetic Device Class """
    def __init__(self, device_connection, *args):
        """ Initialize the Synthetic Device

        Parameters
        ----------
        device_connection : object
            The device connection object
        args : list
            The arguments for the device
        """
        pass
```

(continues on next page)

(continued from previous page)

```
    def move(self, step_size=1.0):
        """ Move the Synthetic Device

        Parameters
        ----------
        step_size : float
            The step size of the movement. Default is 1.0 micron.
        """
        print("*** Synthetic Device receive command: move", step_size)

    def stop(self):
        """ Stop the Synthetic Device """
        print("*** Synthetic Device receive command: stop")

    def rotate(self, angle=0.1):
        """ Turn the Synthetic Device

        Parameters
        ----------
        angle : float
            The angle of the rotation. Default is 0.1 degree.
        """
        print("*** Synthetic Device receive command: turn", angle)

    @property
    def commands(self):
        """ Return the commands for the Synthetic Device.

        Returns
        -------
        dict
            The commands for the Synthetic Device
          """
        return {
            "move_custom_device": lambda *args: self.move(args[0]),
            "stop_custom_device": lambda *args: self.stop(),
            "rotate_custom_device": lambda *args: self.rotate(args[0]),
        }
```

Edit `device_startup_functions.py` to tell **navigate** how to connect to and start the `CustomDevice`. This is the portion of the code that actually makes a connection to the hardware. `load_device()` should return an object that can control the hardware.

**navigate** establishes communication with each device independently, and passes the instance of that device to class that controls it (e.g., in this case, the *CustomDevice* class). This allows **navigate** to be initialized with multiple microscope *configurations*, some of which may share devices.

```
# Standard library imports
import os
from pathlib import Path

# Third party imports
```

(continues on next page)

(continued from previous page)

```
# Local application imports
from navigate.tools.common_functions import load_module_from_file
from navigate.model.device_startup_functions import (
    auto_redial,
    device_not_found,
    DummyDeviceConnection,
)

DEVICE_TYPE_NAME = "custom_device"
DEVICE_REF_LIST = ["type"]

def load_device(configuration, is_synthetic=False):
    """ Load the Custom Device

    Parameters
    ----------
    configuration : dict
        The configuration for the Custom Device
    is_synthetic : bool
        Whether the device is synthetic or not. Default is False.

    Returns
    -------
    object
        The Custom Device object
    """
    return DummyDeviceConnection()

def start_device(microscope_name, device_connection, configuration, is_synthetic=False):
    """ Start the Custom Device

    Parameters
    ----------
    microscope_name : str
        The name of the microscope
    device_connection : object
        The device connection object
    configuration : dict
        The configuration for the Custom Device
    is_synthetic : bool
        Whether the device is synthetic or not. Default is False.

    Returns
    -------
    object
        The Custom Device object
    """
    if is_synthetic:
        device_type = "synthetic"
    else:
        device_type = configuration["configuration"]["microscopes"][microscope_name][
```

(continues on next page)

(continued from previous page)

```
            "custom_device"
        ]["hardware"]["type"]

    if device_type == "CustomDevice":
        custom_device = load_module_from_file(
            "custom_device",
            os.path.join(Path(__file__).resolve().parent, "custom_device.py"),
        )
        return custom_device.CustomDevice(
            microscope_name, device_connection, configuration
        )
    elif device_type == "synthetic":
        synthetic_device = load_module_from_file(
            "custom_synthetic_device",
            os.path.join(Path(__file__).resolve().parent, "synthetic_device.py"),
        )
        return synthetic_device.SyntheticDevice(
            microscope_name, device_connection, configuration
        )
    else:
        device_not_found(microscope_name, device_type)
```

##### 3.3.3 View Code

To add a GUI control window, go to the `view` folder, rename `plugin_name_frame.py` to `custom_device_frame.py`, and edit the code as follows.

```
# Standard library imports
import tkinter as tk
from tkinter import ttk

# Third party imports

# Local application imports
from navigate.view.custom_widgets.LabelInputWidgetFactory import LabelInput

class CustomDeviceFrame(ttk.Frame):
    """ The Custom Device Frame """
    def __init__(self, root, *args, **kwargs):
        """ Initialize the Custom Device Frame

        Parameters
        ----------
        root : object
            The root Tk object
        args : list
            The arguments for the Custom Device Frame
        kwargs : dict
```

(continues on next page)

(continued from previous page)

```
            The keyword arguments for the Custom Device Frame
        """
        ttk.Frame.__init__(self, root, *args, **kwargs)

        # Formatting
        tk.Grid.columnconfigure(self, "all", weight=1)
        tk.Grid.rowconfigure(self, "all", weight=1)

        # Dictionary for widgets and buttons
        #: dict: Dictionary of the widgets in the frame
        self.inputs = {}

        self.inputs["step_size"] = LabelInput(
            parent=self,
            label="Step Size",
            label_args={"padding": (0, 0, 10, 0)},
            input_class=ttk.Entry,
            input_var=tk.DoubleVar(),
        )
        self.inputs["step_size"].grid(row=0, column=0, sticky="N", padx=6)
        self.inputs["step_size"].label.grid(sticky="N")
        self.inputs["angle"] = LabelInput(
            parent=self,
            label="Angle",
            label_args={"padding": (0, 5, 25, 0)},
            input_class=ttk.Entry,
            input_var=tk.DoubleVar(),
        )
        self.inputs["angle"].grid(row=1, column=0, sticky="N", padx=6)
        self.inputs["angle"].label.grid(sticky="N")

        self.buttons = {}
        self.buttons["move"] = ttk.Button(self, text="MOVE")
        self.buttons["rotate"] = ttk.Button(self, text="ROTATE")
        self.buttons["stop"] = ttk.Button(self, text="STOP")
        self.buttons["move"].grid(row=0, column=1, sticky="N", padx=6)
        self.buttons["rotate"].grid(row=1, column=1, sticky="N", padx=6)
        self.buttons["stop"].grid(row=2, column=1, sticky="N", padx=6)

    # Getters
    def get_variables(self):
        variables = {}
        for key, widget in self.inputs.items():
            variables[key] = widget.get_variable()
        return variables

    def get_widgets(self):
        return self.inputs
```

**Tip:** **navigate** comes equipped with a large number of validated widgets, which prevent users from entering invalid values that can crash the program or result in undesirable outcomes. It is highly recommended that you use these, which include the following:

- The `LabelInput` widget conveniently combines a label and an input widget into a single object. It is used to create the `step_size` and `angle` widgets in the code above.
- The `LabelInput` widget can accept multiple types of `input_class` objects, which can include standard tkinter widgets (e.g., spinbox, entry, etc.) or custom widgets. In this example, we use the `ttk.Entry` widget.
- Other examples of validated widgets include a `ValidatedSpinbox`, `ValidatedEntry`, `ValidatedCombobox`, and `ValidatedMixin`.
- Please see the `navigate.view.custom_widgets` module for more details.

---

##### 3.3.4 Controller Code

Now, let's build a controller. Open the `controller` folder, rename `plugin_name_controller.py` to `custom_device_controller.py`, and edit the code as follows.

```
# Standard library imports
import tkinter as tk

# Third party imports

# Local application imports
from navigate.controller.sub_controllers.gui_controller import GUIController

class CustomDeviceController(GUIController):
    """ The Custom Device Controller """
    def __init__(self, view, parent_controller=None):
        """ Initialize the Custom Device Controller

        Parameters
        ----------
        view : object
            The Custom Device View object
        parent_controller : object
            The parent (e.g., main) controller object
        """
        super().__init__(view, parent_controller)

        # Get the variables and buttons from the view
        self.variables = self.view.get_variables()
        self.buttons = self.view.buttons

        # Set the trace commands for the variables associated with the widgets in the␣
↪View.
        self.buttons["move"].configure(command=self.move_device)
        self.buttons["rotate"].configure(command=self.rotate_device)
        self.buttons["stop"].configure(command=self.stop_device)

    def move_device(self, *args):
        """ Listen to the move button and move the Custom Device upon clicking.
```

(continues on next page)

(continued from previous page)

```
        Parameters
        ----------
        args : list
            The arguments for the move_device function. Should be included as the␣
↪tkinter event
            is passed to this function.
        """
        self.parent_controller.execute(
            "move_custom_device", self.variables["step_size"].get()
        )

    def rotate_device(self, *args):
        """ Listen to the rotate button and rotate the Custom Device upon clicking.

        Parameters
        ----------
        args : list
            The arguments for the rotate_device function. Should be included as the␣
↪tkinter event
            is passed to this function.
        """
        self.parent_controller.execute(
            "rotate_custom_device", self.variables["angle"].get()
        )

    def stop_device(self, *args):
        """ Listen to the stop button and stop the Custom Device upon clicking.

        Parameters
        ----------
        args : list
            The arguments for the stop_device function. Should be included as the␣
↪tkinter event
            is passed to this function.
        """
        self.parent_controller.execute("stop_custom_device")
```

In each case above, the sub-controller for the `custom-device` establishes what actions should take place once a button in the view is clicked. In this case, the methods `move_device`, `rotate_device`, and `stop_device`. This triggers a sequence of events:

- The sub-controller passes the command to the parent controller, which is the main controller for the software.
- The parent controller passes the command to the model, which is operating in its own sub-process, using an event queue. This eliminates the need for the controller to know anything about the model and prevents race conditions.
- The model then executes command, and any updates to the controller from the model are relayed using another event queue.

##### 3.3.5 Plugin Configuration

Next, update the `plugin_config.yml` file as follows:

```yaml
name: Custom Device
view: Popup
```

Remove the folder `./model/features`, the file `feature_list.py`, and the file `plugin_acquisition_mode.py`. The plugin folder structure is as follows.

```
custom_device/
    ├── controller/
    │   └── custom_device_controller.py
    |
    ├── model/
    |   └── devices/
    │       └── custom_device/
    │           ├── device_startup_functions.py
    │           ├── custom_device.py
    │           └── synthetic_device.py
    ├── view/
    |   └── custom_device_frame.py
    │
    └── plugin_config.yml
```

**Install the plugin using one of two methods:**

- Install a plugin by putting the whole plugin folder directly into `navigate/plugins/`. In this example, put `custom_device` folder and all its contents into `navigate/plugins`.
- Alernatively, install this plugin through the menu *Plugins* → *Install Plugin* by selecting the plugin folder.

The plugin is ready to use. For this plugin, you can now specify a CustomDevice in the `configuration.yaml` as follows.

```yaml
microscopes:
    microscope_1:
        daq:
            hardware:
                name: daq
                type: NI
        ...
        custom_device:
            hardware:
                type: CustomDevice
        ...
```

The `custom_device` will be loaded when **navigate** is launched, and it can be controlled through the GUI.

### USER INTERFACE WALKTHROUGH

**navigate**'s user interface is modular, and designed to be reconfigurable to a user's preferences. At a high level, it is split into a menu bar, an acquisition bar, settings notebooks, and display notebooks.

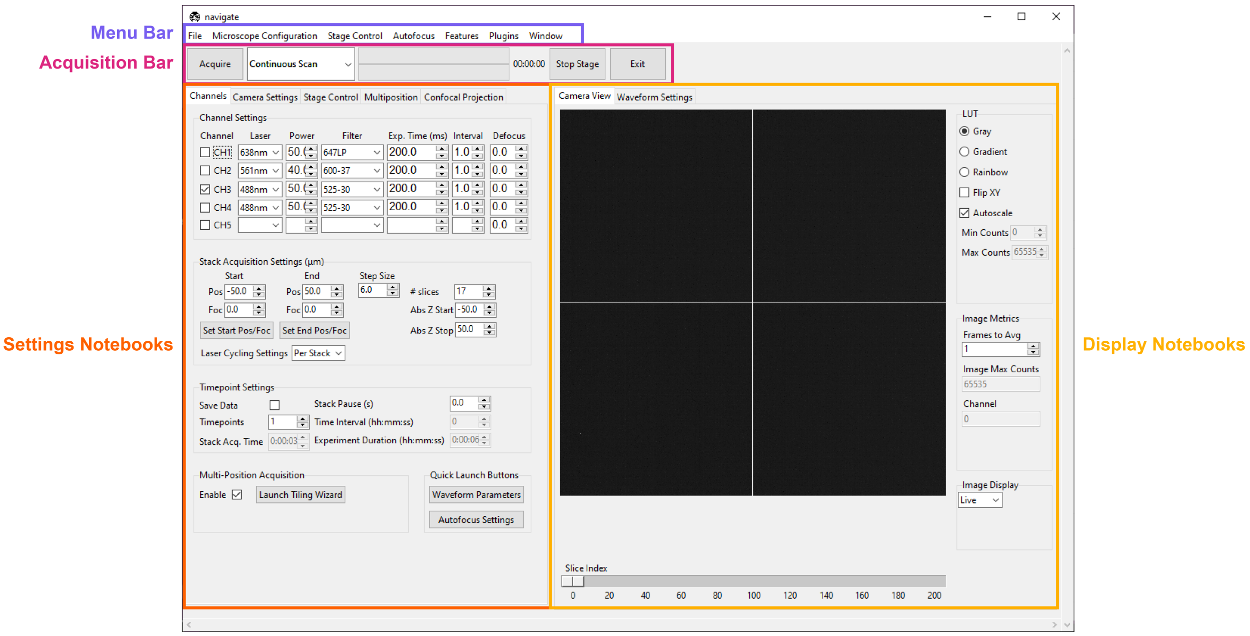

#### 4.1 Menu Bar

The menu bar is an entry point for much of **navigate**.

##### 4.1.1 File

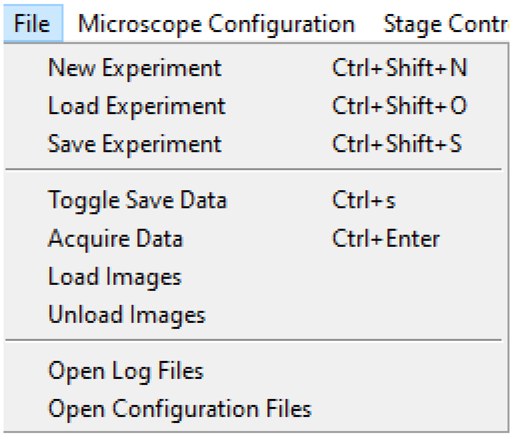

The *File* menu lets a user create, load and save *experiment files*, which store the states of the GUI and the hardware. This is useful if a user wants to perform an experiment with the same parameters multiple times, but close the software in between acquisitions. To facilitate reproducibility, an *experiment.yml* file is always saved with collected image data.

The *File* menu also provides access to toggle the *Save Data* flag, also found under *Timepoint Settings* in the *Channels Settings Notebook*, and to start an acquisition (which can also be done by pressing *Acquire* in the *Acquisition Bar*). Loading and unloading images only works if there is a *synthetic camera*. In this case, the camera loads images to display in lieu of the simulated noise usually generated by the synthetic camera.

*Open Log Files* opens the folder containing the software's log files. This is helpful for debugging code and configuration problems.

*Open Configuration Files* opens the folder containing the software's *configuration file*.

---

##### 4.1.2 Microscope Configuration

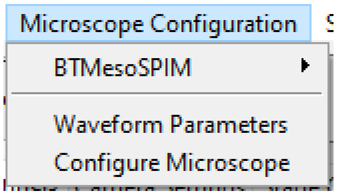

The *Microscope Configuration* menu is split into two parts, above and below the horizontal divider.

Above the horizontal divider it lists the names of all microscopes named in the `microscopes` section of the *configuration file*. This allows users to readily switch between different microscopes, each with their own hardware configurations. Mousing over a microscope name reveals all zoom values available under the *Mechanical Zoom*. Selecting one of these zoom values changes the magnification of the microscope.

Below the horizontal divider is access to the *Waveform Parameters* settings panel and the *Configure Microscope* settings panel.

---

##### 4.1.3 Stage Control

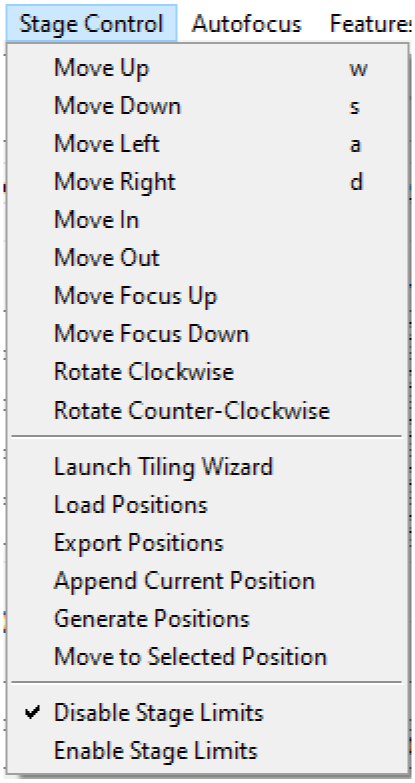

The stage control menu is split by horizontal dividers into three parts.

The top part provides similar functionality to the *Stage Control Settings Notebook*. It allows movement of the stage along `X`, `Y`, `Z`, `focus` and `Theta`. The `w`, `s`, `a` and `d` keys are bound to movement in `X` and `Y`, and these can be used to scroll around a sample.

The middle part provides similar functionality to the *Multiposition Settings Notebook*. Here, a user can launch the *Tiling Wizard*, load and export (save) positions stored in the *Multiposition Settings Notebook*, and add the current stage position to the multiposition table.

The bottom part of the menu is used to enable and disable the stage limits set in the configuration file (see the *stage subsection*).

---

##### 4.1.4 Autofocus

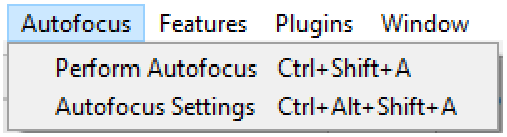

The autofocus menu has two options: *Perform Autofocus*, which autofocus the sample using the current autofocus settings, and *Autofocus Settings*, which launches the *Autofocus Settings* popup.

---

##### 4.1.5 Features

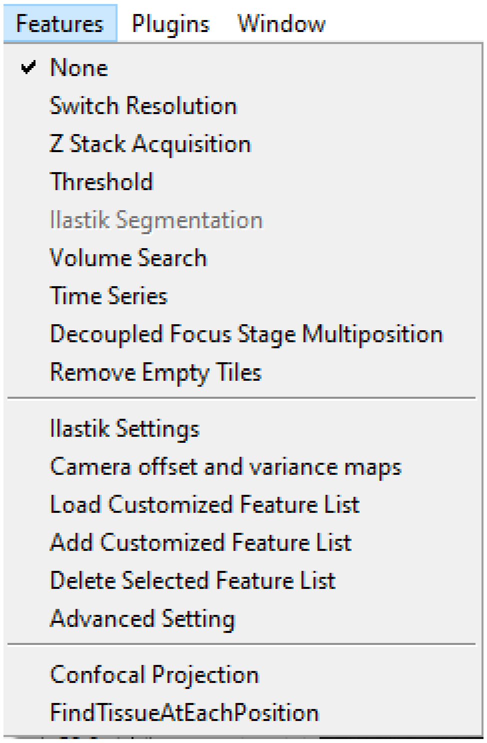

This menu provides access to acquisition feature lists. An explanation of features, feature lists, and the use and operation of this menu is provided under *Reconfigurable Acquisitions Using Features*.

---

##### 4.1.6 Plugins

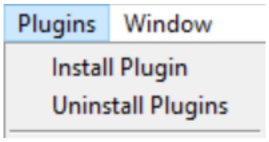

This menu provides an access point for *plugins* that feature a popup GUI.

---

##### 4.1.7 Window

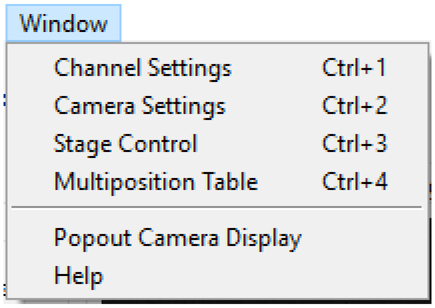

This menu is split into two parts by a horizontal divider and provides some GUI controls.

The top part allows the user to switch between the main *Settings Notebooks*.

The bottom part provides an option to move the camera display to a popup window and *Help* brings the user to the online documentation for **navigate**.

---

#### 4.2 Acquisition Bar

Left-to-right, the acquisition bar provides

- An *Acquire* button, which starts acquisition.
- A drop-down menu providing a selection of acquisition modes.
- A progress bar indicating how far the program has made it through an acquisition. The top bar indicates progress on the current z-stack, whereas the bottom indicates progress for the entire acquisition.
- An approximate estimate of how much time is left in the acquisition.
- An emergency *Stop Stage* button, which instantly halts all stage movement.
- An *Exit Button*, which quits the software.

---

#### 4.3 Settings Notebooks

The settings notebooks are a series of tabs that control microscope settings, including laser power, camera settings and stage positions and many others.

---

##### 4.3.1 Channels

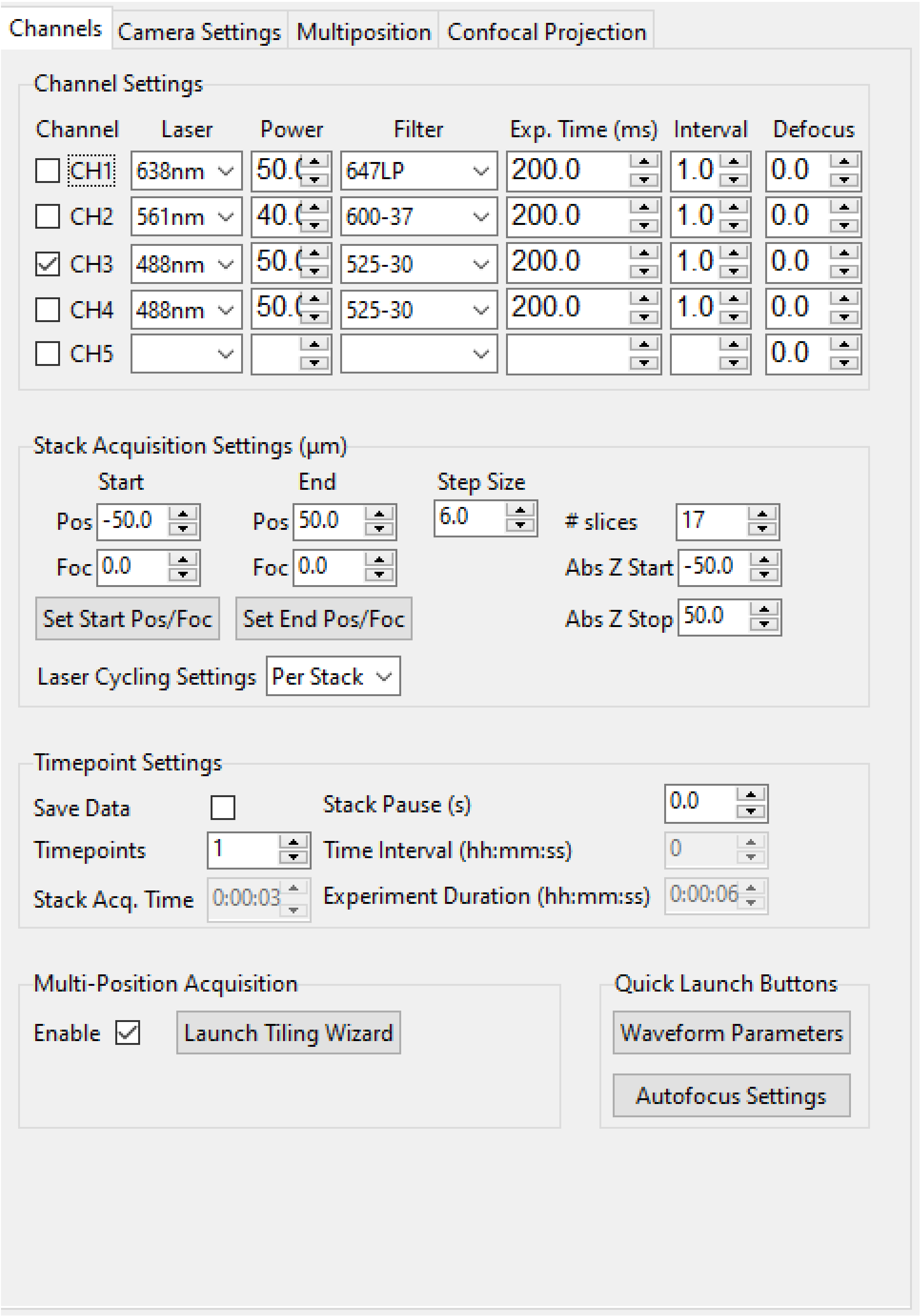

The *Channels* Settings Notebook is a tab (optionally, a popup if right-clicked on) split into five sections: *Channel Settings*, *Stack Acquisition Settings (µm)*, *Timepoint Settings*, *Multi-Position Acquisition* and *Quick Launch Buttons*.

##### Channel Settings

This is used to set up acquisition color channels. A channel is considered to be a combination of an illuminating laser wavelength and a detection filter. Each channel has its own power, exposure time, interval and defocus. The checkbox on the left indicates if a channel should be used (is selected) during acquisition. An acquisition may loop through the channels in sequence.

- *Laser* is the name of the laser, taken from the *configuration file*, and usually expressed in nanometers.
- *Power* is the power of the laser between 0 and 100 percent.
- *Filter* is the name of the filter selected in the detection path filter wheel. Filter names are stored in the configuration file.
- *Exp. Time (ms)* is the exposure time of the camera in milliseconds.
- *Interval* indicates how often this channel should be used in an acquisition. For example, in two-color imaging, CH1 may image a process twice as fast as what is labelled in CH2. Setting the CH2 interval to 2 allows a user to image the processes in both channels at a similar rate. This will be implemented in future releases of the software.
- *Defocus* indicates the defocus between two channels in micrometers. The defocus values are always relative to the focus of the first channel imaged. This setting is useful for compensating for chromatic aberration.

---

##### Stack Acquisition Settings

These are the settings used for a standard Z-Stack Acquisition.

*Pos* indicates z-positions. *Foc* indicates focus positions. The z-stack can optionally ramp through `focus` along with `Z`.

*Start* and *End* are always expressed relative to the center of the z-stack. *Abs Z Start* and *Abs Z Stop* provide true stage positions at the start and end of the z-stack.

The buttons *Set Start Pos/Foc* and *Set End Pos/Foc* grab the current `Z` and `focus` positions from the stage and enter them into the corresponding start and end (stop) GUI boxes.

The *Step Size* is expressed in microns and can be modified by the user. Upon modification, *# slices* will automatically update.

*Laser Cycling Settings* provide the options “Per Stack” and “Per Z”. In “Per Stack” mode, the software will move through all positions before changing to another color channel. In “Per Z” mode, the software will acquire all color channels selected before moving to the next position in the z-stack.

---

##### Timepoint Settings

These are used for acquiring data over multiple timepoints and for toggling the option to save data.

- *Save Data* tells the software to save acquired data to disk when checked. If this is selected, a *saving popup window* will appear when *Acquire* is pressed, unless the user is in “Continuous Scan” mode, which is designed for live previews only.
- *Timepoints* indicates how many time points this acquisition should acquire.
- *Stack Acq Time* provides an estimate of how long a single z-stack will take to acquire.
- *Stack Pause (s)* indicates how much waiting time the software should introduce in between acquisition steps (e.g. in between taking z-stacks).

- *Time Interval (hh:mm:ss)* provides an estimate of how long each time point takes to acquire. This is (stack acquisition + stack pause) x number of channels to image.
- *Experiment Duration (hh:mm:ss)* provides an estimate of how long the full acquisition will take.

**Note:** The *Stack Acq Time* and *Experiment Duration (hh:mm:ss)* do not account for stage movement time. Thus, for stages with serial communication protocols, or stages with slow movement, these estimates will be an underestimate. Future releases will account for stage movement time to provide a more accurate estimate.

##### Multi-Position Acquisition

This contains settings to set up acquisition over multiple positions in the sample, e.g. tiling.

- *Enable* indicates that the software should move through the positions listed in the *Multiposition Settings Notebook* during the acquisition.
- *Launch Tiling Wizard* launches the *Tiling Wizard*.

##### Quick Launch Buttons

This provides access to the *Waveform Parameters* and *Autofocus Settings* popups.

##### 4.3.2 Camera Settings

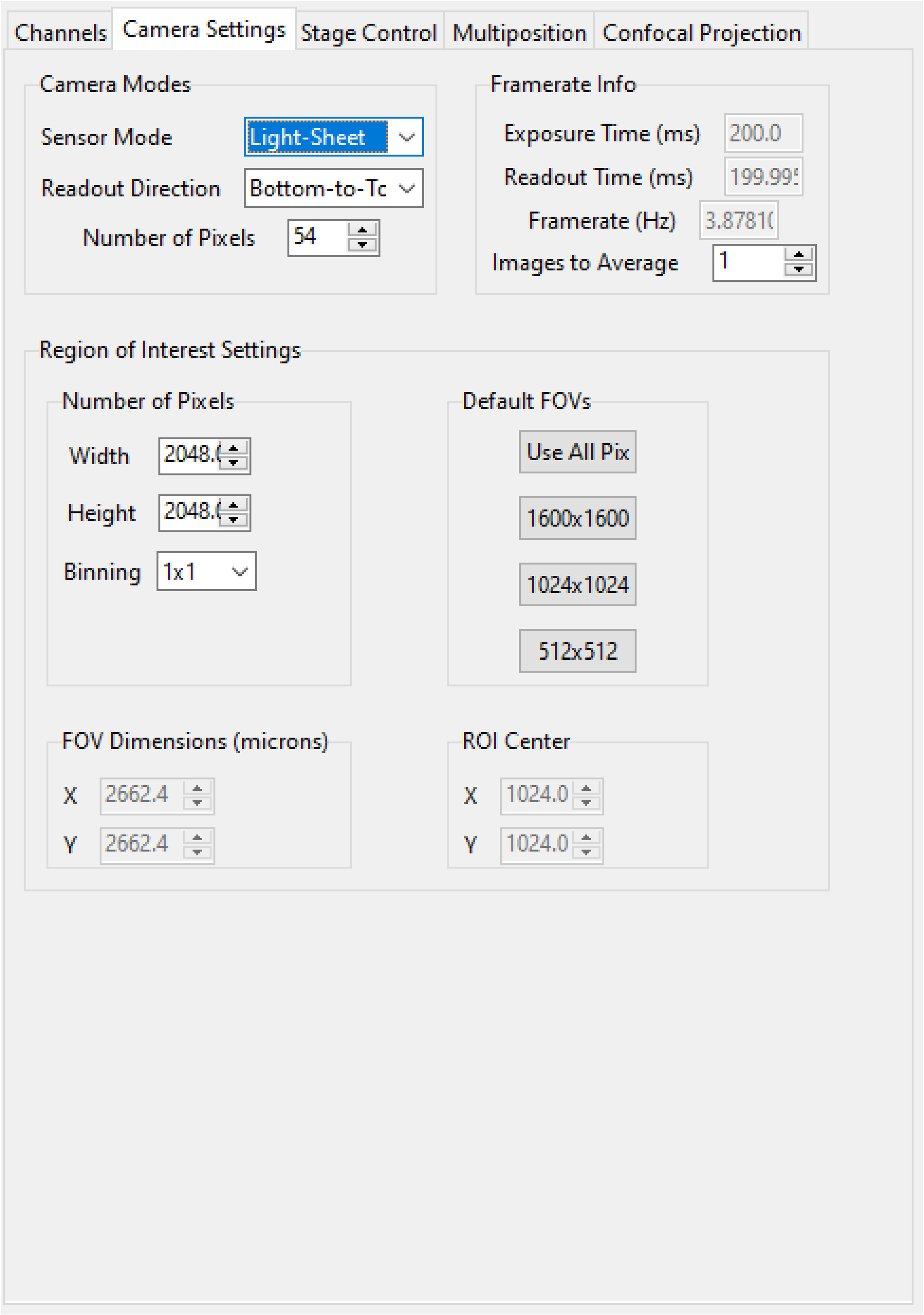

The *Camera Settings* Notebook is a tab (optionally, a popup) that controls the camera. It is split into three sections: *Camera Modes*, *Framerate Info* and *Region of Interest Settings*.

##### Camera Modes

The *Camera Modes* section is designed for switching between normal mode of operation, where the camera exposes all pixels semi-simultaneously, and light-sheet mode (a.k.a rolling shutter mode), where the camera exposes only a few pixels at a time, and progressively images from the top to the bottom of the camera chip or vice versa.

- *Sensor Mode* is used to switch between “Normal” and “Light-Sheet” modes.
- *Readout Direction* indicates if the rolling shutter should move from the bottom to the top of the camera chip or vice versa.
- *Number of Pixels* sets the rolling shutter width of the camera.

---

##### Framerate Info

This displays information concerning the speed of acquisition and optionally allows the user to average these values over multiple images.

- *Exposure Time (ms)* displays the set camera exposure time.
- *Readout Time (ms)* displays how long it takes to read a frame from the camera. This includes exposure time.
- *Framerate (Hz)* displays how long it takes to acquire an image. This is based on an internal “wait ticket” approach, where the software times how long it waits for a frame to come in after receiving the previous frame. This frequency includes not only camera readout time, but, e.g. how long the software had to wait for the stage to finish moving before taking the next image in a z-stack. It is the most accurate time estimate in the software.
- *Images to Average* tells the camera to average frames. This will be implemented in future releases of the software.

---

##### Region of Interest Settings

These allows the user to set the size of the region of interest in pixels. The camera can also be told to bin pixels. The corresponding field of view is displayed by calculating the number of pixels multiplied by the camera’s effective pixel size, which is set in the *Mechanical Zoom* section of the configuration file.

*Default FOVs* includes buttons to quickly change the FOV to preset values.

*ROI center* indicates about what point the pixels crop on the camera.

---

##### 4.3.3 Stage Control

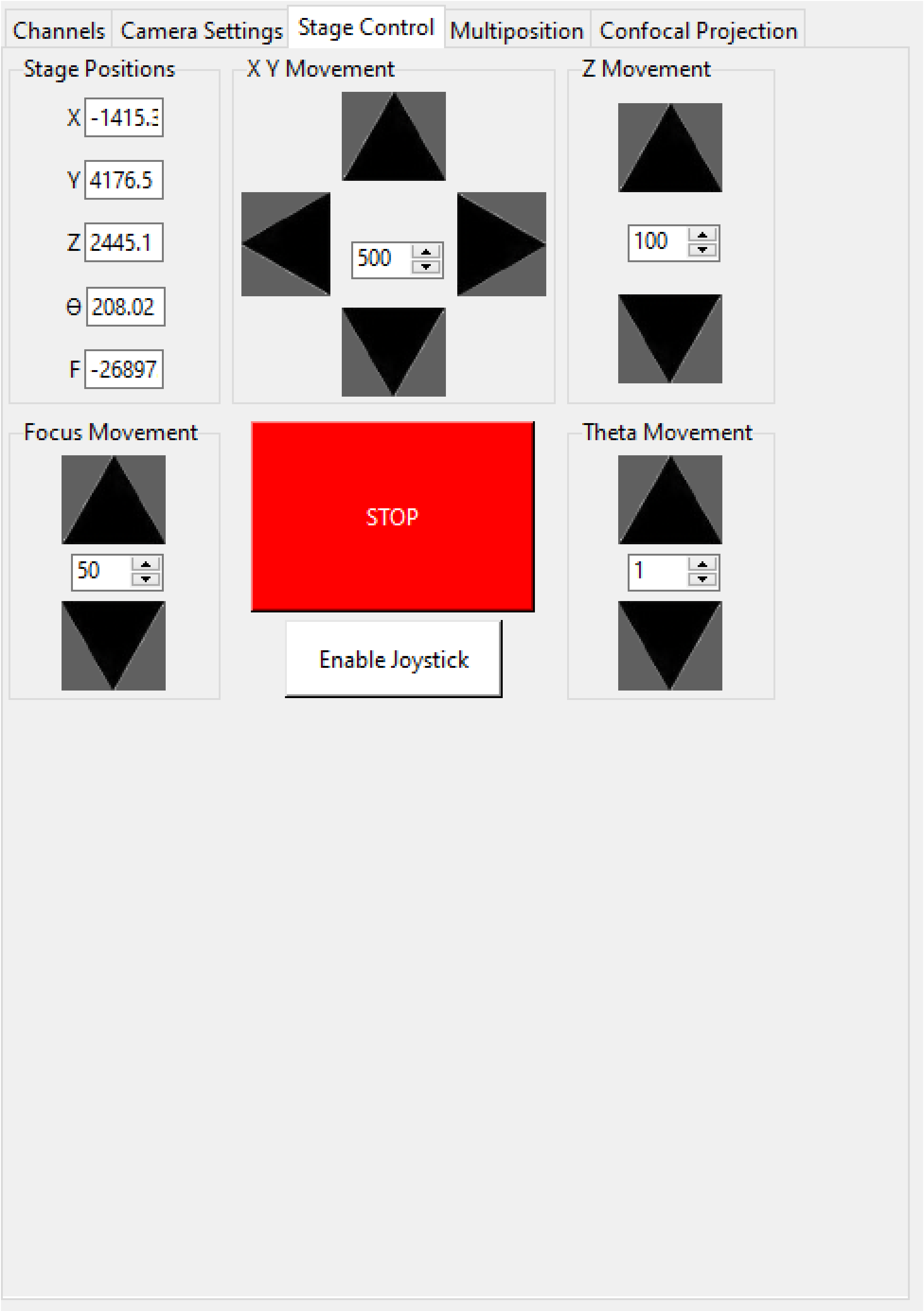

The *Stage Control* Settings Notebook is a tab (optionally, a popup) that controls the stage positions. It is split into six parts: *Stage Positions*, *X Y Movement*, *Z Movement*, *Focus Movement*, *Theta Movement*, and includes an emergency *STOP* button, as well as a button to *Enable Joystick*/*Disable Joystick*.

**Note:**

- **The joystick button will only appear if the `configuration.yaml` file specifies** which axes are controlled by the joystick. For example:

```yaml
stage:
  hardware:
    -
      name: stage
      type: PI
      axes: [x, y, z, theta, f]
      axes_mapping: [1, 2, 3, 4, 5]
  joystick_axes: [x, y, z]
```

- Any stage axes that are loaded as a *synthetic_stage* will have disabled buttons.

---

By default, the stage is expected to have `X`, `Y`, `Z`, `Focus` and `Theta` (rotation) axes. If a stage does not have one of these axes, the user can choose to not use that control. See the *stage subsection* for more information.

---

##### Stage Positions

The entry boxes report the current position of each stage axis. If a user changes a value in an entry box, the stage axis will move to that value (provided it is within the stage bounds if stage limits are enabled, see *here*).

> **Warning:** If the value in the entry box is changed, the stage will move to that value. Such actions may result in the stage crashing into the sample or the objective lens. As such, we highly recommend that you keep the stage limits enabled.

---

##### XY Movement

This includes the movement buttons for the `X` and `Y` axes. The left and right buttons control `X`, while the up and down buttons control `Y`. The entry box in the middle of the buttons indicates the step size along these axes in microns. It can be changed by the user.

---

##### Z Movement

This controls the movement of the `Z` stage. The entry box indicates the step size along this axis and it can be changed by the user.

---

###### Focus Movement

This controls the movement of the `Focus` stage. The entry box indicates the step size along this axis and it can be changed by the user.

---

###### Theta Movement

This controls the movement of the `Theta` stage (i.e., sample rotation). The entry box indicates the step size along this axis and it can be changed by the user.

---

###### Buttons

The *STOP* button halts all stage axes and updates the stage positions to wherever the stage stopped.

The *Enable Joystick* button disables control over the axes associated with the joystick (see the *stage subsection*).

---

**Note:** It is not necessary to press this button to use a joystick. The joystick can be used along with the software controls. However, if a user is running the acquisition in "Continuous Scan" mode and uses the joystick without pressing *Enable Joystick*, the stage positions may not update unless *STOP* is also pressed. In "Continuous Scan" mode, if a user tries to move with the joystick and then the software stage controls without first pressing *STOP*, it is likely the stage will update to the software's position of choice and undo the joystick movement.

---

**Tip:** For a large monitor, it is often helpful to convert the *Stage Control Settings Notebook* to a popup. Right click on the tab and press *Popout Tab*.

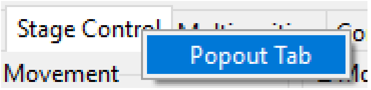

Once this is done, it should be possible to move the stage controls next to the main **navigate** window.

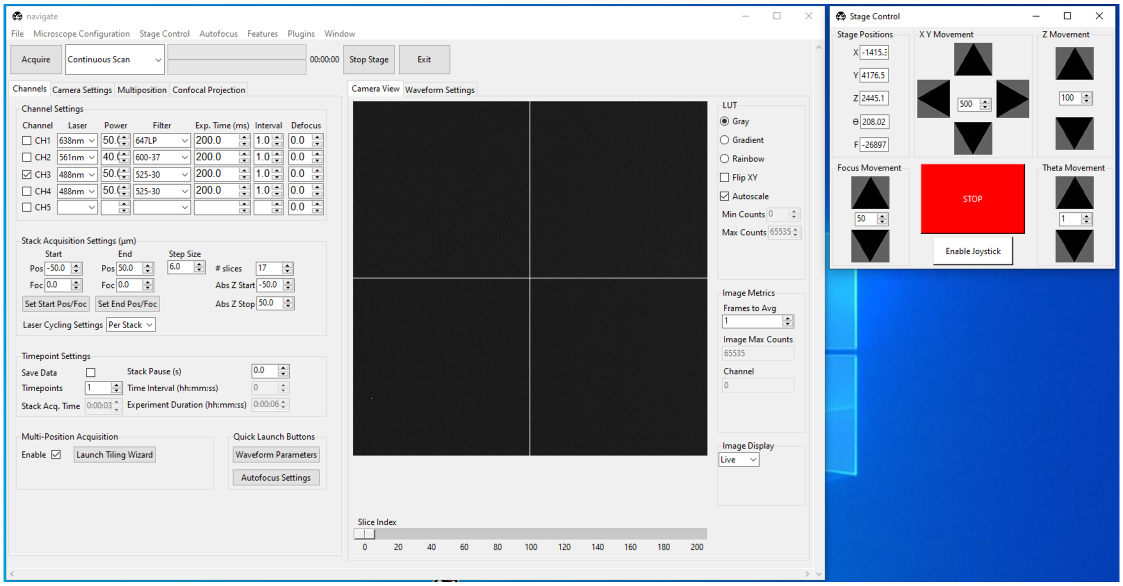
navigate
File Microscope Configuration Stage Control Autofocus Features Plugins Window
Acquire
Continuous Scan
00:00:00
Stop Stage
Exit
Channels Camera Settings Multiposition Confocal Projection
Channel Settings
Channel Laser Power Filter Exp. Time (ms) Interval Defocus
CH1 638nm 647LP 200.0 1.0 0.0
CH2 561nm 600-37 200.0 1.0 0.0
CH3 488nm 525-30 200.0 1.0 0.0
CH4 488nm 525-30 200.0 1.0 0.0
CH5 0.0
Stack Acquisition Settings (µm)
Start End Step Size
Pos -50.0 Pos 50.0 6.0 # slices 17
Foc 0.0 Foc 0.0 Abs Z Start -50.0
Set Start Pos/Foc Set End Pos/Foc Abs Z Stop 50.0
Laser Cycling Settings Per Stack
Timepoint Settings
Save Data Stack Pause (s) 0.0
Timepoints 1 Time Interval (hh:mm:ss) 0
Stack Acq. Time 0:00:03 Experiment Duration (hh:mm:ss) 0:00:06
Multi-Position Acquisition
Enable Launch Tiling Wizard
Quick Launch Buttons
Waveform Parameters
Autofocus Settings
Camera View Waveform Settings
LUT
Gray
Gradient
Rainbow
Flip XY
Autoscale
Min Counts 0
Max Counts 65535
Image Metrics
Frames to Avg
1
Image Max Counts
65535
Channel
0
Image Display
Live
Slice Index
0 20 40 60 80 100 120 140 160 180 200
Stage Control
Stage Positions
X -1415.3
Y 4176.5
Z 2445.1
Θ 208.02
F -26897
X Y Movement
500
Z Movement
100
Focus Movement
50
STOP
Enable Joystick
Theta Movement
1

##### 4.3.4 Multiposition

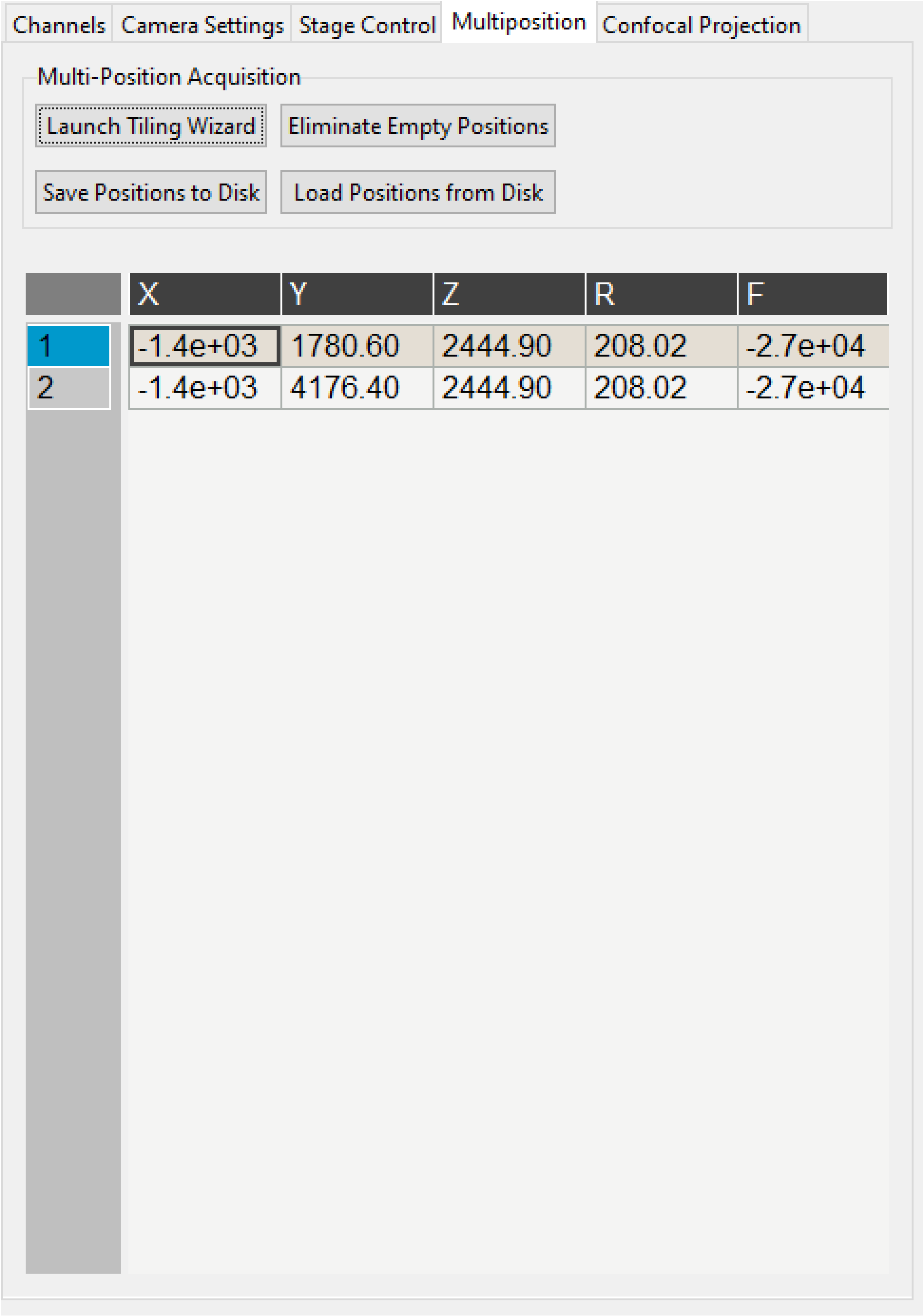

The *Multiposition* Settings Notebook is a tab (optionally, a popup) that helps the user set up and visualize a multi-position acquisition for tiling a large sample. It it split into two parts: buttons and the multi-position table.

##### Buttons

- *Launch Tiling Wizard* launches the *Tiling Wizard*
- *Eliminate Empty Positions* is not implemented and does nothing.
- *Save Positions To Disk* saves the multi-position table to a file.
- *Load Positions From Disk* loads a multi-position file into the table.

---

##### Multi-Position Table

The multi-position table lists stage positions that are included in a multi-position acquisition.

- `Double-clicking` on the integer to the left of a row moves the stage to that position.
- `Double-clicking` on a table cell allows the user to edit the stage position in that cell.
- `Right-clicking` on the integer to the left of a row yields a popup with four options:

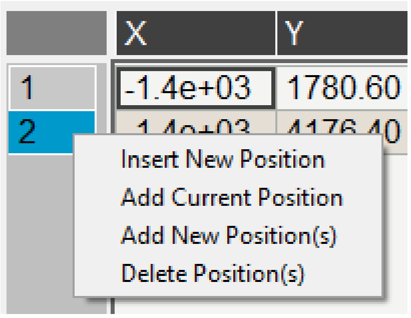

  – *Insert New Position* adds an empy row to the table.
  – *Add Current Position* adds a row containing the current stage position to the table.
  – *Add New Position(s)* yields a popup that asks the user how many new rows to add and then inserts that number of empty rows upon confirmation.
  – *Delete Position(s)* deletes the selected positions. Selection is indicated by a blue highlight of the integer to the left of a row.

---

#### 4.4 Display Notebooks

The display notebooks provide visual feedback of the images taken on the camera and of the galvo and remote focus waveforms sent to the DAQ.

---

##### 4.4.1 Camera View

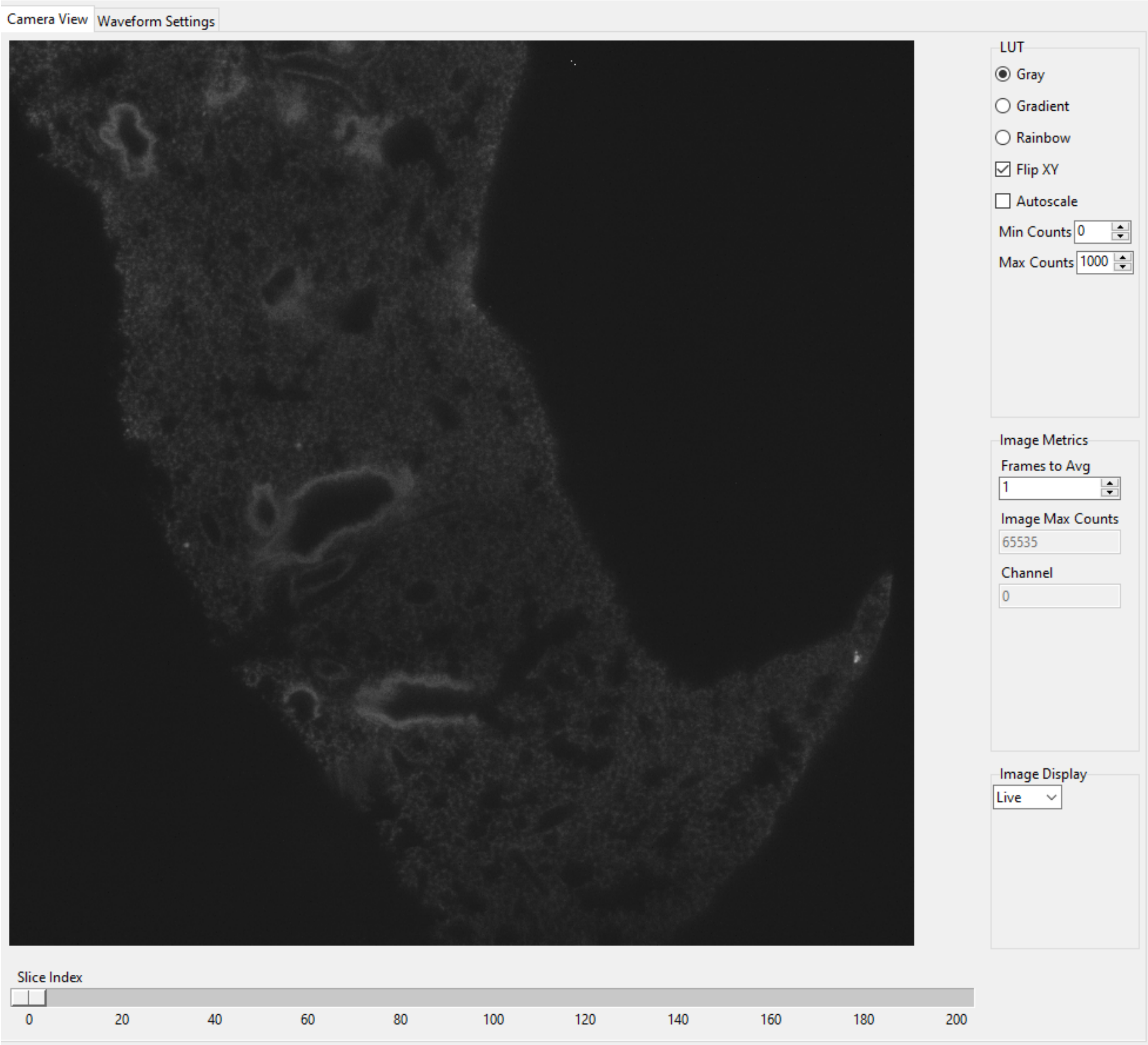

The *Camera View* Notebook is is a tab (optionally, a popup) that is split into two parts. The left part displays the latest image acquired by the camera. The right part modifies this display and is split into *LUT*, *Image Metrics*, and *Image Display*.

`Left-clicking` on the image toggles crosshairs that indicate the center of the field of view.

##### LUT

The *LUT* section of the camera view allows the user to change the lookup table the image uses to display. The options are *Gray*, *Gradient* and *Rainbow*.

*Flip XY* transposes the image in the display. This can produce intuitive results in the display when clicking on the `X` or `Y` stage movements buttons (i.e. with *Flip XY* enabled, the sample moves along the direction expected when a stage movement button is clicked).

*Autoscale* toggles automatic image histogram scaling on and off. When *Autoscale* is enabled, the image automatically scales intensity between the minimum and maximum pixel value in the image produced by the camera. When *Autoscale* is disabled, the image is scaled between *Min Counts* and *Max Counts*.

---

##### Image Metrics

*Frames to Avg* is unimplemented, but should average this many frames coming from the camera and display the average in the viewer. It will be implemented in future releases of the software.

*Image Max Counts* displays the maximum pixel count in the image.

*Channel* indicates which color channel is displayed. It indexes into the selected channels in the *Channel Settings* (i.e. `0` is the first selected channel).

---

##### Image Display

This should toggle in between live mode and maximum projections in multiple dimensions, but it is currently not implemented. This is useful for visual inspection of the data as it is being acquired, and will be implemented in future releases of the software.

---

##### 4.4.2 Waveform Settings

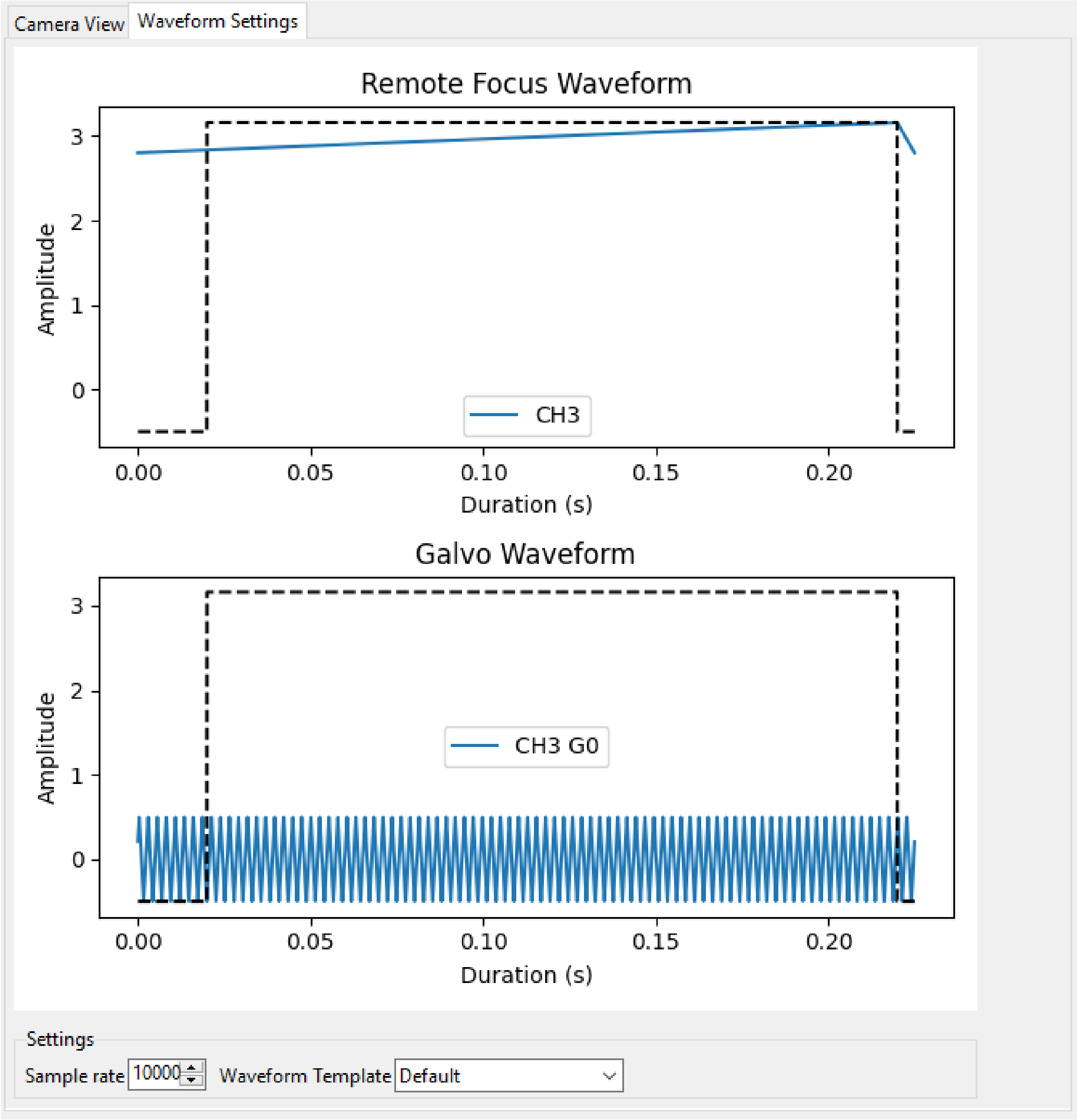

*Waveform Settings* is a tab (optionally, a popup) split into two sections: a waveform display section at the top and a *Settings* section at the bottom.

##### Waveform Display

The waveform display shows the waveforms sent to the remote focus devices (top) and the galvos (bottom). Each channel and each device gets its own color, which is then displayed in the legend. The dotted black line indicates when the camera is acquiring in relation to the waveforms. This can be considered identical to what is sent to the DAQ.

---

##### Settings

*Sample Rate* changes the frequency of the samples sent to the DAQ. It is not recommended that a user change this.

*Waveform Template* changes the *waveform template* used to generate the waveforms.

---

#### 4.5 Additional GUIs

This section includes popups and other non-main sections of the GUI.

---

##### 4.5.1 File Saving Dialog

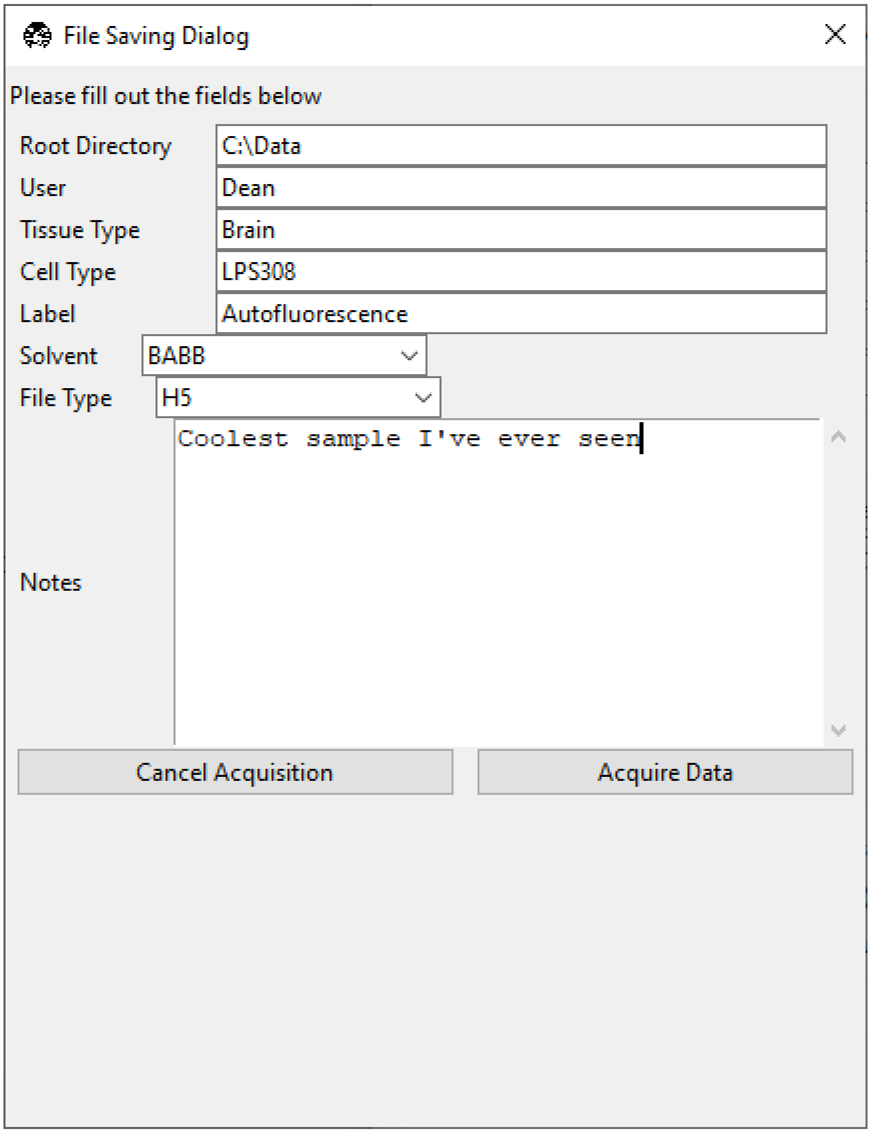

The file saving dialog pops up if an *acquisition mode* other than "Continuous Scan" is selected and *Save Data* is checked.

- *Root Directory* indicates the local directory to which the software will save the data.
- *User* is the name of the user acquiring the data.
- *Tissue Type* is the type of tissue being imaged.
- *Cell Type* is the cell type being imaged.
- *Label* indicates the dyes used in the acquisition.
- *Solvent* indicates the immersion solvent of the tissue/cell.
- *File Type* indicates what type of file to save to.
- *Notes* is for any additional information the user wants to store with the file.

##### 4.5.2 Waveform Parameters

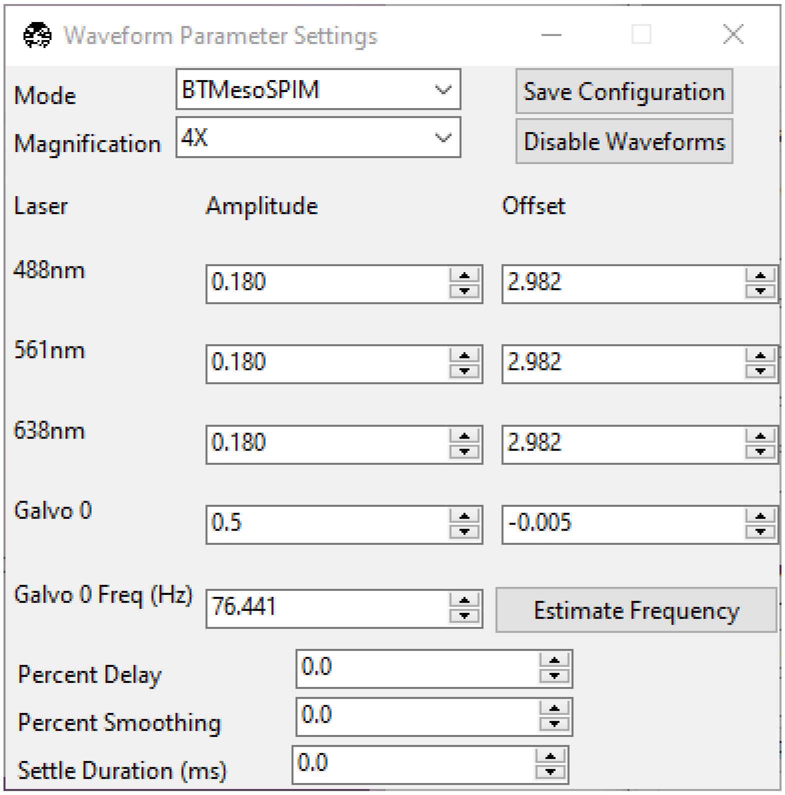

This is used to update the waveforms shown in *Waveform Settings*.

- For each laser, the *Amplitude* and *Offset* correspond to the amplitude and offset of the waveform sent to the remote focus device.
- For each galvo, the *Amplitude* and *Offset* correspond to the amplitude and offset of the waveform sent to the galvo, by default a triangle wave.
  - The *Galvo 0 Frequency (Hz)* sets the frequency of the waveform sent to the galvo. *Estimate Frequency* estimates the frequency needed for a sawtooth wave to sweep over the camera region of interest without aliasing with the light-sheet for a given rolling shutter size and speed (e.g., in a digitally scanned light-sheet format).
  - Additional galvos from the *configuration file* file are incrementally added here (e.g., *Galvo 1 Frequency (Hz)*, . . . ).
- *Percent Delay* introduces a delay before the remote focus waveform starts.
- *Percent Smoothing* smooths the remote focus waveform.
- *Settle Duration (ms)* introduces a delay after the remote focus sawtooth ends.

##### 4.5.3 Configure Microscopes

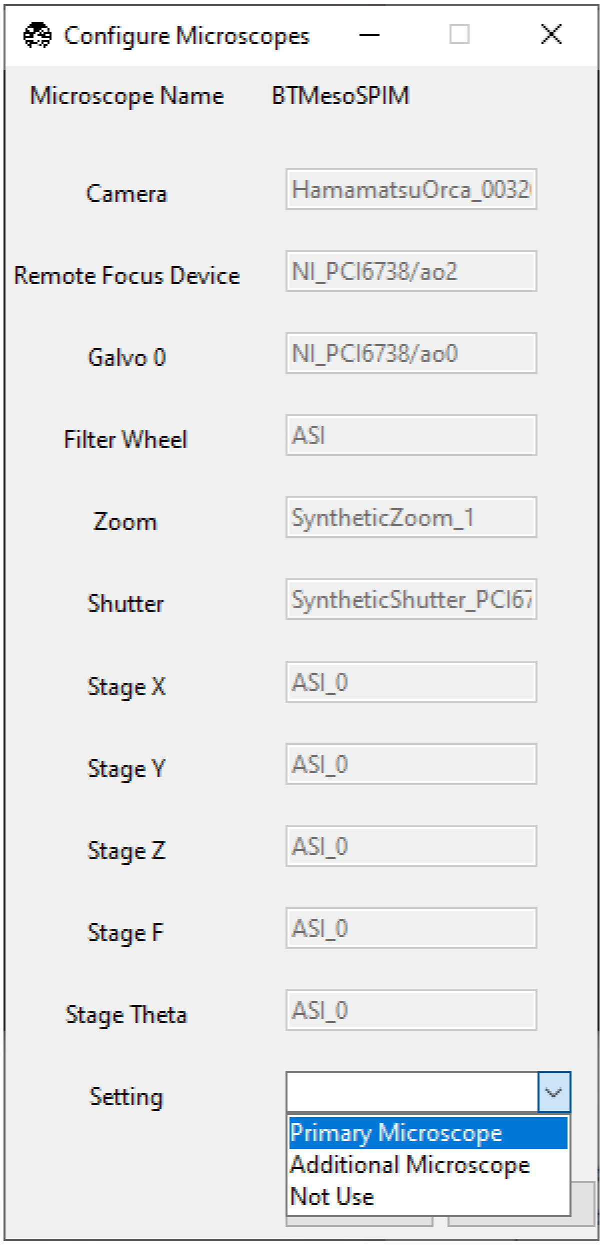

The *Configure Microscopes* window allows a user with multiple microscopes defined in their *configuration file* to select which microscope is primary and launch both microscopes simultaneously. The primary microscope will have control over any hardware shared between both microscopes. This window also provides a GUI interface to look at what hardware is in use.

##### 4.5.4 Multi-Position Tiling Wizard

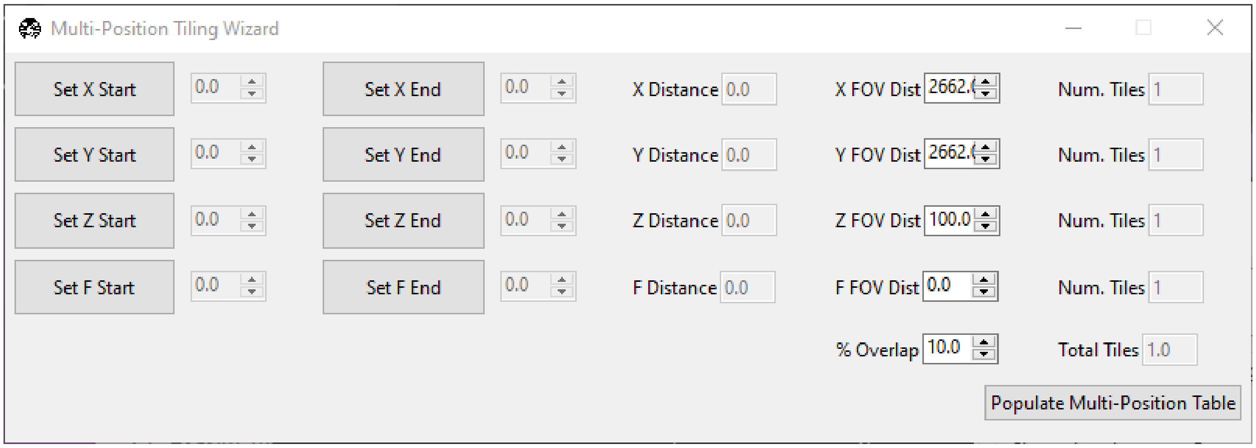

The tiling wizard helps the user set up a tiled acquisition of a sample large enough that it cannot be imaged in a single field of view.

- *Set <axis> Start* indicates the starting position of an axis.
- *Set <axis> End* indicates the end position of an axis.
- *<axis> Distance* indicates difference between the start and end position.
- *<axis> FOV Dist* indicates the field of view along that axis. The Distance between start and end will be split into tiles of this size along this axis.
- *Num. Tiles* indicates how many tiles exist along this axis. It is roughly (End - Start)/FOV dist.
- *% Overlap* indicates the percent of the the that should overlap along each axis. It is a percent of the FOV Dist.
- *Populate Multi-Position Table* puts all of the tiles in the *multi-position table*.

For an example of how to use the tiling wizard, see *Tiling a sample larger than the field of view*.

##### 4.5.5 Autofocus Settings

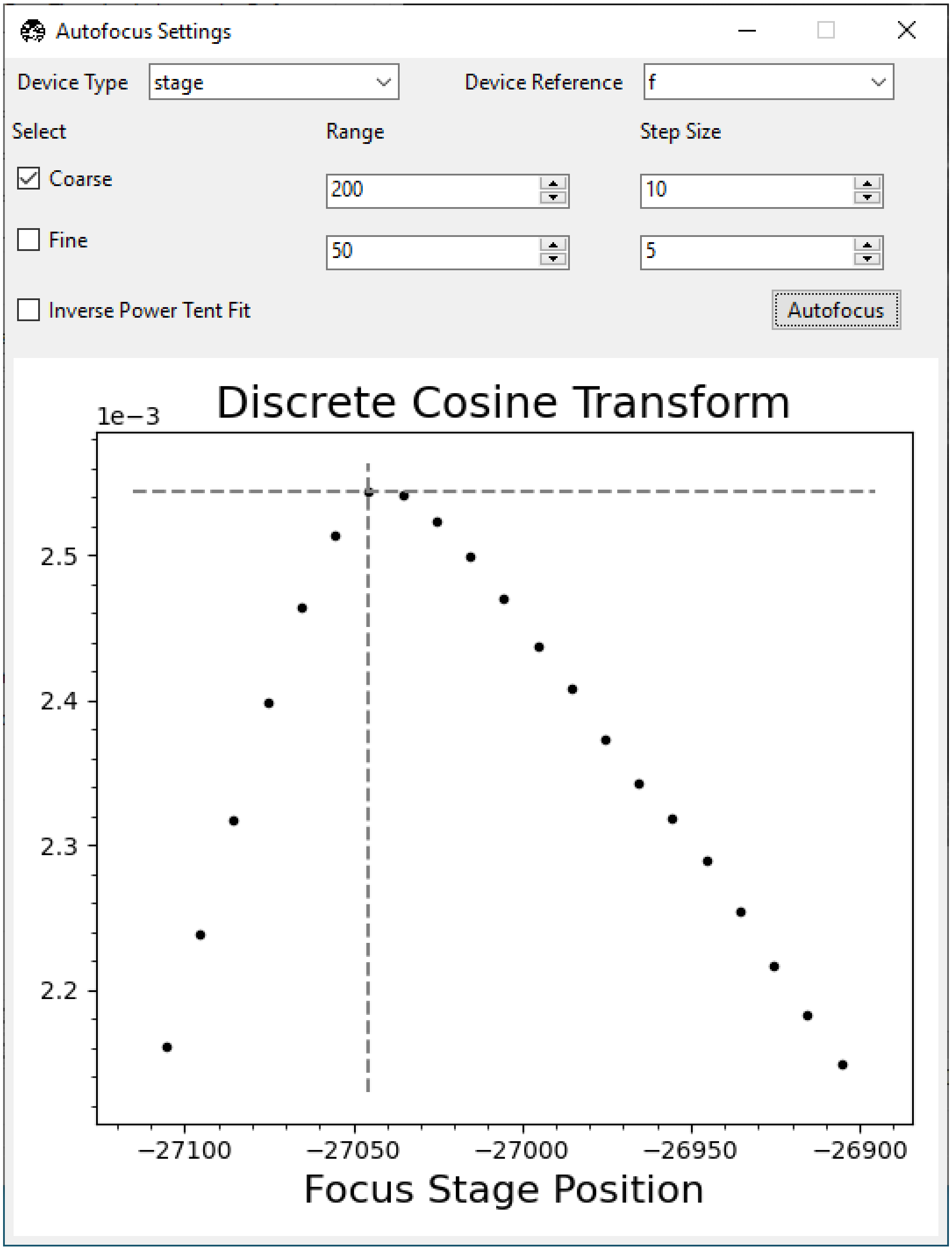

The *Autofocus Settings* panel controls parameters of the autofocus *feature*.

- *Device Type* indicates if the autofocus routine should be applied to a stage or to a remote focus device.
- *Device Reference* indicates the stage axis, or the DAQ analog output for the remote focus device, to use.
- The *Coarse* and *Fine* rows allow users to select a range and step size, both in microns (or volts, if using the remote focus device), over which the autofocus routine should search for an optimal focus value. If coarse and

fine are selected, the coarse search will be performed first and the fine search will be performed about the coarse position with the highest value.

- *Inverse Power Tent Fit* will attempt to find a more accurate position for the optimal focus based on fitting a power tent to the search values. It will only use the fit if its $R^2$ value is higher than `0.9`.
- *Autofocus* runs the autofocus with the set parameters.

Once the settings have been updated here, any run autofocus operation will use the new settings.

CHAPTER

FIVE

### SETTING UP A MICROSCOPE

#### 5.1 Configuration Overview

This section outlines the `configuration.yaml`, `experiment.yml`, `rest_api_config.yml`, `waveform_templates.yml`, and `waveform_constants.yml` files.

---

##### 5.1.1 Configuration File

In order for the **navigate** software to function, you will need to configure the specifications of the various hardware that you will be using. The first time you launch the software, **navigate** will create a copy of the `navigate\config\configuration.yaml` and the rest of the configuration files in `C:\Users\Username\AppData\Local\.navigate\config` on Windows or `~/.navigate` on Mac/Linux. **navigate** uses these local copies of the configuration files to store information specific to the setup attached to the computer it is installed on.

To avoid confusion, we recommend launching the software in the synthetic hardware mode initially. Within your Terminal, or Anaconda Prompt, activate your **navigate** Python environment and launch the software by typing: `navigate -sh`. Thereafter, you should only modify the `configuration.yaml` file in your local `.navigate\config` directory. The local copy avoids conflicts between different microscopes after pulling new changes from GitHub.

**Tip:** Once **navigate** is open in the synthetic hardware mode, you can open the `configuration.yaml` file by going to *File* menu and selecting *Open Configuration Files*.

It may help to open `C:\Users\Username\AppData\Local\.navigate\config\configuration.yaml` and follow along in this file when reading the next sections.

See the *Setting up an Axially Swept Light-Sheet Microscope* case study for a general walkthrough of how to build your own configuration file and see *Implementations* for examples of configuration files.

---

##### Hardware Section

The first section of the `configuration.yaml` file is called `hardware`. It contains all the necessary information to find and connect each hardware device to the computer/software.

Here is an example of what the section will look like:

```yaml
# Configuration in YAML
hardware:
    daq:
        type: NI
    camera:
        -
          type: HamamatsuOrca
          serial_number: 302158
        -
          type: HamamatsuOrca
          serial_number: 302352
    filter_wheel:
        type: SutterFilterWheel
        port: COM19
        baudrate: 9600
        number_of_wheels: 2
    stage:
        -
          type: PI
          controllername: 'C-884'
          stages: L-509.20DG10 L-509.40DG10 L-509.20DG10 M-060.DG M-406.4PD NOSTAGE
          refmode: FRF FRF FRF FRF FRF FRF
          serial_number: 119060508
        -
          type: Thorlabs
          serial_number: 74000375
    zoom:
        type: DynamixelZoom
        servo_id: 1
        port: COM18
        baudrate: 1000000
```

This example specifies that we are connected to

- A National Instruments DAQ (possibly multiple).
- Two Hamamatsu Orca (Flash or Fusion) cameras with different serial numbers for identification.
- A Sutter filter wheel controller connected via USB on port `COM19`. This control two filter wheels.
- A Physik Instrumente controller, identified by serial number, with access to 5 stages in `FRF` refrence mode (see PI's documentation).
- A Thorlabs stage (with a single axis), identified by a serial number.
- A Dynamixel zoom device connected via USB on port `COM18`.

Make sure that the `configuration.yaml` specifies the hardware on your microscope. For example, if you wanted to remove the Thorlabs stage and replace the PI stage with an ASI stage, the `stage` section would instead read:

```yaml
stage:
    type: ASI
    serial_number: 123456789
    port: COM7
    baudrate: 115200
```

Notice that since we are now using a single stage, we no longer have a - above the stage entry. The - indicates a list, and is only needed if we want to load multiple types of a single hardware.

---

**Note:** The type of the device is needed when deciding which Python object to instantiate on startup of the software (eg `type: ASI`). The other fields (eg `port: COM7`) change depending on the manufacturer's API. They help the API communicate with the computer you are using, which in turn allows the **navigate** software to communicate with the device.

---

Running the software with our current microscope setup would fail. It turns out our ASI stage only moves in the `X`, `Y`, `Z` and `F` axes. We need a way to handle the `Theta` axis. To address this, we will change our `stage` block of the YAML to also load a `SyntheticStage`:

```yaml
stage:
    -
        type: ASI
        serial_number: 123456789
        port: COM7
        baudrate: 115200
    -
        type: SyntheticStage
        serial_number: 987654321
```

If your microscope system does not have a device listed in the hardware section using the synthetic typing will allow the software to run without it. Another example would be replacing the zoom type with `SyntheticZoom` in the instance your microscope does not use that hardware. Your system will still run as you expect.

---

##### Microscope Section

The second section of `configuration.yaml` contains the microscope configurations that you will be using with the software. Each microscope is represented as a YAML dictionary, as in the hardware section. This section enables us to load one or more microscopes using the same hardware with varying combinations:

```yaml
microscopes:
    microscope1:
        ...
        ...
    microscope2:
        ...
        ...
```

Where `microscope1` and `microscope2` are names of two different microscopes using different combinations of the hardware listed in the `hardware` section. The names of the microscopes must not include spaces or special characters such as <, \, #, %, or ?.

Each microscope is expected to have a `daq`, `camera`, `remote_focus_device`, `galvo`, `filter_wheel`, `stage`, `zoom`, `shutter` and `lasers` section of the YAML dictionary. As in the hardware section, unused devices can be specified as synthetic.

Most of the information to set up these devices can be found in the *Supported Hardware* section of the documentation. Additional explanations of a few specific sections of the microscope configuration are below. Notably, the `zoom` section of the `configuration.yaml` specifies effective pixel size.

---

##### Stage Subsection

The stage section of the microscope 1) puts the stage control from the `hardware` section into the microscope 2) sets boundaries for stage movement and 3) optionally specifies joystick-controlled axes.

```yaml
microscopes:
    microscope1:
        stage:
        hardware:
            -
                name: stage
                type: ASI
                serial_number: 123456789
                axes: [x, y, z, f] # Software
                axes_mapping: [M, Y, X, Z] # M Shear axis mapping

            -
                name: stage
                type: SyntheticStage
                serial_number: 987654321
                axes: [theta]

        joystick_axes: [x, y, z]
        x_max: 100000
        x_min: -100000
        y_max: 100000
        y_min: -100000
        z_max: 100000
        z_min: -100000
        f_max: 100000
        f_min: -100000
        theta_max: 360
        theta_min: 0

        x_offset: 0
        y_offset: 0
        z_offset: 0
        theta_offset: 0
        f_offset: 0
```

First, we set the axes controlled by each piece of hardware and a mapping from the hardware's API axes to our software's axes. For example, the ASI `M` axis is mapped onto our software's `X` axis below.

As you may recall from the *Hardware Section*, we needed to add the `SyntheticStage` to control `Theta`. We now specify in the microscope that `Theta` is controlled by the synthetic stage in the `hardware` section of `microscope1`.

Below this, we specify that only `X`, `Y` and `Z` axes may be controlled by a joystick and we set the stage bounds for each of the axes.

Finally, we set the offset for each stage axis. This is an offset relative to other microscopes (e.g. `microscope2`) specified in `configuration.yaml`. In this case, `microscope1` is the reference microscope. Additional microscopes may ask the stage to move to a different offset in order to observe the sample at the same position as `microscope1`.

---

##### Stage Axes Definition

Many times, the coordinate system of the stage hardware do not agree with the optical definition of each axes identity. For example, many stages define their vertical dimension as `Z`, whereas optically, we often define this axis as `X`. Thus, there is often a need to map the mechanical axes to the optical axes, and this is done with the `axes_mapping` dictionary entry in the stage hardware section. By default, stage axes are read in as `X`, `Y`, `Z`, `Theta`, `F`, where `Theta` is rotation and `F` is focus, but this can be changed by changing axes mapping.

```
axes: [x, y, z, theta, f]
axes_mapping: [x, y, z, r, f]
```

If, on a certain microscope, the `Z` stage axis corresponds to the optical y-axis, and vice versa, you would then have to import the stages as following:

```
axes: [x, y, z, theta, f]
axes_mapping: [x, z, y, r, f]
```

---

##### Joystick Axes Definition

If you are using a joystick, it is possible to disable GUI control of the stage axes that the joystick can interact with. The axes that the joystick can interact with appear in the stage field as following:

```
joystick_axes: [x, y, z]
```

---

**Note:** These axes should agree with the optical axes. If, on the same microscope as mentioned in the *Stage Axes Definition* section, the joystick were to control the optical y-axis corresponding to the stage z axis, you would have to put `Y` in the joystick axes brackets as following:

---

```
joystick_axes: [y]
```

---

##### Zoom Subsection

The `zoom` section of `configuration.yaml` specifies control over microscope zoom lenses. For example, we use the Dynamixel Smart Actuator to control the rotating zoom wheel on an Olympus MVXPLAPO 1x/0.25.

```yaml
microscopes:
    microscope1:
        zoom:
            hardware:
                name: zoom
                type: DynamixelZoom
                servo_id: 1
            position:
                0.63x: 0
                1x: 627
                2x: 1711
                3x: 2301
                4x: 2710
                5x: 3079
                6x: 3383
            pixel_size:
                0.63x: 9.7
                1x: 6.38
                2x: 3.14
                3x: 2.12
                4x: 1.609
                5x: 1.255
                6x: 1.044
            stage_positions:
                BABB:
                    f:
                        0.63x: 0
                        1x: 1
                        2x: 2
                        3x: 3
                        4x: 4
                        5x: 5
                        6x: 6
```

The `hardware` section connects to the zoom hardware. The `positions` specify the voltage of the actuator at different zoom positions. The `pixel_size` specifies the effective pixel size of the system at each zoom. The `stage_positions` account for focal shifts in between the different zoom values (the MVXPLAPO does not have a consistent focal plane). These may change depending on the immersion media. Here it is specified for a `BABB` (Benzyl Alcohol Benzyl Benzoate) immersion media.

Regardless of whether or not your microscope uses a zoom device, you must have a `zoom` entry, indicating the effective pixel size of your system in micrometers. For example,

```yaml
zoom:
  hardware:
    name: zoom
    type: SyntheticZoom
    servo_id: 1
  position:
```

(continues on next page)

(continued from previous page)

```
  N/A: 0
pixel_size:
  N/A: 0.168
```

##### GUI Section

The third and final section of the `configuration.yaml` file is the GUI parameters.

It will look something like the below:

```
gui:
    channels:
        count: 5
        laser_power:
            min: 0
            max: 100
            step: 10
        exposure_time:
            min: 1
            max: 1000
            step: 5
        interval_time:
            min: 0
            max: 1000
            step: 5
    stack_acquisition:
        step_size:
            min: 0.200
            max: 1000
            step: 0.1
        start_pos:
            min: -5000
            max: 5000
            step: 1
        end_pos:
            min: -5000
            max: 10000
            step: 1
    timepoint:
        timepoints:
            min: 1
            max: 1000
            step: 1
        stack_pause:
            min: 0
            max: 1000
            step: 1
```

The values in each field relate to GUI widgets.

- The `channels` section indicates GUI settings for the channel settings under *Channels*, *Channel Settings*.

- *count* specifies how many channels should be displayed.
- *laser_power*, *exposure_time* and *interval_time* are used to set the minimum, maximum and step size values for *Power*, *Exp. Time (ms)* and *Interval*, respectively.

- The `stack_acquisition` section indicates GUI settings for the stack acquisition settings under *Channels*, *Stack Acquisition Settings (um)*.
  - *step_size*, *start_pos* and *end_pos* are used to set the minimum, maximum and step size values for *Step Size*, *Start* and *End*, respectively.
- The `timepoint` section indicates GUI settings for the timepoint settings under *Channels*, *Timepoint Settings*.
  - *timepoints* and *stack_pause* are used to set the minimum, maximum and step size values for *Timepoints* and *Stack Pause (s)*, respectively.

**Note:** This section is still under development. The plan going forward is to have all widgets be controlled in this manner.

##### 5.1.2 Experiment File

The `experiment.yml` file stores information about the current state of the program. This includes laser and camera parameters, saving options, z-stack settings and much more. This file does not need to be edited by the user. The program will update it automatically and save changes automatically on exit.

##### 5.1.3 Waveform Constants File

The `waveform_constants.yml` file stores the waveform parameters that can be edited by going to *Microscope Configuration* → *Waveform Parameters*. This file does not need to be edited by the user. The program will update it automatically and save changes automatically on exit.

##### 5.1.4 Waveform Templates File

The waveform templates file stores default behavior for the number of repeats for specific waveforms. This file only needs to be edited if the user wishes to introduce a new waveform behavior to the application.

##### 5.1.5 Rest API Configuration File

The REST API configuration file specifies where the REST API should look to get and post data. This is only needed if you are using a plugin that requires the REST API, such as our communication with ilastik.

#### 5.2 Hardware

##### 5.2.1 Supported Hardware

###### Data Acquisition Card

Data acquisition cards deliver and receive analog and digital signals. To acquire an image, the software calculates all of the analog and digital waveforms and queues these waveforms on the data acquisition card. Upon receipt of a trigger (either from the software itself, or an external piece of hardware), all of the analog and digital signals are delivered in parallel. This provides deterministic behavior on a per-frame basis, which is necessary for proper acquisition of light-sheet data. It does not however provide us with deterministic behavior between image frames, and some jitter in timing is anticipated.

---

###### National Instruments (NI)

In principle, most NI-based data acquisition cards should work with the software, so long as there are a sufficient number of analog and digital ports, and the sampling rate (typically 100 kHz) is high enough per port.

Prior to installing the card within the computer, first install the NI-DAQmx drivers. Once installed, connect the PCIe or PXIe-based device to the computer. A functioning system should be recognized by the operating system, and visible in the Windows Device Manager as a **NI Data Acquisition Device**.

---

**Tip: The most important aspect is to wire up the breakout box properly.**

To find the device pin outs for your NI-based data acquisition card, open NI MAX, find the card under devices, right-click and select "device pinouts".

Important: Should you use the SCB-68A breakout box, do not look at the pinout on the back of the cover. This is misleading. You must look at the device pinouts in NI MAX.

---

###### Wiring

- Identify the device name in NI MAX, and change it if you would like. Common names are `Dev1`, `Dev2`, etc. This name must correspond with the pinouts provided in the configuration file.
- Connect the `master_trigger_out_line` to the `trigger_source` with a direct wire, commonly `PXI6259/port0/line1` and `/PXI6259/PFI0`. In this example, the default name for the device (e.g., `Dev1`) has been changed to `PXI6259`.
- Connect the `camera_trigger_out_line` to the `Ext. Trigger` on the camera using the `CTR0Out` pin.
- These values must precisely match those in the configuration file. An example is provided below:

```yaml
hardware:
  daq:
    type: NI

microscopes:
  microscope_name:
    daq:
```

(continues on next page)

(continued from previous page)

```
    hardware:
      name: daq
      type: NI
    sample_rate: 100000
    sweep_time: 0.2
    master_trigger_out_line: PCI6738/port0/line1
    camera_trigger_out_line: /PCI6738/ctr0
    trigger_source: /PCI6738/PFI0
    laser_port_switcher: PCI6738/port0/line0
    laser_switch_state: False
```

**Note:** For NI-based cards, `port0/line1` is the equivalent of `P0.1`. There are multiple pins for each PFIO, including source, out, gate, etc. You must use the out terminal.

The software has been tested with the following NI-based cards:

##### PCIe/PXIe-6738

The PCIe-6738 can only create one software-timed analog task for every four channels. As such, the lasers much be attached to analog output ports outside of the banks (shown as solid lines in the device pinout) used by the galvo/remote focus units. For example, if you use ao0, ao2, and ao6 for the remote focus, galvo, and galvo stage, the lasers should be connected to ao8, ao9, and ao10. In such a configuration, they will not compete with the other analog output ports. Since only one task will be created created on the ao8, ao9, ao10 bank at a time (only one laser is on at a time), only one laser can be on at a time. If we wanted to turn lasers on simultaneously, we could distribute the lasers across independent banks (e.g. ao8, ao14, ao19).

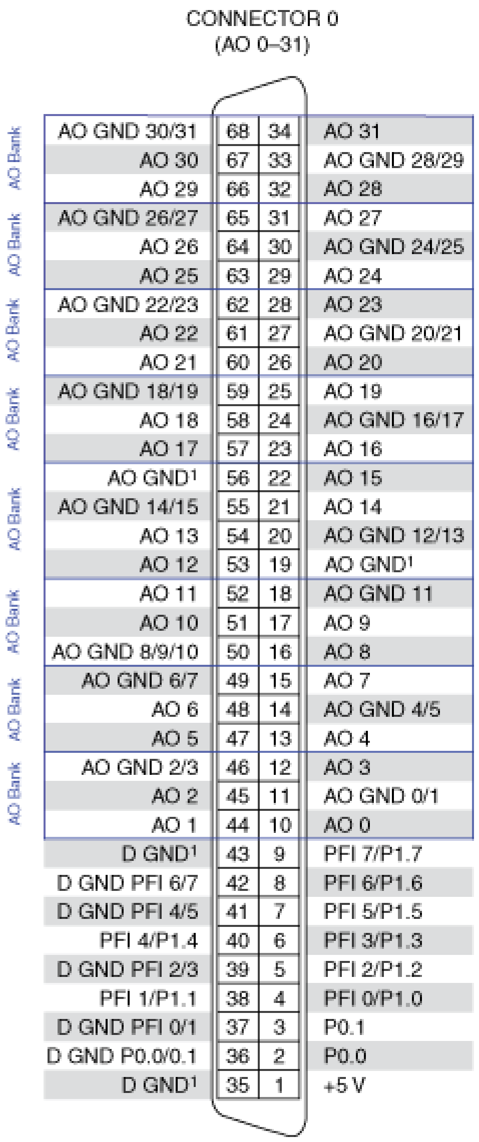

[1] No connect when using the SHC68-68-A2 cable.

#### PCIe/PXIe-6259

The PXI-6259 can create one software-timed analog task per channel. As such, the galvo/remote focus/lasers can be attached to any of the analog output ports. The 6259 has two connectors, and it is important to make sure that the analog and digital ports that you are using are connected to the correct connector. For example, if you are using `ao0`, this is located on `connector 0`.

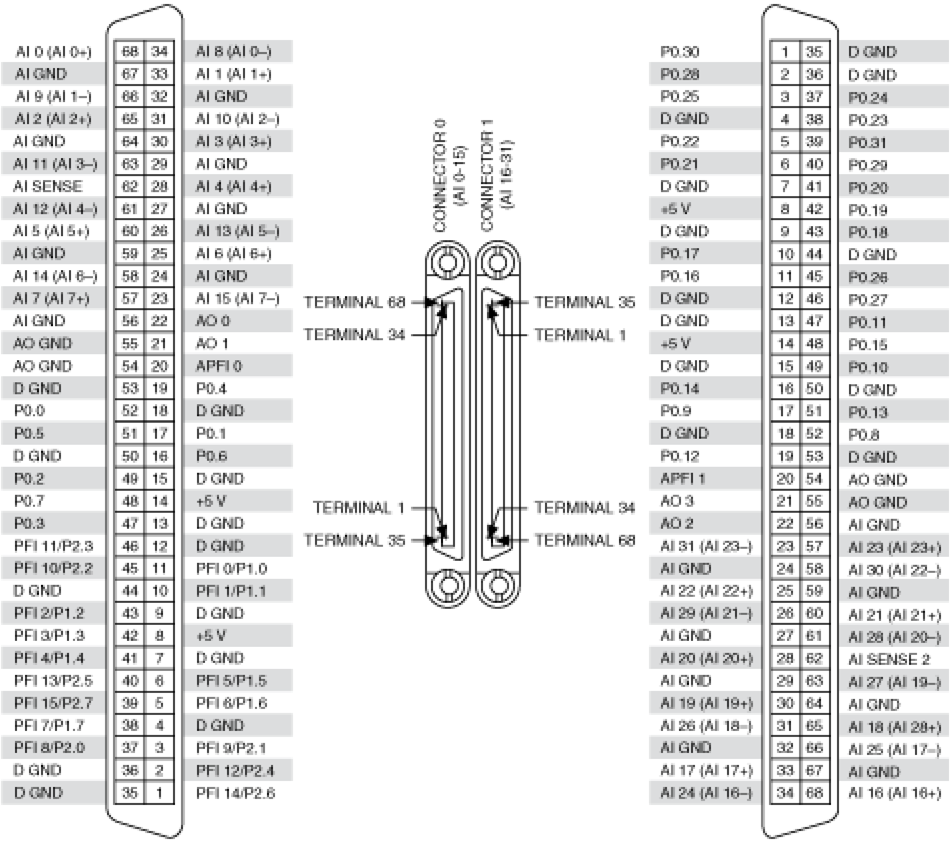

#### PCIe/PXIe-6723

The PXI-6723 can also create one software-timed analog task per channel. As such, the analog outputs can be wired up as is most convenient.

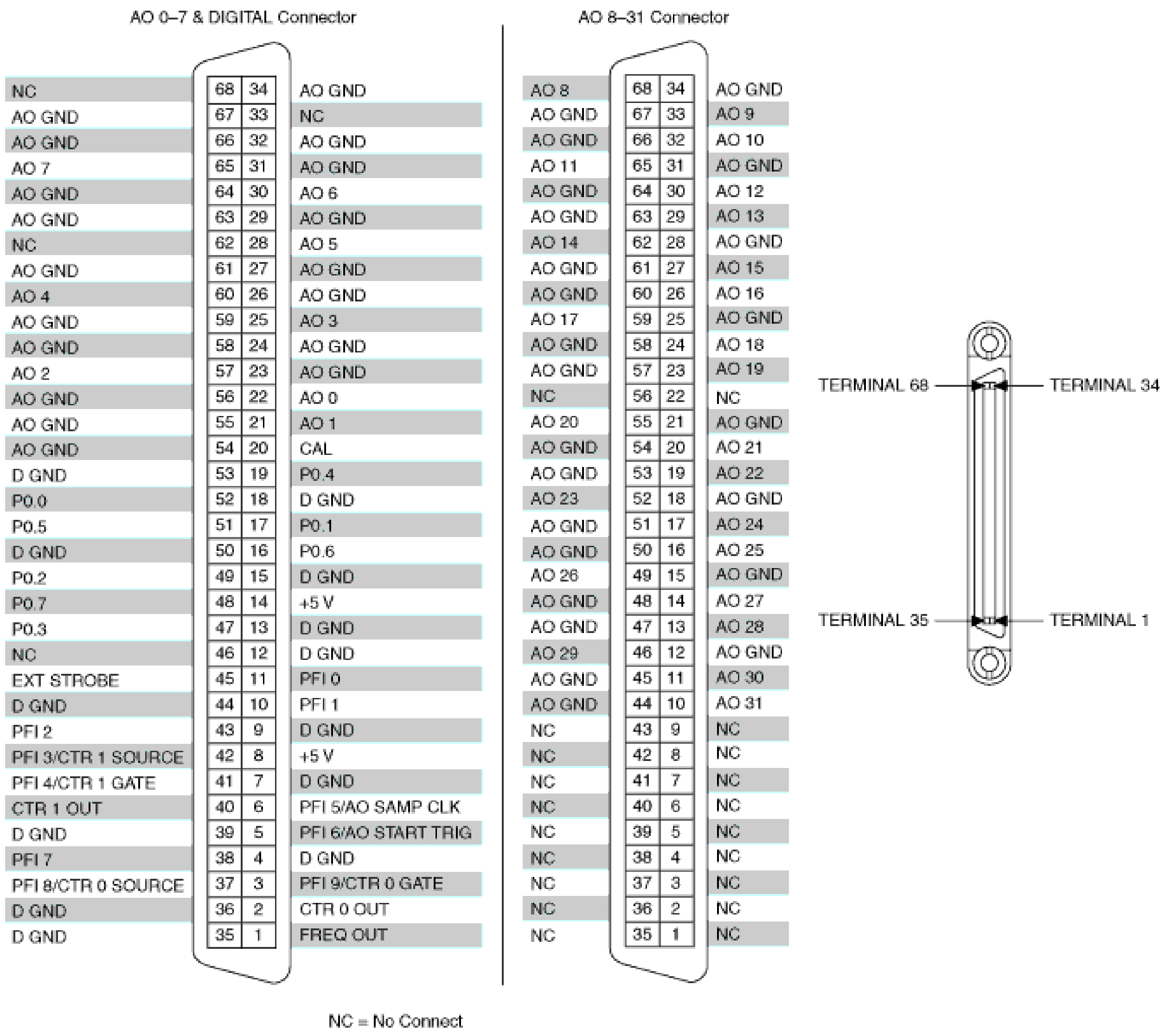

##### Synthetic Data Acquisition Card

If no data acquisition card is present, one must configure the software to use a synthetic data acquisition card.

```yaml
hardware:
  daq:
    type: SyntheticDAQ

microscopes:
  microscope_name:
    daq:
      hardware:
        name: daq
        type: SyntheticDAQ
      sample_rate: 100000
      sweep_time: 0.2
      master_trigger_out_line: PCI6738/port0/line1
```

(continues on next page)

(continued from previous page)

```
      camera_trigger_out_line: /PCI6738/ctr0
      trigger_source: /PCI6738/PFI0
      laser_port_switcher: PCI6738/port0/line0
      laser_switch_state: False
```

---

##### Cameras

The software supports camera-based acquisition. It can run both normal and rolling shutter modes of contemporary scientific CMOS cameras.

###### Hamamatsu ORCA-Flash4.0 V3/Fusion

- Insert the USB that came with the camera into the computer and install HCImageLive. Alternatively, download DCAM-API. The software can be found here.
- When prompted with the DCAM-API Setup
  - If you are going to use the Frame Grabber, install the Active Silicon Firebird drivers.
  - Select ... next to the tools button, and install DCAM tools onto the computer.
- Shutdown the computer and install the Hamamatsu frame grabber into an appropriate PCIe-x16 slot on the motherboard.
- Turn on the computer and the camera, and confirm that it is functioning properly in HCImageLive or Excap (one of the DCAM tools installed)
- Connect the *camera_trigger_out_line* to the External Trigger of the Hamamatsu Camera. Commonly, this is done with a counter port, e.g., `/PXI6259/ctr0`

More about the ORCA-Flash 4.0 v3 and ORCA-Fusion can be found here and here, respectively.

```
hardware:
  camera:
    -
      type: HamamatsuOrca
      serial_number: 302153

microscopes:
  microscope_name:
    camera:
      hardware:
        name: camera
        type: HamamatsuOrca
        serial_number: 302153
      x_pixels: 2048.0
      y_pixels: 2048.0
```

(continues on next page)

(continued from previous page)

```
      flip_x: True
      flip_y: False
      pixel_size_in_microns: 6.5
      subsampling: [1, 2, 4]
      sensor_mode: Normal
      readout_direction: Top-to-Bottom
      lightsheet_rolling_shutter_width: 608
      defect_correct_mode: 1.0
      binning: 1x1
      readout_speed: 2.0
      trigger_active: 1.0
      trigger_mode: 1.0
      trigger_polarity: 2.0
      trigger_source: 2.0
      exposure_time: 20
      delay_percent: 20
      pulse_percent: 1
      line_interval: 0.000075
      display_acquisition_subsampling: 4
      average_frame_rate: 4.969
      frames_to_average: 1
      exposure_time_range:
        min: 1
        max: 1000
        step: 1
      x_pixels_step: 4
      y_pixels_step: 4
      x_pixels_min: 4
      y_pixels_min: 4
```

##### Hamamatsu ORCA-Lightning

The Hamamatsu ORCA-Lightning has a slightly different class than the Flash/Fusion as it reads out 4 rows at a time rather than 1 in rolling shutter mode. Learn more here.

```
hardware:
  camera:
    -
      type:  HamamatsuOrcaLightning
      serial_number: 000035

microscopes:
  microscope_name:
    camera:
      hardware:
        name: camera
        type: HamamatsuOrcaLightning
```

(continues on next page)

(continued from previous page)

```
        serial_number: 000035
      x_pixels: 4608.0
      y_pixels: 2592.0
      pixel_size_in_microns: 5.5
      subsampling: [1, 2, 4]
      sensor_mode: Normal
      readout_direction: Bottom-to-Top
      lightsheet_rolling_shutter_width: 608
      defect_correct_mode: 2.0
      binning: 1x1
      readout_speed: 0x7FFFFFFF
      trigger_active: 1.0
      trigger_mode: 1.0
      trigger_polarity: 2.0
      trigger_source: 2.0
      exposure_time: 20
      delay_percent: 8
      pulse_percent: 1
      line_interval: 0.000075
      display_acquisition_subsampling: 4
      average_frame_rate: 4.969
      frames_to_average: 1
      exposure_time_range:
        min: 1
        max: 1000
        step: 1
```

##### Hamamatsu ORCA-Fire

The Hamamatsu ORCA-Fire is one of the latest releases from Hamamatsu. It is a scientific CMOS camera that offers large 10.5 MPix sensor and greater diversity of rolling shutter readout modes. Learn more here.

```
hardware:
  camera:
    -
      type:  HamamatsuOrcaFire
      serial_number: 000035

microscopes:
  microscope_name:
    camera:
      hardware:
        name: camera
        type: HamamatsuOrcaFire
        serial_number: 000035
      x_pixels: 4432.0
      y_pixels: 2368.0
```

(continues on next page)

(continued from previous page)

```
      pixel_size_in_microns: 4.6
      subsampling: [1, 2, 4]
      sensor_mode: Normal
      readout_direction: Bottom-to-Top
      lightsheet_rolling_shutter_width: 608
      defect_correct_mode: 2.0
      binning: 1x1
      readout_speed: 0x7FFFFFFF
      trigger_active: 1.0
      trigger_mode: 1.0
      trigger_polarity: 2.0
      trigger_source: 2.0
      exposure_time: 20
      delay_percent: 8
      pulse_percent: 1
      line_interval: 0.000075
      display_acquisition_subsampling: 4
      average_frame_rate: 4.969
      frames_to_average: 1
      exposure_time_range:
        min: 1
        max: 1000
        step: 1
```

###### Photometrics Iris 15

- Download the PVCAM software from Photometrics. The PVCAM SDK is also available form this location. You will likely have to register and agree to Photometrics terms.
- Perform the Full Installation of the PVCAM software.
- Should a "Base Device" still show up as unknown in the Windows Device Manager, you may need to install the Broadcom PCI/PCIe Software Development Kit
- Upon successful installation, one should be able to acquire images with the manufacturer-provided PVCamTest software.

```
camera:
  type: Photometrics
  camera_connection: PMPCIECam00
  serial_number: 1

camera:
    hardware:
      name: camera
      type: Photometrics
      serial_number: 1
    x_pixels: 5056.0
    y_pixels: 2960.0
```

(continues on next page)

(continued from previous page)

```
    pixel_size_in_microns: 4.25
    subsampling: [1, 2, 4]
    sensor_mode: Normal
    readout_direction: Bottom-to-Top
    lightsheet_rolling_shutter_width: 608
    defect_correct_mode: 2.0
    binning: 1x1
    readout_speed: 0x7FFFFFFF
    trigger_active: 1.0
    trigger_mode: 1.0
    trigger_polarity: 2.0
    trigger_source: 2.0
    exposure_time: 20
    delay_percent: 25
    pulse_percent: 1
    line_interval: 0.000075
    display_acquisition_subsampling: 4
    average_frame_rate: 4.969
    frames_to_average: 1
    exposure_time_range:
      min: 1
      max: 1000
      step: 1
```

##### Synthetic Camera

The synthetic camera simulates noise images from an sCMOS camera. If no camera is present, the synthetic camera class must be used.

```
camera:
  type: SyntheticCamera
  serial_number: 12345

 microscopes:
  microscope_name:
    camera:
      hardware:
        name: camera
        type: SyntheticCamera
        serial_number: 12345
      x_pixels: 2048.0
      y_pixels: 2048.0
      flip_x: True
      flip_y: False
      pixel_size_in_microns: 6.5
      subsampling: [1, 2, 4]
      sensor_mode: Normal
```

(continues on next page)

(continued from previous page)

```yaml
      readout_direction: Top-to-Bottom
      lightsheet_rolling_shutter_width: 608
      defect_correct_mode: 1.0
      binning: 1x1
      readout_speed: 2.0
      trigger_active: 1.0
      trigger_mode: 1.0
      trigger_polarity: 2.0
      trigger_source: 2.0
      exposure_time: 20
      delay_percent: 20
      pulse_percent: 1
      line_interval: 0.000075
      display_acquisition_subsampling: 4
      average_frame_rate: 4.969
      frames_to_average: 1
      exposure_time_range:
        min: 1
        max: 1000
        step: 1
      x_pixels_step: 4
      y_pixels_step: 4
      x_pixels_min: 4
      y_pixels_min: 4
```

---

##### Remote Focusing Devices

Voice coils, also known as linear actuators, play a crucial role in implementing aberration-free remote focusing in navigate. These electromagnetic actuators are used to control the axial position of the light-sheet and the sample relative to the microscope objective lens. By precisely adjusting the axial position, the focal plane can be shifted without moving the objective lens, thus enabling remote focusing. Focus tunable lenses serve as an alternative to voice coils owing to their simple operation and high bandwidth. Tunable lenses axially scan a beam by introducing defocus into the optical train. Nonetheless, they do not provide the higher-order correction provided by voice coils in an aberration-free remote focusing system.

##### Equipment Solutions

Configuration of the device can be variable. Many voice coils we have received require establishing serial communication with the device to explicitly place it in an analog control mode. In this case, the comport must be specified properly in the configuration file.

More recently, Equipment Solutions has begun delivering devices that automatically initialize in an analog control mode, and thus no longer need the serial communication to be established. For these devices, we recommend using the analog control mode described in the next section.

The LFA-2010 Linear Focus Actuator is controlled with a SCA814 Linear Servo Controller, which accepts a +/- 2.5 Volt analog signal. The minimum and maximum voltages can be set in the configuration file to prevent the device from receiving a voltage outside of its operating range.

```yaml
microscopes:
  microscope_name:
    remote_focus_device:
      hardware:
        name: remote_focus
        type: EquipmentSolutions
        channel: PCI6738/ao2
        comport: COM7
        min: -2.5
        max: 2.5
      delay_percent: 7.5
      ramp_rising_percent: 85
      ramp_falling_percent: 5.0
      amplitude: 0.7
      offset: 2.3
      smoothing: 0.0
```

##### Analog Controlled Voice Coils and Tunable Lenses

In principle, this hardware type can support any analog-controlled voice coil or tunable lens. The BLINK and the Optotune Focus Tunable Lens are controlled with an analog signal from the DAQ. The BLINK is a voice coil that is pneumatically actuated voice coil. it is recommended that you specify the min and max voltages that are compatible with your device to prevent the device from receiving a voltage outside of its operating range.

```yaml
microscopes:
  microscope_name:
    remote_focus_device:
      hardware:
        name: remote_focus
        type: NI
        channel: PCI6738/ao2
        comport: COM7
        min: -2.5
        max: 2.5
      delay_percent: 7.5
      ramp_rising_percent: 85
      ramp_falling_percent: 5.0
      amplitude: 0.7
      offset: 2.3
      smoothing: 0.0
```

##### Synthetic Remote Focus Device

If no remote focus device is present, one must configure the software to use a synthetic remote focus device.

```yaml
microscopes:
  microscope_name:
    remote_focus_device:
      hardware:
        name: remote_focus
        type: SyntheticRemoteFocus
        channel: PCI6738/ao2
        comport: COM7
        min: -2.5
        max: 2.5
      delay_percent: 7.5
      ramp_rising_percent: 85
      ramp_falling_percent: 5.0
      amplitude: 0.7
      offset: 2.3
      smoothing: 0.0
```

---

##### Stages

Our software empowers users with a flexible solution for configuring multiple stages, catering to diverse microscope modalities. Each stage can be customized to suit the specific requirements of a particular modality or shared across various modalities. Our unique approach allows seamless integration of stages from different manufacturers, enabling users to mix and match components for a truly versatile and optimized setup tailored to their research needs.

**Note:** The software provides configure specific hardware axes to software axes. This is specified in the configuration file. For example, if specified as follows, the software x, y, z, and f axes can be mapped to the hardware axes M, Y, X, and Z, respectively.

```yaml
axes: [x, y, z, f]
axes_mapping: [M, Y, X, Z]
```

##### ASI Tiger Controller

The ASI Tiger Controller. is a multi-purpose controller for ASI stages, filter wheels, dichroic sliders, and more. We communicate with Tiger Controllers via a serial port. It is recommended that you first establish communication with the device using ASI provided software. For stages in particular, there is a `feedback_alignment` configuration option option corresponds to the Tiger Controller AA Command.

**Tip:** If you are using the FTP-2000 stage, do not change the F stage axis. This will differentially drive the two vertical posts, causing them to torque and potentially damage one another.

```yaml
hardware:
  stage:
    type: ASI
    serial_number: 123456789
    port: COM8
    baudrate: 115200

microscopes:
  microscope:
    stage:
      hardware:
        name: stage
        type: ASI
        serial_number: 123456789
        axes: [x, y, z, f]
        axes_mapping: [M, Y, X, Z]
        feedback_alignment: [90, 90, 90, 90]
```

##### Sutter MP-285

The Sutter MP-285 communicates via serial port and is quite particular. We have done our best to ensure the communication is stable, but occasionally the stage will send or receive an extra character, throwing off communication. In this case, the MP-285's screen will be covered in 0s, 1s or look garbled. If this happens, simply turn off the software, power cycle the stage, and press the "MOVE" button on the MP-285 controller once. When the software is restarted, it should work.

**Tip:** Sometimes the Coherent Connection software messes with the MP-285 serial communication if it is connected to the lasers.

```yaml
hardware:
  stage:
  -
    type: MP285
    port: COM2
    timeout: 0.25
```

(continues on next page)

(continued from previous page)

```
    baudrate: 9600
    serial_number: 0000
    stages: None

microscopes:
  microscope_name:
    stage:
      hardware:
        name: stage1
        type: MP285
        serial_number: 0000
        axes: [y, x, f]
        axes_mapping: [z, y, x]
        volts_per_micron: None
        axes_channels: None
        max: 25000
        min: 0
```

##### Physik Instrumente

These stages are controlled by PI's own Python code and are quite stable. They include a special `hardware` option, `refmode`, which corresponds to how the PI stage chooses to self-reference. Options are `REF`, `FRF`, `MNL`, `FNL`, `MPL` or `FPL`. These are PI's GCS commands, and the correct reference mode for your stage should be found by launching PIMikroMove, which should come with your stage. Stage names (e.g. `L-509.20DG10`) can also be found in PIMikroMove or on a label on the side of your stage.

**Note:** PI L-509.20DG10 has a unidirectional repeatability of 100 nm, bidirectional repeatability of 2 microns, and a minimum incremental motion of 100 nm. This is potentially too coarse.

```
hardware:
  stage:
    -
      type: PI
      controllername: C-884
      stages: L-509.20DG10 L-509.40DG10 L-509.20DG10 M-060.DG M-406.4PD NOSTAGE
      refmode: FRF FRF FRF FRF FRF FRF
      serial_number: 119060508
    -
microscopes:
  microscope_name:
    stage:
      hardware:
        name: stage
        type: PI
        serial_number: 119060508
        axes: [x, y, z, theta, f]
```

(continues on next page)

(continued from previous page)

```
    y_unload_position: 10000
    y_load_position: 90000

    startfocus: 75000
    x_max: 100000
    x_min: -100000
    y_max: 100000
    y_min: -100000
    z_max: 100000
    z_min: -100000
    f_max: 100000
    f_min: 0
    theta_max: 360
    theta_min: 0

    x_rot_position: 2000
    y_rot_position: 2000
    z_rot_position: 2000

    x_step: 500
    y_step: 500
    z_step: 500
    theta_step: 30
    f_step: 500

    position:
      x_pos: 25250
      y_pos: 40000
      z_pos: 40000
      f_pos: 70000
      theta_pos: 0
    velocity: 1000

    x_offset: 0
    y_offset: 0
    z_offset: 0
    f_offset: 0
    theta_offset: 0
```

##### Thorlabs

We currently support the KIM001 controller. Importantly, this device shows significant hysterisis, and thus we do not recommend it for precise positioning tasks (e.g., autofocusing). It serves as a cost-effective solution for manual, user-driven positioning.

```yaml
hardware:
  stage:
    -
      type: Thorlabs
      serial_number: 74000375

microscopes:
  microscope_name:
    stage:
        hardware:
          -
            name: stage
            type: Thorlabs
            serial_number: 74000375
            axes: [f]
            axes_mapping: [1]
            volts_per_micron: None
            axes_channels: None
            max: None
            min: None
```

##### Analog-Controlled Galvo/Piezo

We sometimes control position via a galvo or piezo with no software API. In this case, we treat a standard galvo mirror or piezo as a stage axis. We control the "stage" via voltages sent to the galvo or piezo. The `volts_per_micron` setting allows the user to pass an equation that converts position in microns `X`, which is passed from the software stage controls, to a voltage. Note that we use `GalvoNIStage` whether or not the device is a galvo or a piezo since the logic is identical. The voltage signal is delivered via the data acquisition card specified in the `axes_mapping` entry.

**Note:** The parameters `distance_threshold` and `settle_duration_ms` are used to provide a settle time for large moves. if the move is larger than the `distance_threshold`, then a wait duration of `settle_duration_ms` is used to allow the stage to settle before the image is acquired.

```yaml
hardware:
  stage:
  -
    type: GalvoNIStage
    port: COM9999
    timeout: 0.25
    baudrate: 9600
    serial_number: 0000
```

(continues on next page)

(continued from previous page)

```
    stages: None
    distance_threshold: 20
    settle_duration_ms: 5

microscopes:
  microscope_name:
    stage:
      hardware:
          name: stage3
          type: GalvoNIStage
          serial_number: 0000
          axes: [z]
          axes_mapping: [PCI6738/ao6]
          volts_per_micron: 0.05*x
          max: 10
          min: 0
          distance_threshold: 5
          settle_duration_ms: 5
```

##### Synthetic Stage

If no stage is present for a particular axis, one must configure the software to use a synthetic stage. For example, not all microscopes have a theta axis.

```
hardware:
  stage:
    -
      type: SyntheticStage
      serial_number: 74000375

microscopes:
  microscope_name:
    stage:
        hardware:
          -
            name: stage
            type: SyntheticStage
            serial_number: 74000375
            axes: [theta]
            axes_mapping: [purple]
            volts_per_micron: None
            axes_channels: None
            max: None
            min: None
```

##### Filter Wheels

Filter wheels can be used in both illumination and detection paths. Dichroic turrets are controlled via the same code as filter wheels. The user is expected to change the names of available filters to match what is in the filter wheel or turret.

###### Sutter

We typically communicate with Sutter Lambda 10-3 controllers via serial port. It is recommended that you first establish communication with the device using manufacturer provided software. Alternatively, one can use MicroManager. For some filter wheel types, the filter_wheel_delay is calculated according to the size of the move and model of the filter wheel. For other filter wheel types, the filter_wheel_delay is a fixed value, which is specified as the `filter_wheel_delay` entry in the configuration file. The number of filter wheels connected to the controller is specified as `wheel_number` in the configuration file. Currently, both wheels are moved to the same position, but future implementations will enable control of both filter wheels independently.

```yaml
hardware:
  filter_wheel:
    type: SutterFilterWheel
    port: COM10
    baudrate: 9600
    number_of_wheels: 1

microscopes:
  microscope_name:
    filter_wheel:
    hardware:
      name: filter_wheel
      type: SutterFilterWheel
      wheel_number: 1
    filter_wheel_delay: .030
    available_filters:
      Empty-1: 0
      525-30: 1
      600-52: 2
      670-30: 3
      647-LP: 4
      Empty-2: 5
      Empty-3: 6
      Empty-4: 7
```

##### ASI

The ASI filter wheel is controlled by the ASI Tiger Controller. Thus, you should provide the same `comport` entry as you did for the stage. A single communication instance is used for both the stage and filter wheel.

```yaml
hardware:
  filter_wheel:
    type: ASI
    port: COM10
    baudrate: 115200
    number_of_wheels: 1

microscopes:
  microscope_name:
    filter_wheel:
      hardware:
        name: filter_wheel
        type: ASI
        wheel_number: 1
      filter_wheel_delay: .030
      available_filters:
        BLU - FF01-442/42-32: 0
        GFP - FF01-515/30-32: 1
        RFP - FF01-595/31-32: 2
        Far-Red - FF01-670/30-32: 3
        Blocked1: 4
        Empty: 5
        Blocked3: 6
        Blocked4: 7
        Blocked5: 8
        Blocked6: 9
```

##### Synthetic Filter Wheel

If no filter wheel is present, one must configure the software to use a synthetic filter wheel.

```yaml
hardware:
  filter_wheel:
    type: SyntheticFilterWheel
    port: COM10
    baudrate: 115200
    number_of_wheels: 1

microscopes:
  microscope_name:
    filter_wheel:
      hardware:
        name: filter_wheel
```

(continues on next page)

(continued from previous page)

```
      type: SyntheticFilterWheel
      wheel_number: 1
    filter_wheel_delay: .030
    available_filters:
      BLU - FF01-442/42-32: 0
      GFP - FF01-515/30-32: 1
      RFP - FF01-595/31-32: 2
      Far-Red - FF01-670/30-32: 3
      Blocked1: 4
      Empty: 5
      Blocked3: 6
      Blocked4: 7
      Blocked5: 8
      Blocked6: 9
```

---

##### Galvanometers

Galvo mirrors are used for fast scanning, shadow reduction, and occasionally as stages (see *Analog-Controlled Galvo/Piezo*).

##### Analog-Controlled Galvo

Multiple types of galvanometers have been used, including Cambridge Technologies/Novanta, Thorlabs, and Scanner-MAX Each of these devices are externally controlled via analog signals delivered from a data acquisition card.

```
microscopes:
  microscope_name:
    galvo:
      -
        hardware:
          name: daq
          type: NI
          channel: PCI6738/ao0
          min: -5
          max: 5
        waveform: sawtooth
        frequency: 99.9
        amplitude: 2.5
        offset: 0.5
        duty_cycle: 50
        phase: 1.57079
```

##### Synthetic Galvo

If no galvo is present, one must configure the software to use a synthetic galvo.

```yaml
microscopes:
  microscope_name:
    galvo:
      -
        hardware:
          name: daq
          type: SynthticGalvo
          channel: PCI6738/ao0
          min: -5
          max: 5
        waveform: sawtooth
        frequency: 99.9
        amplitude: 2.5
        offset: 0.5
        duty_cycle: 50
        phase: 1.57079
```

#### Lasers

We currently support laser control via voltage signals. In the near-future, we will consider implementing laser control via serial communication for power control, but digital modulation will still be controlled via voltage signals.

##### Omicron LightHUB Ultra

**Note:** Omicron laser source includes both Coherent- and LuxX lasers, which vary according to wavelength. LuxX lasers should be operated in an ACC operating mode with the analog modulation option enabled. The Coherent Obis lasers should be set in the mixed modulation mode.

##### Coherent Obis

**Note:** Coherent Obis lasers should be set in the mixed modulation mode. It is not uncommon for the slew rate from the data acquisition card to be insufficient to drive the modulation of the laser if the laser is set to an analog modulation mode.

##### Analog/Digital-Controlled Lasers

Most lasers are controlled externally via mixed analog and digital modulation. The `onoff` entry is for digital modulation. The `power` entry is for analog modulation.

```yaml
microscopes:
  microscope_name:
    lasers:
      - wavelength: 488
        onoff:
          hardware:
            name: daq
            type: NI
            channel: PCI6738/port1/line5
            min: 0
            max: 5
        power:
          hardware:
            name: daq
            type: NI
            channel: PCI6738/ao8
            min: 0
            max: 5
        type: Obis
        index: 0
        delay_percent: 10
        pulse_percent: 87
      - wavelength: 561...
```

---

#### Shutters

Shutters automatically open at the start of acquisition and close upon finish.

##### Analog/Digital-Controlled Shutters

Thorlabs shutters are controlled via a digital on off voltage.

```yaml
microscopes:
  microscope_name:
    shutter:
      hardware:
        name: daq
        type: NI
        channel: PXI6259/port0/line0
        min: 0
        max: 5
```

##### Synthetic Shutter

If no shutter is present, one must configure the software to use a synthetic shutter.

```yaml
hardware:
  shutter:
    hardware:
      name: daq
      type: synthetic
      channel: PCIE6738/port0/line0
      min: 0
      max: 5
```

---

##### Mechanical Zoom

Zoom devices control the magnification of the microscope. If such control is not needed, the software expects a *Synthetic Zoom* to provide the fixed magnification and the effective pixel size of the microscope.

##### Dynamixel Zoom

This software supports the Dynamixel Smart Actuator.

**Note:** The `positions` specify the voltage of the actuator at different zoom positions. The `stage_positions` account for focal shifts in between the different zoom values (the MVXPLAPO does not have a consistent focal plane). These may change depending on the immersion media. Here it is specified for a `BABB` (Benzyl Alcohol Benzyl Benzoate) immersion media. The `pixel_size` specifies the effective pixel size of the system at each zoom.

```yaml
hardware:
  zoom:
    type: DynamixelZoom
    servo_id: 1
    port: COM18
    baudrate: 1000000

microscopes:
  microscope_name:
    zoom:
      hardware:
          name: zoom
          type: DynamixelZoom
```

(continues on next page)

(continued from previous page)

```yaml
        servo_id: 1
      position:
        0.63x: 0
        1x: 627
        2x: 1711
        3x: 2301
        4x: 2710
        5x: 3079
        6x: 3383
      pixel_size:
        0.63x: 9.7
        1x: 6.38
        2x: 3.14
        3x: 2.12
        4x: 1.609
        5x: 1.255
        6x: 1.044
      stage_positions:
        BABB:
          f:
            0.63x: 0
            1x: 1
            2x: 2
            3x: 3
            4x: 4
            5x: 5
            6x: 6
```

##### Synthetic Zoom

```yaml
hardware:
  zoom:
    type: synthetic
    servo_id: 1
    port: COM18
    baudrate: 1000000

microscopes:
  microscope_name:
    zoom:
      hardware:
        name: zoom
        type: synthetic
        servo_id: 1
      position:
        36X: 0
      pixel_size:
```

(continues on next page)

(continued from previous page)

```
    36X: 0.180
  stage_positions:
    BABB:
      f:
        36X: 0
```

##### Deformable Mirrors

###### Imagine Optic

We currently have support for a Mirao 52E. The `flat_path` provides a path to a system correction `.wcs` file, an Imagine Optic proprietary file that stores actuator voltages and corresponding Zernike coefficients.

```
mirror:
  type: ImagineOpticsMirror

mirror:
    hardware:
      name: mirror
      type: ImagineOpticsMirror
      flat_path: D:\WaveKitX64\MirrorFiles\BeadsCoverslip_20231212.wcs
    n_modes: 32
```

###### Synthetic Mirror

It is not necessary to have a deformable mirror to run the software. If no deformable mirror is present, but one wants to evaluate the deformable mirror correction features, one must configure the software to use a synthetic deformable mirror.

```
mirror:
  type: SyntheticMirror

mirror:
    hardware:
      name: mirror
      type: ImagineOpticsMirror
      flat_path: D:\WaveKitX64\MirrorFiles\BeadsCoverslip_20231212.wcs
    n_modes: 32
```

##### 5.2.2 Microscope Implementations

For reference, we describe several microscope implementations that are currently in use with the **navigate** software. Each of these implementations is described in the following sections, including a list of the equipment used and the configuration file used to operate the microscope.

---

###### Multiscale Microscope

| Equipment | Description |
|---|---|
| Lasers | Omicron LightHUB Ultra with 488, 561, and 642 nm lasers. |
| Stages | PI L-509.20DG10, L-509.40DG10, L-509.20DG10, M-060.DG, M-406.4PD, PI P726.1CD |
| Stage Controllers | C-884, E-709 |
| Cameras | Hamamatsu Flash 4.0, Hamamatsu Fusion |
| Filter Wheel | Sutter Lambda 10-3 with 2x 32mm High-Speed Filter Wheels |
| Remote Focusing Units | Optotune Electrotunable Lens (EL-16-40-TC-VIS-5D-1-C) and Equipment Solutions LFA-2004 |
| Data Acquisition Cards | National Instruments PXIe-1073 chassis equipped with PXI6733 and PXI6259 |
| Galvo | Novanta CRS 4 KHz Resonant Galvo and Thorlabs GVS112 Linear Galvo |
| Zoom | Dynamixel MX-28R |
| Other | NA |

```yaml
# Specify all necessary information to find and connect to each hardware
# device that will be used on any of the scopes.
hardware:
  daq:
    type: NI
  camera:
    -
      type: HamamatsuOrca
      serial_number: 500502
    -
      type: HamamatsuOrca
      serial_number: 302352
  filter_wheel:
    type: SutterFilterWheel
    port: COM2
    baudrate: 9600
    number_of_wheels: 2
  stage:
    -
      type: PI
      controllername: C-884
      stages: L-509.20DG10 L-509.40DG10 L-509.20DG10 M-060.DG M-406.4PD NOSTAGE
      refmode: FRF FRF FRF FRF FRF FRF
      serial_number: 119060508
    -
      type: PI
      controllername: E-709
```

(continues on next page)

(continued from previous page)

```yaml
      stages: P-726.1CD
      refmode: ATZ
      serial_number: 0116049747
    # -
    #   type: MCL
    #   serial_number: 4011
  zoom:
    type: DynamixelZoom
    servo_id: 1
    port: COM9
    baudrate: 1000000

# Only one microscope can be active in the GUI at a time, but all microscopes will be␣
↪accessible
microscopes:
  Mesoscale:
    daq:
      hardware:
        name: daq
        type: NI

      # NI PCIe-1073 Chassis with PXI-6259 and PXI-6733 DAQ Boards.
      # Sampling rate in Hz
      sample_rate: 100000
      sweep_time: 0.2

      # triggers
      master_trigger_out_line: PXI6259/port0/line1
      camera_trigger_out_line: /PXI6259/ctr0
      trigger_source: /PXI6259/PFI0

      # Digital Laser Outputs
      laser_port_switcher: PXI6733/port0/line0
      laser_switch_state: False

    camera:
      hardware:
        name: camera
        type: HamamatsuOrca
        serial_number: 302352
      x_pixels: 2048.0
      y_pixels: 2048.0
      pixel_size_in_microns: 6.5
      subsampling: [1, 2, 4]
      sensor_mode: Normal  # 12 for progressive, 1 for normal.
      readout_direction: Top-to-Bottom  # Top-to-Bottom', 'Bottom-to-Top'
      lightsheet_rolling_shutter_width: 608
      defect_correct_mode: 2.0
      binning: 1x1
      readout_speed: 0x7FFFFFFF
      trigger_active: 1.0
      trigger_mode: 1.0 # external light-sheet mode
```

(continues on next page)

(continued from previous page)

```
    trigger_polarity: 2.0  # positive pulse
    trigger_source: 2.0  # 2 = external, 3 = software.
    exposure_time: 20 # Use milliseconds throughout.
    delay_percent: 10
    pulse_percent: 1
    line_interval: 0.000075
    display_acquisition_subsampling: 4
    average_frame_rate: 4.969
    frames_to_average: 1
    exposure_time_range:
      min: 1
      max: 1000
      step: 1
    flip_x: False
    flip_y: False
    x_pixels_step: 4
    y_pixels_step: 4
    x_pixels_min: 4
    y_pixels_min: 4

  remote_focus_device:
    hardware:
      name: daq
      type: NI
      channel: PXI6259/ao2
      min: -5
      max: 5
    # Optotune EL-16-40-TC-VIS-5D-1-C
    delay_percent: 7.5
    ramp_rising_percent: 85
    ramp_falling_percent: 2.5
    amplitude: 0.7
    offset: 2.3
    smoothing: 0.0
  galvo:
    -
      hardware:
        name: daq
        type: NI
        channel: PXI6259/ao0
        min: -5
        max: 5
      frequency: 99.9
      amplitude: 2.5
      offset: 0
      duty_cycle: 50
      phase: 1.57079 # pi/2
  filter_wheel:
    hardware:
      name: filter_wheel
      type: SutterFilterWheel
      wheel_number: 1
```

(continues on next page)

(continued from previous page)

```
    filter_wheel_delay: .030 # in seconds
    available_filters:
      Empty-Alignment: 5
      GFP - FF01-515/30-32: 6
      RFP - FF01-595/31-32: 7
      Far-Red - BLP01-647R/31-32: 8
      Blocked1: 4
      Blocked2: 0
      Blocked3: 1
      Blocked4: 2
      Blocked5: 3
      Blocked6: 9
  stage:
    hardware:
      name: stage
      type: PI
      serial_number: 119060508
      axes: [x, y, z, theta, f]
    y_unload_position: 10000
    y_load_position: 90000

    startfocus: 75000
    x_max: 100000
    x_min: -100000
    y_max: 100000
    y_min: -100000
    z_max: 100000
    z_min: -100000
    f_max: 100000
    f_min: 0
    theta_max: 360
    theta_min: 0

    x_rot_position: 2000
    y_rot_position: 2000
    z_rot_position: 2000

    x_step: 500
    y_step: 500
    z_step: 500
    theta_step: 30
    f_step: 500

    position:
      x_pos: 25250
      y_pos: 40000
      z_pos: 40000
      f_pos: 70000
      theta_pos: 0
    velocity: 1000

    x_offset: 0
```

(continues on next page)

(continued from previous page)

```
    y_offset: 0
    z_offset: 0
    f_offset: 0
    theta_offset: 0

    flip_x: False
    flip_y: False
    flip_z: False
  zoom:
    hardware:
      name: zoom
      type: DynamixelZoom
      servo_id: 1
    position:
      0.63x: 0
      1x: 627
      2x: 1711
      3x: 2301
      4x: 2710
      5x: 3079
      6x: 3383
    pixel_size:
      0.63x: 9.7
      1x: 6.38
      2x: 3.14
      3x: 2.12
      4x: 1.609
      5x: 1.255
      6x: 1.044
    stage_positions:
      BABB:
        f:
          0.63x: 67410
          1x: 70775
          2x: 72455
          3x: 72710
          4x: 72795
          5x: 72850
          6x: 72880
      CUBIC:
        f:
          0.63x: 67410
          1x: 70775
          2x: 72455
          3x: 72710
          4x: 72795
          5x: 72850
          6x: 72880
  shutter:
    hardware:
      name: daq
      type: NI
```

(continues on next page)

(continued from previous page)

```yaml
      channel: PXI6259/port0/line0
      min: 0
      max: 5
  lasers:
    # Omicron LightHub Ultra
    # 488 and 640 are LuxX+ Lasers
    # 561 is a Coherent OBIS Laser
    # Digital Laser Outputs
    - wavelength: 488
      onoff:
        hardware:
          name: daq
          type: NI
          channel: PXI6733/port0/line2
          min: 0
          max: 5
      power:
        hardware:
          name: daq
          type: NI
          channel: PXI6733/ao0
          min: 0
          max: 5
      type: LuxX
      index: 0
      delay_percent: 10
      pulse_percent: 87
    - wavelength: 562
      onoff:
        hardware:
          name: daq
          type: NI
          channel: PXI6733/port0/line3
          min: 0
          max: 5
      power:
        hardware:
          name: daq
          type: NI
          channel: PXI6733/ao1
          min: 0
          max: 5
      type: Obis
      index: 1
      delay_percent: 10
      pulse_percent: 87
    - wavelength: 642
      onoff:
        hardware:
          name: daq
          type: NI
          channel: PXI6733/port0/line4
```

(continues on next page)

(continued from previous page)

```yaml
          min: 0
          max: 5
      power:
        hardware:
          name: daq
          type: NI
          channel: PXI6733/ao2
          min: 0
          max: 5
      type: LuxX
      index: 2
      delay_percent: 10
      pulse_percent: 87

Nanoscale:
  daq:
    hardware:
      name: daq
      type: NI

    # NI PCIe-1073 Chassis with PXI-6259 and PXI-6733 DAQ Boards.
    # Sampling rate in Hz
    sample_rate: 100000
    sweep_time: 0.2

    # triggers
    master_trigger_out_line: PXI6259/port0/line1
    camera_trigger_out_line: /PXI6259/ctr0
    trigger_source: /PXI6259/PFI0

    # Digital Laser Outputs
    laser_port_switcher: PXI6733/port0/line0
    laser_switch_state: True

  camera:
    hardware:
      type: HamamatsuOrca
      serial_number: 500502
    x_pixels: 2304.0
    y_pixels: 2304.0
    pixel_size_in_microns: 6.5
    subsampling: [1, 2, 4]
    sensor_mode: Normal  # 12 for progressive, 1 for normal.
    readout_direction: Top-to-Bottom  # Top-to-Bottom', 'Bottom-to-Top'
    lightsheet_rolling_shutter_width: 100
    defect_correct_mode: 2.0
    binning: 1x1
    readout_speed: 0x7FFFFFFF
    trigger_active: 1.0
    trigger_mode: 1.0 # external light-sheet mode
    trigger_polarity: 2.0  # positive pulse
    trigger_source: 2.0  # 2 = external, 3 = software.
```

(continues on next page)

(continued from previous page)

```
    exposure_time: 20 # Use milliseconds throughout.
    delay_percent: 20
    pulse_percent: 1
    line_interval: 0.000075
    display_acquisition_subsampling: 4
    average_frame_rate: 4.969
    frames_to_average: 1
    exposure_time_range:
      min: 1
      max: 1000
      step: 1
    x_pixels_step: 4
    y_pixels_step: 4
    x_pixels_min: 4
    y_pixels_min: 4

  remote_focus_device:
    hardware:
      name: daq
      type: EquipmentSolutions  #NI
      channel: PXI6259/ao3
      comport: COM6
      min: -5
      max: 5
    # ThorLabs BLINK
    delay_percent: 0
    ramp_rising_percent: 85
    ramp_falling_percent: 2.5
    amplitude: 0.7
    offset: 2.3
    smoothing: 0.0
    # waveform: trig_remote_focus_ramp
  galvo:
    -
      hardware:
        name: daq
        type: NI
        channel: PXI6259/ao1
        min: -5
        max: 5
      offset: 0.5
  filter_wheel:
    hardware:
      name: filter_wheel
      type: SutterFilterWheel
      wheel_number: 2
    filter_wheel_delay: .030 # in seconds
    available_filters:
      Empty-Alignment: 0
      GFP - FF01-515/30-32: 1
      RFP - FF01-595/31-32: 2
      Far-Red - BLP01-647R/31-32: 3
```

(continues on next page)

(continued from previous page)

```yaml
      Blocked1: 4
      Blocked2: 5
      Blocked3: 6
      Blocked4: 7
      Blocked5: 8
      Blocked6: 9
  stage:
    hardware:
      -
        name: stage
        type: PI
        serial_number: 119060508
        axes: [x, y, z, theta]
      -
        name: stage2
        type: PI
        serial_number: 0116049747
        axes: [f]
      # -
      #   name: stage2
      #   type: MCL
      #   serial_number: 4011
      #   axes: [f]
    y_unload_position: 10000
    y_load_position: 90000

    startfocus: 50
    x_max: 100000
    x_min: -100000
    y_max: 100000
    y_min: -100000
    z_max: 100000
    z_min: -100000
    f_max: 100000
    f_min: 0
    theta_max: 360
    theta_min: 0

    x_rot_position: 2000
    y_rot_position: 2000
    z_rot_position: 2000

    x_step: 500
    y_step: 500
    z_step: 500
    theta_step: 30
    f_step: 5

    position:
      x_pos: 25250
      y_pos: 40000
      z_pos: 40000
```

(continues on next page)

(continued from previous page)

```yaml
        f_pos: 0
        theta_pos: 0
      velocity: 1000

      x_offset: 500  # -1000
      y_offset: 300  # -70
      z_offset: -18396  # -17842
      # x_offset: 0
      # y_offset: 0
      # z_offset: 0
      f_offset: 0
      theta_offset: 0
    zoom:
      position:
        N/A: 0
      pixel_size:
        N/A: 0.167
    shutter:
      hardware:
        name: daq
        type: NI
        channel: PXI6259/port2/line0
      shutter_min_do: 0
      shutter_max_do: 5
    lasers:
      # Omicron LightHub Ultra
      # 488 and 640 are LuxX+ Lasers
      # 561 is a Coherent OBIS Laser
      # Digital Laser Outputs
      - wavelength: 488
        onoff:
          hardware:
            name: daq
            type: NI
            channel: PXI6733/port0/line2
            min: 0
            max: 5
        power:
          hardware:
            name: daq
            type: NI
            channel: PXI6733/ao0
            min: 0
            max: 5
        type: LuxX
        index: 0
        delay_percent: 10
        pulse_percent: 87
      - wavelength: 562
        onoff:
          hardware:
            name: daq
```

(continues on next page)

(continued from previous page)

```
          type: NI
          channel: PXI6733/port0/line3
          min: 0
          max: 5
      power:
        hardware:
          name: daq
          type: NI
          channel: PXI6733/ao1
          min: 0
          max: 5
      type: Obis
      index: 1
      delay_percent: 10
      pulse_percent: 87
    - wavelength: 642
      onoff:
        hardware:
          name: daq
          type: NI
          channel: PXI6733/port0/line4
          min: 0
          max: 5
      power:
        hardware:
          name: daq
          type: NI
          channel: PXI6733/ao2
          min: 0
          max: 5
      type: LuxX
      index: 2
      delay_percent: 10
      pulse_percent: 87

gui:
  channels:
    count: 5
    laser_power:
      min: 0
      max: 100
      step: 10
    exposure_time:
      min: 1
      max: 1000
      step: 5
    interval_time:
      min: 0
      max: 1000
      step: 5
  stack_acquisition:
    step_size:
```

(continues on next page)

(continued from previous page)

```
      min: 0.200
      max: 1000
      step: 0.1
    start_pos:
      min: -5000
      max: 5000
      step: 1
    end_pos:
      min: -5000
      max: 10000
      step: 1
  timepoint:
    timepoints:
      min: 1
      max: 1000
      step: 1
    stack_pause:
      min: 0
      max: 1000
      step: 1
```

##### Expansion ASLM

| Equipment | Description |
|---|---|
| Lasers | Omicron LightHUB Ultra with 405, 488, 561, and 642 nm lasers. |
| Stages | ASI FTP-2000 with Linear Encoders in X and Y, and 3x LS-50 Linear Stages |
| Stage Controllers | ASI Tiger Controller |
| Cameras | Hamamatsu Lightning and Photometrics Iris15 |
| Filter Wheel | 2x ASI 6-Position 32 mm Filter Wheels |
| Remote Focusing Units | ThorLabs BLINK |
| Data Acquisition Cards | National Instruments PXIe-1073 chassis equipped with PXI6733 and PXI6259 |
| Galvo | Novanta CRS 4 KHz Resonant Galvo |
| Zoom | N/A |
| Other | NA |

```
# Specify all necessary information to find and connect to each hardware
# device that will be used on any of the scopes.
hardware:
  daq:
    type: NI
  camera:
    -
      type:  HamamatsuOrcaLightning #SyntheticCamera
```

(continues on next page)

(continued from previous page)

```yaml
      serial_number: 000035
    -
      type: Photometrics
      camera_connection: PMPCIECam00
      serial_number: 1
  filter_wheel:
    type: ASI #SyntheticFilterWheel
    port: COM8
    baudrate: 115200
    number_of_wheels: 2
  stage:
    -
      type: ASI
      serial_number: 123456789
      port: COM8
      baudrate: 115200
    -
      type: SyntheticStage
      serial_number: 987654321
  zoom:
    type: SyntheticZoom
    servo_id: 0
    port: 0
    baudrate: 0

# Only one microscope can be active in the GUI at a time, but all microscopes will be␣
↪accessible
microscopes:
  Nanoscale:
    daq:
      hardware:
        name: daq
        type: NI

      # NI PCIe-1073 Chassis with PXI-6259 and PXI-6733 DAQ Boards.
      # Sampling rate in Hz
      sample_rate: 100000
      sweep_time: 0.2

      # triggers
      master_trigger_out_line: PXI6259/port0/line1
      camera_trigger_out_line: /PXI6259/ctr0
      trigger_source: /PXI6259/PFI0

      # Digital Laser Outputs
      laser_port_switcher: PXI6733/port0/line1
      laser_switch_state: False

    camera:
      hardware:
        name: camera
        type: HamamatsuOrcaLightning #SyntheticCamera
```

(continues on next page)

(continued from previous page)

```
      serial_number: 000035
    x_pixels: 4608.0
    y_pixels: 2592.0
    pixel_size_in_microns: 5.5
    subsampling: [1, 2, 4]
    sensor_mode: Normal  # 12 for progressive, 1 for normal.
    readout_direction: Bottom-to-Top  # Top-to-Bottom', 'Bottom-to-Top'
    lightsheet_rolling_shutter_width: 608
    defect_correct_mode: 2.0
    binning: 1x1
    readout_speed: 0x7FFFFFFF
    trigger_active: 1.0
    trigger_mode: 1.0 # external light-sheet mode
    trigger_polarity: 2.0  # positive pulse
    trigger_source: 2.0  # 2 = external, 3 = software.
    exposure_time: 20 # Use milliseconds throughout.
    delay_percent: 30 #30 #25 #8 #5.0
    pulse_percent: 1
    line_interval: 0.000075
    display_acquisition_subsampling: 4
    average_frame_rate: 4.969
    frames_to_average: 1
    exposure_time_range:
      min: 1
      max: 1000
      step: 1
  remote_focus_device:
    hardware:
      name: daq
      type: NI
      channel: PXI6259/ao3
      min: -0.5
      max: 0.5
    # Optotune EL-16-40-TC-VIS-5D-1-C
    delay_percent: 0 #1.5 #7.5
    ramp_rising_percent: 85
    ramp_falling_percent: 1.5 #2.5
    amplitude: 0.7
    offset: 2.3
  galvo:
    -
      hardware:
        name: daq
        type: NI
        channel: PXI6259/ao1
        min: 0
        max: 5
      frequency: 99.9
      amplitude: 2.5
      offset: 0
      duty_cycle: 50
      phase: 1.57079 # pi/2
```

(continues on next page)

(continued from previous page)

```yaml
    filter_wheel:
      hardware:
        name: filter_wheel
        type: ASI #SyntheticFilterWheel
        wheel_number: 2
      filter_wheel_delay: .030 # in seconds
      available_filters:
        BLU - FF01-442/42-32: 0
        GFP - FF01-515/30-32: 1
        RFP - FF01-595/31-32: 2
        Far-Red - FF01-670/30-32: 3
        Blocked1: 4
        Empty: 5
        Blocked3: 6
        Blocked4: 7
        Blocked5: 8
        Blocked6: 9
    stage:
      hardware:
        -
          name: stage
          type: ASI
          serial_number: 123456789
          axes: [x, y, z, f] # Software
          axes_mapping: [M, Y, X, Z] # M Shear axis mapping
          #axes_mapping: [M, X, Y, Z] #testing y
          #axes_mapping: [M, X, Z, Y] #testing Z
          # axes_mapping: [Z, Y, X, M]
          feedback_alignment: [90, 90, 90, 90]

        -
          name: stage
          type: SyntheticStage
          serial_number: 987654321
          axes: [theta]

      startfocus: -16000
      x_max: 0 # Swapped from Z
      x_min: -22708.3 # Swapped from Z
      y_max: 1361.3
      y_min: -3496.3
      z_max: 3521.9 # Swapped from X
      z_min: -4551.1 # Swapped from X
      f_max: 3233.0 #=m
      f_min: -9382.0 #=m
      theta_max: 360
      theta_min: 0
      external_trigger: /PXI6259/PFI1
      # joystick_axes: [x, y, z, f]

      x_rot_position: 2000
      y_rot_position: 2000
```

(continues on next page)

(continued from previous page)

```
    z_rot_position: 2000

    x_step: 50
    y_step: 50
    z_step: 50
    theta_step: 30
    f_step: 50

    position:
      x_pos: 5 # Swapped from Z initial stage position
      y_pos: 1
      z_pos: 1 # Swapped from X
      f_pos: 0
      theta_pos: 0
    velocity: 1000

    x_offset: 0
    y_offset: 0
    z_offset: 0
    f_offset: 0
    theta_offset: 0
  zoom:
    hardware:
      name: zoom
      type: SyntheticZoom
      servo_id: 1
    position:
      N/A: 0
    pixel_size:
      N/A: 0.168
  shutter:
    hardware:
      name: daq
      type: NI
      channel: PXI6259/port0/line0
      min: 0
      max: 5
  lasers:
    # Omicron LightHub Ultra
    # 488 and 640 are LuxX+ Lasers
    # 561 is a Coherent OBIS Laser
    # Digital Laser Outputs
    - wavelength: 405
      onoff:
        hardware:
          name: daq
          type: NI
          channel: PXI6733/port0/line2
          min: 0
          max: 5
      power:
        hardware:
```

(continues on next page)

(continued from previous page)

```
          name: daq
          type: NI
          channel: PXI6733/ao0
          min: 0
          max: 5
      type: LuxX
      index: 0
      delay_percent: 10
      pulse_percent: 87
    - wavelength: 488
      onoff:
        hardware:
          name: daq
          type: NI
          channel: PXI6733/port0/line3
          min: 0
          max: 5
      power:
        hardware:
          name: daq
          type: NI
          channel: PXI6733/ao1
          min: 0
          max: 5
      type: LuxX
      index: 1
      delay_percent: 10
      pulse_percent: 87
    - wavelength: 561
      onoff:
        hardware:
          name: daq
          type: NI
          channel: PXI6733/port0/line4
          min: 0
          max: 5
      power:
        hardware:
          name: daq
          type: NI
          channel: PXI6733/ao2
          min: 0
          max: 5
      type: Obis
      index: 2
      delay_percent: 10
      pulse_percent: 87
    - wavelength: 642
      onoff:
        hardware:
          name: daq
          type: NI
```

(continues on next page)

(continued from previous page)

```yaml
        channel: PXI6733/port0/line5
        min: 0
        max: 5
    power:
      hardware:
        name: daq
        type: NI
        channel: PXI6733/ao3
        min: 0
        max: 5
    type: LuxX
    index: 3
    delay_percent: 10
    pulse_percent: 87
  - wavelength: LED
    power:
      hardware:
        name: daq
        type: NI
        channel: PXI6733/ao4
        min: 0
        max: 5
    index: 4

Macroscale:
  daq:
    hardware:
      name: daq
      type: NI

    # NI PCIe-1073 Chassis with PXI-6259 and PXI-6733 DAQ Boards.
    # Sampling rate in Hz
    sample_rate: 100000
    sweep_time: 0.2

    # triggers
    master_trigger_out_line: PXI6259/port0/line1
    camera_trigger_out_line: /PXI6259/ctr0
    trigger_source: /PXI6259/PFI0

    # Digital Laser Outputs
    laser_port_switcher: PXI6733/port0/line1
    laser_switch_state: True

  camera:
    hardware:
      name: camera
      type: Photometrics #SyntheticCamera
      serial_number: 1
    x_pixels: 5056.0
    y_pixels: 2960.0
    pixel_size_in_microns: 4.25
```

(continues on next page)

(continued from previous page)

```yaml
    subsampling: [1, 2, 4]
    sensor_mode: Normal  # 12 for progressive, 1 for normal.
    readout_direction: Bottom-to-Top  # Top-to-Bottom', 'Bottom-to-Top'
    lightsheet_rolling_shutter_width: 608
    defect_correct_mode: 2.0
    binning: 1x1
    readout_speed: 0x7FFFFFFF
    trigger_active: 1.0
    trigger_mode: 1.0 # external light-sheet mode
    trigger_polarity: 2.0  # positive pulse
    trigger_source: 2.0  # 2 = external, 3 = software.
    exposure_time: 20 # Use milliseconds throughout.
    delay_percent: 25 #8 #5.0
    pulse_percent: 1
    line_interval: 0.000075
    display_acquisition_subsampling: 4
    average_frame_rate: 4.969
    frames_to_average: 1
    exposure_time_range:
      min: 1
      max: 1000
      step: 1
  remote_focus_device:
    hardware:
      name: daq
      type: NI
      channel: PXI6259/ao3
      min: -0.5
      max: 0.5
    # Optotune EL-16-40-TC-VIS-5D-1-C
    delay_percent: 0 #1.5 #7.5
    ramp_rising_percent: 85
    ramp_falling_percent: 2.5
    amplitude: 0.7
    offset: 2.3
  galvo:
    -
      hardware:
        name: daq
        type: NI
        channel: PXI6259/ao1
        min: 0
        max: 5
      frequency: 99.9
      amplitude: 2.5
      offset: 0
      duty_cycle: 50
      phase: 1.57079 # pi/2
  filter_wheel:
    hardware:
      name: filter_wheel
      type: ASI #SyntheticFilterWheel
```

(continues on next page)

(continued from previous page)

```
      wheel_number: 2
    filter_wheel_delay: .030 # in seconds
    available_filters:
      BLU - FF01-442/42-32: 0
      GFP - FF01-515/30-32: 1
      RFP - FF01-595/31-32: 2
      Far-Red - FF01-670/30-32: 3
      Blocked1: 4
      Empty: 5
      Blocked3: 6
      Blocked4: 7
      Blocked5: 8
      Blocked6: 9
  stage:
    hardware:
      -
        name: stage
        type: ASI
        serial_number: 123456789
        axes: [x, y, z, f] #Software
        # axes_mapping: [M, Y, X, Z]
        axes_mapping: [M, Y, X, Z] #M  shear
        feedback_alignment: [90, 90, 90, 90]

      -
        name: stage
        type: SyntheticStage
        serial_number: 987654321
        axes: [theta]

    startfocus: -16000
    x_max: 0 # Swapped from Z
    x_min: -22708.3 # Swapped from Z
    y_max: 1361.3
    y_min: -3496.3
    z_max: 3521.9 # Swapped from X
    z_min: -4651.1 # Swapped from X
    f_max: 3233.0 #=m
    f_min: -9382.0 #=m
    theta_max: 360
    theta_min: 0
    external_trigger: /PXI6259/PFI1
    # joystick_axes: [x, y, z, f]

    x_rot_position: 2000
    y_rot_position: 2000
    z_rot_position: 2000

    x_step: 50
    y_step: 50
    z_step: 50
    theta_step: 30
```

(continues on next page)

(continued from previous page)

```
      f_step: 50

      position:
        x_pos: 5 # Swapped from Z initial stage position
        y_pos: 1
        z_pos: 1 # Swapped from X
        f_pos: 0
        theta_pos: 0
      velocity: 1000

      x_offset: 0
      y_offset: 0
      z_offset: 0
      f_offset: 0
      theta_offset: 0
    zoom:
      hardware:
        name: zoom
        type: SyntheticZoom
        servo_id: 1
      position:
        N/A: 0
      pixel_size:
        N/A: 1.06
    shutter:
      hardware:
        name: daq
        type: NI
        channel: PXI6259/port0/line0
        min: 0
        max: 5
    lasers:
      # Omicron LightHub Ultra
      # 488 and 640 are LuxX+ Lasers
      # 561 is a Coherent OBIS Laser
      # Digital Laser Outputs
      - wavelength: 405
        onoff:
          hardware:
            name: daq
            type: NI
            channel: PXI6733/port0/line2
            min: 0
            max: 5
        power:
          hardware:
            name: daq
            type: NI
            channel: PXI6733/ao0
            min: 0
            max: 5
        type: LuxX
```

(continues on next page)

(continued from previous page)

```
      index: 0
      delay_percent: 10
      pulse_percent: 87
    - wavelength: 488
      onoff:
        hardware:
          name: daq
          type: NI
          channel: PXI6733/port0/line3
          min: 0
          max: 5
      power:
        hardware:
          name: daq
          type: NI
          channel: PXI6733/ao1
          min: 0
          max: 5
      type: LuxX
      index: 1
      delay_percent: 10
      pulse_percent: 87
    - wavelength: 561
      onoff:
        hardware:
          name: daq
          type: NI
          channel: PXI6733/port0/line4
          min: 0
          max: 5
      power:
        hardware:
          name: daq
          type: NI
          channel: PXI6733/ao2
          min: 0
          max: 5
      type: Obis
      index: 2
      delay_percent: 10
      pulse_percent: 87
    - wavelength: 642
      onoff:
        hardware:
          name: daq
          type: NI
          channel: PXI6733/port0/line5
          min: 0
          max: 5
      power:
        hardware:
          name: daq
```

(continues on next page)

(continued from previous page)

```
            type: NI
            channel: PXI6733/ao3
            min: 0
            max: 5
        type: LuxX
        index: 3
        delay_percent: 10
        pulse_percent: 87
      - wavelength: LED
        power:
          hardware:
            name: daq
            type: NI
            channel: PXI6733/ao4
            min: 0
            max: 5
        index: 4

gui:
  channels:
    count: 5
    laser_power:
      min: 0
      max: 100
      step: 10
    exposure_time:
      min: 1
      max: 1000
      step: 5
    interval_time:
      min: 0
      max: 1000
      step: 5
  stack_acquisition:
    step_size:
      min: 0.1
      max: 1000
      step: 0.1
    start_pos:
      min: -5000
      max: 5000
      step: 1
    end_pos:
      min: -5000
      max: 10000
      step: 1
  timepoint:
    timepoints:
      min: 1
      max: 1000
      step: 1
```

(continues on next page)

(continued from previous page)

```yaml
    stack_pause:
      min: 0
      max: 1000
      step: 1
```

#### OPM-V2

| Equipment | Description |
|---|---|
| Lasers | Coherent Galaxy with 488, 561, and 642 nm lasers. |
| Stages | ASI FTP-2000 with MS-2000 XY stage, and a Galvo for acquisition of z-stacks. |
| Stage Controllers | ASI Tiger Controller |
| Cameras | Hamamatsu Flash 4.0 |
| Filter Wheel | 2x ASI 6-Position 32 mm Filter Wheels |
| Remote Focusing Units | Optotune Electrotunable Lens (EL-16-40-TC-VIS-5D-1-C) |
| Data Acquisition Cards | National Instruments PCIe-6738 |
| Galvo | Novanta CRS 4 KHz Resonant Galvo, and 2x Novanta Linear Galvos for shearing and tiling. |
| Zoom | N/A |
| Other | NA |

```yaml
# Specify all necessary information to find and connect to each hardware
# device that will be used on any of the scopes.
hardware:
  daq:
    type: NI
  camera:
    -
      type: HamamatsuOrca
      serial_number: 304064
  filter_wheel:
    type: SyntheticFilterWheel
    port: COM6
    baudrate: 9600
    number_of_wheels: 1
  stage:
    -
      type: SyntheticStage
      serial_number: 123
    - type: GalvoNIStage
      serial_number: 123
  zoom:
    type: SyntheticZoom
    servo_id: 1
```

(continues on next page)

(continued from previous page)

```yaml
# Only one microscope can be active in the GUI at a time, but all microscopes will be␣
↪accessible
microscopes:
  OPMv2:
    daq:
      hardware:
        name: daq
        type: NI

      # NI PCIe-1073 Chassis with PXI-6259 and PXI-6733 DAQ Boards.
      # Sampling rate in Hz
      sample_rate: 100000
      sweep_time: 0.2

      # triggers
      master_trigger_out_line: /Dev5/port0/line1
      camera_trigger_out_line: /Dev5/ctr0
      trigger_source: /Dev5/PFI0

    zoom:
      hardware:
        name: zoom
        type: SyntheticZoom
        servo_id: 1
      position:
        1x: 0
      pixel_size:
        1x: 0.15
    shutter:
      hardware:
        name: shutter
        type: SyntheticShutter
        channel: none/line0
      shutter_min_do: 0
      shutter_max_do: 5
    camera:
      hardware:
        name: camera
        type: HamamatsuOrca
        serial_number: 304064
      x_pixels: 2048.0
      y_pixels: 2048.0
      pixel_size_in_microns: 6.5
      subsampling: [1, 2, 4]
      sensor_mode: Normal  # 12 for progressive, 1 for normal.
      readout_direction: Bottom-to-Top  # Top-to-Bottom', 'Bottom-to-Top'
      lightsheet_rolling_shutter_width: 10
      defect_correct_mode: 2.0
      binning: 1x1
      readout_speed: 1.0
      trigger_active: 1.0
```

(continues on next page)

(continued from previous page)

```
    trigger_mode: 1.0 # external light-sheet mode
    trigger_polarity: 2.0  # positive pulse
    trigger_source: 2.0  # 2 = external, 3 = software.
    exposure_time: 20 # Use milliseconds throughout.
    delay_percent: 7.5
    pulse_percent: 1
    line_interval: 0.000075
    display_acquisition_subsampling: 4
    average_frame_rate: 4.969
    frames_to_average: 1
    exposure_time_range:
      min: 1
      max: 1000
      step: 1
  remote_focus_device:
    hardware:
      name: daq
      type: NI
      channel: Dev5/ao3
      min: 0
      max: 5
    # Optotune EL-16-40-TC-VIS-5D-1-C
    delay_percent: 5
    ramp_rising_percent: 92.5
    ramp_falling_percent: 2.5
    amplitude: 0.7
    offset: 2.3
  galvo:
    # -
    #   name: xgalvo
    #   hardware:
    #     name: daq
    #     type: NI
    #     channel: Dev5/ao0
    #     min: -5
    #     max: 5
    #   frequency: 99.9
    #   amplitude: 2.5
    #   offset: 0
    #   duty_cycle: 50
    #   phase: 1.57079 # pi/2
    -
      name: ygalvo
      hardware:
        name: daq
        type: NI
        channel: Dev5/ao1
        min: -5
        max: 5
      frequency: 99.9
      amplitude: 2.5
      offset: 0
```

(continues on next page)

(continued from previous page)

```
        duty_cycle: 50
        phase: 1.57079 # pi/2
      -
        name: sheargalvo
        hardware:
          name: daq
          type: NI
          channel: Dev5/ao2
          min: -5
          max: 5
        frequency: 99.9
        amplitude: 2.5
        offset: 0
        duty_cycle: 50
        phase: 1.57079 # pi/2
    stage:
      hardware:
        -
          name: fake_stage
          type: SyntheticStage
          serial_number: 123
          axes: [x,y,theta,f]
        -
          name: galvo-stage
          type: GalvoNIStage
          serial_number: 123
          axes: [z]
          axes_mapping: [Dev5/ao0]
          min: -5
          max: 5
          volts_per_micron: 0.01*x + 0
      y_unload_position: 10000
      y_load_position: 90000

      startfocus: 75000
      x_max: 500
      x_min: -500
      y_max: 500
      y_min: -500
      z_max: 500
      z_min: -500
      f_max: 100000
      f_min: -100000
      theta_max: 360
      theta_min: 0

      x_rot_position: 2000
      y_rot_position: 2000
      z_rot_position: 2000

      x_step: 500
      y_step: 500
```

(continues on next page)

(continued from previous page)

```
    z_step: 10
    theta_step: 30
    f_step: 500

    position:
      x_pos: 25250
      y_pos: 40000
      z_pos: 0
      f_pos: 70000
      theta_pos: 0
    velocity: 1000

    x_offset: 0
    y_offset: 0
    z_offset: 0
    f_offset: 0
    theta_offset: 0
  filter_wheel:
    hardware:
      name: filter_wheel
      type: SyntheticFilterWheel
      wheel_number: 1
    filter_wheel_delay: .030 # in seconds
    available_filters:
      FRFP - BLP01-664R-25: 0
      RFP - FF01-598/25-25: 1
      GFP - 527/20: 2
      GFPRFP - ZET488/561m: 3
      Empty-Alignment: 4
      Blocked2: 5
      Blocked3: 6
      Blocked4: 7
      Blocked5: 8
      Blocked6: 9
  lasers:
    # Omicron LightHub Ultra
    # 488 and 640 are LuxX+ Lasers
    # 561 is a Coherent OBIS Laser
    # Digital Laser Outputs
    - wavelength: 488
      power:
        hardware:
          name: daq
          type: NI
          channel: Dev5/ao12
          min: 0
          max: 5
      type: LuxX
      index: 0
      delay_percent: 10
      pulse_percent: 87
    - wavelength: 562
```

(continues on next page)

(continued from previous page)

```
        onoff:
          hardware:
            name: daq
            type: NI
            channel: Dev5/port1/line5
            min: 0
            max: 5
        power:
          hardware:
            name: daq
            type: NI
            channel: Dev5/ao13
            min: 0
            max: 5
        type: Obis
        index: 1
        delay_percent: 10
        pulse_percent: 87
      - wavelength: 642
        power:
          hardware:
            name: daq
            type: NI
            channel: Dev5/ao14
            min: 0
            max: 5
        type: LuxX
        index: 2
        delay_percent: 10
        pulse_percent: 87

gui:
  channels:
    count: 5
    laser_power:
      min: 0
      max: 100
      step: 10
    exposure_time:
      min: 1
      max: 1000
      step: 5
    interval_time:
      min: 0
      max: 1000
      step: 5
  stack_acquisition:
    step_size:
      min: 0.200
      max: 1000
      step: 0.1
    start_pos:
```

(continues on next page)

(continued from previous page)

```
    min: -5000
    max: 5000
    step: 1
  end_pos:
    min: -5000
    max: 10000
    step: 1
timepoint:
  timepoints:
    min: 1
    max: 1000
    step: 1
  stack_pause:
    min: 0
    max: 1000
    step: 1
confocal_projection:
  scanrange:
    min: 0
    max: 600
    step: 1
  offset_start:
    min: -300
    max: 300
    step: 0.1
  offset_end:
    min: -300
    max: 300
    step: 0.1
  n_plane:
    min: 1
    max: 200
    step: 1
```

##### OPM-V3

| Equipment | Description |
|---|---|
| Lasers | Omicron LightHUB Ultra with 488 and 561 nm lasers. |
| Stages | A piezo for adjusting the position of the tertiary objective, and a galvo for acquisition of z-stacks. |
| Stage Controllers | N/A |
| Cameras | Hamamatsu Flash 4.0 |
| Filter Wheel | N/A |
| Remote Focusing Units | N/A |
| Data Acquisition Cards | National Instruments PCIe-6738 |
| Galvo | Two Novanta galvos for shearing and lateral sweeping of the illumination beam. |
| Zoom | N/A |
| Other | VAST large object flow cytometry system and Imagine Optics deformable mirror for wavefront correction. |
| Other | NA |

```yaml
# Specify all necessary information to find and connect to each hardware
# device that will be used on any of the scopes.
hardware:
  daq:
    type: NI
  camera:
    -
      type: HamamatsuOrca
      serial_number: 001301
  filter_wheel:
    type: SyntheticFilterWheel
    port: COM6
    baudrate: 9600
    number_of_wheels: 1
  stage:
    -
      type: SyntheticStage
      serial_number: 123
    -
      type: GalvoNIStage
      serial_number: 124
    -
      type: GalvoNIStage
      serial_number: 125
  zoom:
    type: SyntheticZoom
    servo_id: 1
  mirror:
    type: ImagineOpticsMirror

# Only one microscope can be active in the GUI at a time, but all microscopes will be
↪accessible
microscopes:
  ProjectionScope:
    daq:
```

(continues on next page)

(continued from previous page)

```
    hardware:
      name: daq
      type: NI

    # NI PCIe-1073 Chassis with PXI-6259 and PXI-6733 DAQ Boards.
    # Sampling rate in Hz
    sample_rate: 100000
    sweep_time: 0.2

    # triggers
    master_trigger_out_line: /PCIe-6738/port0/line1
    camera_trigger_out_line: /PCIe-6738/ctr0 #PFI7 Camera trigger
    trigger_source: /PCIe-6738/PFI0

  mirror:
    hardware:
      name: mirror
      type: ImagineOpticsMirror
      flat_path: D:\WaveKitX64\MirrorFiles\BeadsCoverslip_20231212.wcs
    n_modes: 32

  zoom:
    hardware:
      name: zoom
      type: SyntheticZoom
      servo_id: 1
    position:
      1x: 0
    pixel_size:
      1x: 0.15
  shutter:
    hardware:
      name: shutter
      type: SyntheticShutter
      channel: none/line0
    shutter_min_do: 0
    shutter_max_do: 5
  camera:
    hardware:
      name: camera
      type: HamamatsuOrca
      serial_number: 001301
    x_pixels: 2048.0
    y_pixels: 2048.0
    pixel_size_in_microns: 6.5
    subsampling: [1, 2, 4]
    sensor_mode: Normal  # 12 for progressive, 1 for normal.
    readout_direction: Top-to-Bottom  # Top-to-Bottom', 'Bottom-to-Top'
    lightsheet_rolling_shutter_width: 608
    defect_correct_mode: 2.0
    binning: 1x1
    readout_speed: 1.0
```

(continues on next page)

(continued from previous page)

```
    trigger_active: 1.0
    trigger_mode: 1.0 # external light-sheet mode
    trigger_polarity: 2.0  # positive pulse
    trigger_source: 2.0  # 2 = external, 3 = software.
    exposure_time: 20 # Use milliseconds throughout.
    delay_percent: 10
    pulse_percent: 1
    line_interval: 0.000075
    display_acquisition_subsampling: 4
    average_frame_rate: 4.969
    frames_to_average: 1
    exposure_time_range:
      min: 1
      max: 1000
      step: 1
  remote_focus_device:
    hardware:
      name: daq
      type: SyntheticRemoteFocus
      channel: none
      min: 0
      max: 5
    # Optotune EL-16-40-TC-VIS-5D-1-C
    delay_percent: 0
    ramp_rising_percent: 50
    ramp_falling_percent: 2.5
    amplitude: -0.5
    offset: 0
  galvo:
    -
      # shear galvo measured: 351.04 um/V
      name: sheargalvo
      hardware:
        name: daq
        type: NI # SyntheticGalvo
        channel: PCIe-6738/ao12
        min: -5
        max: 5
      # waveform: halfsaw
      waveform: sawtooth
      frequency: 0.5
      amplitude: -1
      offset: 0
      duty_cycle: 50
      phase: 1.57079
    -
      name: xgalvo
      hardware:
        name: daq
        type: NI # SyntheticGalvo
        channel: PCIe-6738/ao0
        min: -5
```

(continues on next page)

(continued from previous page)

```
        max: 5
      waveform: sawtooth
      frequency: 0.5
      amplitude: 0.931
      offset: 0
      duty_cycle: 50
      phase: 1.57079
  stage:
    hardware:
      -
        name: fake_stage
        type: SyntheticStage
        serial_number: 123
        axes: [x,y,theta,z]
      -
        name: snouty_piezo
        type: GalvoNIStage
        serial_number: 124
        axes: [f]
        axes_mapping: [PCIe-6738/ao14]
        min: 0
        max: 10
        volts_per_micron: (10/15.4)*x + 5.0
    y_unload_position: 10000
    y_load_position: 90000

    startfocus: 0
    x_max: 50
    x_min: -50
    y_max: 50
    y_min: -50
    z_max: 500
    z_min: -500
    f_max: 100000
    f_min: -100000
    theta_max: 360
    theta_min: 0

    x_rot_position: 2000
    y_rot_position: 2000
    z_rot_position: 2000

    x_step: 500
    y_step: 500
    z_step: 500
    theta_step: 30
    f_step: 500

    position:
      x_pos: 25250
      y_pos: 40000
      z_pos: 0
```

(continues on next page)

(continued from previous page)

```yaml
      f_pos: 70000
      theta_pos: 0
    velocity: 1000

    x_offset: 0
    y_offset: 0
    z_offset: 0
    f_offset: 0
    theta_offset: 0
  filter_wheel:
    hardware:
      name: filter_wheel
      type: SyntheticFilterWheel
      wheel_number: 1
    filter_wheel_delay: .030 # in seconds
    available_filters:
      FRFP - BLP01-664R-25: 0
      RFP - FF01-598/25-25: 1
      GFP - 527/20: 2
      GFPRFP - ZET488/561m: 3
      Empty-Alignment: 4
      Blocked2: 5
      Blocked3: 6
      Blocked4: 7
      Blocked5: 8
      Blocked6: 9
  lasers:
    # Omicron LightHub Ultra
    # 488 and 640 are LuxX+ Lasers
    # 561 is a Coherent OBIS Laser
    # Digital Laser Outputs
    - wavelength: 488
      onoff:
        hardware:
          name: daq
          type: NI
          channel: PCIe-6738/port1/line2
          min: 0
          max: 5
      power:
        hardware:
          name: daq
          type: NI
          channel: PCIe-6738/ao5
          min: 0
          max: 5
      type: Obis
      index: 0
      delay_percent: 10
      pulse_percent: 100
    - wavelength: 561
      onoff:
```

(continues on next page)

(continued from previous page)

```
          hardware:
            name: daq
            type: NI
            channel: PCIe-6738/port1/line3
            min: 0
            max: 5
        power:
          hardware:
            name: daq
            type: NI
            channel: PCIe-6738/ao11
            min: 0
            max: 5
        type: Obis
        index: 1
        delay_percent: 10
        pulse_percent: 100

  StackingScope:
    daq:
      hardware:
        name: daq
        type: NI

      # NI PCIe-1073 Chassis with PXI-6259 and PXI-6733 DAQ Boards.
      # Sampling rate in Hz
      sample_rate: 100000
      sweep_time: 0.2

      # triggers
      master_trigger_out_line: /PCIe-6738/port0/line1
      camera_trigger_out_line: /PCIe-6738/ctr0 #PFI7 Camera trigger
      trigger_source: /PCIe-6738/PFI0

    mirror:
      hardware:
        name: mirror
        type: ImagineOpticsMirror
        flat_path: D:\WaveKitX64\MirrorFiles\BeadsCoverslip_20231212.wcs
      n_modes: 32

    zoom:
      hardware:
        name: zoom
        type: SyntheticZoom
        servo_id: 1
      position:
        1x: 0
      pixel_size:
        1x: 0.15
    shutter:
      hardware:
```

(continues on next page)

(continued from previous page)

```
      name: shutter
      type: SyntheticShutter
      channel: none/line0
    shutter_min_do: 0
    shutter_max_do: 5
  camera:
    hardware:
      name: camera
      type: HamamatsuOrca
      serial_number: 001301
    x_pixels: 2048.0
    y_pixels: 2048.0
    pixel_size_in_microns: 6.5
    subsampling: [1, 2, 4]
    sensor_mode: Normal  # 12 for progressive, 1 for normal.
    readout_direction: Top-to-Bottom  # Top-to-Bottom', 'Bottom-to-Top'
    lightsheet_rolling_shutter_width: 608
    defect_correct_mode: 2.0
    binning: 1x1
    readout_speed: 1.0
    trigger_active: 1.0
    trigger_mode: 1.0 # external light-sheet mode
    trigger_polarity: 2.0  # positive pulse
    trigger_source: 2.0  # 2 = external, 3 = software.
    exposure_time: 20 # Use milliseconds throughout.
    delay_percent: 10
    pulse_percent: 1
    line_interval: 0.000075
    display_acquisition_subsampling: 4
    average_frame_rate: 4.969
    frames_to_average: 1
    exposure_time_range:
      min: 1
      max: 1000
      step: 1
  remote_focus_device:
    hardware:
      name: daq
      type: SyntheticRemoteFocus
      channel: none
      min: 0
      max: 10
    # Optotune EL-16-40-TC-VIS-5D-1-C
    delay_percent: 7.5
    ramp_rising_percent: 85
    ramp_falling_percent: 2.5
    amplitude: 0.7
    offset: 2.3
  galvo:
    -
      name: sheargalvo
      hardware:
```

(continues on next page)

(continued from previous page)

```
        name: daq
        type: NI # SyntheticGalvo
        channel: PCIe-6738/ao12
        min: -5
        max: 5
      frequency: 0.5
      amplitude: 0
      offset: 0
      duty_cycle: 50
      phase: 1.57079
  stage:
    hardware:
      -
        name: fake_stage
        type: SyntheticStage
        serial_number: 123
        axes: [x,y,theta]
      -
        name: snouty_piezo
        type: GalvoNIStage
        serial_number: 124
        axes: [f]
        axes_mapping: [PCIe-6738/ao14]
        min: 0
        max: 10
        volts_per_micron: (10/15.4)*x + 5.0
      -
        name: z_galvo
        type: GalvoNIStage
        serial_number: 125
        axes: [z]
        # axes_channels: [PCIe-6738/ao0]
        axes_mapping: [PCIe-6738/ao0]
        min: -3.5
        max: 3.5
        volts_per_micron: 0.007*x
    y_unload_position: 10000
    y_load_position: 90000

    startfocus: 0
    x_max: 50
    x_min: -50
    y_max: 50
    y_min: -50
    z_max: 500
    z_min: -500
    f_max: 100000
    f_min: -100000
    theta_max: 360
    theta_min: 0

    x_rot_position: 2000
```

(continues on next page)

(continued from previous page)

```
    y_rot_position: 2000
    z_rot_position: 2000

    x_step: 500
    y_step: 500
    z_step: 500
    theta_step: 30
    f_step: 500

    position:
      x_pos: 25250
      y_pos: 40000
      z_pos: 0
      f_pos: 70000
      theta_pos: 0
    velocity: 1000

    x_offset: 0
    y_offset: 0
    z_offset: 0
    f_offset: 0
    theta_offset: 0
  filter_wheel:
    hardware:
      name: filter_wheel
      type: SyntheticFilterWheel
      wheel_number: 1
    filter_wheel_delay: .030 # in seconds
    available_filters:
      FRFP - BLP01-664R-25: 0
      RFP - FF01-598/25-25: 1
      GFP - 527/20: 2
      GFPRFP - ZET488/561m: 3
      Empty-Alignment: 4
      Blocked2: 5
      Blocked3: 6
      Blocked4: 7
      Blocked5: 8
      Blocked6: 9
  lasers:
    # Omicron LightHub Ultra
    # 488 and 640 are LuxX+ Lasers
    # 561 is a Coherent OBIS Laser
    # Digital Laser Outputs
    - wavelength: 488
      onoff:
        hardware:
          name: daq
          type: NI
          channel: PCIe-6738/port1/line2
          min: 0
          max: 5
```

(continues on next page)

(continued from previous page)

```
        power:
          hardware:
            name: daq
            type: NI
            channel: PCIe-6738/ao5
            min: 0
            max: 5
        type: Obis
        index: 0
        delay_percent: 10
        pulse_percent: 100
      - wavelength: 561
        onoff:
          hardware:
            name: daq
            type: NI
            channel: PCIe-6738/port1/line3
            min: 0
            max: 5
        power:
          hardware:
            name: daq
            type: NI
            channel: PCIe-6738/ao11
            min: 0
            max: 5
        type: Obis
        index: 1
        delay_percent: 10
        pulse_percent: 100

gui:
  channels:
    count: 5
    laser_power:
      min: 0
      max: 100
      step: 10
    exposure_time:
      min: 1
      max: 1000
      step: 5
    interval_time:
      min: 0
      max: 1000
      step: 5
  stack_acquisition:
    step_size:
      min: 0.200
      max: 1000
      step: 0.1
    start_pos:
```

(continues on next page)

(continued from previous page)

```
    min: -5000
    max: 5000
    step: 1
  end_pos:
    min: -5000
    max: 10000
    step: 1
timepoint:
  timepoints:
    min: 1
    max: 1000
    step: 1
  stack_pause:
    min: 0
    max: 1000
    step: 1
confocal_projection:
  scanrange:
    min: 0
    max: 600
    step: 1
  offset_start:
    min: -300
    max: 300
    step: 0.1
  offset_end:
    min: -300
    max: 300
    step: 0.1
  n_plane:
    min: 1
    max: 200
    step: 1
```

##### CT-ASLM-V1

| Equipment | Description |
|---|---|
| Lasers | Coherent Obis lasers with emission at 488, 561, and 642 nm. |
| Stages | MP-285 and Piezo Jena 200-micron piezo for acquisition of z-stacks via sample scanning. |
| Stage Controllers | Sutter MP-285 |
| Cameras | Hamamatsu Flash 4.0 |
| Filter Wheel | Sutter Lambda 10-3 with 1x 25mm Filter Wheel |
| Remote Focusing Units | Equipment Solutions LFA-2010 Linear Focus Actuator |
| Data Acquisition Cards | National Instruments PCIe-6738 |
| Galvo | Novanta CRS 4 KHz Resonant Galvo |
| Zoom | N/A |
| Other | NA |

```yaml
hardware:
  daq:
    type: NI
  camera:
    -
      type: HamamatsuOrca
      serial_number: 000420
  filter_wheel:
    type: SutterFilterWheel
    port: COM34
    baudrate: 9600
    number_of_wheels: 1
  stage:
    -
      type: MP285
      port: COM2
      timeout: 0.25
      baudrate: 9600
      serial_number: 0000
      stages: None
    -
      type: syntheticstage
      port: COM9999
      timeout: 0.25
      baudrate: 9600
      serial_number: 0000
      stages: None
    -
      type: GalvoNIStage
      port: COM9999
      timeout: 0.25
      baudrate: 9600
      serial_number: 0000
      stages: None
  zoom:
    type: synthetic
    servo_id: 1
```

(continues on next page)

(continued from previous page)

```
    port: COM18
    baudrate: 1000000

microscopes:
  CTASLMv1:
    daq:
      hardware:
        name: daq
        type: NI
      sample_rate: 100000
      sweep_time: 0.2

      # triggers
      master_trigger_out_line: PCI6738/port0/line1
      camera_trigger_out_line: /PCI6738/ctr0
      trigger_source: /PCI6738/PFI0

      # Digital Laser Outputs
      laser_port_switcher: PCI6738/port0/line0
      laser_switch_state: False

    camera:
      hardware:
        name: camera
        type: HamamatsuOrca
        serial_number: 000420
      x_pixels: 2048.0
      y_pixels: 2048.0
      pixel_size_in_microns: 6.5
      subsampling: [1, 2, 4]
      sensor_mode: Normal  # 12 for progressive, 1 for normal. Normal/Light-Sheet
      readout_direction: Top-to-Bottom  # Top-to-Bottom', 'Bottom-to-Top'
      lightsheet_rolling_shutter_width: 608
      defect_correct_mode: 2.0
      binning: 1x1
      readout_speed: 1.0
      trigger_active: 1.0
      trigger_mode: 1.0 # external light-sheet mode
      trigger_polarity: 2.0  # positive pulse
      trigger_source: 2.0  # 2 = external, 3 = software.
      exposure_time: 20 # Use milliseconds throughout.
      delay_percent: 2 #10
      pulse_percent: 1
      line_interval: 0.000075
      display_acquisition_subsampling: 4
      average_frame_rate: 4.969
      frames_to_average: 1
      exposure_time_range:
        min: 1
        max: 1000
        step: 1
    remote_focus_device:
```

(continues on next page)

(continued from previous page)

```
    hardware:
      name: remote_focus
      type: EquipmentSolutions # NI
      channel: PCI6738/ao2  #45/46
      comport: COM1
      min: -5
      max: 5
    delay_percent: 7.5
    ramp_rising_percent: 85
    ramp_falling_percent: 2.5
    amplitude: 0.7
    offset: 2.3
    smoothing: 0.0
  galvo:
    -
      hardware:
        name: daq
        type: NI
        channel: PCI6738/ao0  #10/11
        min: -5
        max: 5
      waveform: sawtooth
      frequency: 99.9
      amplitude: 2.5
      offset: 0.5
      duty_cycle: 50
      phase: 1.57079 # pi/2
  filter_wheel:
    hardware:
      name: filter_wheel
      type: SutterFilterWheel
      wheel_number: 1
    filter_wheel_delay: .030 # in seconds
    available_filters:
      445-20: 6
      525-30: 0
      550-49: 9 # switched
      600-53: 7
      665LP: 8
      EMPTY: 1
      BLOCKED1: 2
      BLOCKED2: 3
      BLOCKED3: 4
      BLOCKED4: 5
      # 665LP: 0
      # 550-49: 1
      # 525-30: 2
      # 445-20: 3
      # Blocked1: 4
      # Blocked2: 5
      # Blocked3: 6
      # Blocked4: 7
```

(continues on next page)

(continued from previous page)

```
      # Blocked5: 8
      # Blocked6: 9
  stage:
    hardware:
      -
        name: stage1
        type: MP285
        serial_number: 0000
        axes: [y, x, f]
        axes_mapping: [z, y, x]
        volts_per_micron: None
        axes_channels: None
        max: 25000
        min: 0
      -
        name: stage2
        type: syntheticstage
        serial_number: 0000
        axes: [theta]
        axes_mapping: [theta]
        volts_per_micron: PCI6738/ao0
        # axes_channels: f
        max: 360
        min: 0
      -
        name: stage3
        type: GalvoNIStage
        serial_number: 0000
        axes: [z]
        axes_mapping: [PCI6738/ao6]
        volts_per_micron: 0.02*x
        max: 10
        min: 0
    # joystick_axes: [x, y, f]
    x_max: 12500
    x_min: -12500
    y_max: 12500
    y_min: -12500
    z_max: 500
    z_min: 0
    f_max: 100000
    f_min: -100000
    theta_max: 360
    theta_min: 0

    x_step: 500
    y_step: 500
    z_step: 5
    theta_step: 30
    f_step: 500
    velocity: 1000
```

(continues on next page)

(continued from previous page)

```
      x_offset: 0
      y_offset: 0
      z_offset: 0
      theta_offset: 0
      f_offset: 0
    zoom:
      hardware:
        name: zoom
        type: synthetic
        servo_id: 1
      position:
        16X: 0
      pixel_size:
        16X: 0.425
      stage_positions:
        BABB:
          f:
            16X: 0
    shutter:
      hardware:
        name: daq
        type: synthetic
        channel: PCI6738/port0/line10
        min: 0
        max: 5
    lasers:
      - wavelength: 642
        onoff:
          hardware:
            name: daq
            type: NI
            channel: PCI6738/port1/line5 # 38/35
            min: 0
            max: 5
        power:
          hardware:
            name: daq
            type: NI
            channel: PCI6738/ao8 #16/50 #ao1  # 44/11
            min: 0
            max: 5
        type: Obis
        index: 0
        delay_percent: 10
        pulse_percent: 87
      - wavelength: 561
        onoff:
          hardware:
            name: daq
            type: NI
            channel: PCI6738/port1/line2 # 5/39
            min: 0
```

(continues on next page)

(continued from previous page)

```
          max: 5
      power:
        hardware:
          name: daq
          type: NI
          channel: PCI6738/ao9 #17/50 #ao3  # 12/46
          min: 0
          max: 5
      type: Obis
      index: 1
      delay_percent: 10
      pulse_percent: 87
    - wavelength: 488
      onoff:
        hardware:
          name: daq
          type: NI
          channel: PCI6738/port1/line3 # 6/39
          min: 0
          max: 5
      power:
        hardware:
          name: daq
          type: NI
          channel: PCI6738/ao10 #51/50 #ao4 13/14
          min: 0
          max: 5
      type: Obis
      index: 2
      delay_percent: 10
      pulse_percent: 87
    - wavelength: 405
      onoff:
        hardware:
          name: daq
          type: NI
          channel: PCI6738/port1/line4 # 40/41
          min: 0
          max: 5
      power:
        hardware:
          name: daq
          type: NI
          channel: PCI6738/ao11 #52/18 #ao5  #47/14
          min: 0
          max: 5
      type: Obis
      index: 2
      delay_percent: 10
      pulse_percent: 87

gui:
```

(continues on next page)

(continued from previous page)

```
channels:
  count: 5
  laser_power:
    min: 0
    max: 100
    step: 10
  exposure_time:
    min: 1
    max: 1000
    step: 5
  interval_time:
    min: 0
    max: 1000
    step: 5
stack_acquisition:
  step_size:
    min: 0.100
    max: 1000
    step: 0.1
  start_pos:
    min: -5000
    max: 5000
    step: 1
  end_pos:
    min: -5000
    max: 10000
    step: 1
timepoint:
  timepoints:
    min: 1
    max: 1000
    step: 1
  stack_pause:
    min: 0
    max: 1000
    step: 1
```

##### CT-ASLM-V2

| Equipment | Description |
|---|---|
| Lasers | Coherent Obis lasers with emission at 405, 488, 561, and 642 nm. |
| Stages | Sutter MP-285 and Mad City Lab 500-micron piezo for acquisition of z-stacks via sample scanning. |
| Stage Controllers | Sutter MP-285 |
| Cameras | Hamamatsu Flash 4.0 |
| Filter Wheel | Sutter Lambda 10-3 with 1x 25mm Filter Wheel |
| Remote Focusing Units | Equipment Solutions LFA-2010 Linear Focus Actuator |
| Data Acquisition Cards | National Instruments PCIe-6738 |
| Galvo | Novanta CRS 4 KHz Resonant Galvo |
| Zoom | N/A |
| Other | NA |

```yaml
# Specify all necessary information to find and connect to each hardware
# device that will be used on any of the scopes.
hardware:
  daq:
    type: NI
  camera:
    -
      type: HamamatsuOrca
      serial_number: 302153
  filter_wheel:
    type: SutterFilterWheel
    port: COM10
    baudrate: 9600
    number_of_wheels: 1
  stage:
    -
      type: MP285
      port: COM2
      timeout: 0.25
      baudrate: 9600
      serial_number: 0000
      stages: None
    -
      type: syntheticstage
      port: COM9999
      timeout: 0.25
      baudrate: 9600
      serial_number: 0000
      stages: None
    -
      type: GalvoNIStage
      port: COM9999
      timeout: 0.25
      baudrate: 9600
      serial_number: 0000
      stages: None
  zoom:
```

(continues on next page)

(continued from previous page)

```
    type: synthetic
    servo_id: 1
    port: COM18
    baudrate: 1000000

# Only one microscope can be active in the GUI at a time, but all microscopes will be␣
↪accessible
microscopes:
  CTASLMv2:
    daq:
      hardware:
        name: daq
        type: NI
      sample_rate: 100000
      sweep_time: 0.2

      # triggers
      master_trigger_out_line: PCI6738/port0/line1 #3
      camera_trigger_out_line: /PCI6738/ctr0 #9/42
      trigger_source: /PCI6738/PFI0 #4

      # Digital Laser Outputs
      laser_port_switcher: PCI6738/port0/line0
      laser_switch_state: False

    camera:
      hardware:
        name: camera
        type: HamamatsuOrca
        serial_number: 302153
      x_pixels: 2048.0
      y_pixels: 2048.0
      flip_x: True
      flip_y: False
      pixel_size_in_microns: 6.5
      subsampling: [1, 2, 4]
      sensor_mode: Normal  # 12 for progressive, 1 for normal. Normal/Light-Sheet
      readout_direction: Top-to-Bottom  # Top-to-Bottom', 'Bottom-to-Top'
      lightsheet_rolling_shutter_width: 608
      defect_correct_mode: 1.0
      binning: 1x1
      readout_speed: 2.0
      trigger_active: 1.0
      trigger_mode: 1.0 # external light-sheet mode
      trigger_polarity: 2.0  # positive pulse
      trigger_source: 2.0  # 2 = external, 3 = software.
      exposure_time: 20 # Use milliseconds throughout.
      delay_percent: 20
      pulse_percent: 1
      line_interval: 0.000075
      display_acquisition_subsampling: 4
      average_frame_rate: 4.969
```

(continues on next page)

(continued from previous page)

```
    frames_to_average: 1
    exposure_time_range:
      min: 1
      max: 1000
      step: 1
    x_pixels_step: 4
    y_pixels_step: 4
    x_pixels_min: 4
    y_pixels_min: 4
  remote_focus_device:
    hardware:
      name: remote_focus
      type: EquipmentSolutions # NI
      channel: PCI6738/ao2  #45/46
      comport: COM7
      min: -5
      max: 5
    delay_percent: 7.5
    ramp_rising_percent: 85
    ramp_falling_percent: 5.0
    amplitude: 0.7
    offset: 2.3
    smoothing: 0.0
  galvo:
    -
      hardware:
        name: daq
        type: NI
        channel: PCI6738/ao0  #10/11
        min: -5
        max: 5
      waveform: sawtooth
      frequency: 99.9
      amplitude: 2.5
      offset: 0.5
      duty_cycle: 50
      phase: 1.57079 # pi/2
  filter_wheel:
    hardware:
      name: filter_wheel
      type: SutterFilterWheel
      wheel_number: 1
    filter_wheel_delay: .030 # in seconds
    available_filters:
      Empty-1: 0
      525-30: 1
      600-52: 2
      670-30: 3
      647-LP: 4
      Empty-2: 5
      Empty-3: 6
      Empty-4: 7
```

(continues on next page)

(continued from previous page)

```
    stage:
      hardware:
        -
          name: stage1
          type: MP285
          serial_number: 0000
          axes: [y, x, f]
          axes_mapping: [z, y, x]
          volts_per_micron: None
          axes_channels: None
          max: 25000
          min: 0
        -
          name: stage2
          type: syntheticstage
          serial_number: 0000
          axes: [theta]
          axes_mapping: [theta]
          max: 360
          min: 0
        -
          name: stage3
          type: GalvoNIStage
          serial_number: 0000
          axes: [z]
          axes_mapping: [PCI6738/ao6] #48/49
          volts_per_micron: 0.05*x
          max: 10
          min: 0
          distance_threshold: 20
          settle_duration_ms: 100
      x_max: 12500
      x_min: -12500
      y_max: 12500
      y_min: -12500
      z_max: 200
      z_min: 0
      f_max: 12500
      f_min: -12500
      theta_max: 360
      theta_min: 0

      x_step: 500
      y_step: 500
      z_step: 5
      theta_step: 30
      f_step: 500
      velocity: 1000

      x_offset: 0
      y_offset: 0
      z_offset: 0
```

(continues on next page)

(continued from previous page)

```yaml
      theta_offset: 0
      f_offset: 0
      coupled_axes:
        z: f
    zoom:
      hardware:
        name: zoom
        type: synthetic
        servo_id: 1
      position:
        36X: 0
      pixel_size:
        36X: 0.180
      stage_positions:
        BABB:
          f:
            36X: 0
    shutter:
      hardware:
        name: daq
        type: SyntheticShutter
        channel: PCI6738/port0/line10
        min: 0
        max: 5
    lasers:
      - wavelength: 488
        onoff:
          hardware:
            name: daq
            type: NI
            channel: PCI6738/port1/line5 # 7/41
            min: 0
            max: 5
        power:
          hardware:
            name: daq
            type: NI
            channel: PCI6738/ao8 #1  # 44/11
            min: 0
            max: 5
        type: Obis
        index: 0
        delay_percent: 10
        pulse_percent: 87
      - wavelength: 561
        onoff:
          hardware:
            name: daq
            type: NI
            channel: PCI6738/port1/line2 # 5/39
            min: 0
            max: 5
```

(continues on next page)

(continued from previous page)

```yaml
        power:
          hardware:
            name: daq
            type: NI
            channel: PCI6738/ao9 # 3  # 12/46
            min: 0
            max: 5
        type: Obis
        index: 1
        delay_percent: 10
        pulse_percent: 87
      - wavelength: 647
        onoff:
          hardware:
            name: daq
            type: NI
            channel: PCI6738/port1/line3 # 6/39
            min: 0
            max: 5
        power:
          hardware:
            name: daq
            type: NI
            channel: PCI6738/ao10 # 4  # 13/14
            min: 0
            max: 5
        type: Obis
        index: 2
        delay_percent: 10
        pulse_percent: 87

gui:
  channels:
    count: 5
    laser_power:
      min: 0
      max: 100
      step: 10
    exposure_time:
      min: 1
      max: 1000
      step: 5
    interval_time:
      min: 0
      max: 1000
      step: 5
  stack_acquisition:
    step_size:
      min: 0.100
      max: 1000
      step: 0.1
    start_pos:
```

(continues on next page)

(continued from previous page)

```
      min: -5000
      max: 5000
      step: 1
    end_pos:
      min: -5000
      max: 10000
      step: 1
  timepoint:
    timepoints:
      min: 1
      max: 1000
      step: 1
    stack_pause:
      min: 0
      max: 1000
      step: 1
```

---

##### Spectral TIRF

| Equipment | Description |
|---|---|
| Lasers | Omicron LightHUB Ultra with 405, 457, 488, 514, 532, 561, and 642 nm lasers. |
| Stages | ASI LS-50 linear stage and MS-2000 XY stage. |
| Stage Controllers | ASI Tiger Controller |
| Cameras | 2x Hamamatsu Flash 4.0 |
| Filter Wheel | 2x ASI 6-Position 32 mm Filter Wheels, and 1x motorized ASI dichroic slider. |
| Remote Focusing Units | N/A |
| Data Acquisition Cards | National Instruments PCIe-1073 chassis equipped with PCIe-6259 and PCIe-6738 |
| Galvo | 2x Novanta Linear Galvos. |
| Zoom | N/A |
| Other | NA |

```
# Specify all necessary information to find and connect to each hardware
# device that will be used on any of the scopes.
hardware:
  daq:
    type: NI
  camera:
    -
      type: HamamatsuOrca
      serial_number: 001480
#    -
#      type: HamamatsuOrca
#      serial_number: 001480
```

(continues on next page)

(continued from previous page)

```yaml
  filter_wheel:
    type: ASI
    port: COM17
    baudrate: 115200
    number_of_wheels: 2
  stage:
    -
      type: ASI
      port: COM17
      baudrate: 115200
      controllername: 'C-884'
      stages: L-509.20DG10 L-509.40DG10 L-509.20DG10 M-060.DG M-406.4PD NOSTAGE
      refmode: FRF FRF FRF FRF FRF FRF
      serial_number: 119060508
    -
      type: SyntheticStage
      port: COM17
      baudrate: 115200
      controllername: 'C-884'
      stages: L-509.20DG10 L-509.40DG10 L-509.20DG10 M-060.DG M-406.4PD NOSTAGE
      refmode: FRF FRF FRF FRF FRF FRF
      serial_number: 119060508
  zoom:
    type: synthetic
    servo_id: 1
    port: COM18
    baudrate: 1000000

# Only one microscope can be active in the GUI at a time, but all microscopes will be␣
↪accessible
microscopes:
  SpectralTIRF:
    daq:
      hardware:
        name: daq
        type: NI

      # NI PCIe-1073 Chassis with PXI-6259 and PXI-6733 DAQ Boards.
      # Sampling rate in Hz
      sample_rate: 100000
      sweep_time: 0.2

      # triggers
      master_trigger_out_line: PCIE6738/port0/line1
      camera_trigger_out_line: /PCIE6738/ctr0
      trigger_source: /PCIE6738/PFI0

      # Digital Laser Outputs
      laser_port_switcher: PCIE6738/port0/line0
      laser_switch_state: False

    camera:
```

(continues on next page)

(continued from previous page)

```
      hardware:
        name: camera
        type: HamamatsuOrca
        serial_number: 001480
      x_pixels: 2048.0
      y_pixels: 2048.0
      pixel_size_in_microns: 6.5
      subsampling: [1, 2, 4]
      sensor_mode: Normal  # 12 for progressive, 1 for normal. Normal/Light-Sheet
      readout_direction: Top-to-Bottom  # Top-to-Bottom', 'Bottom-to-Top'
      lightsheet_rolling_shutter_width: 608
      defect_correct_mode: 2.0
      binning: 1x1
      readout_speed: 1.0
      trigger_active: 1.0
      trigger_mode: 1.0 # external light-sheet mode
      trigger_polarity: 2.0  # positive pulse
      trigger_source: 2.0  # 2 = external, 3 = software.
      exposure_time: 20 # Use milliseconds throughout.
      delay_percent: 10
      pulse_percent: 1
      line_interval: 0.000075
      display_acquisition_subsampling: 4
      average_frame_rate: 4.969
      frames_to_average: 1
      exposure_time_range:
        min: 1
        max: 1000
        step: 1
    remote_focus_device:
      hardware:
        name: daq
        type: NI
        channel: PCIE6738/ao2
        min: 0
        max: 5
      # Optotune EL-16-40-TC-VIS-5D-1-C
      delay_percent: 7.5
      ramp_rising_percent: 85
      ramp_falling_percent: 2.5
      amplitude: 0.7
      offset: 2.3
    galvo:
      -
        hardware:
          name: daq
          type: NI
          channel: PCIE6738/ao2  #galvo-x
          min: -5
          max: 5
        waveform: sine
        frequency: 99.9
```

(continues on next page)

(continued from previous page)

```
        period: 10
        amplitude: 2.5
        offset: 0
        duty_cycle: 50
        phase: 0
      -
        hardware:
          name: daq
          type: NI
          channel: PCIE6738/ao3  #galvo-y
          min: -5
          max: 5
        waveform: sine
        frequency: 99.9
        period: 10
        amplitude: 2.5
        offset: 0
        duty_cycle: 50
        phase: 90
    filter_wheel:
      hardware:
        name: emission
        type: ASI
        wheel_number: 1
      filter_wheel_delay: .030 # in seconds
      available_filters:
        CFP - FF01-482/35: 0
        YFP - FF01-250/15: 1
        Blocked 1: 2
        Empty Position 1: 3
        Blocked 2: 4
        Blocked 3: 5
        Empty Position 2: 6
        Blocked 4: 7
        Blocked 5: 8
        Blocked 6: 9
    stage:
      hardware:
        -
          name: ASI
          type: ASI
          serial_number: 119060508
          axes: [x, y, z]
          axes_mapping: [x, y, z]
          volts_per_micron: None
          axes_channels: None
          max: None
          min: None
        -
          name: Synthetic
          type: SyntheticStage
          serial_number: 119060508
```

(continues on next page)

(continued from previous page)

```
        axes: [theta, f]
        axes_mapping: [theta, f]
        volts_per_micron: None
        axes_channels: None
        max: None
        min: None

    x_max: 50000
    x_min: -50000
    y_max: 50000
    y_min: -50000
    z_max: 50000
    z_min: -50000
    f_max: 50000
    f_min: 0
    theta_max: 360
    theta_min: 0

    x_step: 500
    y_step: 500
    z_step: 500
    theta_step: 30
    f_step: 500
    velocity: 1000

  zoom:
    hardware:
      name: zoom
      type: synthetic
      servo_id: 1
    position:
      110x: 0
    pixel_size:
      110x: 0.059
  shutter:
    hardware:
      name: daq
      type: synthetic
      channel: PCIE6738/port0/line0
      min: 0
      max: 5
  lasers:
    - wavelength: 445
      onoff:
        hardware:
          name: daq
          type: NI
          channel: PCIE6738/ao5
          min: 0
          max: 5
      power:
        hardware:
```

(continues on next page)

(continued from previous page)

```
        name: daq
        type: NI
        channel: PCIE6738/ao4
        min: 0
        max: 5
    type: LuxX
    index: 0
    delay_percent: 10
    pulse_percent: 87

  - wavelength: 488
    onoff:
      hardware:
        name: daq
        type: NI
        channel: PCIE6738/ao7
        min: 0
        max: 5
    power:
      hardware:
        name: daq
        type: NI
        channel: PCIE6738/ao6
        min: 0
        max: 5
    type: LuxX
    index: 0
    delay_percent: 10
    pulse_percent: 87

  - wavelength: 514
    onoff:
      hardware:
        name: daq
        type: NI
        channel: PCIE6738/ao9
        min: 0
        max: 5
    power:
      hardware:
        name: daq
        type: NI
        channel: PCIE6738/ao8
        min: 0
        max: 5
    type: LuxX
    index: 0
    delay_percent: 10
    pulse_percent: 87
  - wavelength: 532
    onoff:
      hardware:
```

(continues on next page)

(continued from previous page)

```
          name: daq
          type: NI
          channel: PCIE6738/ao11
          min: 0
          max: 5
      power:
        hardware:
          name: daq
          type: NI
          channel: PCIE6738/ao10
          min: 0
          max: 5
      type: LuxX
      index: 0
      delay_percent: 10
      pulse_percent: 87

    - wavelength: 562
      onoff:
        hardware:
          name: daq
          type: NI
          channel: PCIE6738/ao13
          min: 0
          max: 5
      power:
        hardware:
          name: daq
          type: NI
          channel: PCIE6738/ao12
          min: 0
          max: 5
      type: Obis
      index: 1
      delay_percent: 10
      pulse_percent: 87

    - wavelength: 592
      onoff:
        hardware:
          name: daq
          type: NI
          channel: PCIE6738/ao15
          min: 0
          max: 5
      power:
        hardware:
          name: daq
          type: NI
          channel: PCIE6738/ao14
          min: 0
          max: 5
```

(continues on next page)

(continued from previous page)

```
        type: LuxX
        index: 0
        delay_percent: 10
        pulse_percent: 87

      - wavelength: 642
        onoff:
          hardware:
            name: daq
            type: NI
            channel: PCIE6738/ao17
            min: 0
            max: 5
        power:
          hardware:
            name: daq
            type: NI
            channel: PCIE6738/ao16
            min: 0
            max: 5
        type: LuxX
        index: 2
        delay_percent: 10
        pulse_percent: 87

gui:
  channels:
    count: 5
    laser_power:
      min: 0
      max: 100
      step: 10
    exposure_time:
      min: 1
      max: 1000
      step: 5
    interval_time:
      min: 0
      max: 1000
      step: 5
  stack_acquisition:
    step_size:
      min: 0.200
      max: 1000
      step: 0.1
    start_pos:
      min: -5000
      max: 5000
      step: 1
    end_pos:
      min: -5000
```

(continues on next page)

(continued from previous page)

```
      max: 10000
      step: 1
  timepoint:
    timepoints:
      min: 1
      max: 1000
      step: 1
    stack_pause:
      min: 0
      max: 1000
      step: 1
```

##### Live-Cell ASLM

| Equipment | Description |
|---|---|
| Lasers | Coherent Obis with emission at 405, 457, 488, 514, 561, and 642 nm. |
| Stages | MP-285, PI P-726 PIFOC High-Load piezo, and a galvo for acquisition of z-stacks. |
| Stage Controllers | Sutter MP-285 and PI E-709 |
| Cameras | 2x Hamamatsu Flash 4.0 |
| Filter Wheel | Sutter Lambda 10-3 with 1x 25mm Filter Wheels |
| Remote Focusing Units | Equipment Solutions LFA-2010 Linear Focus Actuator |
| Data Acquisition Cards | National Instruments PCIe-6738 |
| Galvo | Novanta CRS 4 KHz Resonant Galvo |
| Zoom | N/A |
| Other | NA |

```
# Specify all necessary information to find and connect to each hardware
# device that will be used on any of the scopes.
hardware:
  daq:
    type: NI
  camera:
    -
      type: HamamatsuOrca
      serial_number: 100803
#    -
#      type: HamamatsuOrca
#      serial_number: 003054
  filter_wheel:
    type: SutterFilterWheel
    port: COM1
    baudrate: 9600
    number_of_wheels: 1
```

(continues on next page)

(continued from previous page)

```
  stage:
    -
      type: synthetic #MP285
      port: COM6
      timeout: 0.25
      baudrate: 9600
      serial_number: 0001
      stages: None
    -
      type: syntheticstage #syntheticstage
      port: COM9999
      timeout: 0.25
      baudrate: 9600
      serial_number: 0000
      stages: None
    -
      type: GalvoNIStage
      port: COM9999
      timeout: 0.25
      baudrate: 9600
      serial_number: 0000
      stages: None
  zoom:
    type: synthetic
    servo_id: 1
    port: COM18
    baudrate: 1000000

# Only one microscope can be active in the GUI at a time, but all microscopes will be␣
↪accessible
microscopes:
  CTASLMv2:
    daq:
      hardware:
        name: daq
        type: NI
      sample_rate: 100000
      sweep_time: 0.2

      # triggers
      master_trigger_out_line: PCI6738/port0/line1 #3
      camera_trigger_out_line: /PCI6738/ctr0 #9/42
      trigger_source: /PCI6738/PFI0 #4

    camera:
#       -
      hardware:
        name: camera
        type: HamamatsuOrca
        serial_number: 100803
      x_pixels: 2048.0
      y_pixels: 2048.0
```

(continues on next page)

(continued from previous page)

```
      pixel_size_in_microns: 6.5
      subsampling: [1, 2, 4]
      sensor_mode: Normal  # 12 for progressive, 1 for normal. Normal/Light-Sheet
      readout_direction: Top-to-Bottom  # Top-to-Bottom', 'Bottom-to-Top'
      lightsheet_rolling_shutter_width: 50
      defect_correct_mode: 2.0
      binning: 1x1
      readout_speed: 2.0
      trigger_active: 1.0
      trigger_mode: 1.0 # external light-sheet mode
      trigger_polarity: 2.0  # positive pulse
      trigger_source: 2.0  # 2 = external, 3 = software.
      exposure_time: 20 # Use milliseconds throughout.
      delay_percent: 20
      pulse_percent: 1
      line_interval: 0.000075
      display_acquisition_subsampling: 4
      average_frame_rate: 4.969
      frames_to_average: 1
      exposure_time_range:
        min: 1
        max: 1000
        step: 1
#     -
#       hardware:
#         name: camera
#         type: HamamatsuOrca
#         serial_number: 003054
#       x_pixels: 2048.0
#       y_pixels: 2048.0
#       pixel_size_in_microns: 6.5
#       subsampling: [1, 2, 4]
#       sensor_mode: Normal  # 12 for progressive, 1 for normal. Normal/Light-Sheet
#       readout_direction: Top-to-Bottom  # Top-to-Bottom', 'Bottom-to-Top'
#       lightsheet_rolling_shutter_width: 608
#       defect_correct_mode: 2.0
#       binning: 1x1
#       readout_speed: 2.0
#       trigger_active: 1.0
#       trigger_mode: 1.0 # external light-sheet mode
#       trigger_polarity: 2.0  # positive pulse
#       trigger_source: 2.0  # 2 = external, 3 = software.
#       exposure_time: 20 # Use milliseconds throughout.
#       delay_percent: 20
#       pulse_percent: 1
#       line_interval: 0.000075
#       display_acquisition_subsampling: 4
#       display_acquisition_subsampling: 4
#       average_frame_rate: 4.969
#       frames_to_average: 1
#       exposure_time_range:
#         min: 1
```

(continues on next page)

(continued from previous page)

```
#          max: 1000
#          step: 1

    remote_focus_device:
      hardware:
        name: remote_focus
        type: NI #synthetic #EquipmentSolutions # NI
        channel: PCI6738/ao2  #45/46
        comport: COM700
        min: -5
        max: 5
      delay_percent: 7.5
      ramp_rising_percent: 85
      ramp_falling_percent: 5.0
      amplitude: 0.03
      offset: 0.007
      smoothing: 0.0
    galvo:
      -
        hardware:
          name: daq
          type: NI
          channel: PCI6738/ao0  #10/11
          min: -5
          max: 5
        waveform: sawtooth
        frequency: 99.9
        amplitude: 2.5
        offset: 0.5
        duty_cycle: 50
        phase: 1.57079 # pi/2
    filter_wheel:
      hardware:
        name: filter_wheel
        type: SutterFilterWheel
        wheel_number: 1
      filter_wheel_delay: .030 # in seconds
      available_filters:
        647-LP: 0
        600-50: 1
        525-50: 2
        480-40: 3
        Empty-1: 4
        Empty-2: 5
        Empty-3: 6
        Empty-4: 7
    stage:
      hardware:
        -
          name: stage1
          type: synthetic #MP285
          serial_number: 0001
```

(continues on next page)

(continued from previous page)

```
        axes: [y, x, f]
        axes_mapping: [z, y, x]
        max: 25000
        min: 0
      -
        name: stage2
        type: syntheticstage
        serial_number: 0000
        axes: [theta]
        axes_mapping: [theta]
        max: 360
        min: 0
      -
        name: PIFOC
        type: GalvoNIStage
        serial_number: 0000
        axes: [z]
        axes_mapping: [PCI6738/ao6] #48/49
        volts_per_micron: 0.1*x
        max: 10
        min: 0
    x_max: 12500
    x_min: -12500
    y_max: 12500
    y_min: -12500
    z_max: 200
    z_min: 0
    f_max: 12500
    f_min: -12500
    theta_max: 360
    theta_min: 0

    x_step: 500
    y_step: 500
    z_step: 5
    theta_step: 30
    f_step: 500
    velocity: 1000

    x_offset: 0
    y_offset: 0
    z_offset: 0
    theta_offset: 0
    f_offset: 0
  zoom:
    hardware:
      name: zoom
      type: synthetic
      servo_id: 1
    position:
      62.5X: 0.104
    pixel_size:
```

(continues on next page)

(continued from previous page)

```
      62.5X: 0.104
    stage_positions:
      BABB:
        f:
          62.5X: 0
  shutter:
    hardware:
      name: daq
      type: SyntheticShutter
      channel: PCI6738/port0/line10
      min: 0
      max: 5
  lasers:
    - wavelength: 405
      onoff:
        hardware:
          name: daq
          type: NI
          channel: PCI6738/port1/line5 # 7/41
          min: 0
          max: 5
      power:
        hardware:
          name: daq
          type: NI
          channel: PCI6738/ao8 # 16/50
          min: 0
          max: 5
      type: Obis
      index: 0
      delay_percent: 10
      pulse_percent: 87
    - wavelength: 445
      onoff:
        hardware:
          name: daq
          type: NI
          channel: PCI6738/port1/line2 # 5/39
          min: 0
          max: 5
      power:
        hardware:
          name: daq
          type: NI
          channel: PCI6738/ao9 # 17/50
          min: 0
          max: 5
      type: Obis
      index: 1
      delay_percent: 10
      pulse_percent: 87
    - wavelength: 488
```

(continues on next page)

(continued from previous page)

```
      onoff:
        hardware:
          name: daq
          type: NI
          channel: PCI6738/port1/line3 # 6/39
          min: 0
          max: 5
      power:
        hardware:
          name: daq
          type: NI
          channel: PCI6738/ao10 # 51/50
          min: 0
          max: 5
      type: Obis
      index: 2
      delay_percent: 10
      pulse_percent: 87
    - wavelength: 514
      onoff:
        hardware:
          name: daq
          type: NI
          channel: PCI6738/port1/line4 #40/41
          min: 0
          max: 5
      power:
        hardware:
          name: daq
          type: NI
          channel: PCI6738/ao11  #52/18
          min: 0
          max: 5
      type: Obis
      index: 2
      delay_percent: 10
      pulse_percent: 87
    - wavelength: 561
      onoff:
        hardware:
          name: daq
          type: NI
          channel: PCI6738/port1/line6 # 8/42
          min: 0
          max: 5
      power:
        hardware:
          name: daq
          type: NI
          channel: PCI6738/ao12 #53/20
          min: 0
          max: 5
```

(continues on next page)

(continued from previous page)

```
        type: Obis
        index: 2
        delay_percent: 10
        pulse_percent: 87
      - wavelength: 640
        onoff:
          hardware:
            name: daq
            type: NI
            channel: PCI6738/port0/line0 #9/42
            min: 0
            max: 5
        power:
          hardware:
            name: daq
            type: NI
            channel: PCI6738/ao13 #54/20
            min: 0
            max: 5
        type: Obis
        index: 2
        delay_percent: 10
        pulse_percent: 87
gui:
  channels:
    count: 5
    laser_power:
      min: 0
      max: 100
      step: 10
    exposure_time:
      min: 1
      max: 1000
      step: 5
    interval_time:
      min: 0
      max: 1000
      step: 5
  stack_acquisition:
    step_size:
      min: 0.100
      max: 1000
      step: 0.1
    start_pos:
      min: -5000
      max: 5000
      step: 1
    end_pos:
      min: -5000
      max: 10000
      step: 1
  timepoint:
```

(continues on next page)

(continued from previous page)

```
    timepoints:
      min: 1
      max: 1000
      step: 1
    stack_pause:
      min: 0
      max: 1000
      step: 1
```

##### BioFrontiers OPM

| Equipment | Description |
|---|---|
| Lasers | 3i LaserStack with 405, 488, 561, and 642 nm lasers. |
| Stages | ASI FTP-2000 with MS-2000 XY stage, and a Galvo for acquisition of z-stacks. |
| Stage Controllers | ASI Tiger Controller |
| Cameras | Hamamatsu Flash 4.0 |
| Filter Wheel | ASI 8-Position 25 mm Filter Wheel |
| Remote Focusing Units | N/A |
| Data Acquisition Cards | National Instruments PCIe-6723 |
| Galvo | Thorlabs GVS112 Linear Galvo |
| Zoom | N/A |
| Other | NA |

```
# Specify all necessary information to find and connect to each hardware
# device that will be used on any of the scopes.
hardware:
  daq:
    type: NI #SyntheticDAQ or NI
  camera:
    -
      type: HamamatsuOrca
      serial_number: 000646

  filter_wheel:
    type: ASI #SyntheticFilterWheel or ASI
    port: COM13
    baudrate: 115200
    number_of_wheels: 1

  stage:
    -
      type: ASI
      port: COM13
```

(continues on next page)

(continued from previous page)

```
      baudrate: 115200
      serial_number: 123456789
    -
      type: GalvoNIStage
      serial_number: 987654321
      timeout: 0.25
      baudrate: 9600
      stages: None
    -
      type: SyntheticStage
      serial_number: 123
      timeout: 0.25
      baudrate: 9600
      stages: None
  zoom:
    type: SyntheticZoom
    servo_id: 1
    port: COM18
    baudrate: 1000000

microscopes:
  OPM:
    daq:
      hardware:
        name: daq
        type: NI
      sample_rate: 100000
      sweep_time: 0.2
      master_trigger_out_line: PXI6723/port0/line1
      camera_trigger_out_line: /PXI6723/ctr0
      trigger_source: /PXI6723/PFI0
      laser_port_switcher: PXI6723/port0/line0
      laser_switch_state: False

    camera:
      hardware:
        name: camera
        type: HamamatsuOrca
        serial_number: 000646
      lightsheet_rolling_shutter_width: 608
      defect_correct_mode: 2.0
      delay_percent: 10
      pulse_percent: 1
      x_pixels_step: 4
      y_pixels_step: 4
      x_pixels_min: 4
      y_pixels_min: 4
      exposure_time_range:
        min: 1
        max: 1000
        step: 1
      flip_x: False
```

(continues on next page)

(continued from previous page)

```
    flip_y: False

  remote_focus_device:
    hardware:
      name: daq
      type: SyntheticRemoteFocus
      channel: PXI6723/ao2
      min: 0
      max: 5
    delay_percent: 7.5
    ramp_rising_percent: 85
    ramp_falling_percent: 2.5
    amplitude: 0.7
    offset: 2.3
    smoothing: 0.0

  galvo:
    -
      hardware:
        name: daq
        type: NI
        channel: PXI6723/ao0
        min: -10
        max: 10
      waveform: sine
      frequency: 99.9
      amplitude: 2.5
      offset: 0.5
      duty_cycle: 50
      phase: 1.57079

  filter_wheel:
    hardware:
      name: filter_wheel
      type: ASI
      wheel_number: 1
    filter_wheel_delay: .030
    available_filters:
      Empty-Alignment: 0
      GFP - FF01-515/30-32: 1
      RFP - FF01-595/31-32: 2
      Far-Red - BLP01-647R/31-32: 3
      Blocked1: 4
      Blocked2: 5
      Blocked3: 6
      Blocked4: 7
      Blocked5: 8
      Blocked6: 9

  stage:
    hardware:
      -
```

(continues on next page)

(continued from previous page)

```
        name: ASI
        type: ASI
        serial_number: 123456789
        axes: [x, y, f]
        axes_mapping: [X, Y, Z]
        volts_per_micron: None
        axes_channels: None
        max: None
        min: None
      -
        name: GalvoNIStage
        type: GalvoNIStage
        serial_number: 987654321
        axes: [z]
        axes_mapping: [ PXI6723/ao1 ]
        volts_per_micron: 0.05*x
        max: 10
        min: -10
      -
        name: SyntheticStage
        type: SyntheticStage
        serial_number: 123
        axes: [theta]
        axes_mapping: [theta]
        max: 360
        min: 0
    joystick_axes: [x, y, z]
    x_max: 100000
    x_min: -100000
    y_max: 100000
    y_min: -100000
    z_max: 100000
    z_min: -100000
    f_max: 100000
    f_min: -100000
    theta_max: 360
    theta_min: 0
    x_step: 500
    y_step: 500
    z_step: 500
    theta_step: 30
    f_step: 500
    velocity: 1000

    x_offset: 0
    y_offset: 0
    z_offset: 0
    theta_offset: 0
    f_offset: 0

    flip_x: False
    flip_y: False
```

(continues on next page)

(continued from previous page)

```yaml
      flip_z: False

    zoom:
      hardware:
        name: zoom
        type: SyntheticZoom
        servo_id: 1
      position:
        0.63x: 0
        1x: 627
        2x: 1711
        3x: 2301
        4x: 2710
        5x: 3079
        6x: 3383
      pixel_size:
        0.63x: 9.7
        1x: 6.38
        2x: 3.14
        3x: 2.12
        4x: 1.609
        5x: 1.255
        6x: 1.044
      stage_positions:
        BABB:
          f:
            0.63x: 0
            1x: 1
            2x: 2
            3x: 3
            4x: 4
            5x: 5
            6x: 6

    shutter:
      hardware:
        name: daq
        type: SyntheticShutter #NI
        channel: PXI6723/port0/line0
        min: 0
        max: 5

    lasers:
      - wavelength: 405
        onoff:
          hardware:
            name: daq
            type: NI
            channel: PCI6321/port0/line3
            min: 0
            max: 5
        power: #analog
```

(continues on next page)

(continued from previous page)

```yaml
        hardware:
          name: daq
          type: SyntheticLaser
          channel: PCI6321/ao1
          min: 0
          max: 5
      type: LuxX
      index: 0
      delay_percent: 10
      pulse_percent: 87

    - wavelength: 488
      onoff: #digital
        hardware:
          name: daq
          type: NI
          channel: PCI6321/port0/line5
          min: 0
          max: 5
      power: #analog
        hardware:
          name: daq
          type: SyntheticLaser
          channel: PCI6321/ao0
          min: 0
          max: 5
      type: LuxX
      index: 0
      delay_percent: 10
      pulse_percent: 87

    - wavelength: 561
      onoff:
        hardware:
          name: daq
          type: NI
          channel: PCI6321/port0/line7
          min: 0
          max: 5
      power: #analog
        hardware:
          name: daq
          type: SyntheticLaser
          channel: PCI6321/ao1
          min: 0
          max: 5
      type: Obis
      index: 1
      delay_percent: 10
      pulse_percent: 87

    - wavelength: 640
```

(continues on next page)

(continued from previous page)

```
        onoff: #digital
          hardware:
            name: daq
            type: NI
            channel: PCI6321/port0/line1
            min: 0
            max: 5
        power: #analog
          hardware:
            name: daq
            type: SyntheticLaser
            channel: PCI6321/ao2
            min: 0
            max: 5
        type: LuxX
        index: 2
        delay_percent: 10
        pulse_percent: 87

gui:
  channels:
    count: 5
    laser_power:
      min: 0
      max: 100
      step: 10
    exposure_time:
      min: 1
      max: 1000
      step: 5
    interval_time:
      min: 0
      max: 1000
      step: 5
  stack_acquisition:
    step_size:
      min: 0.100
      max: 1000
      step: 0.1
    start_pos:
      min: -5000
      max: 5000
      step: 1
    end_pos:
      min: -5000
      max: 10000
      step: 1
  timepoint:
    timepoints:
      min: 1
      max: 1000
      step: 1
```

(continues on next page)

(continued from previous page)

```
    stack_pause:
      min: 0
      max: 1000
      step: 1
```

CHAPTER

SIX

### ACQUIRING DATA

#### 6.1 Acquiring Data

This provides detailed descriptions of **navigate**'s acquisition modes and saving capabilities. For a how to on acquiring data, please see *Acquiring an Image*.

---

##### 6.1.1 Standard Acquisition Modes

**navigate** features standard acquisition modes including Continuous/Live, Single Frame and Z-Stack, which can be saved to TIFF, H5 and N5 data formats. Saving is toggled under the GUI's *timepoint settings*.

These modes (and other custom modes) can be selected in the program's *acquisition bar* dropdown list.

Each acquisition mode is implemented as a *feature list* and can be used in sequence with other features that can, for example, *make smart decisions*.

---

###### Continuous Scan

This creates a live view of what is on the camera. It is not possible to save data in this mode, only to preview what is in focus. This mode is helpful for alignment, parameter tuning, and scrolling around the sample with the stage.

It is implemented as a *feature list*, shown in its *textual form* below.

```
[
  (
    {"name": PrepareNextChannel},
    {
        "name": LoopByCount,
        "args": ("experiment.MicroscopeState.selected_channels",),
    },
  )
]
```

The sequence begins with the *PrepareNextChannel* feature and loops over *experiment.MicroscopeState.selected_channels*. As such, continuous mode will display a live preview of all *selected color channels* in sequence, then return the first color channel and start again.

---

##### Single Acquisition

This takes a single image of each *selected channel* and optionally saves them to a file. Its feature list is identical to that of "Continuous Scan".

---

##### Z-Stack Acquisition

This takes an image stack over the range and at the step size defined by the *stack acquisition settings* and optionally saves the stack to a file. The color channels will appear as in "Continuous Scan" and "Single Acquisition" if *Laser Cycling Settings* is set to "Per Z" in the stack acquisition settings. A single z-stack will be taken for each color channel, one channel at a time, if *Laser Cycling Settings* is set to "Per Stack".

Z-Stack acquisition is implemented as the feature list below.

```
[
    (
        {"name": ZStackAcquisition},
        {"name": StackPause},
        {
            "name": LoopByCount,
            "args": ("experiment.MicroscopeState.timepoints",),
        },
    )
]
```

Note that in the z-stack the color channel looping is abstracted into `ZStackAcquisition`, but we will take one set of z-stacks at each *timepoint*. It is also worth noting that `ZStackAcquisition` handles moving through *multiple positions*. `ZStackAcquisition` will loop over `Z` or `C` first, as decided by "Per Stack" or "Per Z", and then will loop over positions.

---

##### Customized

The customized acquisition mode can be used to run any feature list of the user's choosing. Data acquisition with **navigate** is almost infinitely reconfigurable with the either the *feature container*, if a desired acquisition can be performed using a reconfiguration of existing features and saving formats, or the *plugin architecture*, if new features or saving formats are required. We strongly recommend the reader check through the available features and see if they can be combined into a acquisition feature list before writing a new acquisition feature.

---

#### 6.1.2 Saving Formats

**navigate** comes pre-packaged with TIFF, OME-TIFF, and H5/N5 (BigDataViewer) file saving formats. The performance of these saving data sources is limited by write speed to disk. To achieve maximal saving speed, we recommend saving all data to a local SSD. See *Hardware Considerations* for more information.

---

###### TIFF/OME-TIFF

**navigate** uses the tifffile package to write TIFF, BigTIFF, and OME-TIFF data to file. The **navigate** package creates a custom OME-TIFF XML to store metadata.

---

###### BigDataViewer H5/N5

**navigate** uses h5py (H5) and zarr (N5) to store data in a BigDataViewer file format. This is a pyramidal format, necessitating the saving of both the original data and down sampled versions of this data. The additional data slows down the write speed. The N5 format is faster than H5 because it allows multithreaded writes.

##### 6.1.3 Image Pipeline

Images are stored from the camera onto a circular buffer of size `databuffer_size`, a setting under `experiment.CameraParameters` in the *software configuration*. By default, this buffer is 100 frames in length.

Image processing and saving operations (see the *feature container* data operations) are performed on frames in this buffer. These operations must take less time than it takes to add a new frame to the buffer, or the buffer will eventually overflow. This is, in part, why saving to an SSD (as opposed to HDD) is critical.

#### 6.2 Reconfigurable Acquisitions Using Features

##### 6.2.1 What are features?

**navigate** allows users to reconfigure acquisition routines within the GUI by chaining so-called "features" together in sequence. A feature is the name given to a single acquisition unit such as `Snap`, which snaps an image, or `MoveToNextPositionInMultiPositionTable`, which moves the stage to the next imaging position indicated in the *Multiposition Table*. Some acquisition units, such as `Autofocus` or `ZStackAcquisition` have a bit more going on under the hood, but can be used in the same way.

---

##### 6.2.2 Customizing Feature Functionality in the GUI

Features can be optionally customized within the GUI. For example instead of reprogramming a feature and loading it again, we can swap Python functions in and out of features. This can be helpful, e.g., when you are prototyping a function to automatically detect an object within an image and want to try a few different options.

---

##### Loading Custom Functions

1. You can load customized functions in the software by selecting the menu *Features* → *Advanced Setting*.

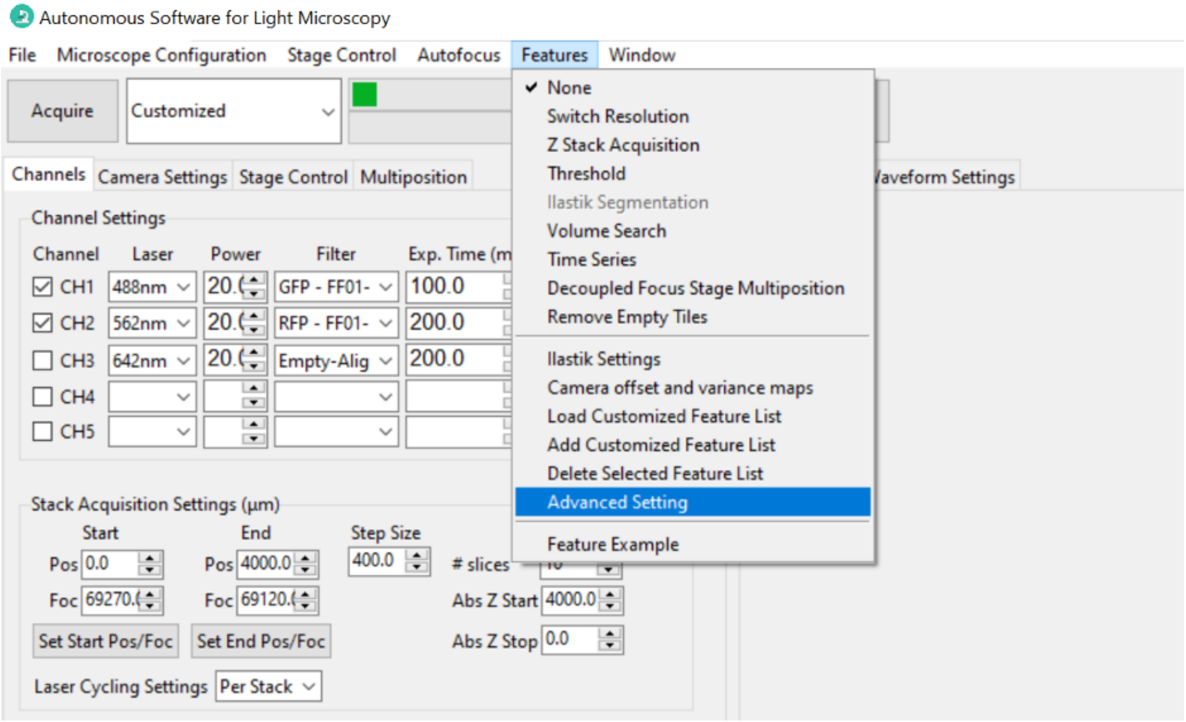

2. In the *Advanced Setting* popup window, choose the feature name with which you want to use the dynamic customized functions as feature parameters.

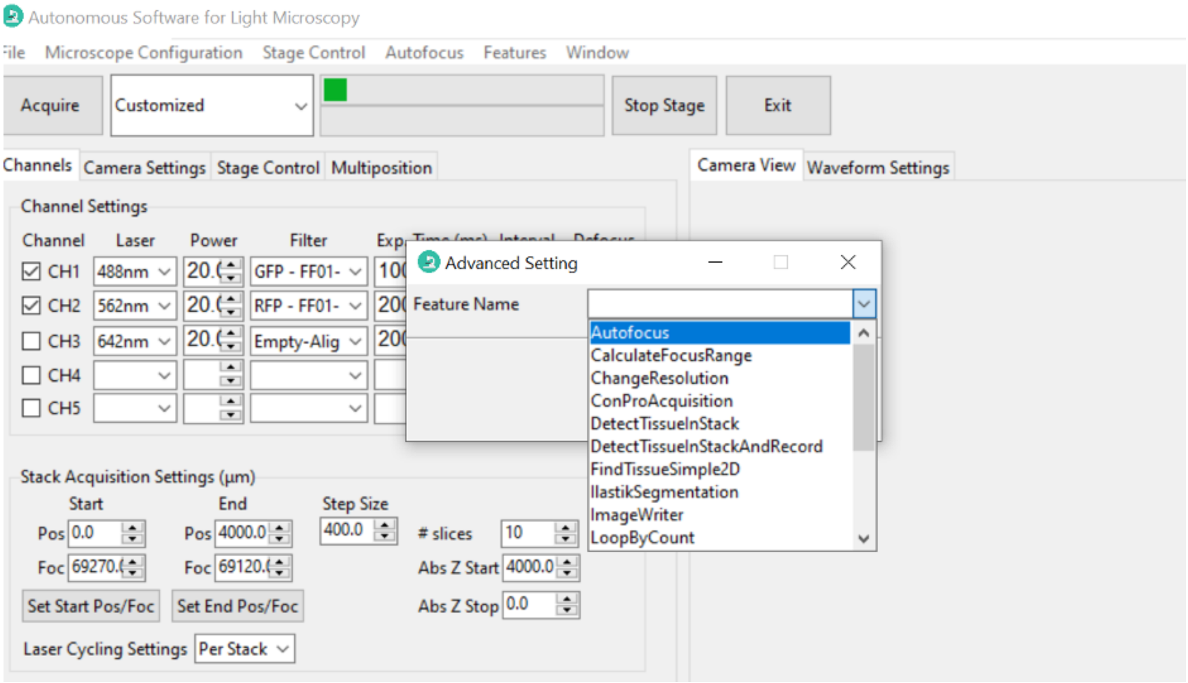

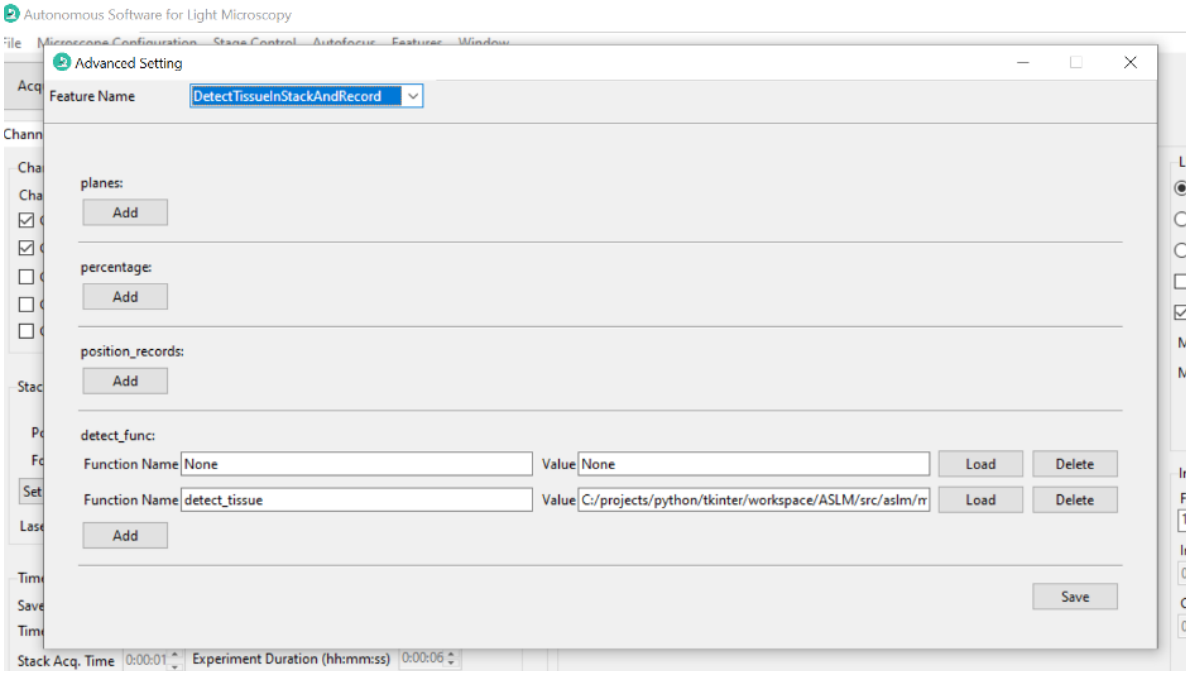

3. Click *Add*, A new line will appear and allow you to edit the parameter options. Type the function name which is defined in your python file.

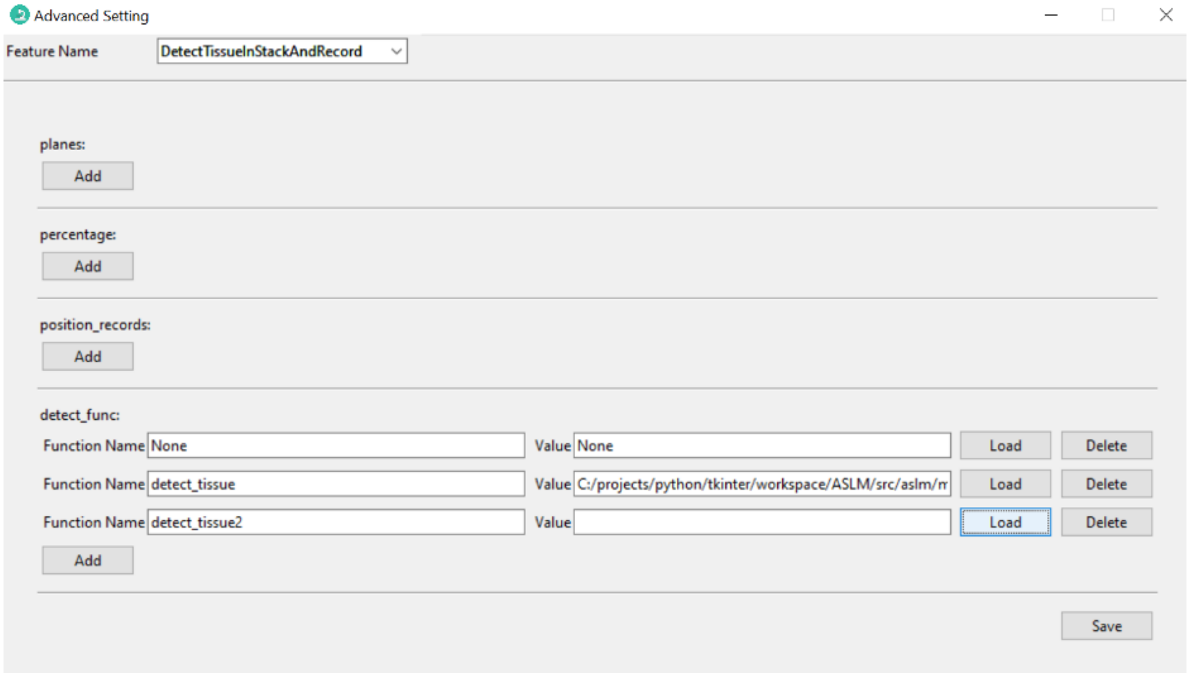

4. Then click *Load* to choose your Python file.

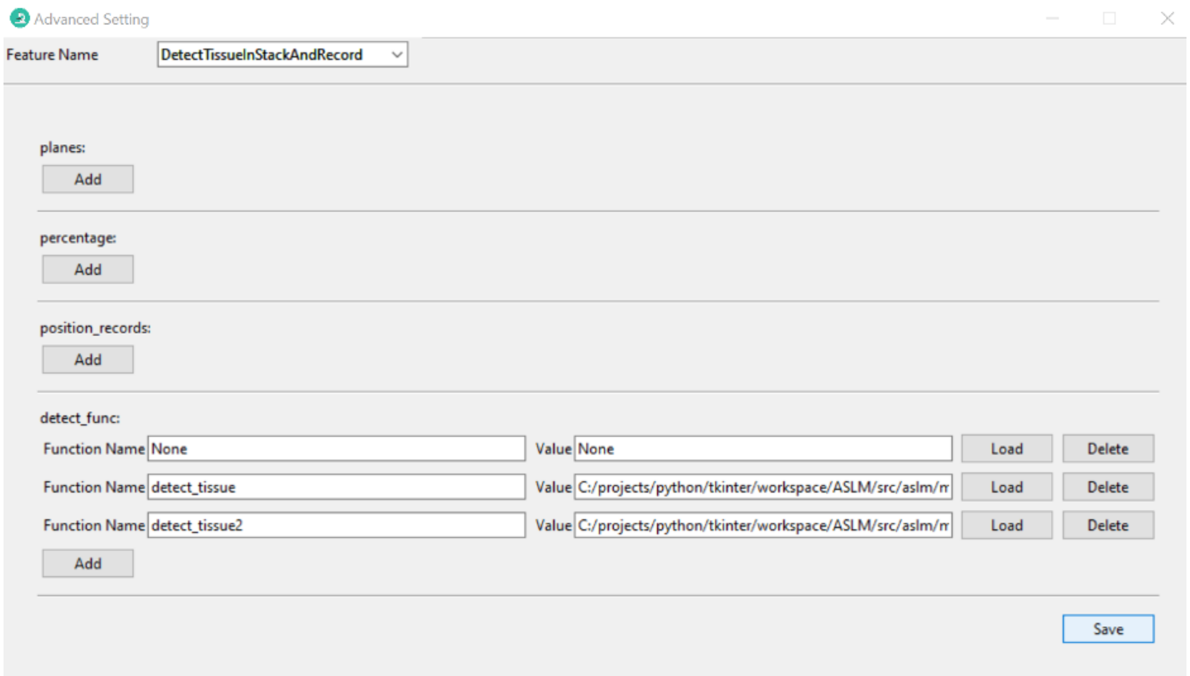

5. When you run a feature list containing the feature you just set, the new function name will appear and you can choose the one you just added.

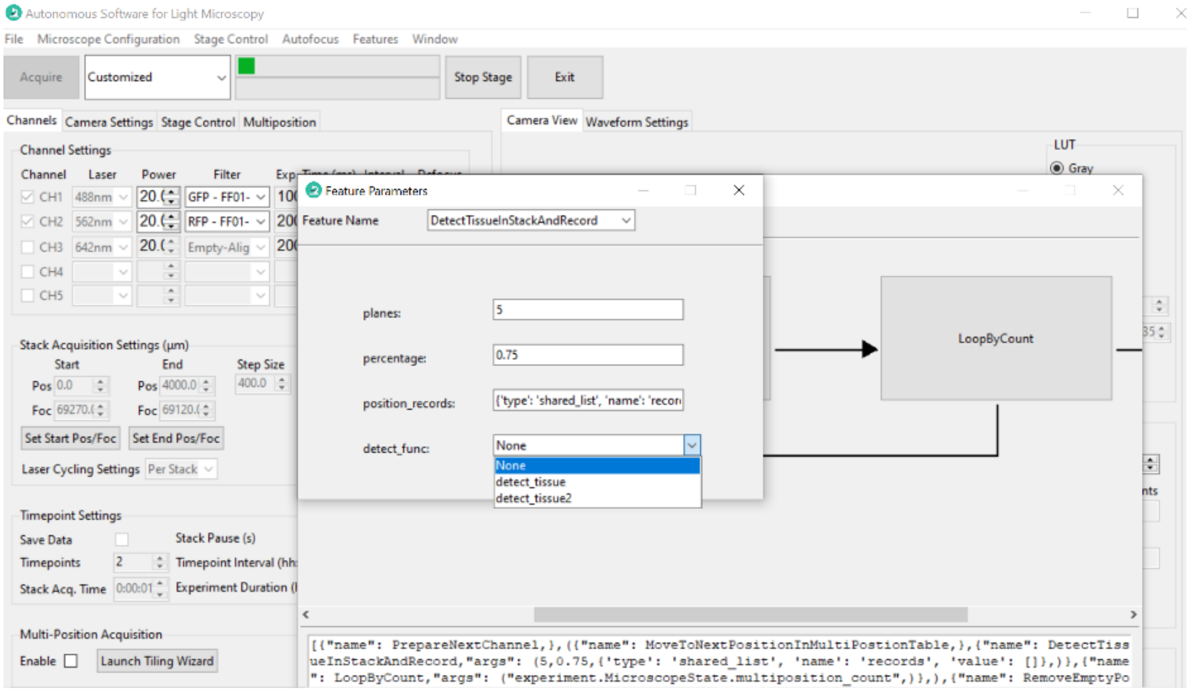

##### 6.2.3 Creating a Custom Feature List in the GUI

Once you have loaded your feature list, the next step is to use it in combination with other features to create an intelligent acquisition workflow. To do this, you will need to create a new feature list that combines your custom feature with other features:

1. Select *Features* → *Add Customized Feature List*. This will open a new dialog box that allows you to create a new feature list.

2. Provide the feature list with a *Feature List Name* of your choice, and type the feature list content (which must be a list object). The feature list content could be the whole feature list or just a simple feature name. In this example, the feature list name is `Feature Example 2`, and the content is a simple feature name:

```
[{"name": PrepareNextChannel}]
```

Once you select *Preview*, the feature list will be displayed in the *Preview* window. If you are satisfied with the feature list, select *OK* to save it.

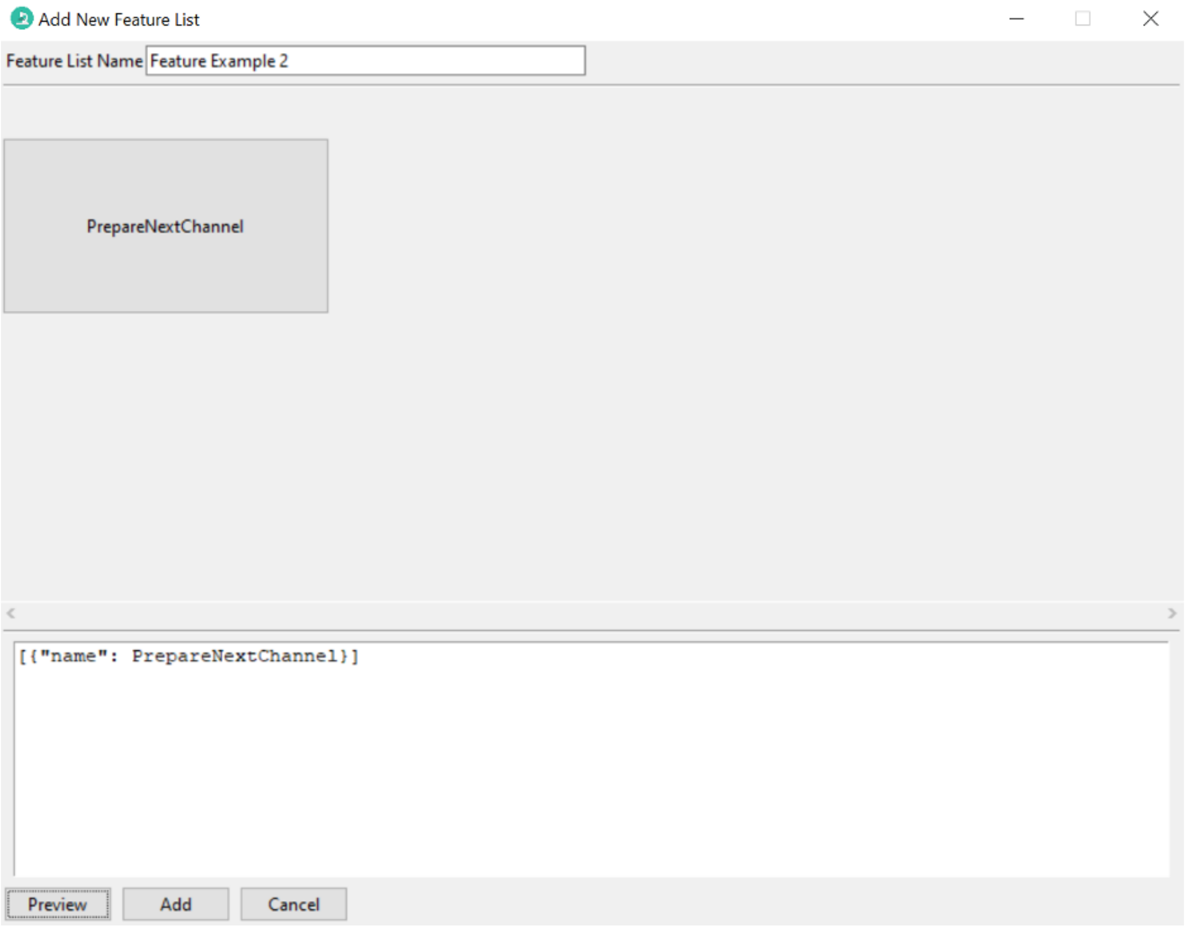

3. You can edit the list of features directly by modifying the text, or through a popup menu that is available by right clicking the feature tile. The popup menu allows you to add a new feature, delete a feature, or edit a feature. In this example, click *Insert After*, and a new feature `PrepareNextChannel` will be inserted by default.

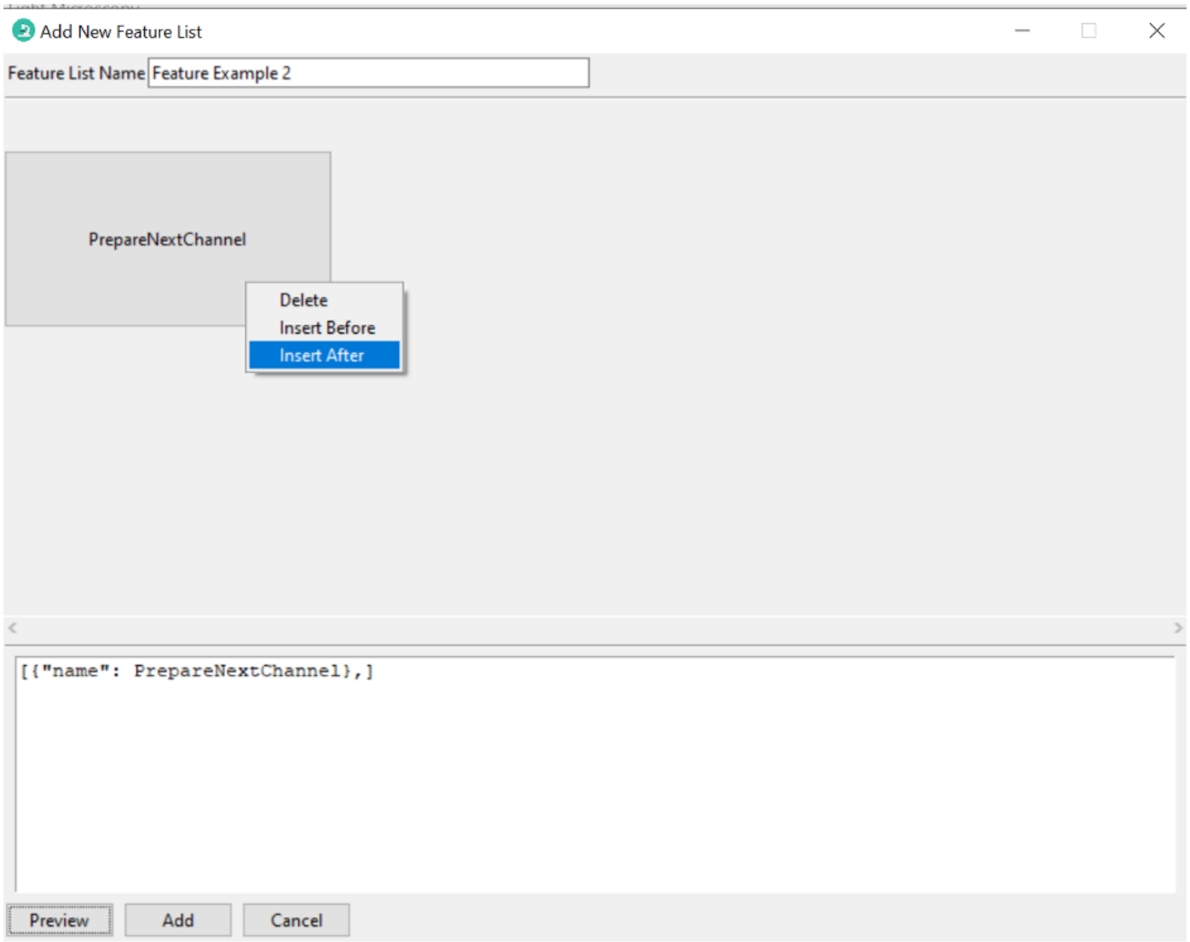
Add New Feature List
Feature List Name
Feature Example 2
PrepareNextChannel
Delete
Insert Before
Insert After
[{"name": PrepareNextChannel},]
Preview
Add
Cancel

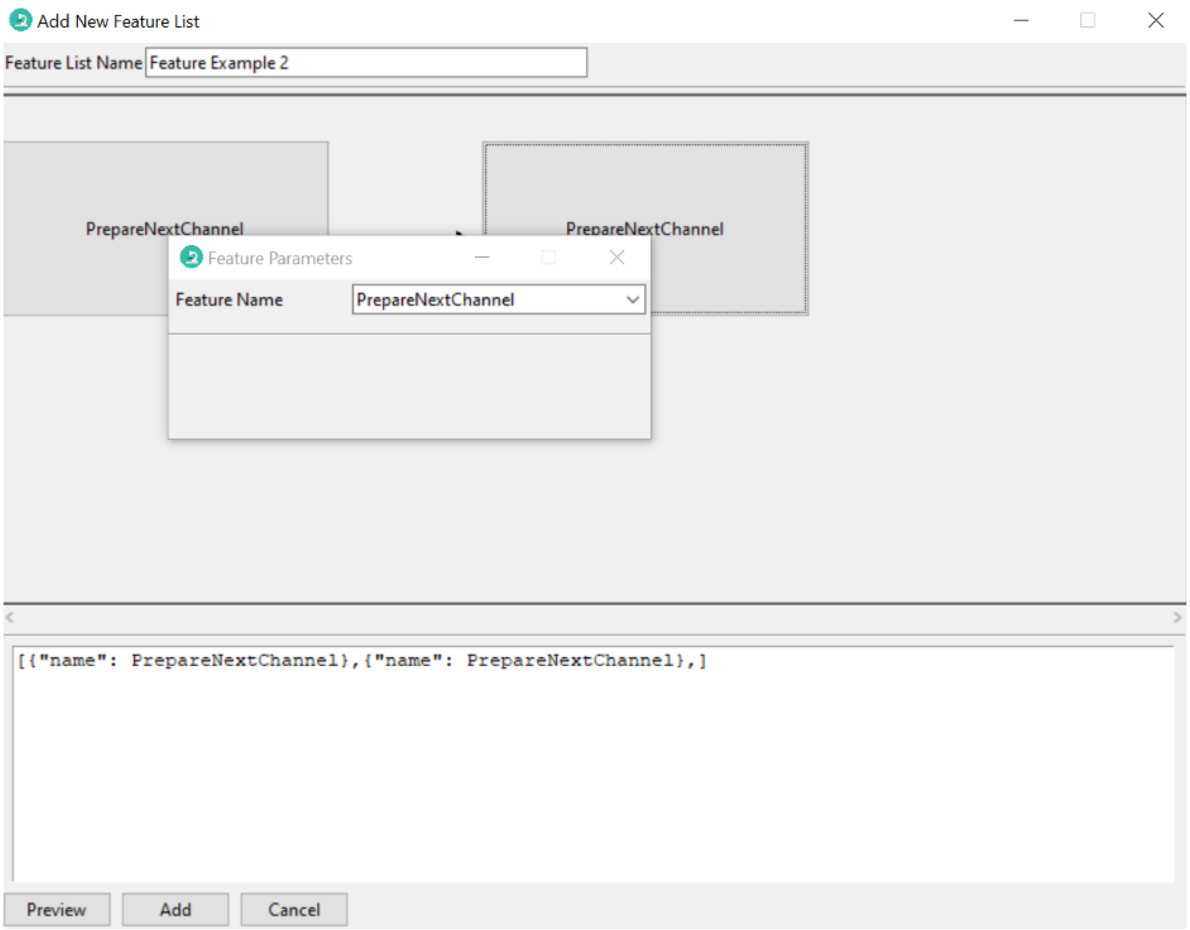

4. To change the identity of the inserted feature, you can select a different feature form the drop-down menu. For example, the feature can be changed from `PrepareNextChannel` to `LoopByCount`. The parameters of the feature can be changed automatically in the popup window.

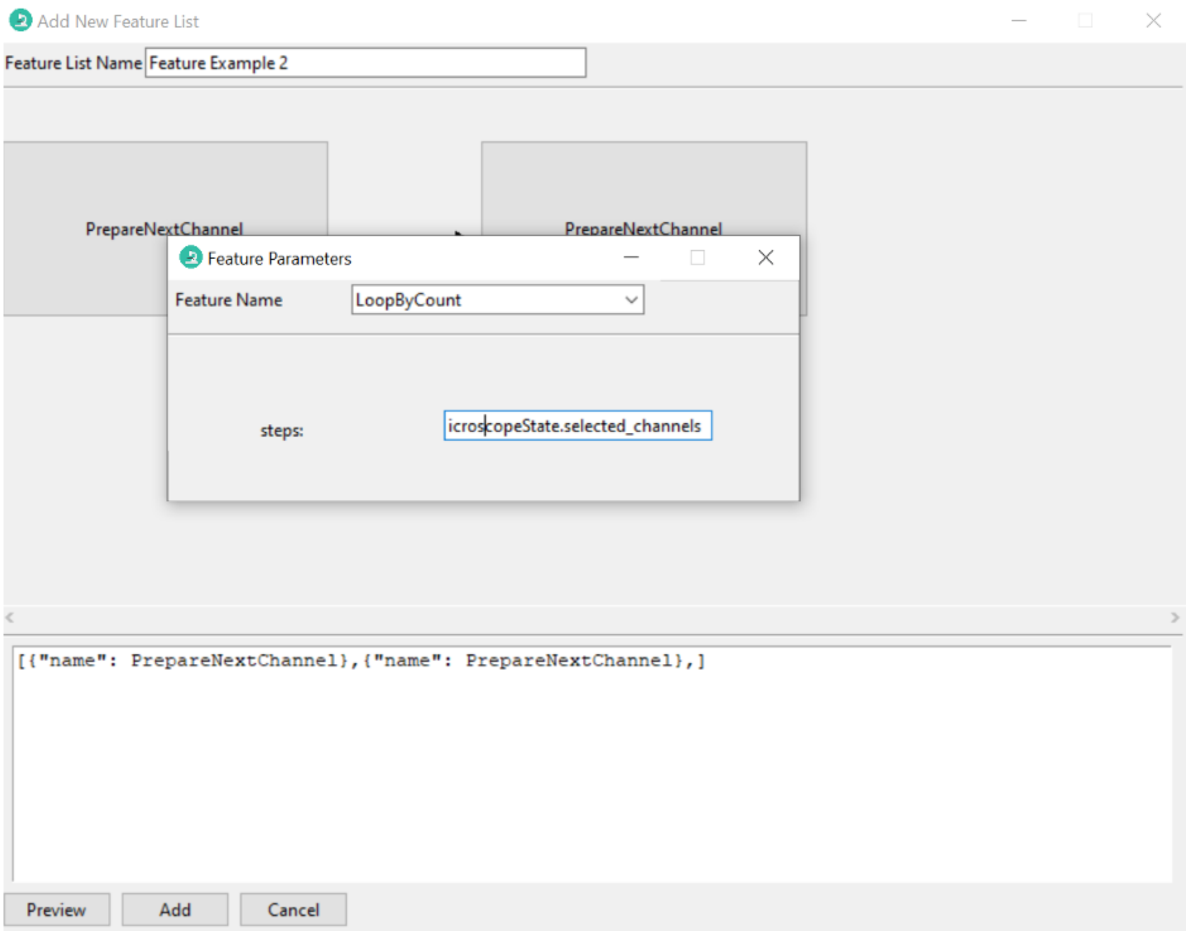

5. If you click the preview button, a graphical representation of the feature list will be displayed.

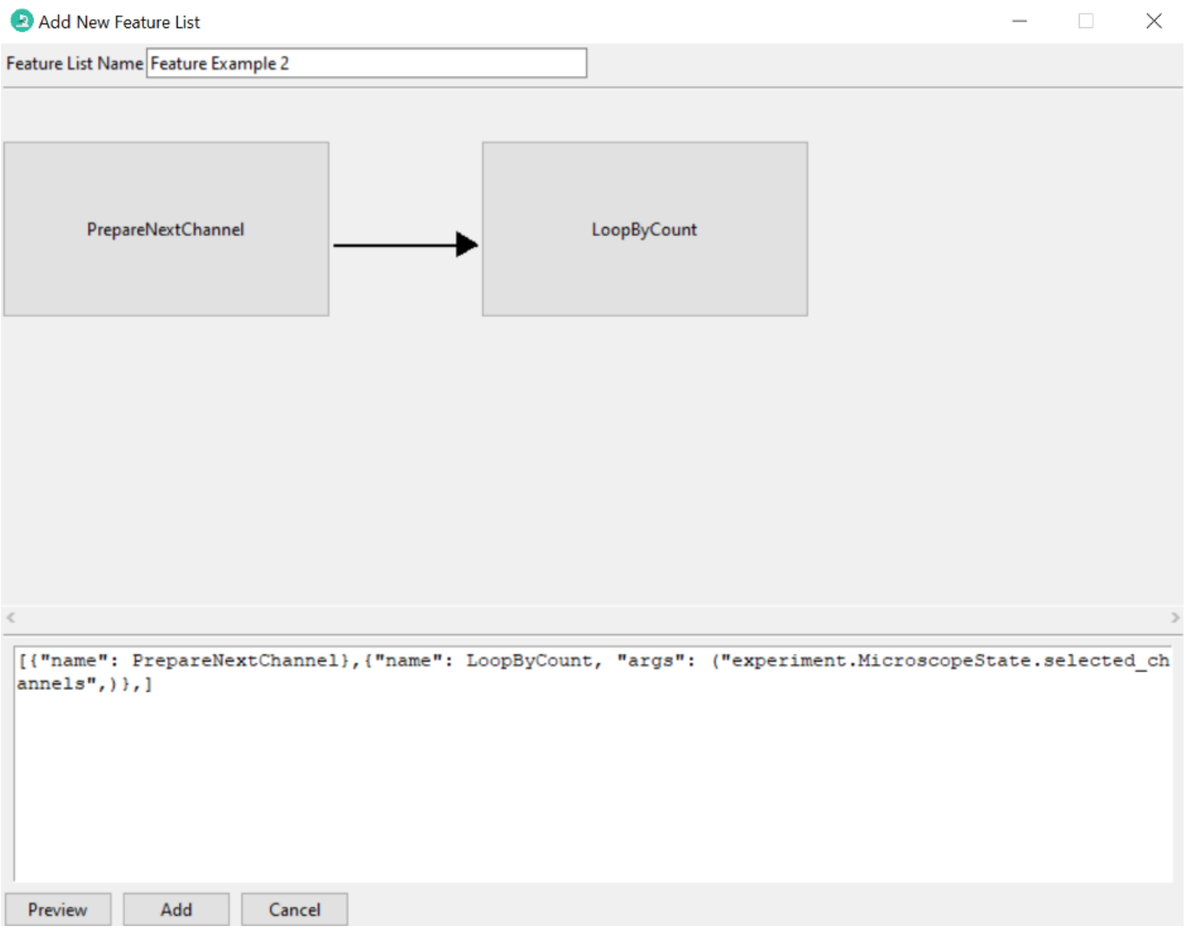

6. If you want a loop structure, type a pair of parentheses around the features, then click *Preview*. Given this design, you can loop through arbitrary features in a user-selected format.

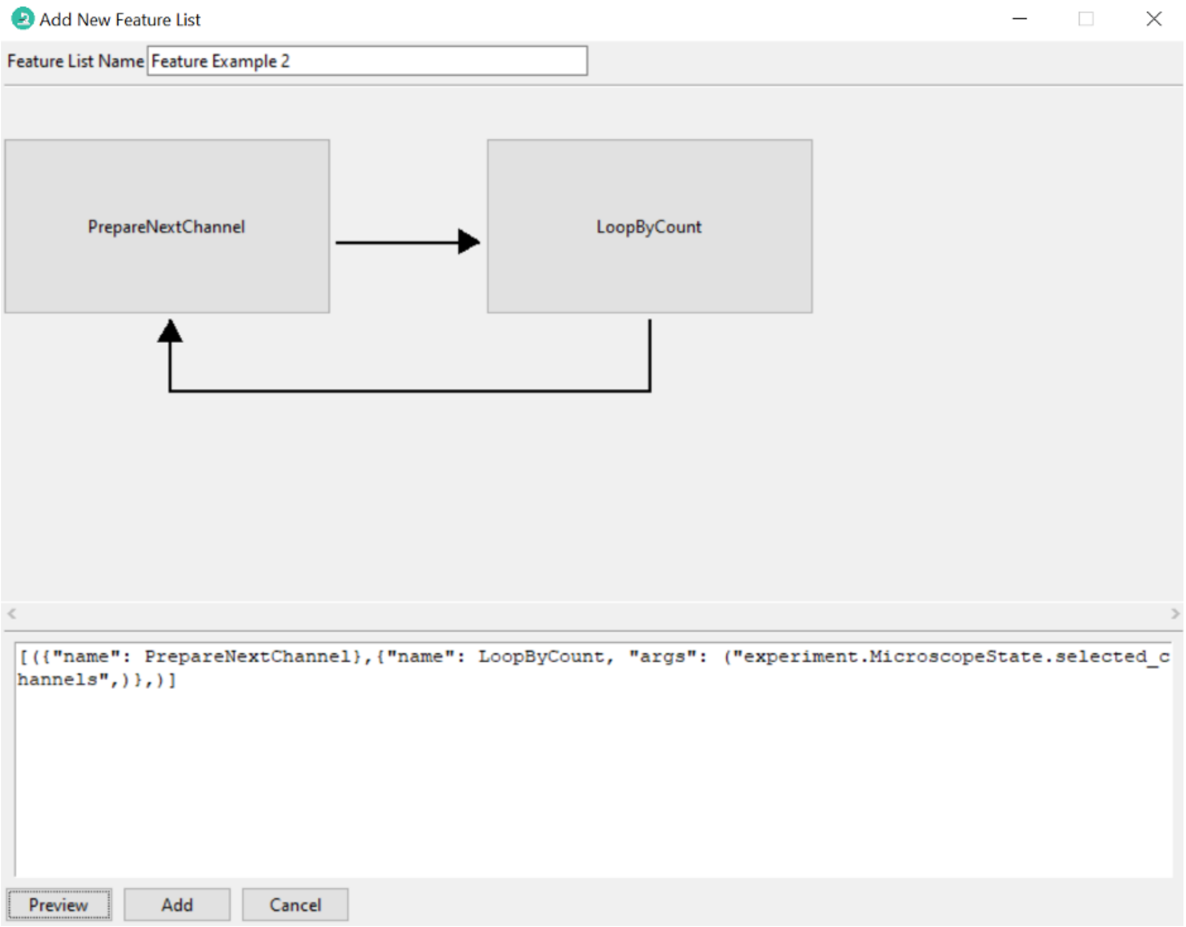

7. After editing the feature list, click *Add*. The new feature list will show up under the *Features* menu.

##### 6.2.4 Editing Feature Lists on the Fly

1. Select the feature list you want to run, choose "Customized" acquisition mode, and then click *Acquire*. A *Feature List Configuration* popup window will show up. In this popup window, you can see the structure of the selected feature list.

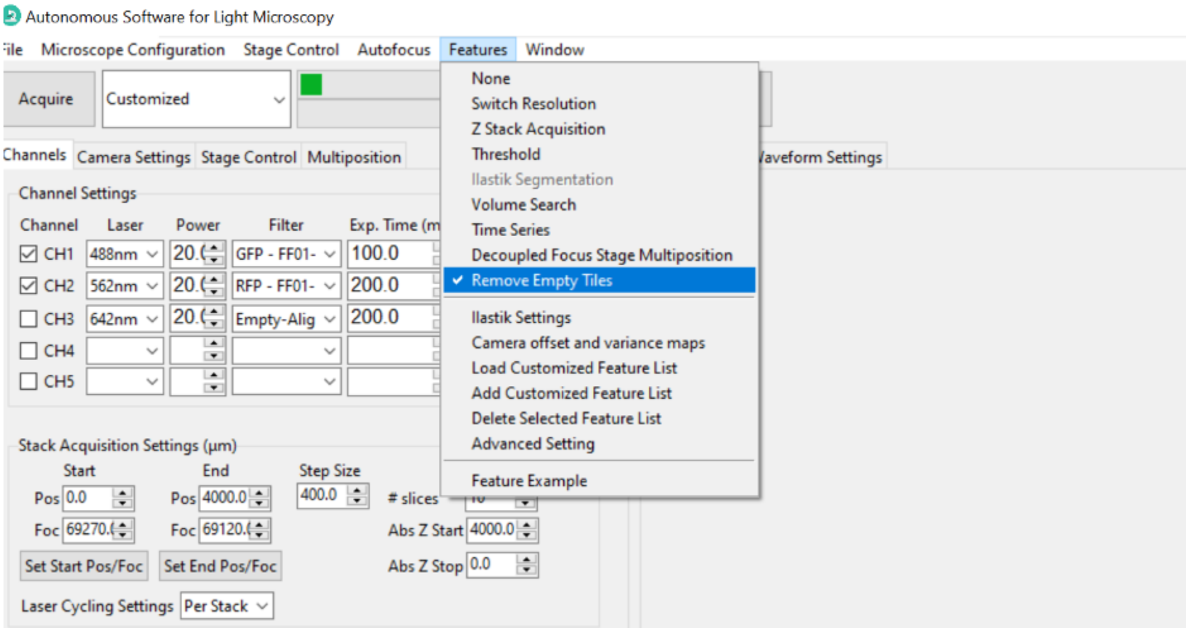
Autonomous Software for Light Microscopy
File Microscope Configuration Stage Control Autofocus Features Window
Acquire
Customized
None
Switch Resolution
Z Stack Acquisition
Threshold
Ilastik Segmentation
Volume Search
Time Series
Decoupled Focus Stage Multiposition
Remove Empty Tiles
Ilastik Settings
Camera offset and variance maps
Load Customized Feature List
Add Customized Feature List
Delete Selected Feature List
Advanced Setting
Feature Example
Channels Camera Settings Stage Control Multiposition
Waveform Settings
Channel Settings
Stack Acquisition Settings (µm)
Set Start Pos/Foc
Set End Pos/Foc
Laser Cycling Settings Per Stack

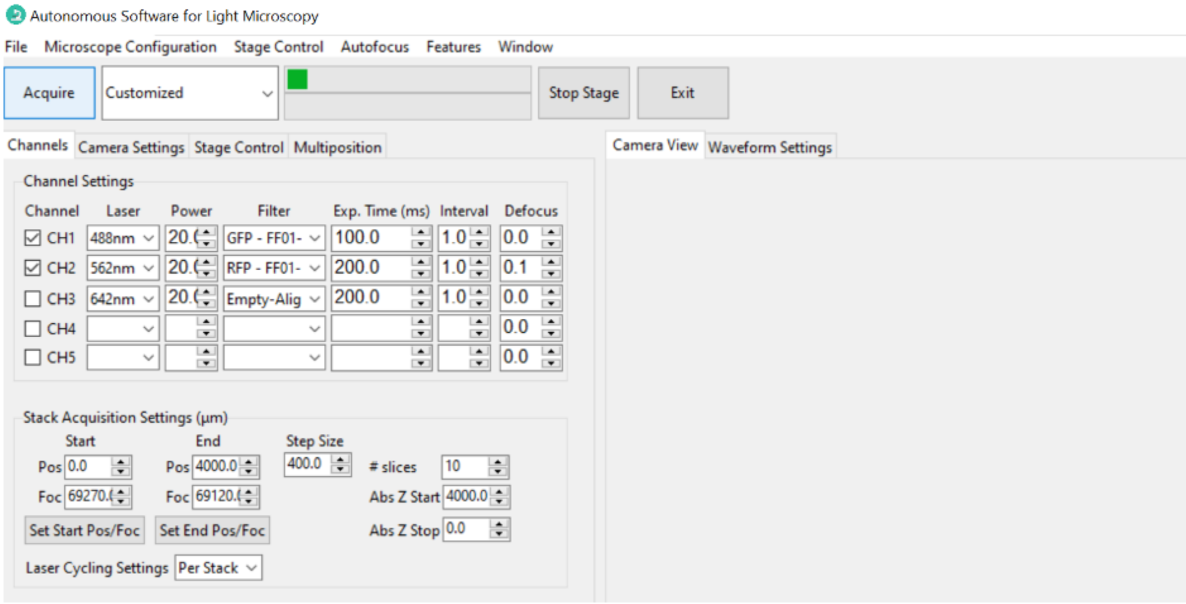
Autonomous Software for Light Microscopy
File Microscope Configuration Stage Control Autofocus Features Window
Acquire
Customized
Stop Stage
Exit
Channels Camera Settings Stage Control Multiposition
Camera View Waveform Settings
Channel Settings
Channel Laser Power Filter Exp. Time (ms) Interval Defocus
Stack Acquisition Settings (µm)
Set Start Pos/Foc
Set End Pos/Foc
Laser Cycling Settings Per Stack

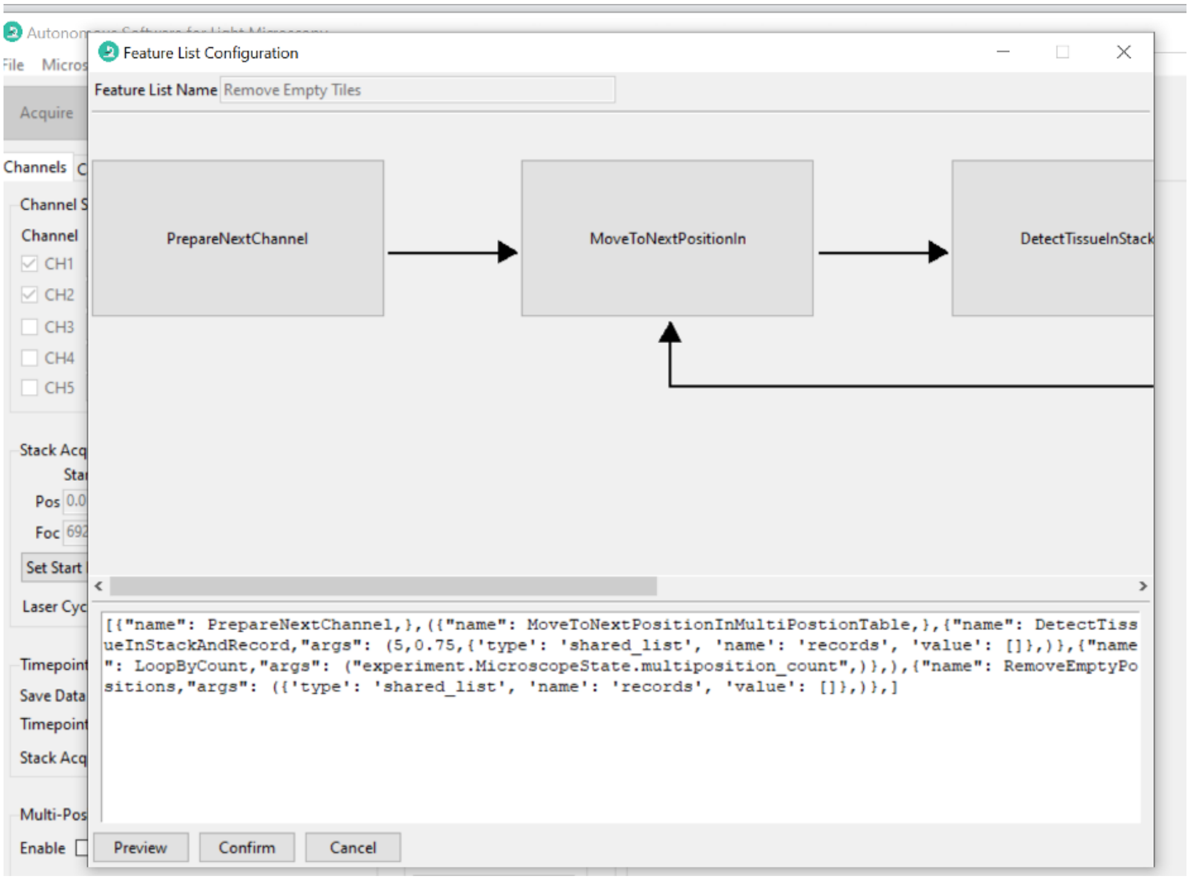

2. Click one feature in the preview window, a *Feature Parameters* window will show up. Then set the desired parameters (e.g., *planes* in this screenshot). Close the *Feature Parameters* window.

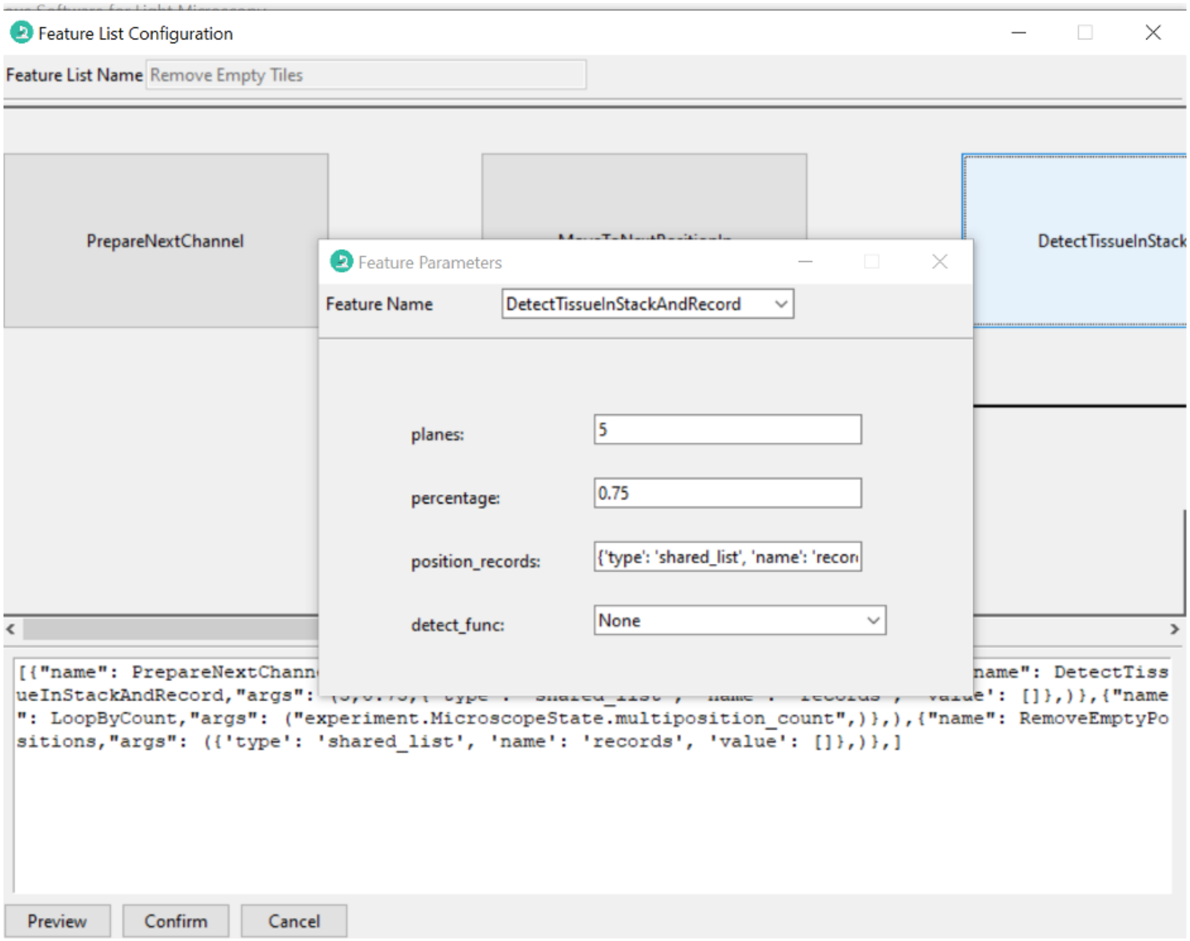

3. Click *Confirm*. The feature list will start to run.

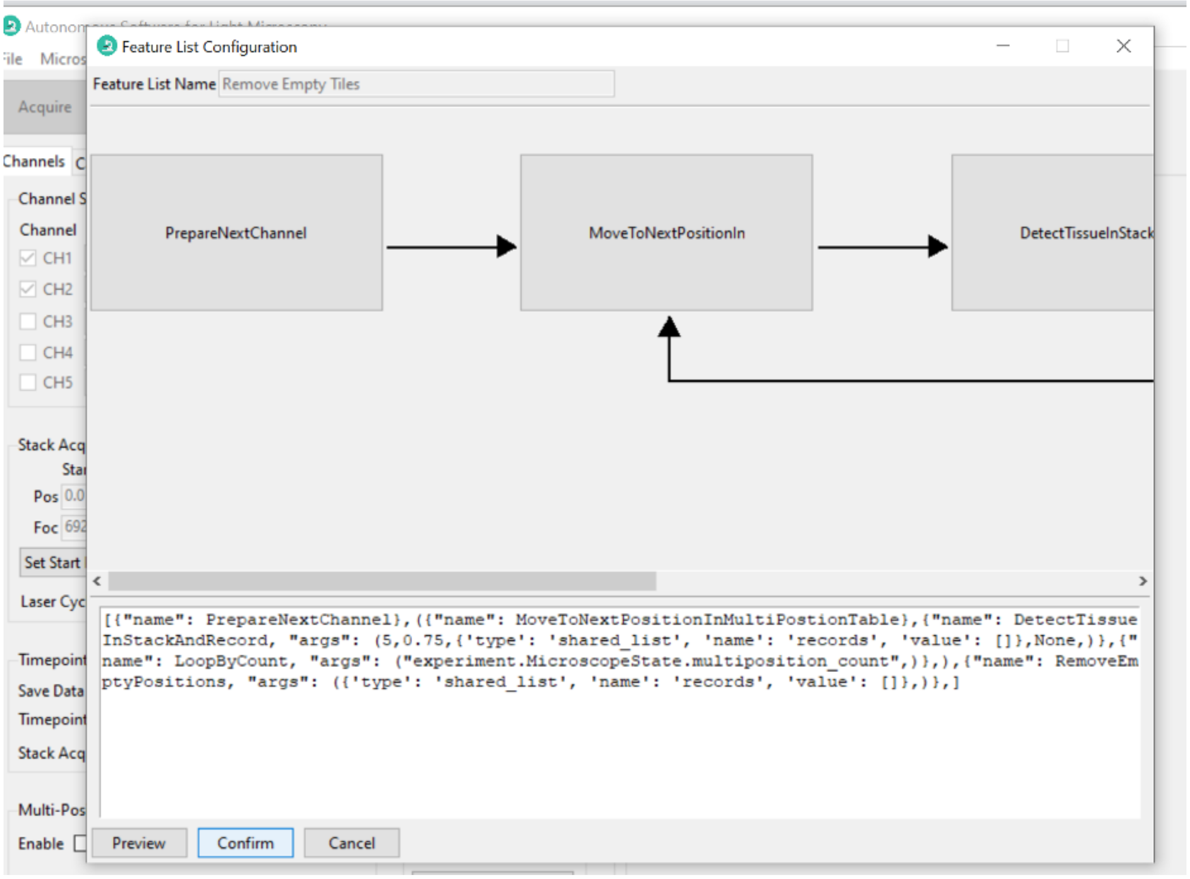

##### 6.2.5 Deleting Feature Lists

1. Select the feature list you want to delete in the *Features* menu.
2. Then, go back to the *Features* menu and select *Delete Selected Feature* The feature list will be removed from the menu and the software.

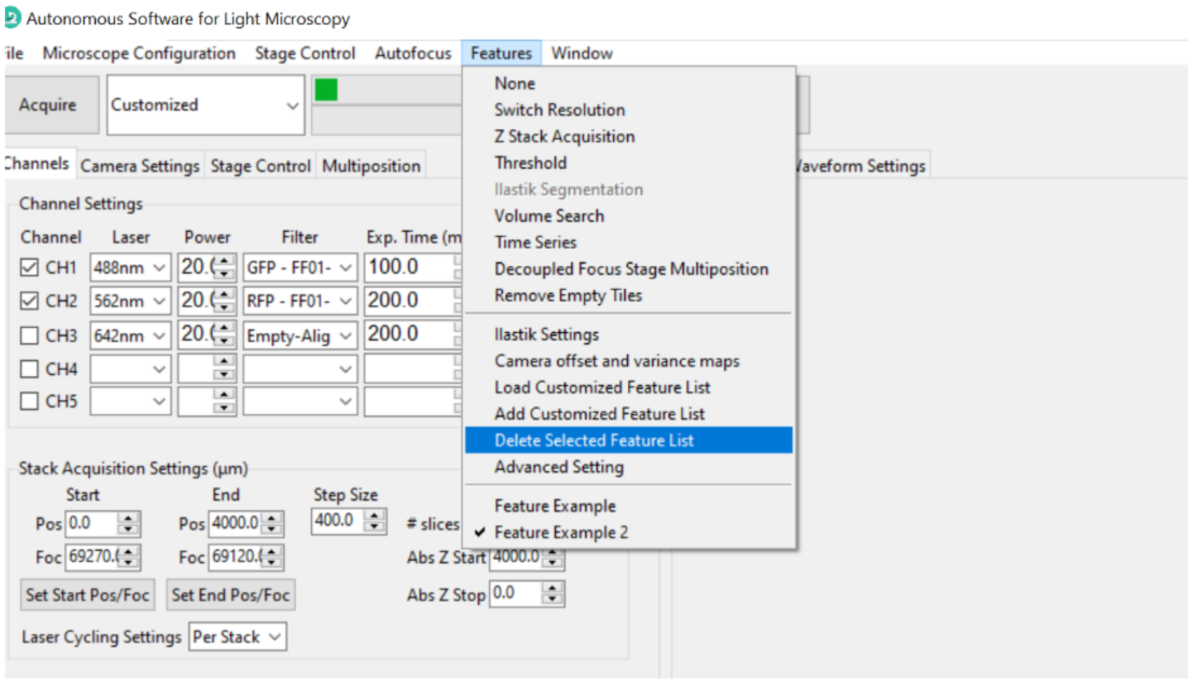

#### 6.2.6 Text Representation of Feature Lists

At the bottom of each of the *Feature List Configuration* frames above, there is a text box with a textual representation of the feature list. As an alternative to point-and-click editing, a user can update feature lists by editing this textual representation and then pressing *Preview*.

The square brackets `[]` create a sequence of events to run in the feature container. The `{}` braces contain features. The parentheses `()` indicate a loop.

As an example, let's look at the feature list that describes the *Continuous Scan* mode:

```
[
  (
    {"name": PrepareNextChannel},
    {
        "name": LoopByCount,
        "args": ("experiment.MicroscopeState.selected_channels",),
    },
  )
]
```

Here, we have a sequence defined by `[]` containing one element, a loop, indicated by the closed parentheses. There are two features within this loop. One feature has the name PrepareNextChannel and the other LoopByCount. The parentheses indicate we will keep looping through both of these features until stopping criteria is met. In this case, the looping will stop when `LoopByCount` returns `False` due to running out of `selected_channels` to loop through. That is, it will end once all *selected channel* have been imaged.

CHAPTER

SEVEN

### CASE STUDIES

Light sheet microscopy is a very versatile technique. Here we present some case studies that demonstrate how **navigate** can be used to acquire data from different types of light sheet microscopes. These include:

- An Axially Swept Light-Sheet Microscope that scans the beam in both the laser propagation (`Y`) and detection (`Z`) directions synchronously with a piezo mounted objective. Tiling in `X`, `Y`, and `Z` is provided by a motorized stage.
- A Digitally Scanned, Axially Swept Light-Sheet Microscope that scans the beam laterally (`X`) with galvanometric mirrors to create a virtual sheet of light, and axially (`Y`) with an electronically tunable lens. The sample is moved in the detection direction (`Z`) to acquire a stack. Tiling in `X`, `Y`, and `Z` is provided by a motorized stage.
- An Axially Swept Light-Sheet Microscope that scans the beam in the laser propagation direction (`Y`), but acquires a stack by moving the sample in the detection direction (`Z`. Tiling in `X`, `Y`, and `Z` is provided by a motorized stage.
- An Axially Swept Light-Sheet Microscope that is configured in an upright, di-SPIM-like, geometry. The beam is scanned in the laser propagation direction (`Y`), but the sample is scanned at a constant velocity in a direction that is 45 degrees to the detection and laser propagation directions (`X`). Acquiring data in this format permits imaging of thinner (e.g., ~2 mm) specimens with very large lateral extents without the overhead associated with stepping and settling the sample stage. Tiling in `X`, `Y`, and `Z` is provided by a motorized stage.

---

#### 7.1 Setting up an Axially Swept Light-Sheet Microscope

##### 7.1.1 Important points

The key to properly setting up the navigate software is to sequentially enable select devices, and troubleshoot each one independently. By carefully and methodically adding devices, and checking that they are functional, one can be confident that the entire system is working as intended. In this example, we will be using a Hamamatsu Flash 4.0 camera. However, you may also use another Hamamatsu sCMOS model, or a Photometrics camera, as long as the drivers are installed and the camera is recognized by the computer. The microscope will operate in a sample scanning format for volumetric image acquisition.

---

##### 7.1.2 First steps

1. Launch the software in synthetic hardware mode. This requires that the conda environment has been established, and that the software has been installed. Once the conda environment has been activated, the software can be launched by typing `navigate -sh` in the terminal.
2. Once the software has been launched, the GUI will appear. Open the folder containing the configuration files by selecting to the menu *File → Open Configuration Files*. This will open the `.navigate` folder in the file explorer, which is where local configuration files are stored.
3. Open the `configuration.yaml` file in your preferred integrated development environment (e.g., PyCharm, VSCode, etc.).
4. For every device in the `hardware` and `microscopes` sections of the `configuration.yaml` file, you will need to change the type to `synthetic`. For convenience, we provide a `synthetic_configuration.yaml` file in `navigate/src/navigate/config` that can be used to replace the `configuration.yaml` file that already has synthetic for each device. You will need to replace the `configuration.yaml` file that is located in the `.navigate` folder with the `synthetic_configuration.yaml` file.
5. Restart the software, but this time launch it in a standard operating mode by typing **navigate** in the terminal. This confirms that the base configuration file is functional. If any problems are encountered, please submit a ticket on GitHub under the "Issues" tab.

---

##### 7.1.3 Sequentially adding devices

###### Data Acquisition Card

1. We will now begin sequentially adding non-synthetic devices to the configuration file. The first device to add is the NI data acquisition card. Of course, the data acquisition card's drivers must be installed and functioning. To confirm that it is functioning, it is best to use the *NI MAX* software and evaluate the card's functionality with an oscilloscope.
2. In the `hardware` and `microscopes` sections of the `configuration.yaml` file, change the `type` to `NI`. You will also need to hard-wire the `master_trigger_out_line` to the `trigger_source`. The identity of these pins can be found in the NI MAX software by right-clicking on the device and selecting *Device Pinouts*. You will also need to make sure that the identity of the pinouts is correct in the `configuration.yaml` file. Most commonly, NI cards default to a device name such as "Dev1". You can change this name in NI MAX, or leave it as is, but whatever you do it has to match the name in the `configuration.yaml` file (e.g., `PXI6259/port0/line1` if the name of the device is "PXI6259" and the pinout is "port0/line1").
3. Open the **navigate** software in the standard operating mode, select the "Continuous Scan" mode, and press *Acquire*. If the software is operating as expected, it should display a synthetically generated image of noise. If it does not, double-check the configuration file and make sure that the `master_trigger_out_line` is connected to the `trigger_source`.

---

##### Camera

1. Next, we will add the camera. The camera must be connected to the computer via a USB or the dedicated frame-grabber cable. The camera drivers must also be installed. The camera drivers can be found on the DCAM-API Website.
2. Update the camera type to `HamamatsuOrca` and input the correct serial number, which can be found on the camera label or through the DCAMConfigurator or HCImage. You will also need to connect the `camera_trigger_out_line` on the data acquisition card to the BNC port labeled "Ext. Trig." on the back of the camera.
3. Restart the software and begin an acquisition in the "Continuous Scan" mode. The camera should now be delivering frames to the software.

---

##### Filter Wheel

1. Set up the Filter Wheel. First, identify the comport via the device manager on Windows. Change the `filter_wheel` type to `SutterFilterWheel` or equivalent in the `hardware` and `microscopes` sections of the configuration file, and provide the necessary information for that filter wheel device (e.g., `baudrate`). At this point, you can also name each filter if desired under the `available_filters` section.
2. Restart the software and select multiple channels in the *Channel Settings* tab, each with different filters. Run the software in "Continuous Scan" mode again and ensure that the filter wheel is changing filters between each acquisition.

---

##### Lasers

1. Set up the lasers. Ideally, the lasers will operate in a mixed modulation mode, which requires that the NI card provides both analog and digital signals to each laser. This allows blanking of the laser, as well as control of its intensity. Open the control software for the lasers, and configure them in a mixed modulation mode. Next, connect the analog output of the NI card to the analog input of the laser. Finally, connect the digital output of the NI card to the digital input of the laser. A common port for the analog and digital outputs are `PXI6733/ao0` and `PXI6733/port0/line2`, respectively.
2. Configure the lasers in the `hardware` and `microscopes` sections of the configuration to type `NI`. Here, you can specify the wavelength of each laser, as well as the minimum and maximum volts to deliver to the laser in both the analog (`power`) and digital (`onoff`) sections.

---

##### Remote focusing unit

1. Configure the Voice Coil. Most voice coils only require an analog signal to control, which can be delivered via the type `NI` in the `hardware` and `microscopes`. However, some voice coils must be configured to accept an analog signal upon each power cycle (e.g., `EquipmentSolutions`). In this case, you will also need to specify the COM port.

---

###### Galvos

1. Set up the galvos. Galvos can be used for a wide variety of tasks, including shadow reduction, digitally scanned light-sheet formation, and also for stepping the beam in z during the acquisition of a z-stack. If the galvo will be used for a z-stack, it should be configured in the `stage` section. All other galvos are placed in `galvo` section. For a sample-scanning ASLM, we use a resonant galvo to perform shadow reduction.

---

###### Stages

1. Install and configure the Stages. You will need to specify stages for the `X`, `Y`, `Z`, `Theta`, and `F` axes. If you do not need one of these stages, it should remain specified as a `SyntheticStage`. It is also important to make sure that you map the stage coordinates to the software coordinates. For example, with the Sutter MP285, the vertical movement of the stage is its z axis. However, for light-sheet microscopes that are laid out horizontally, this axis is the x axis. Thus, we must map the hardware z-axis to the software x-axis. This is done with the `axes` and `axes_mapping` entries, which for the example provided, would be as follows:

```
axes: [x] # software axes
axes_mapping: [z] #hardware axes
```

Importantly, any stage you designate as `Z` will be used for acquisition of a z-stack.

#### 7.2 Imaging on a mesoSPIM BT

This is a case study in using the software to image with a mesoSPIM BT microscope.

---

##### 7.2.1 Setting the beam parameters

Make sure the imaging chamber is empty or, if a sample is mounted, the sample is not in the beam path.

1. Select "Continuous Scan" from the dropdown next to the *Acquire* button. Press *Acquire*. This will launch a live acquisition mode.
2. Go to the *Channels* tab. Choose the wavelength you want to align. Set the laser's *Power* to `100.0`. Change *Filter* to an "Empty" option.

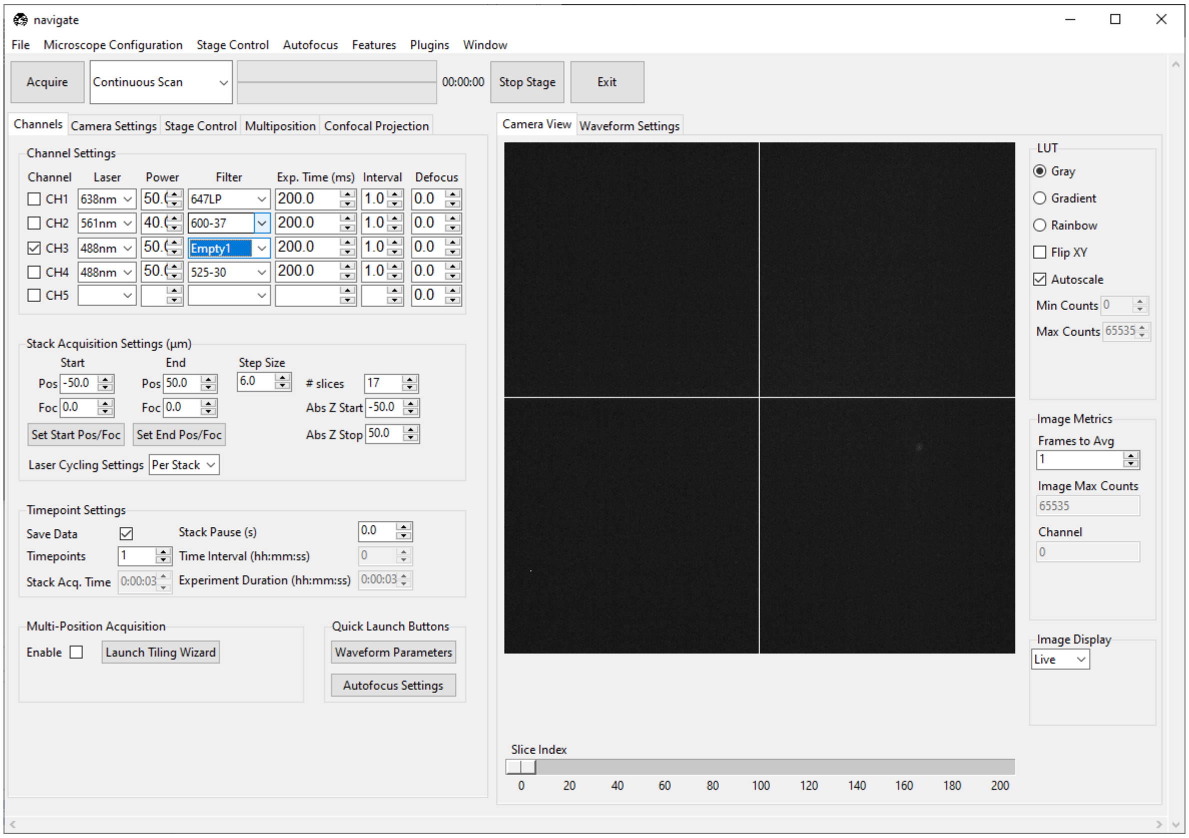

3. Go to the *Microscope Configuration* → *Waveform Parameters*. A popup named *Waveform Parameter Settings* will appear. Make sure the *Mode* matches "mesoSPIM BT" and the *Magnification* matches the magnification of the objective you are using.

---

**Note:** The *Mode* is the name of the microscope as defined in the `configuration.yaml` file. Likewise, the *Magnification* is the magnification(s) for that microscope as defined in the `configuration.yaml` file.

The mesoSPIM BT largely operates at a fixed magnification. However, the original mesoSPIM used a variable magnification research grade macro zoom microscope.

---

1. *Galvo 0* digitally sweeps the beam across the field of view in the `X` direction. To align the axially-swept light sheet parameters, set the *Galvo 0 Amplitude* to `0.0`.

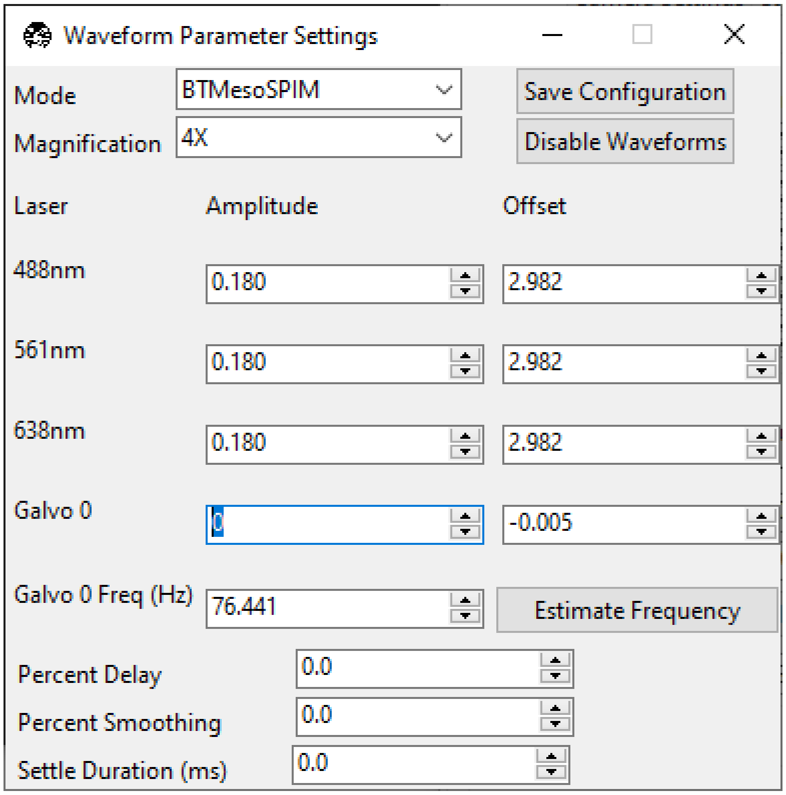

2. The empty filter makes us susceptible to seeing particles scattering light in the chamber. This can effect the software's autoscaling routine. To ensure we are looking at the beam correctly, uncheck *Autoscale* and set the *Min Counts* and *Max Counts* so the beam is visible, but not saturating the display.
3. Set the wavelength's *Amplitude* to `0.0`. Set the wavelength's *Offset* so that the beam is focused in the center of the field of view.

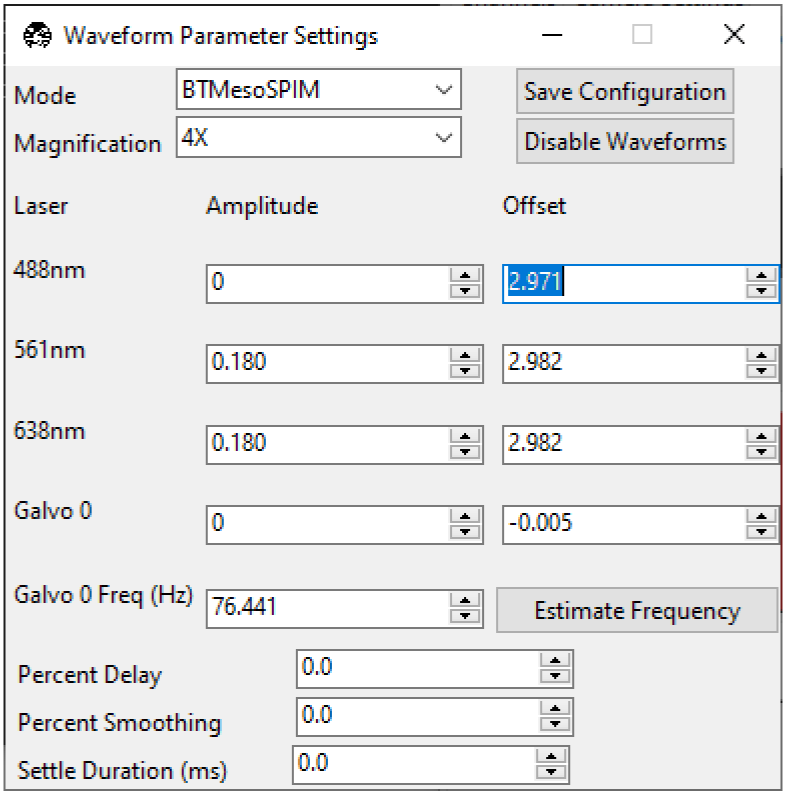

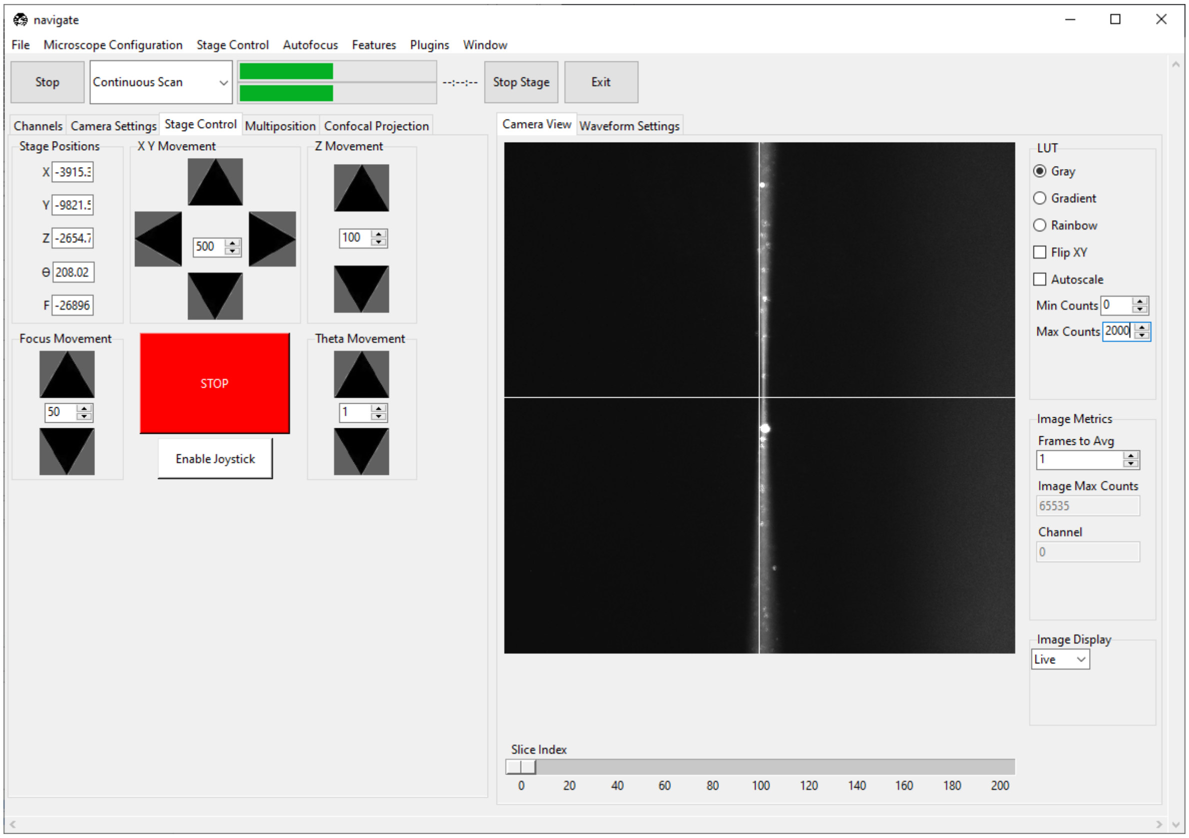

4. Set the *Galvo 0 Offset* so that the beam is centered in the field of view. Click the *Camera View* to toggle the crosshairs, which indicate the center of the field of view.

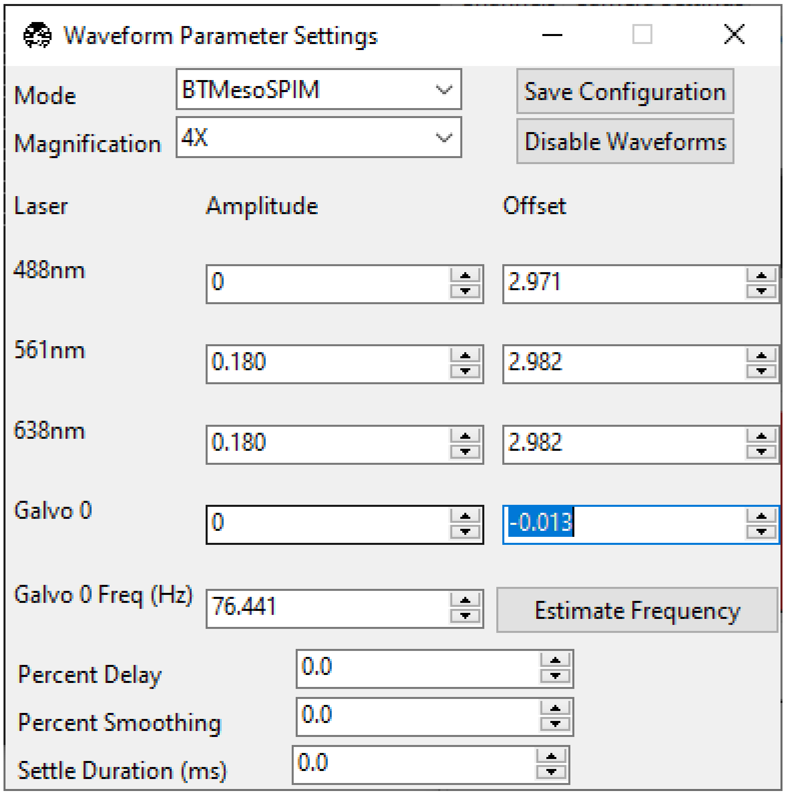

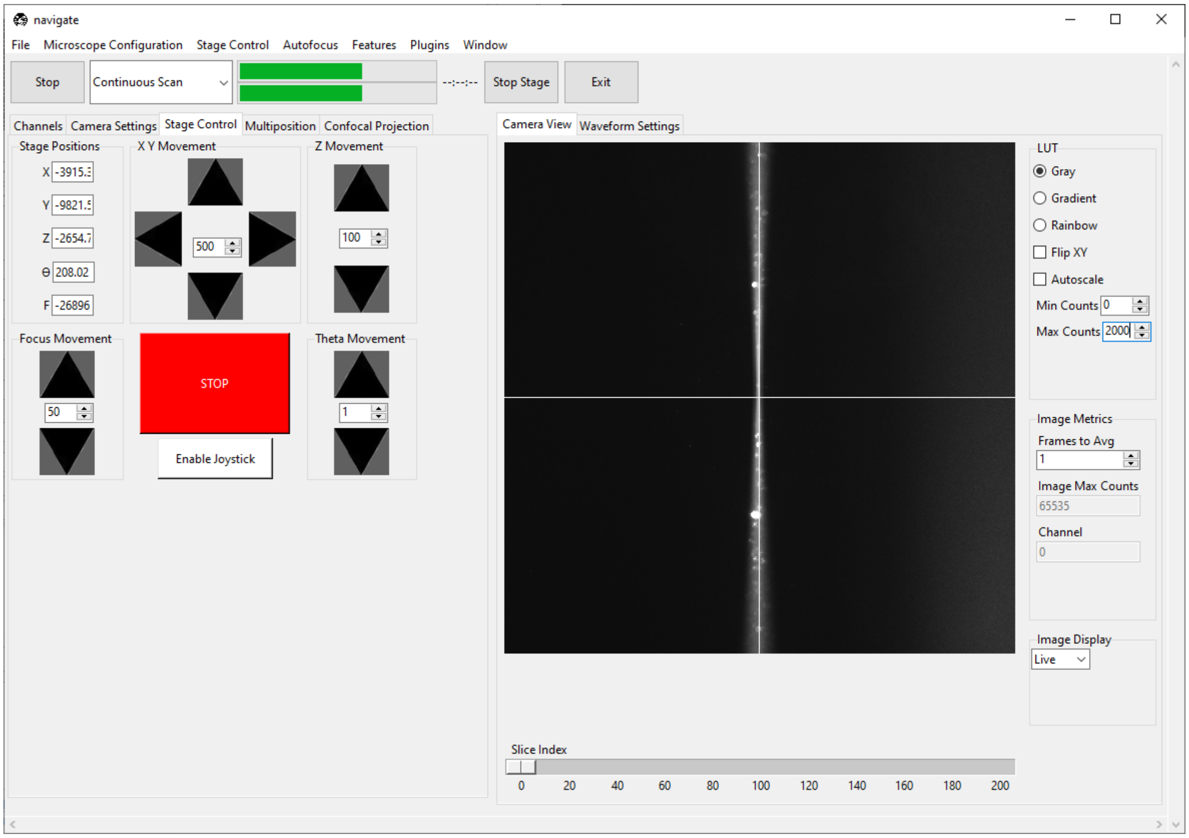

5. Go to *Camera Settings* and ensure that *Light-Sheet* is selected under *Sensor Mode*. Slowly increase the wavelength's *Amplitude* until the beam becomes a straight line across the screen. If the beam does not become

straighter, try changing the camera's *Readout Direction*.

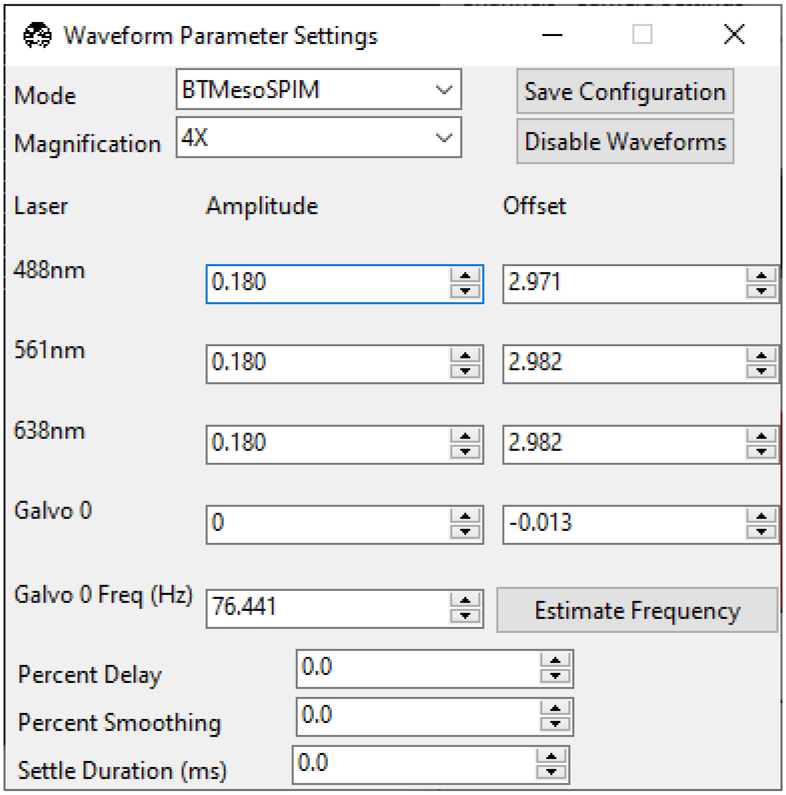

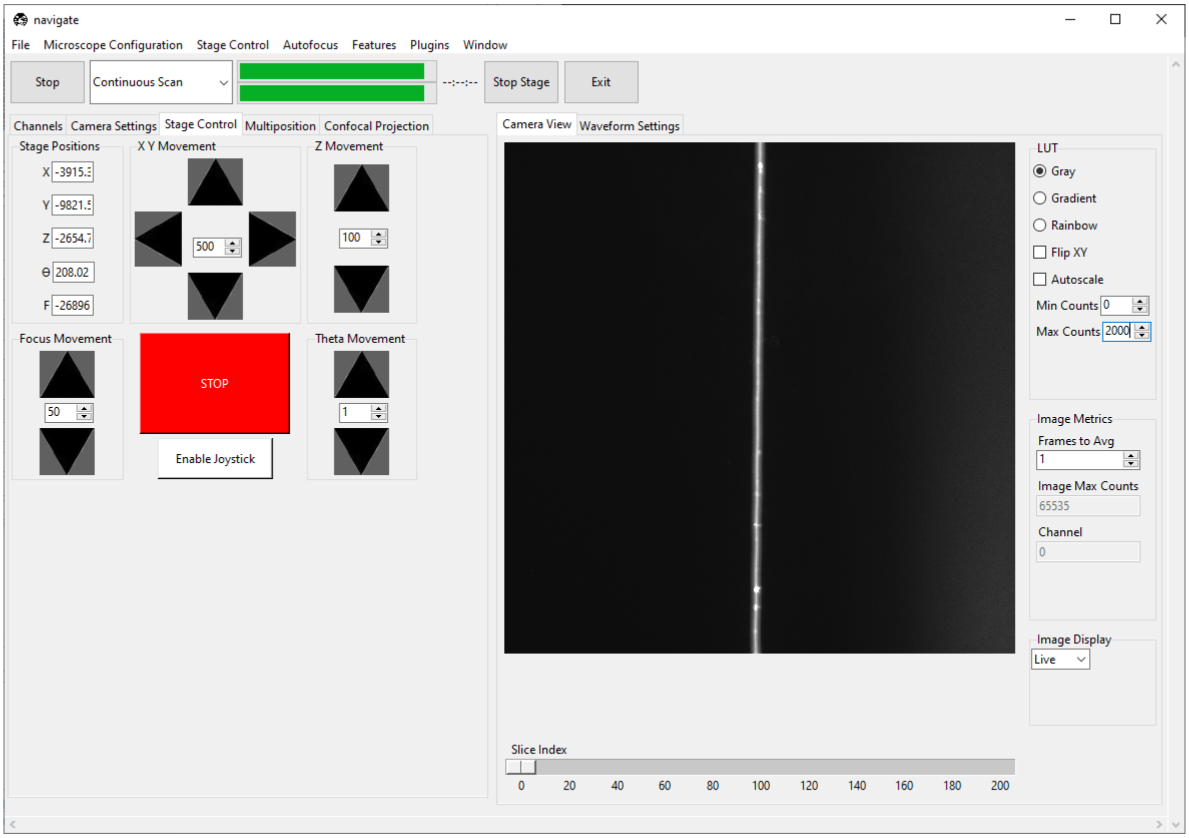

Adjust the *F* (focus) value in the *Stage Control* panel until the beam is as thin/focused as possible.

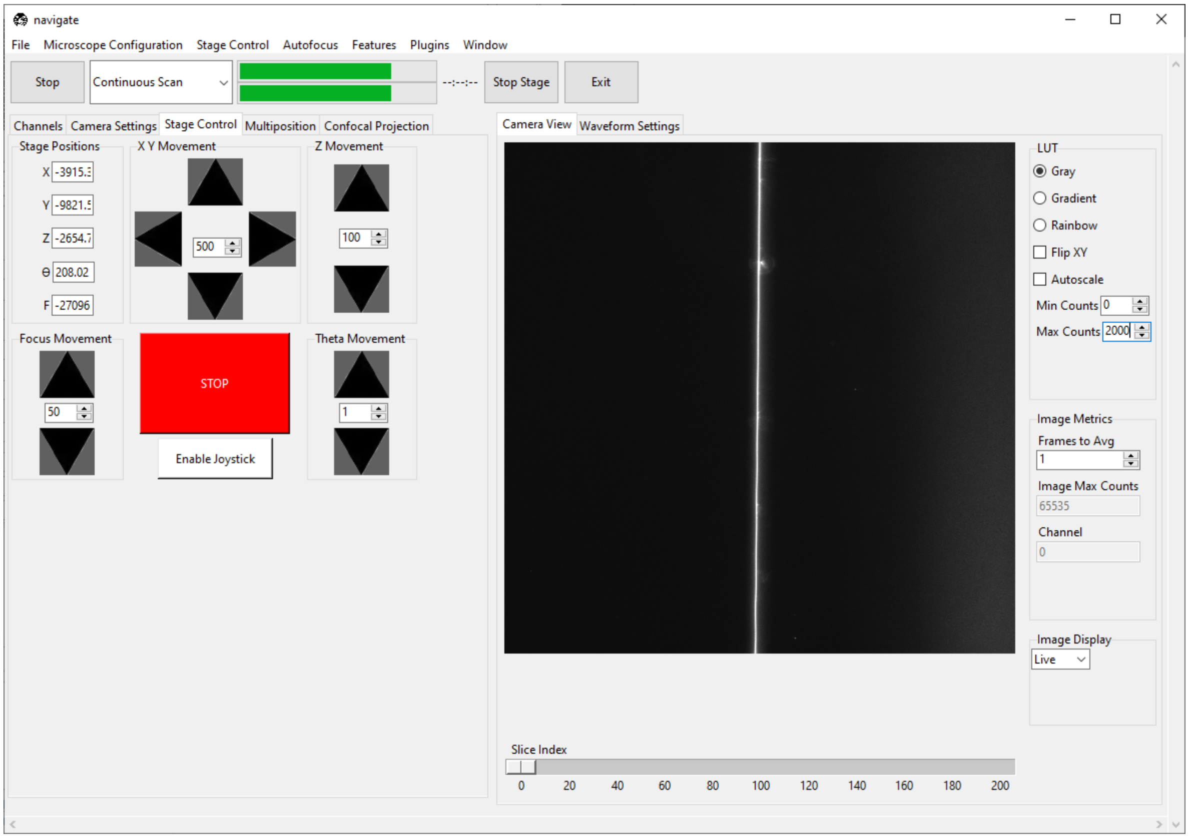

6. Once the beam is straight, slowly change the wavelength's *Offset* until the beam has an even thickness across the field of view. This will also make the beam a bit thinner.

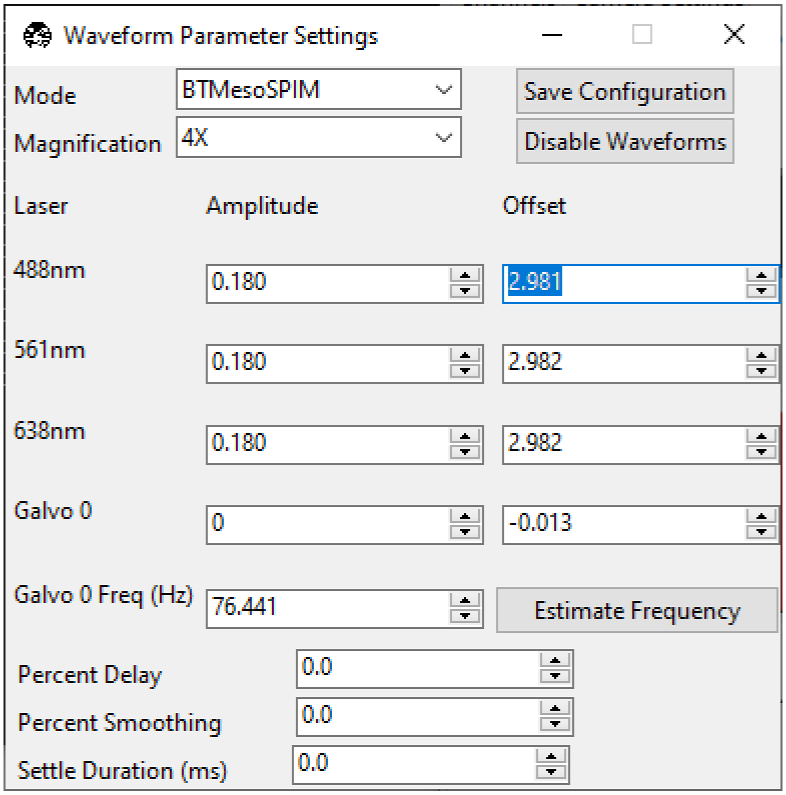
Waveform Parameter Settings
Mode
BTMesoSPIM
Save Configuration
Magnification
4X
Disable Waveforms
Laser
Amplitude
Offset
488nm
0.180
2.981
561nm
0.180
2.982
638nm
0.180
2.982
Galvo 0
0
-0.013
Galvo 0 Freq (Hz)
76.441
Estimate Frequency
Percent Delay
0.0
Percent Smoothing
0.0
Settle Duration (ms)
0.0

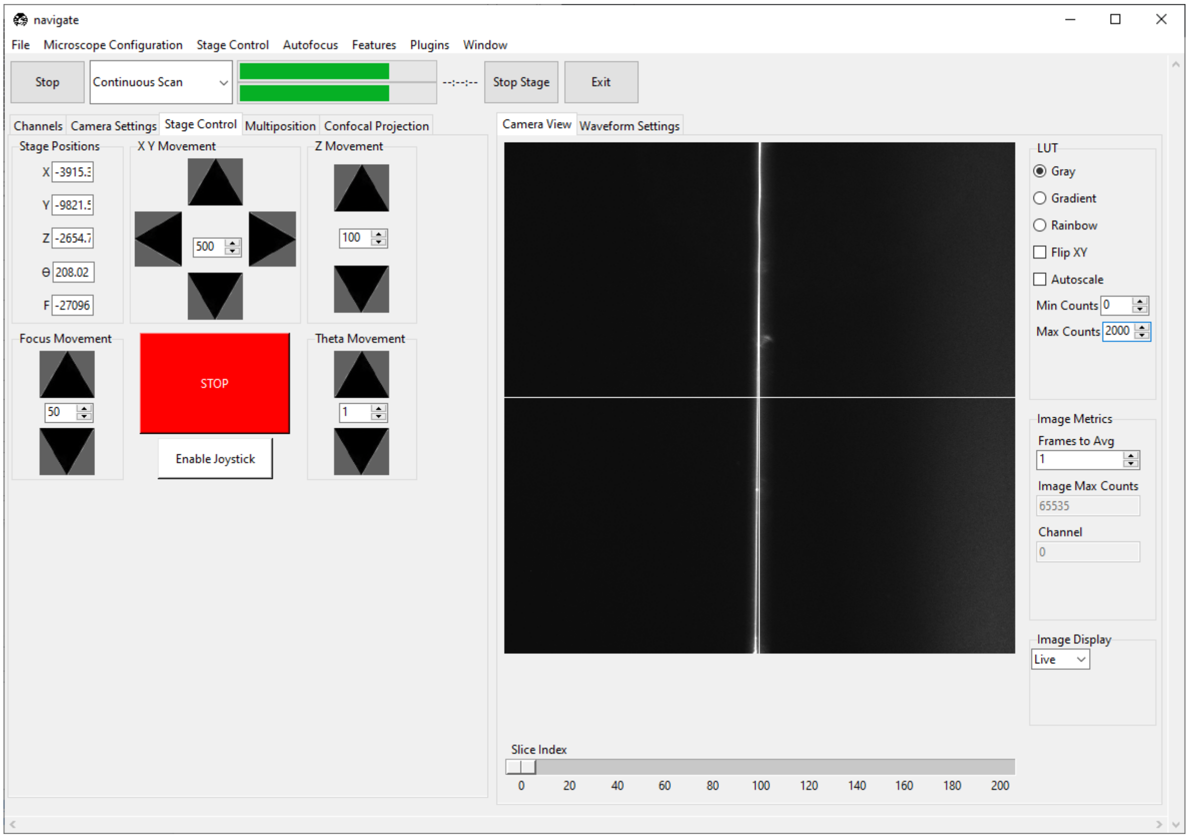
navigate
File
Microscope Configuration
Stage Control
Autofocus
Features
Plugins
Window
Stop
Continuous Scan
Stop Stage
Exit
Channels
Camera Settings
Stage Control
Multiposition
Confocal Projection
Camera View
Waveform Settings
Stage Positions
X Y Movement
Z Movement
Focus Movement
STOP
Enable Joystick
Theta Movement
LUT
Gray
Gradient
Rainbow
Flip XY
Autoscale
Min Counts
Max Counts
Image Metrics
Frames to Avg
Image Max Counts
65535
Channel
Image Display
Live
Slice Index

> **Warning:** Proper alignment of the ASLM scan is critical to the quality of the image. We recommend iterating the *Amplitude*, *Offset*, and *F* until the beam is uniformly as thin as possible throughout the entire field of view.

1. Slowly increase *Galvo 0*'s *Amplitude* until the entire field of view is just covered by the digitally scanned beam. Over-scanning the beam will result in a loss of light, but also provide a more uniform illumination for tiling applications.

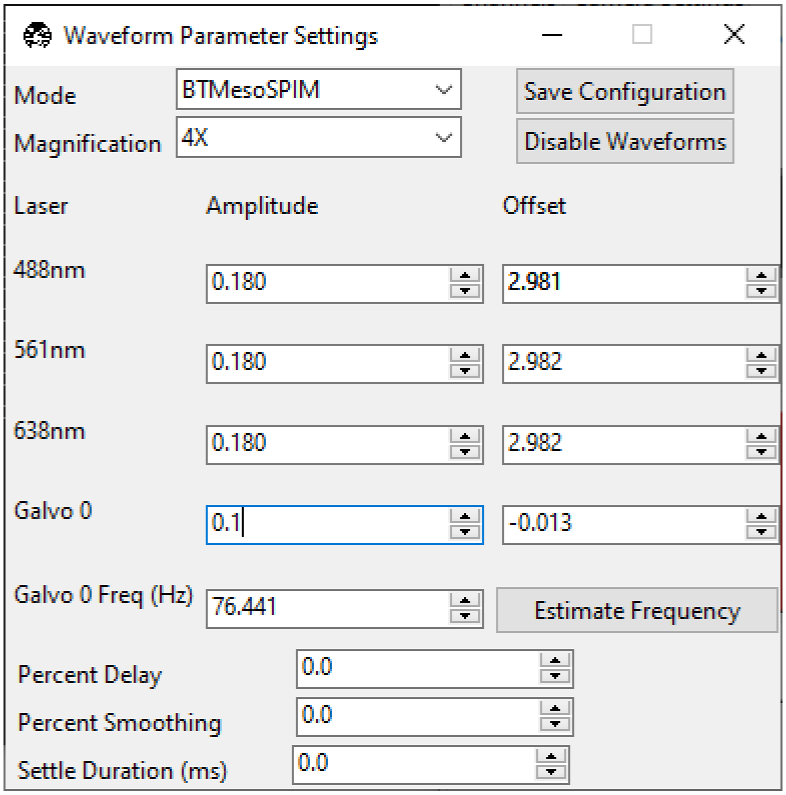

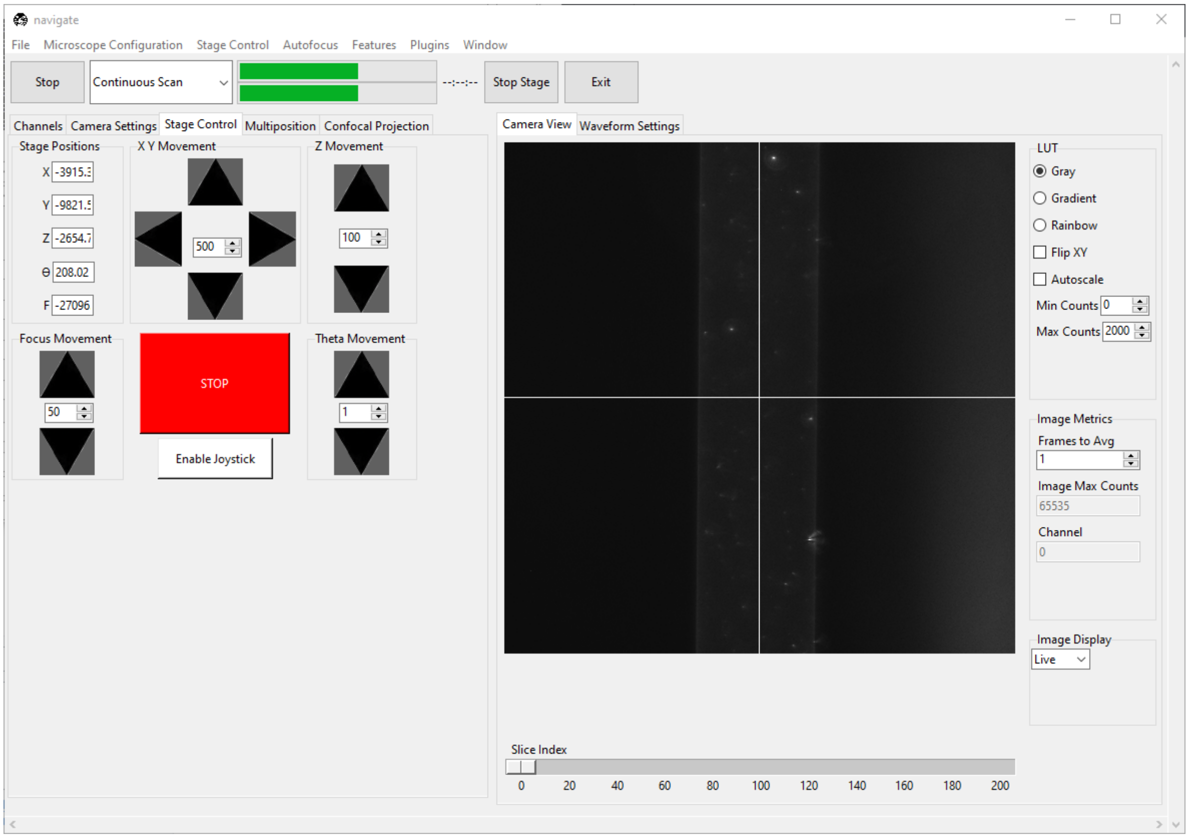
navigate
File
Microscope Configuration
Stage Control
Autofocus
Features
Plugins
Window
Stop
Continuous Scan
Stop Stage
Exit
Channels
Camera Settings
Stage Control
Multiposition
Confocal Projection
Camera View
Waveform Settings
Stage Positions
X Y Movement
Z Movement
Focus Movement
STOP
Enable Joystick
Theta Movement
LUT
Gray
Gradient
Rainbow
Flip XY
Autoscale
Min Counts
Max Counts
Image Metrics
Frames to Avg
Image Max Counts
Channel
Image Display
Live
Slice Index

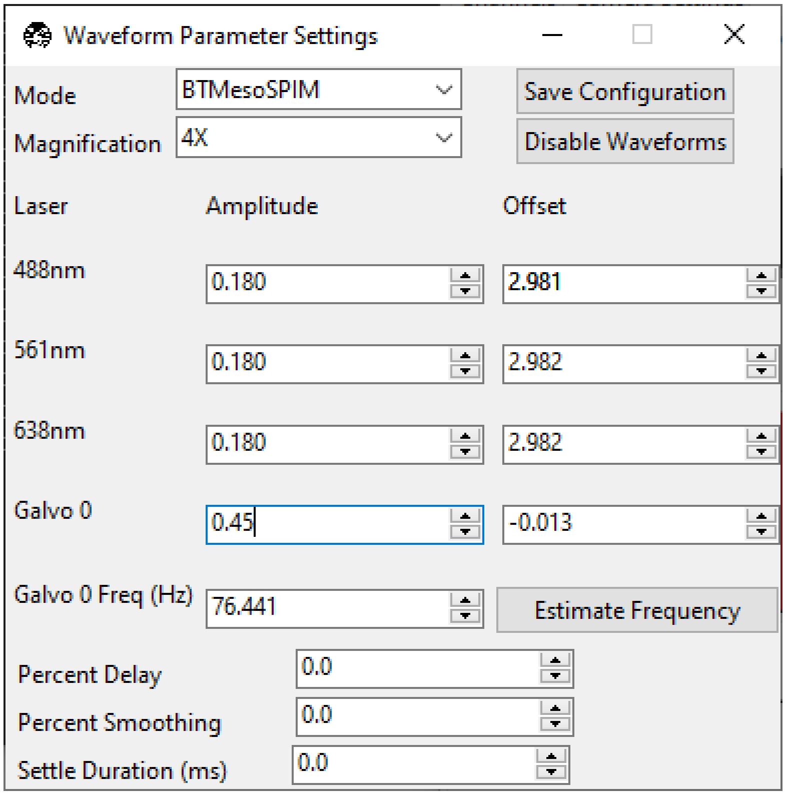
Waveform Parameter Settings
Mode
BTMesoSPIM
Save Configuration
Magnification
4X
Disable Waveforms
Laser
Amplitude
Offset
488nm
0.180
2.981
561nm
0.180
2.982
638nm
0.180
2.982
Galvo 0
0.45
-0.013
Galvo 0 Freq (Hz)
76.441
Estimate Frequency
Percent Delay
0.0
Percent Smoothing
0.0
Settle Duration (ms)
0.0

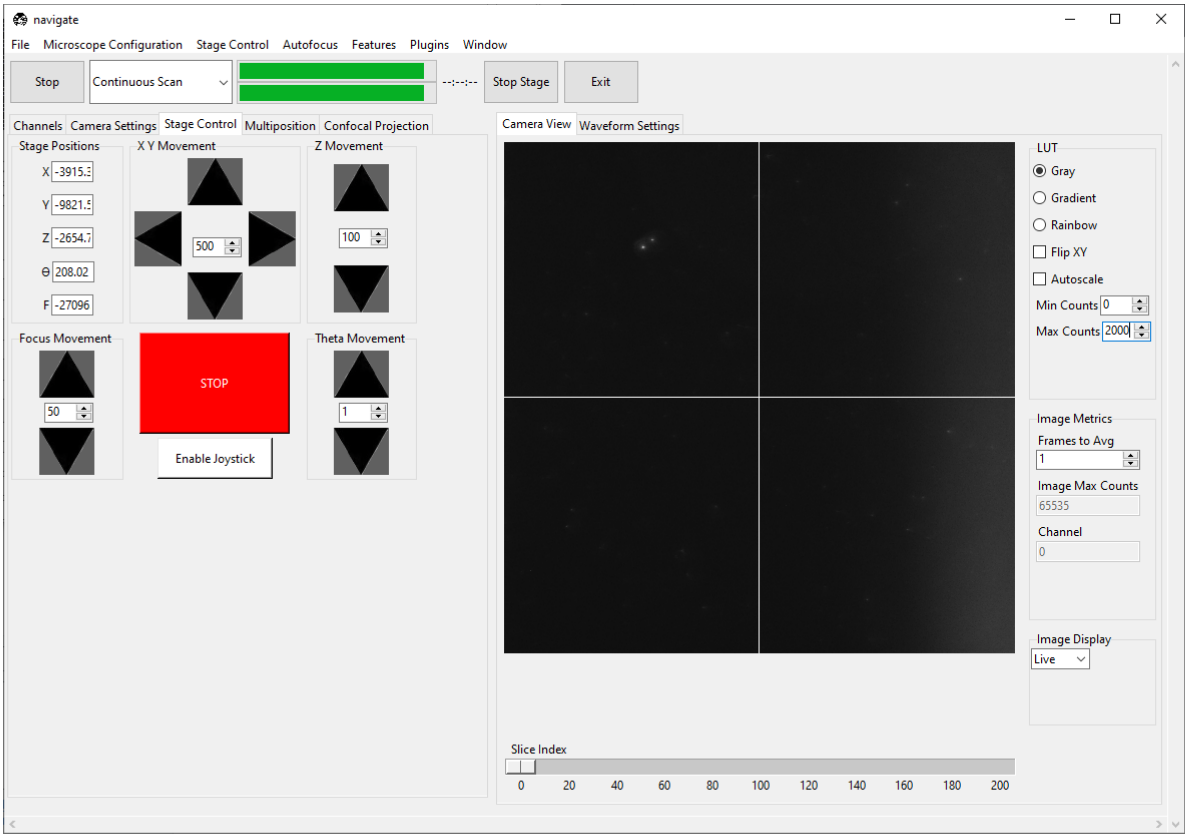

2. Under *Waveform Parameter Settings*, press *Save Configuration*.
3. Under the *Channels* tab, restore the filter to its non-empty position.

##### 7.2.2 Loading and finding the sample

1. Load the sample on the microscope.
2. Select “Continuous Scan” from the dropdown next to the *Acquire* button. Press *Acquire*. This will launch a live acquisition mode.
3. Scroll around with the stage either via joystick or using the controls in the *Stage Control* tab until the sample comes into view.

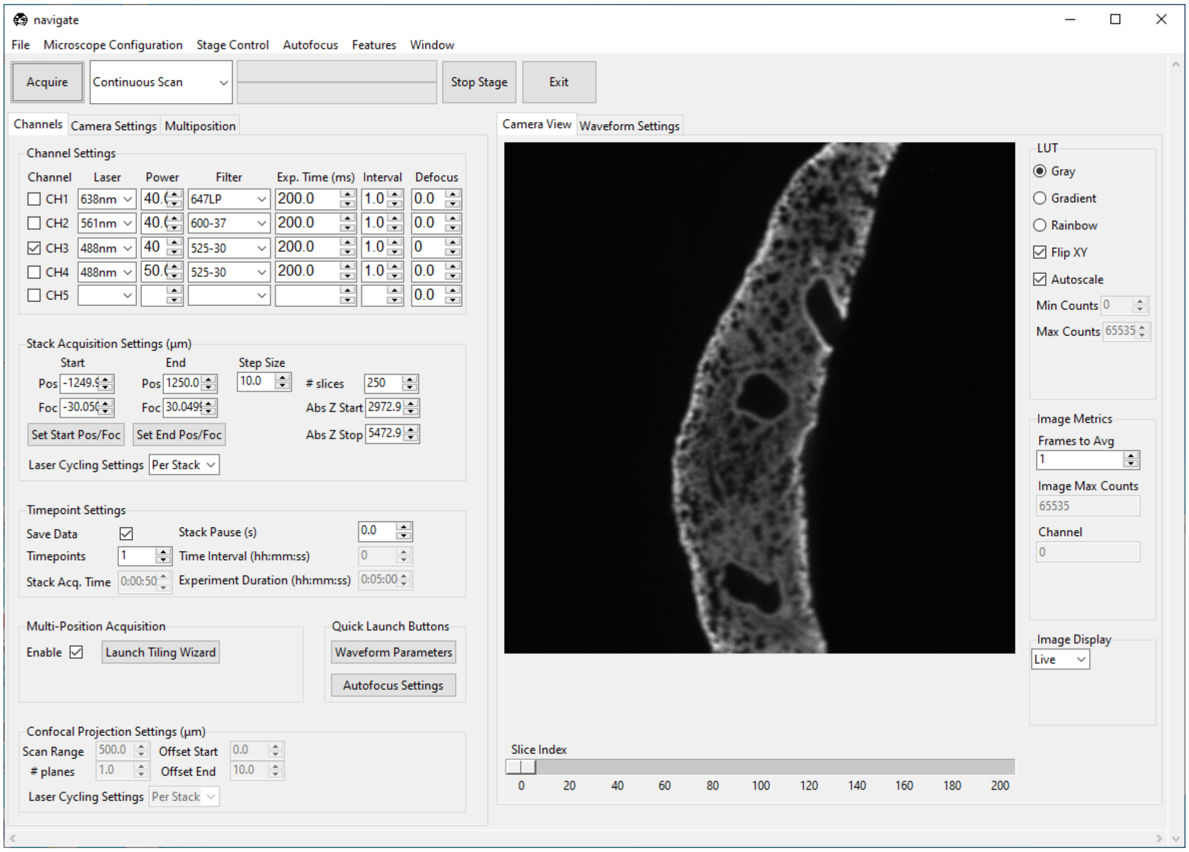

4. Focus on the sample using the F axis. Optionally, use Autofocus by going to *Autofocus* → *Autofocus Settings*. Press *Autofocus*. Ensure there is a clear peak in the resulting plot.

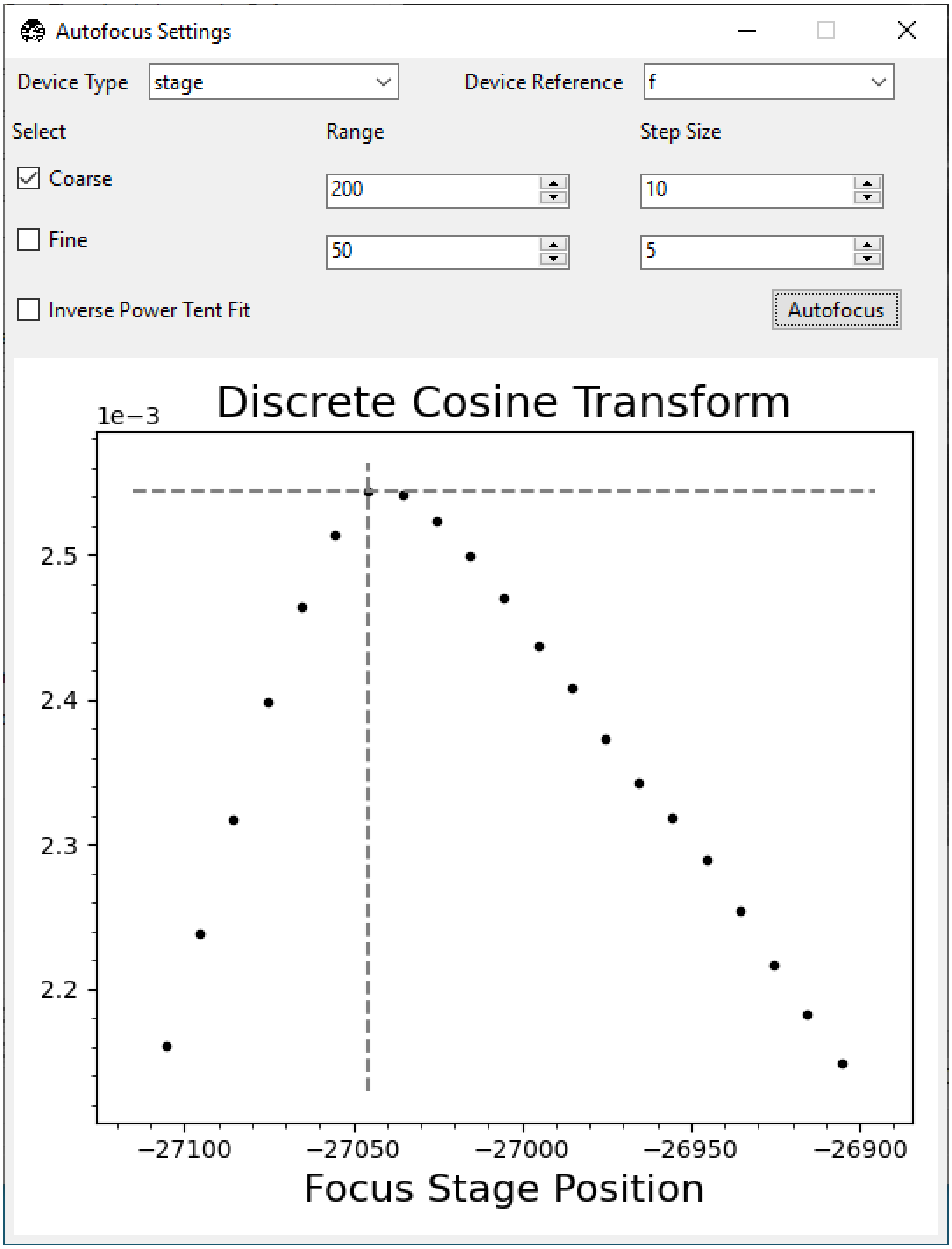

If there is not a clear peak, the autofocusing routine did not work. Try increasing the laser power and/or bringing the sample more into focus manually. If it did work, the sample should now be in focus.

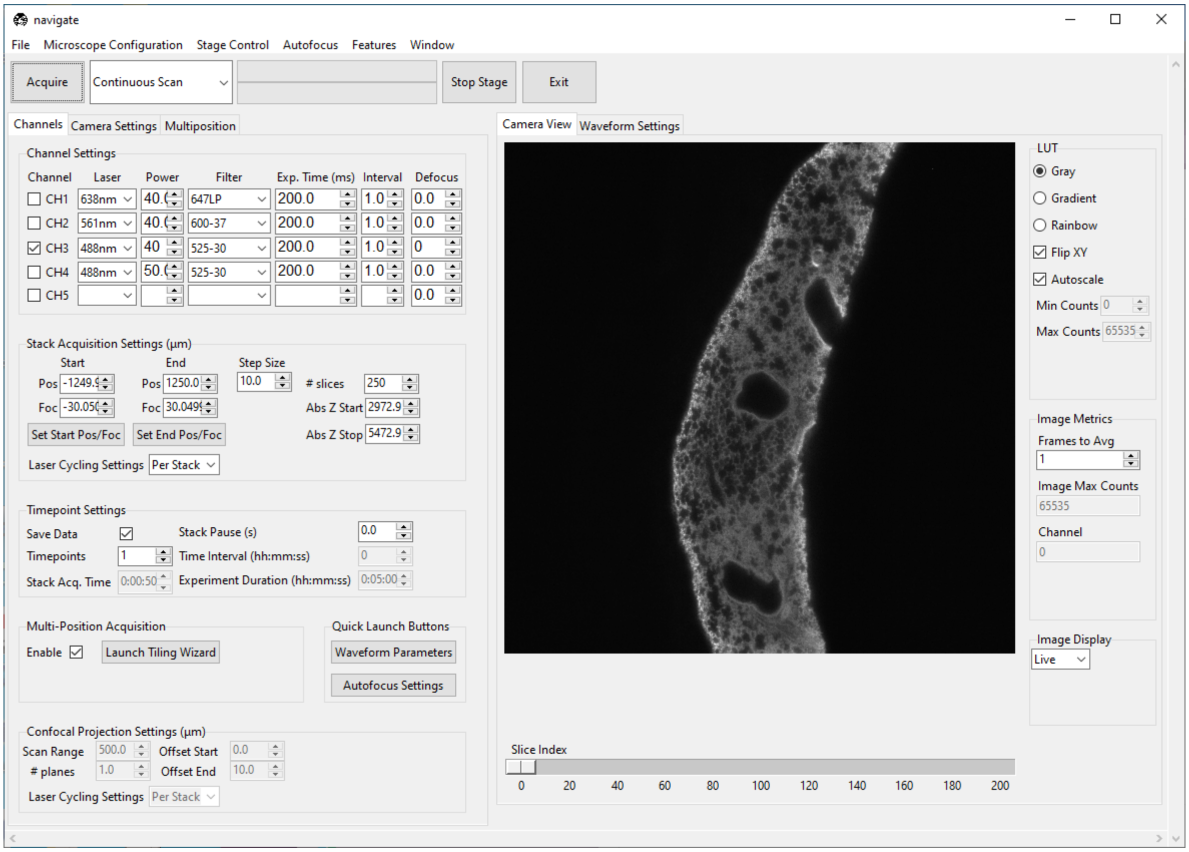

**Note:** Sometimes there isn't a clear peak, but there is a clear trend toward a peak. In this case, the autofocus is converging, but the true focus position is outside the range of your search. Run autofocus again to achieve convergence.

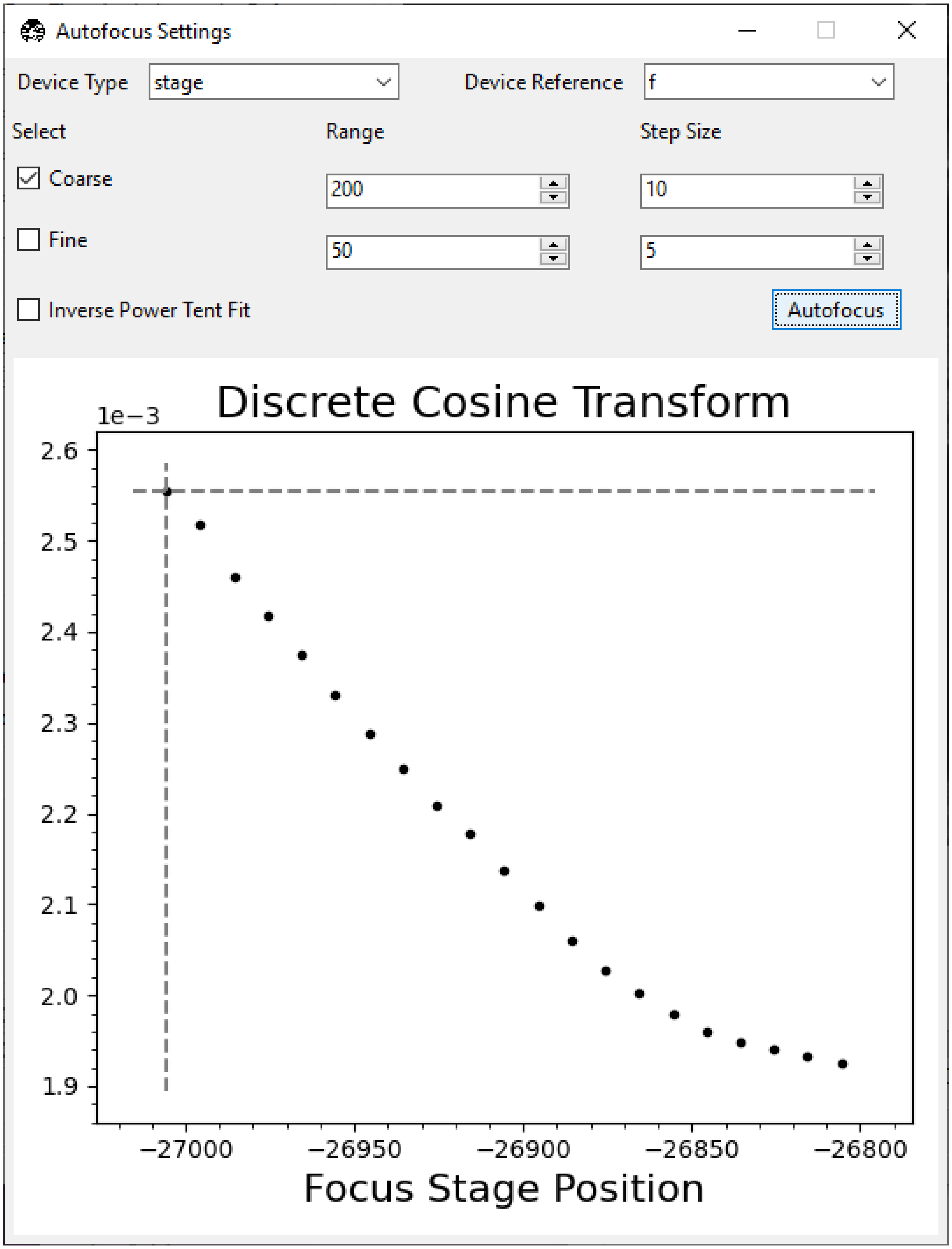

Autofocus Settings
Device Type
stage
Device Reference
f
Select
Range
Step Size
Coarse
200
10
Fine
50
5
Inverse Power Tent Fit
Autofocus
Discrete Cosine Transform
1e−3
2.6
2.5
2.4
2.3
2.2
2.1
2.0
1.9
−27000
−26950
−26900
−26850
−26800
Focus Stage Position

##### 7.2.3 Imaging a z-stack

1. Select "Continuous Scan" from the dropdown next to the *Acquire* button. Press *Acquire*. This will launch a live acquisition mode.
2. Using the *Stage Control*, go to a shallow Z-position in the sample. Under the *Channels* tab, in *Stack Acquisition Settings (um)* press *Set Start Pos/Foc*.

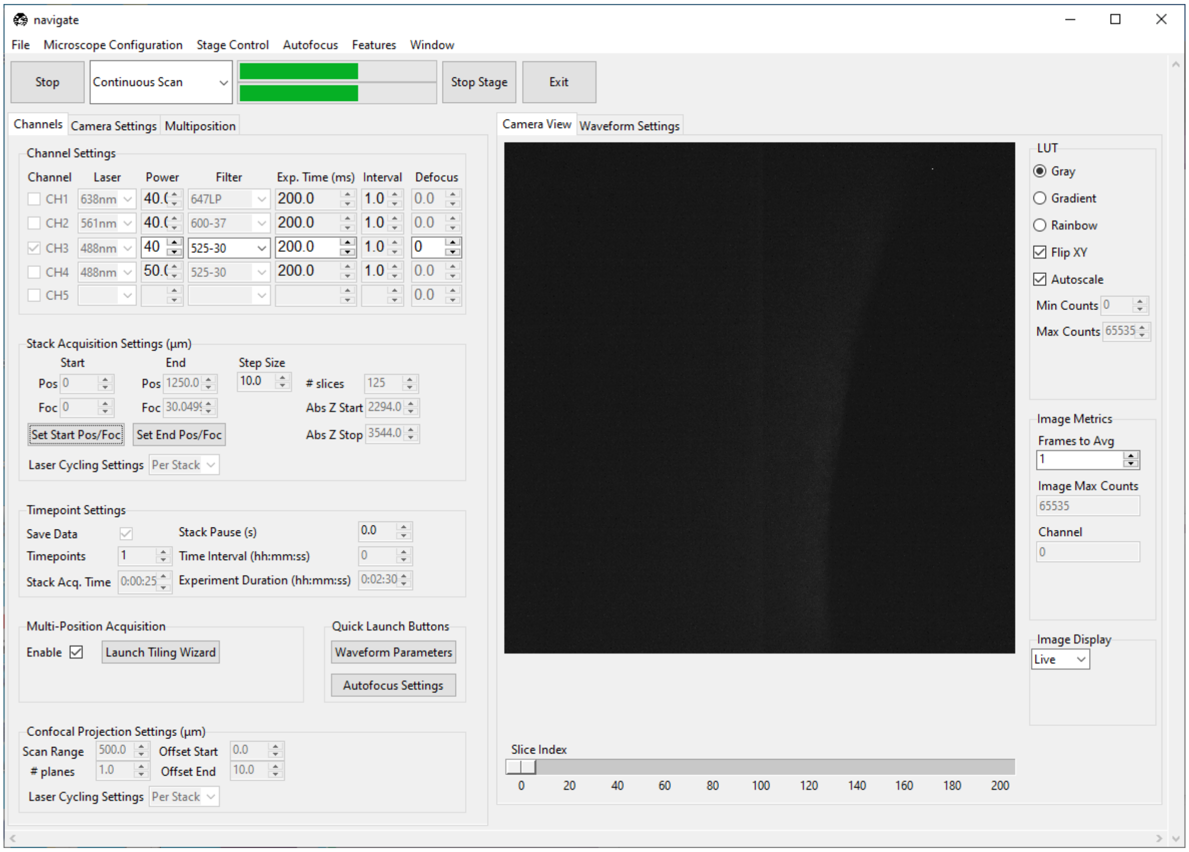

3. Go to a deep Z-position in the sample. Press *Set End Pos/Foc*.

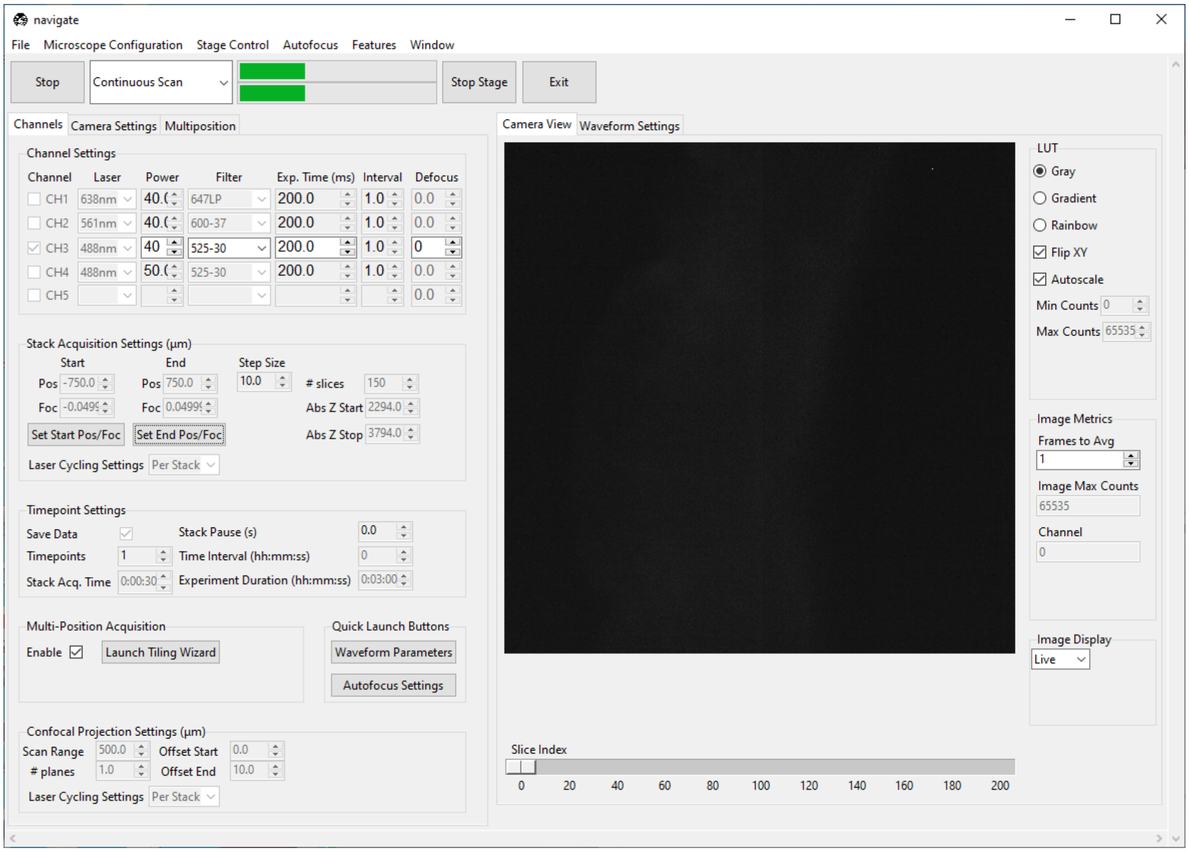

4. Select “Z-Stack” from the dropdown next to the *Acquire* button. Press *Acquire*.
5. Enter the sample parameters in the *File Saving Dialog* that pops up. Press *Acquire Data*.

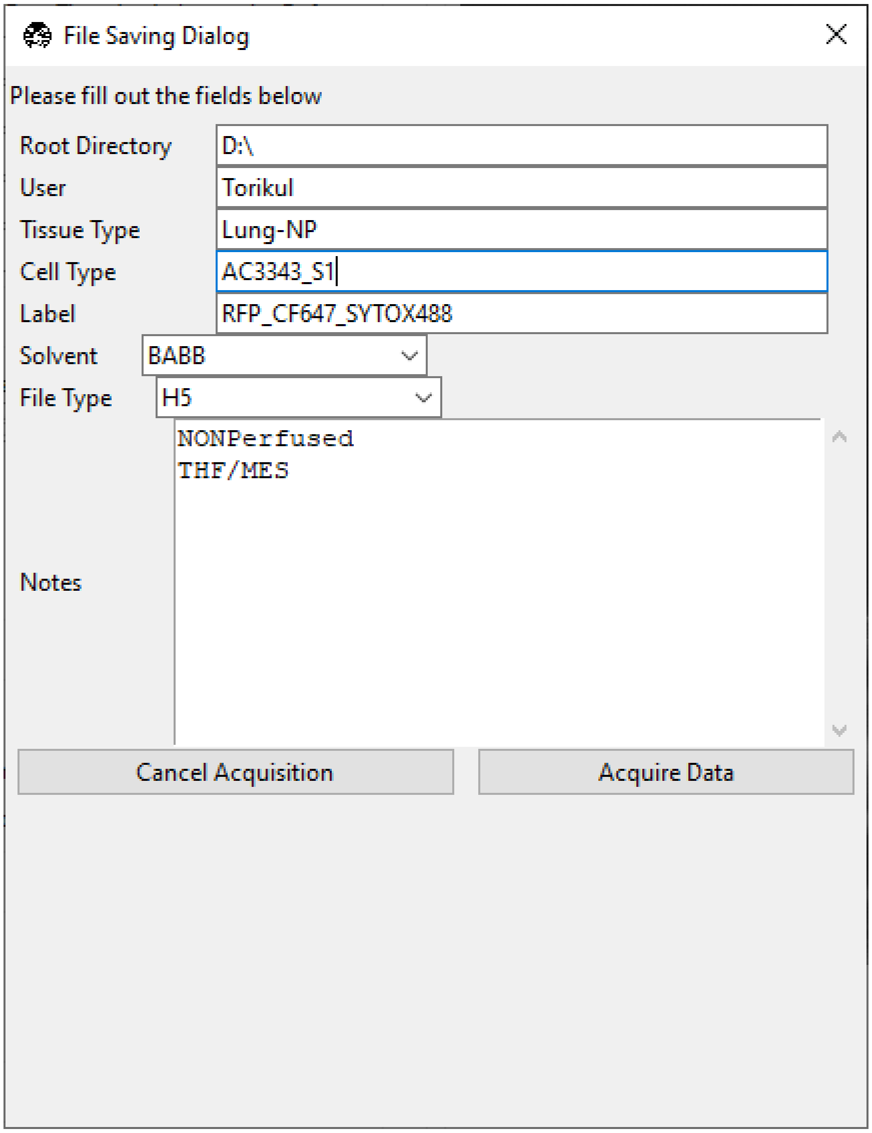

##### 7.2.4 Tiling a sample larger than the field of view

This assumes you have already set the start and end positions in *Stack Acquisition Settings (um)* (see *Imaging a Z-Stack*).

1. Under the *Channels* tab, press *Launch Tiling Wizard*.

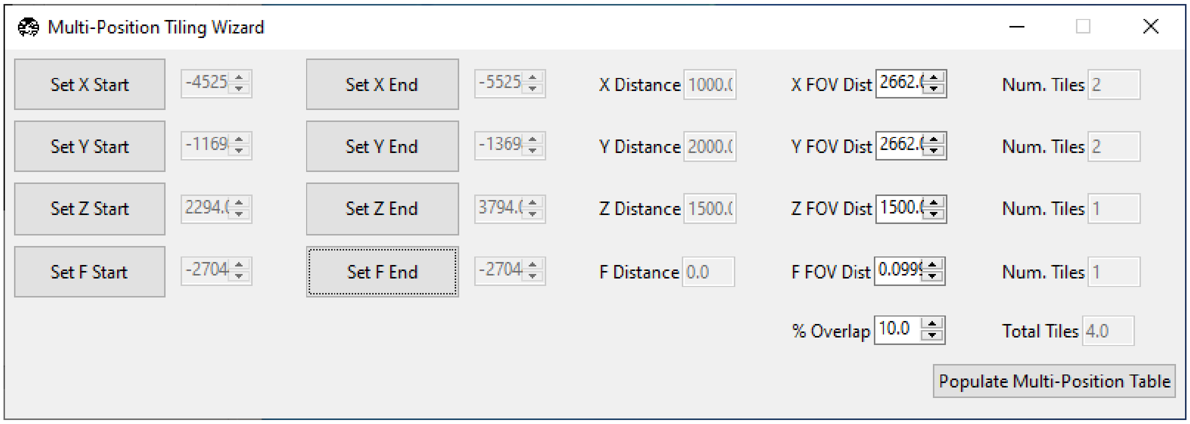

2. Go to thickest part of the sample. Go to the lower bound of the `X` axis and press *Set X Start*. Go to the upper bound of the `X` axis and press *Set X End*. Repeat for all axes except for focus.
3. Ensure the sample is in focus and press *Set F Start* and *Set F End* without changing the focus position.
4. Press *Populate Multi-Position Table*. Navigate to the *Multiposition* tab and ensure the locations populated.

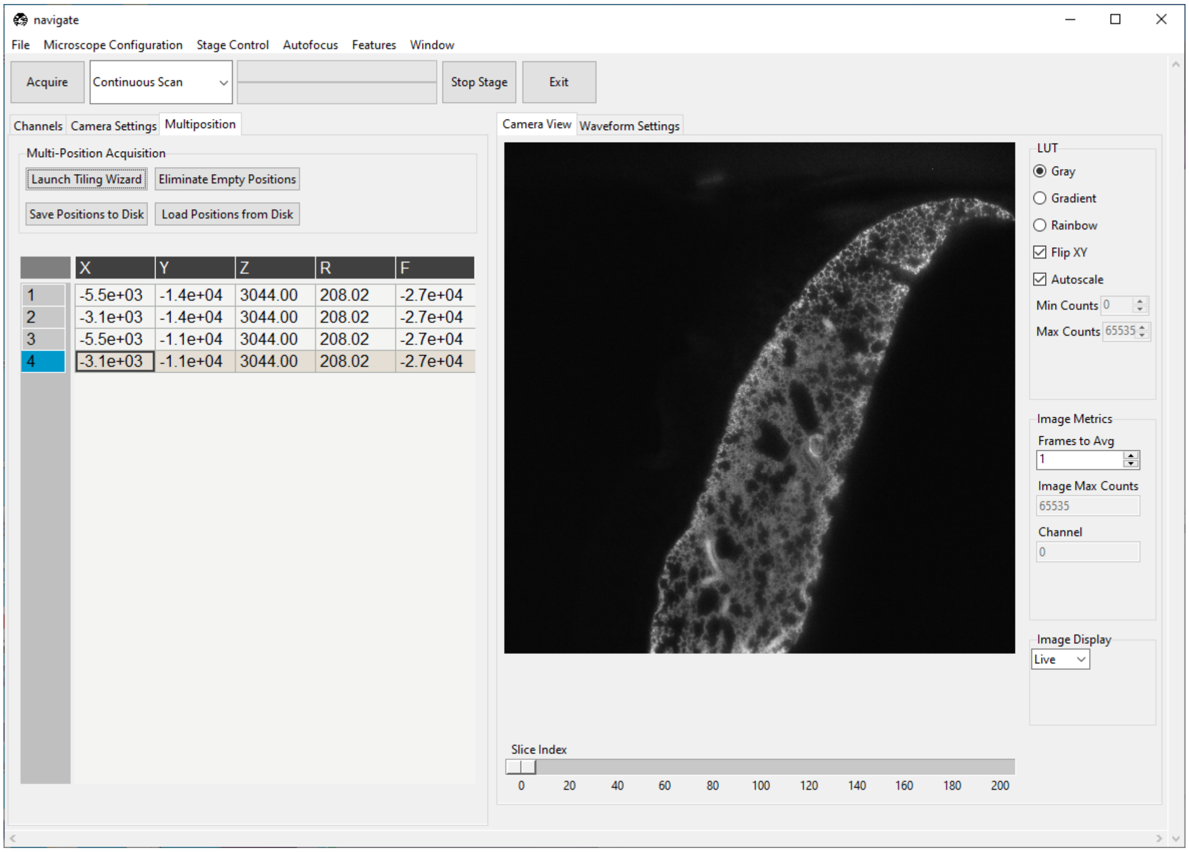

5. Under the *Channels*, make sure *Enable* is checked under *Multi-Position Acquisition*.
6. Under the *Channels*, make sure *Save Data* is checked under *Timepoint Settings*.
7. Select "Z-Stack" from the dropdown next to the *Acquire* button. Press *Acquire*.
8. Enter the sample parameters in the *File Saving Dialog* that pops up. Press *Acquire Data*.

#### 7.3 Imaging on the CT-ASLM-V1 and CT-ASLM-V2

This is a case study in using the software to image with a CT-ASLM-V1 and CT-ASLM-V2 microscopes.

---

##### 7.3.1 Setting up the chamber

Make sure the chamber is clean and dry. If in doubt, fill the chamber with deionized water to see if there is any residue. To clean the chamber, rinse it with deionized water and ethanol, then gently clean the chamber with Q-tips. Repeat the process a few times and end the process with a rinse of 100% ethanol. Gently clean the objectives with lens paper and 100% ethanol. Finally, let the chamber air dry. To speed up the drying process, one can gently blow some air into the chamber.

Once the chamber is completely dry, fill the chamber with imaging media.

---

##### 7.3.2 Sample loading and finding the samples

It's recommended to start the software before loading the sample on the stage.

1. Mount the sample on custom cut glass slide with silicon or super glue.
2. Mount the glass slide onto the sample holder.
3. Mount the sample holder onto the stage with the sample facing the illumination and detection objective so the glass slide is 45 degrees with both objectives.
4. Decrease the numerical aperture of the illumination beam such that it covers the entire field of view. Typically this is achieved with a magnetic mounted slit aperture that can be readily adjusted.
5. Go to *Camera Settings*. Select *Normal* under *Sensor Mode*.
6. Go to the *Microscope Configuration* → *Waveform Parameters*. A popup named *Waveform Parameter Settings* will appear.
7. Set the wavelength's *Amplitude* and *Offset* to `0.0`.
8. Select the channel with a proper laser under the *Channels* tab and set the laser power to an appropriate value.
9. Select "Continuous Scan" from the dropdown next to the *Acquire* button. Press *Acquire*. This will launch a live acquisition mode.
10. Scroll around with the stage either via joystick or using the controls in the *Stage Control* tab until the sample comes into view.
11. Focus on the sample in the center of the beam. Zoom in by placing the mouse over the image and scrolling the mouse wheel. Slowly adjusting the focus by scrolling the piezo controller to move the detection objective along the z axis. Lower the laser power if the image is saturated.

---

##### 7.3.3 Imaging a Z-Stack with Stelzer mode

Stelzer mode is the normal non-ASLM light sheet mode, it gives more signal while offering around 1040 nm (CT-ASLM-V1) and 500 nm (CT-ASLM-V2) axial resolution.

1. Go to the *Microscope Configuration* → *Waveform Parameters*. A popup named *Waveform Parameter Settings* will appear.
2. Set the wavelength's *Amplitude* and *Offset* to `0.0`.
3. Go to *Camera Settings*, select *Normal* under *Sensor Mode*.
4. Put a slit into the setup.
5. Select the channel with a proper laser under the *Channels* tab and set the laser power to an appropriate value.
6. Select "Continuous Scan" from the dropdown next to the *Acquire* button. Press *Acquire*. This will launch a live acquisition mode.
7. If needed, slowly adjust the slit opening until the image sharpness looks uniform across the whole field of view. Uncheck *Autoscale* in *Camera View* under LUT and adjust the *Min Counts* and *Max Counts* if needed.
8. Go to *Stage Control*, set the Z position in *Stage Positions* to be `0`.
9. Find the region of interest by using the joystick or using the controls in the *Stage Control* tab.
10. Move along the Z axis with the joystick or the "Focus" in the *Stage Control* tab to one end of the region of interest. Under the *Channels* tab, in *Stack Acquisition Settings (um)*, press *Set Start Pos/Foc*.
11. Go to *Stage Control*, change the Z position in *Stage Positions* to set the scan range. Be aware the range for z-piezo is 0 - 200. Going outside of the range will cause the stage to have issues.
12. Go back to *Channels* tab, in *Stack Acquisition Settings (um)*, press *Set End Pos/Foc*.
13. Setup *Step Size* under the *Channels*, recommend 3.0 (CT-ASLM-V1) and 1.0 (CT-ASLM-V2).
14. Under the *Channels*, make sure *Enable* is unchecked under *Multi-Position Acquisition*.
15. Under the *Channels*, make sure *Save Data* is checked under *Timepoint Settings*.
16. Select "Z-Stack" from the dropdown next to the *Acquire* button. Press *Acquire*. A popup named *File Saving Dialog* will appear.
17. Fill out the fields and press *Acquire Data*.

---

##### 7.3.4 Imaging a Z-Stack with ASLM mode

ASLM mode is the high-resolution light sheet mode, it gives leas signal but offering around 950 nm (CT-ASLM-V1) and 480 nm (CT-ASLM-V2) isotropic resolution.

1. Switch the slit out of the setup.
2. Go to *Camera Settings*, select "Light-Sheet" under *Sensor Mode*.
3. Select the channel with a proper laser under the *Channels* tab and set the laser power to an appropriate value.
4. Select "Continuous Scan" from the dropdown next to the *Acquire* button. Press *Acquire*. This will launch a live acquisition mode.
5. Go to the *Microscope Configuration* → *Waveform Parameters*. A popup named *Waveform Parameter Settings* will appear.
6. Uncheck *Autoscale* in *Camera View* under LUT and adjust the *Min Counts* and *Max Counts* if needed.

7. Set the wavelength's *Amplitude* to `0.0`.
8. Adjust the wavelength's *Offset* so the focus part of the image can be located perfectly in the center of the field of view.
9. Slowly adjust the wavelength's *Amplitude* so it will be uniform across the whole field of view.
10. Adjust the wavelength's *Offset* again slightly and make sure it is uniformly in focus across the whole field of view.
11. Go to *Stage Control*, set the Z position in *Stage Positions* to be `0`.
12. Find the region of interest by using the joystick or using the controls in the *Stage Control* tab.
13. Move along the Z axis with the joystick or the "Focus" in the *Stage Control* tab to one end of the region of interest. Under the *Channels* tab, in *Stack Acquistion Settings (um)*, press *Set Start Pos/Foc*.
14. Go to *Stage Control*, change the Z position in *Stage Positions* to set the scan range. Be aware the range for z-piezo is 0 - 200. Going outside of the range will cause the stage to have issues.
15. Go back to *Channels* tab, in *Stack Acquisition Settings (um)*, press *Set End Pos/Foc*.
16. Setup *Step Size* under the *Channels*, recommend 0.46 (CT-ASLM-V1) and 0.2 (CT-ASLM-V2) for isotropic imaging.
17. Under the *Channels*, make sure *Enable* is unchecked under *Multi-Position Acquisition*.
18. Under the *Channels*, make sure *Save Data* is checked under *Timepoint Settings*.
19. Select "Z-Stack" from the dropdown next to the *Acquire* button. Press *Acquire*. A popup named *File Saving Dialog* will appear.
20. Fill out the fields and press *Acquire Data*.

---

##### 7.3.5 Tiling a sample larger than the field of view

This assumes you have already found the samples and ready to acquire data in either Stelzer mode or ASLM mode. (see *Imaging a Z-Stack with Stelzer mode* and *Imaging a Z-Stack with ASLM mode*).

1. Under *Channels* tab, press *Launch Tiling Wizard*. A popup named *Multi-Position Tiling Wizard* will appear.
2. Follow *Imaging a Z-Stack with Stelzer mode* to set up the start and end positions in *Stack Acquisition Settings (um)*. At the same time, when pressing *Set Start Pos/Foc* to set up the start position, go to *Multi-Position Tiling Wizard* and press *Set Z Start*. When pressing *Set End Pos/Foc* to set up the end position, go to *Multi-Position Tiling Wizard* and press *Set Z End*.
3. Move the joystick or the "X Movement" in the *Stage Control* tab to the lower bound of the x-axis and press *Set X Start* in the *Multi-PositionTiling Wizard* popup. Navigate to the upper bound of the x-axis and press *Set X End* in the *Multi-Position Tiling Wizard* popup. Repeat for all axes except for z.
4. Press *Populate Multi-Position Table*. Navigate to the *Multiposition* tab and ensure the locations populated.
5. Under the *Channels*, make sure Enable is checked under *Multi-Position Acquisition*.
6. Under the *Channels*, make sure *Save Data* is checked under *Timepoint Settings*.
7. Select "Z-Stack" from the dropdown next to the *Acquire* button. Press *Acquire*.
8. Enter the sample parameters in the *File Saving Dialog* that pops up. Press *Acquire Data*.

#### 7.4 Imaging on an Upright ASLM

This is a case study in using the software to image with an upright ASLM microscope. The upright ASLM equipped with an ASI FTP2000 and an ASLM microscope in an upright configuration. This microscope configuration allows for imaging across large scan ranges and imaging during scanning which we term constant velocity acquisition. This tutorial aims to show how it is possible to image the sample while imaging in both ASLM mode and normal lightsheet mode.

##### 7.4.1 Loading and finding the sample

1. Load the sample on the microscope.
2. Select "Continuous Scan" from the dropdown next to the *Acquire* button. Press *Acquire*. This will launch a live acquisition mode.
3. Scroll around with the stage either via joystick or using the controls in the *Stage Control* tab until the sample comes into view.
4. Set the resonant galvo *Galvo 0* to 0.3 to mitigate any striping artifacts during imaging.

##### 7.4.2 Imaging a Z-Stack using Stop and Settle Mode

1. Select *Continuous Scan* from the dropdown next to the *Acquire* button. Press *Acquire*. This will launch a live acquisition mode.
2. Using the *Stage Control*, go to a shallow z-position in the sample. Under the *Channels* tab, in *Stack Acquisition Settings (um)* press *Set Start Pos*.
3. Go to a deep z-position in the sample. Press *Set End Pos*.
4. Make sure *Set Foc* is `0` for both the *Set Start Pos* and *End Pos*.
5. Type the desired step size (units um) in the *Step Size* dialog box in *Stack Acquisition Settings (um)*. Step size can only be in increments of 0.1 and the minimum is 0.2.
6. Select the number of color channels needed imaging in the *Channel tab* under *Channel Settings*. Select the correct filter for each channel by using the dropdown menu after each channel under the *Filter*.
7. Change the exposure time by changing number in the *Exp. Time (ms)* for each channel. For the ORCA Lightning camera using ASLM mode, the minimum frame rate is 75 ms and the maximum is 100 ms.
8. Set *Interval* to be `1.0` for each channel.
9. Set *Defocus* to be `0` for each channel.
10. Select "Z-Stack" from the dropdown next to the *Acquire* button. Press *Acquire*.
11. Enter the sample parameters in the *File Saving Dialog* that pops up. Press *Acquire Data*.

##### 7.4.3 Imaging a Z-Stack using Constant Velocity Acquisition Mode

1. Select "Continuous Scan" from the dropdown next to the *Acquire* button. Press *Acquire*. This will launch a live acquisition mode.
2. Using the *Stage Control*, go to a shallow Z-position in the sample. Under the *Channels* tab, in *Stack Acquisition Settings (um)* press *Set Start Pos*.
3. Go to a deep Z-position in the sample. Press *Set End Pos*.
4. Make sure *Set Foc* is `0` for both the *Set Start Pos* and *End Pos*.
5. Type the desired step size (units um) in the *Step Size* dialog box in *Stack Acquisition Settings (um)*. Step size can only be increments of 0.1 and the minimum is 0.2.
6. Select the number of color channels needed imaging in the *Channel tab* under *Channel Settings*. Select the correct filter for each channel by using the dropdown menu after each channel under the *Filter*.
7. Change the exposure time by changing number in the *Exp. Time (ms)* for each channel. For the ORCA Lightning camera using ASLM mode, the minimum frame rate is 75 ms and the maximum is 100 ms.
8. Set *Interval* to be `1.0` for each channel.
9. Set *Defocus* to be *0* for each channel.
10. Select "Constant Velocity Acquisition" from the dropdown next to the *Acquire* button. Press *Acquire*.
11. Enter the sample parameters in the *File Saving Dialog* that pops up. Make sure to save to SSD drive or change buffer size in configuration file to prevent any overwriting of images. Then Press *Acquire Data*.
12. To change frame buffer size, in the *CameraParameters* section in the *experiment.yaml* file in your local navigate directory in the *config* folder, change *databuffer_size* to desired number of frames. Make sure the size of the desired number of frames isn't above the available RAM in the computer.

CHAPTER

EIGHT

### CONTRIBUTING GUIDELINES

We welcome contributions in the form of bug reports, bug fixes, new features and documentation. If you are contributing code, please create it in a fork and branch separate from the main `develop` branch and then make a pull request to the `develop` branch for code review. Some best practices for new code are outlined below.

If you are considering refactoring part of the code, please reach out to us prior to starting this process. We are happy to invite you to our regular software development meeting.

---

#### 8.1 General Principles

- We use a model-view-controller architecture. New functionality should keep this strong separation.
  - The model operates in its own subprocess and is responsible for communicating with hardware and performing image handling and processing tasks.
  - The view is responsible for displaying the user interface and communicating with the controller.
  - The controller is responsible for managing the user interface and communicating with the model. It relays user input in the form of traces and commands to the model and relays model output in the form of images and data to the view.
- Please do not create new configuration variables unless absolutely necessary, especially in the `configuration.yaml` and `experiment.yaml` files. A new variable is necessary only if no variable stores similar information or there is no way to use the most similar variable without disrupting another part of the code base.
- We are happy to discuss code refactors for improved clarity and speed. However, please do not modify something that is already working without discussing this with the software team in advance.
- All code that modifies microscope control behavior must be reviewed and tested on a live system prior to merging into the `develop` branch.

---

#### 8.2 Coding Style

We follow the PEP8 code style guide. All class names are written in `CamelCase` and all variable names are `lowercase_and_separated_by_underscores`.

---

#### 8.3 Documentation

We use Sphinx to generate documentation from documented methods, attributes, and classes. Please document all new methods, attributes, and classes using a Sphinx compatible version of Numpydoc.

---

#### 8.4 Scientific Units

Please express quantities in the following units when they are in the standard model/ view/controller code. Deviations from this can occur where it is necessary to pass a different unit to a piece of hardware.

- Time - Milliseconds
- Distance - Micrometers
- Voltage - Volts
- Rotation - Degrees

---

#### 8.5 Pre-Commit Hooks

We use pre-commit hooks to enforce consistent code formatting and automate some of the code review process. In some rare cases, the linter may complain about a line of code that is actually fine. For example, in the example code below, Ruff linter complains that the start_stage class is imported but not used. However, it is actually used in as part of an `exec` statement.

```
from navigate.model.device_startup_functions import start_stage
device_name = stage
exec(f"self.{device_name} = start_{device_name}(name, device_connection, configuration,␣
↪i, is_synthetic)")
```

To avoid this error, add a `# noqa` comment to the end of the line to tell Ruff to ignore the error.

```
from navigate.model.device_startup_functions import start_stage  # noqa
```

---

#### 8.6 Dictionary Parsing

The *configuration file* is loaded as a large dictionary object, and it is easy to create small errors in the dictionary that can crash the program. To avoid this, when getting properties from the configuration dictionary, it is best to use the `.get()` command, which provides you with the opportunity to also have a default value should the key provided not be found. For example,

```
# Galvo Waveform Information
self.galvo_waveform = self.device_config.get("waveform", "sawtooth")
```

Here, we try to retrieve the `waveform` key from a the `self.device_config` dictionary. In the case that this key is not available, it then by default returns `sawtooth`. If however the `waveform` key is found, it will provide the value associated with it.

---

#### 8.7 Unit Tests

Each line of code is unit tested to ensure it behaves appropriately and alert future coders to modifications that break expected functionality. Guidelines for writing good unit tests can be found here and over here, or in examples of unit tests in this repository's `test` folder. We use the pytest library to evaluate unit tests. Please check that unit tests pass on your machine before making a pull request.

---

#### 8.8 Developing with a Mac

Many of us have Apple products and use them for development. However, there are some issues that you may encounter when developing on a Mac. Below are some of the issues we have encountered and how to resolve them.

---

##### 8.8.1 Shared memory limits

```
OSError: You tried to simultaneously open more SharedNDArrays than are
allowed by your system!
```

This results from a limitation in the number of shared memory objects that can be created on a Mac. To figure out how many objects can open, open a terminal and run the following command

```
ulimit -n
```

To increase this number, simply add an integer value after it. In our hands, 1000 typically works:

```
ulimit -n 1000
```

### CHAPTER NINE

### FEATURE CONTAINER

**navigate** includes a **feature container** that enables reconfigurable acquisition and analysis workflows. The feature container runs a tree of **features**, where each feature may perform a *signal* operation, which modifies the state of the microscope's hardware, or a *data* operation, where it performs an analysis on acquired image data, or both.

Once a feature is executed, any features dependent on this feature's execution will execute (for example, move the stage, then snap a picture). Following this, the next set of features in sequence will be executed.

Examples of some existing features include `navigate.model.features.common_features.ZStackAcquisition`, which acquires a z-stack, and `navigate.model.features.autofocus.Autofocus`, which finds the ideal plane of focus of a sample using a discrete cosine transform.

#### 9.1 Feature Objects

Each feature is an object that accepts a pointer to `navigate.model.model` in its `__init__()` arguments and contains a configuration dictionary that dictates feature behavior in its `__init__()` function. A complete configuration dictionary is shown below. As few or as many of these options can be specified as needed. Each function is considered a leaf or node of the feature tree.

```
self.config_table = {'signal': {'init': self.pre_func_signal,
                                'main': self.in_func_signal,
                                'end': self.end_func_signal,
                                'cleanup': self.cleanup_func_signal},
                        'data': {'init': self.pre_func_data,
                                 'main': self.in_func_data,
                                 'end': self.end_func_data,
                                 'cleanup': self.cleanup_func_data},
                        'node': {'node_type': 'multi-step',
                                 'device_related': True,
                                 'need_response': True },
                      }
```

Both `signal` and `data` configuration entries are themselves dictionaries that can contain `init`, `main`, `end` and/or `cleanup` entries.

- `init` entries dictate pre-processing steps that must be run before the main function of the feature starts.
- `main` entries dictate the primary operation of the feature, and are run once per acquisition step. They return `True` if the acquisition should proceed and `False` if the acquisition should be ended.

- end entries are run once per main function returning True. They check if the acquisition should end, if we are at any boundary points of the main function (e.g. if we need to change positions in a multi-position z-stack acquisition), and describe any closing operations that must be performed when exiting the feature
- cleanup entries dictate what happens if the node fails. This is for failsafe controls such as "turn off all lasers".

The node configuration dictionary contains general properties of feature nodes. node_type can be one-step or multi-step, the latter indicating we have an init, a main and an end. device_related is set to True if we have a multi-step signal container. need_response is set to true if the signal node waits on hardware (e.g. waits for a stage to confirm it has indeed moved) before proceeding.

Each of the functions that are the value entries in self.config_table dictionaries are methods of the feature object.

---

##### 9.1.1 Creating A Custom Feature Object

Each feature object is defined as a class. Creating a new feature is the same as creating any Python class, but with a few requirements. The first parameter of the __init__ function (after self) must be model, which gives the feature object full access to the **navigate** model. All the other parameters are keyword arguments and must have default values. The __init__ function should always have a config_table attribute (see *above* for a description of the config_table).

In the example below, we will create a custom feature that moves to a specified position in **navigate**'s multi-position table and calculates the sharpness of the image at this position using the Normalized DCT Shannon Entropy metric. An example __init__() function for our FeatureExample class is below.

```
from navigate.model.analysis.image_contrast import fast_normalized_dct_shannon_entropy

class FeatureExample:

    def __init__(self, model, position_id=0):
        self.model = model
        self.position_id = position_id

        self.config_table = {
            "signal": {
                    "init": self.pre_func_signal,
                    "main": self.in_func_signal,
            },
            "data": {
                    "main": self.in_func_data,
            },
            "node": {
                "device_related": True,
            }
        }
```

1. Get multi-position table position from the GUI.

   All the GUI parameters are stored in model.configuration["experiment"] during runtime. Below, we create a function to get all the position stored at position_id from the multi-position table in the GUI when we launch our feature.

```
def pre_func_signal(self):
    positions = self.model.configuration["experiment"]["MultiPositions"]
```

(continues on next page)

(continued from previous page)

```
    if self.position_id < len(positions):
        self.target_position = positions[self.position_id]
    else:
        current_position = self.model.get_stage_position()
        self.target_position = dict([(axis[:-4], value) for axis, value in current_
↪position.items()])
```

More GUI parameters can be found in experiment.yml

2. Use the stage to move to this position.

   Now, we move stage to the `target_position` we grabbed from the multi-position table.

```
def in_func_signal(self):
    pos = dict([(f"{axis}_abs", value) for axis, value in self.target_position.
↪items()])
    self.model.move_stage(pos, wait_until_done=True)
```

3. Take a picture and process the resulting image.

   In parallel with our signal function call, the camera will acquire an image. The image captured by the camera will be stored in the `model data_buffer`. The `data` functions run after an image is acquired. We add code to deal with this image in the `"main"` data function. Here, we calculate the Shannon entropy of the image.

```
def in_func_data(self, frame_ids):
    for id in frame_ids:
        image = self.model.data_buffer[id]
        entropy = fast_normalized_dct_shannon_entropy(image,
            psf_support_diameter_xy=3)
        print("entropy of image:", id, entropy)
```

Now, we've create a whole new feature and can use it as we wish.

##### 9.1.2 How to interact with other devices

We interact with all devices through `self.model.active_microscope`. Here is an example to open shutter:

```
self.model.active_microscope.shutter.open_shutter()
```

##### 9.1.3 How to pause and resume data threads in the model

The image data acquired from the camera are handled in an independent thread. As such, the *signal* and *data* operations by default run in parallel and do not block each other. Sometimes, we want to be sure a device is ready or has moved. For example, in `FeatureExample`, we have no guarantee that the stage finished moving before the image was taken. The `wait_until_done` call only blocks the signal thread from progressing before the stage finishes its move. To ensure the data thread also waits, we need to pause the data thread until the stage is ready.

Here is an example of how we can pause and resume the data thread:

```
self.model.pause_data_thread()

```

(continues on next page)

(continued from previous page)

```
self.model.move_stage(pos, wait_until_done=True)
# ...

self.model.resume_data_thread()
```

We can of course replace `self.model.move_stage(pos, wait_until_done=True)` with whatever task we want to wait for before resuming image acquisition.

Model functions can be found in the API.

#### 9.2 Custom Feature Lists

The **navigate** software allows you to chain feature objects into lists to build acquisition workflows.

##### 9.2.1 Creating a Custom Feature List in Python

To create a customized feature list, follow these steps:

1. Import the necessary modules:

   ```
   from navigate.tools.decorators import FeatureList
   from navigate.model.features.feature_related_functions import *
   ```

   `FeatureList` is a decorator that registers the list of features. `feature_related_functions` contains convenience imports that allow us to call `PrepareNextChannel` instead of `navigate.model.features.common_features.PrepareNextChannel`. These functions make for more readable code.

1. Create the feature list.

   ```
   @FeatureList
   def feature_example():
       return [
           (
               {"name": PrepareNextChannel},
               {
                   "name": LoopByCount,
                   "args": ("experiment.MicroscopeState.selected_channels",),
               },
           )
       ]
   ```

   In this example, the feature list takes one image per selected channel in the GUI. `PrepareNextChannel` sets up the channel and `LoopByCount` call this setup once per selected channel.

1. Now, open **navigate**.

2. Go to the *Features* menu.

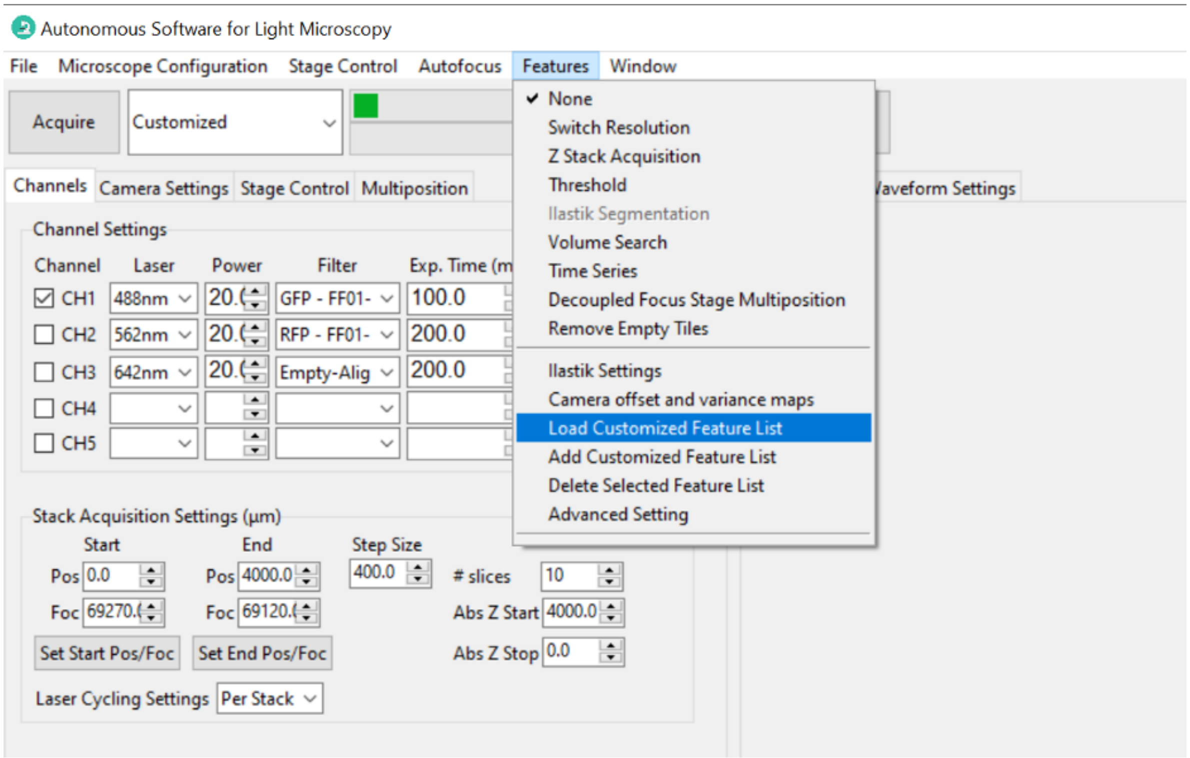

3. Import the customized feature. Select *Add Custom Feature List* from the *Features* menu. A dialog box will appear, allowing you to select the Python file containing your customized feature list function.

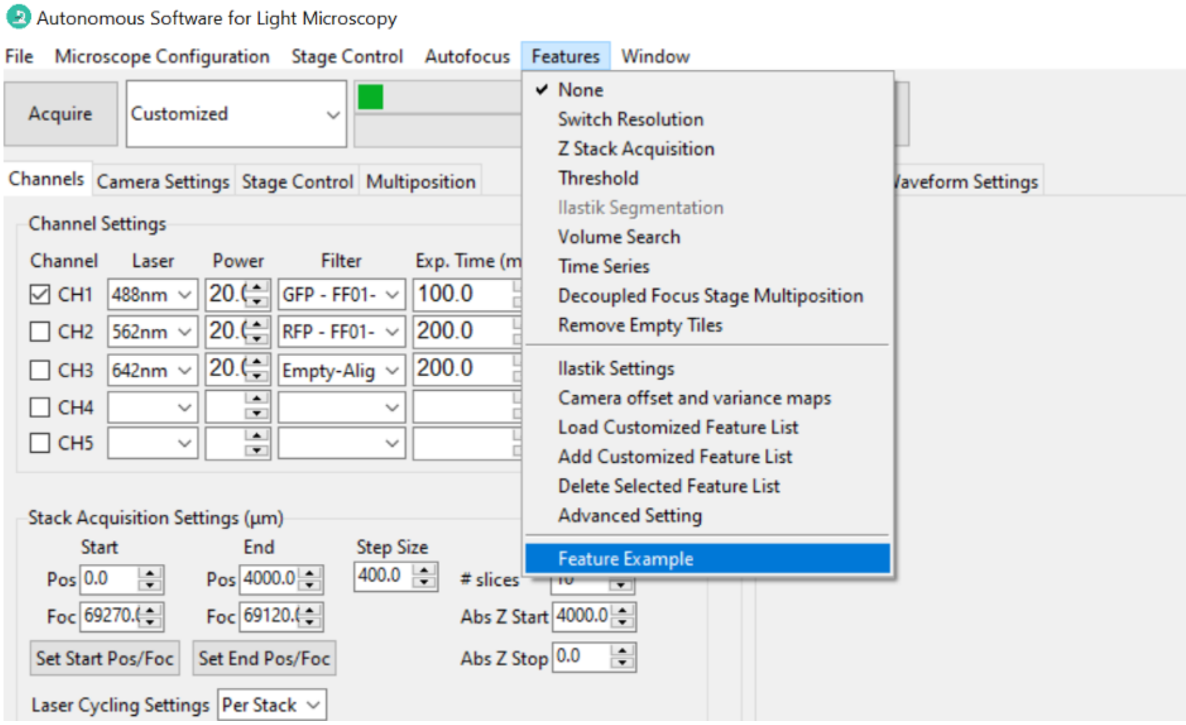

4. Choose the Python file containing your customized feature list function. **navigate** will load the specified feature list, making it available for use in your experiments and analyses. It will appear at the bottom of the *Features* menu.

CHAPTER

TEN

### PLUGIN ARCHITECTURE

**navigate** is designed with extensibility. Users can seamlessly integrate custom GUI plugins, device plugins, new feature plugins, and even define specific acquisition modes.

#### 10.1 Introduction to Plugins

The **navigate plugin system** gives users the flexibility to extend its functionality according to their specific needs. **navigate** will load plugins automatically and users can use their plugins with **navigate** seamlessly.

#### 10.2 Installing a Plugin

Once you've built a plugin or downloaded a **navigate** plugin, you can easily install it. Here, we have downloaded the Navigate Confocal-Projection Plugin.

1. You can install the **navigate-confocal-projection** plugin by selecting the menu *Plugins* → *Install Plugin*.

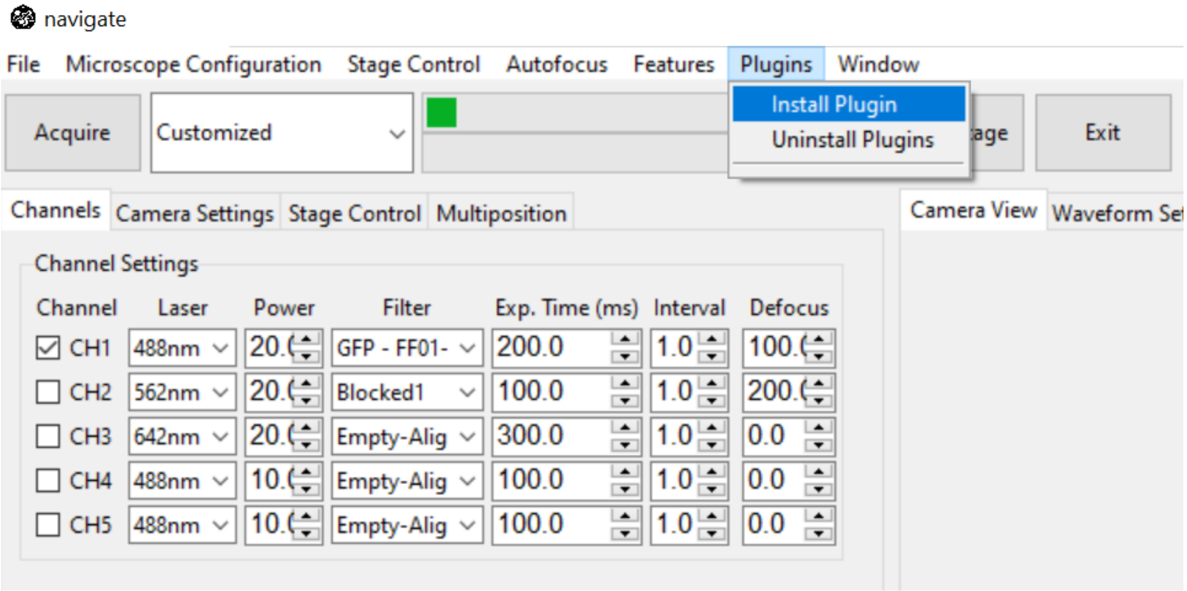

2. Select the folder *ConfocalProjectionPlugin* and click *Select*. The plugin is now installed.

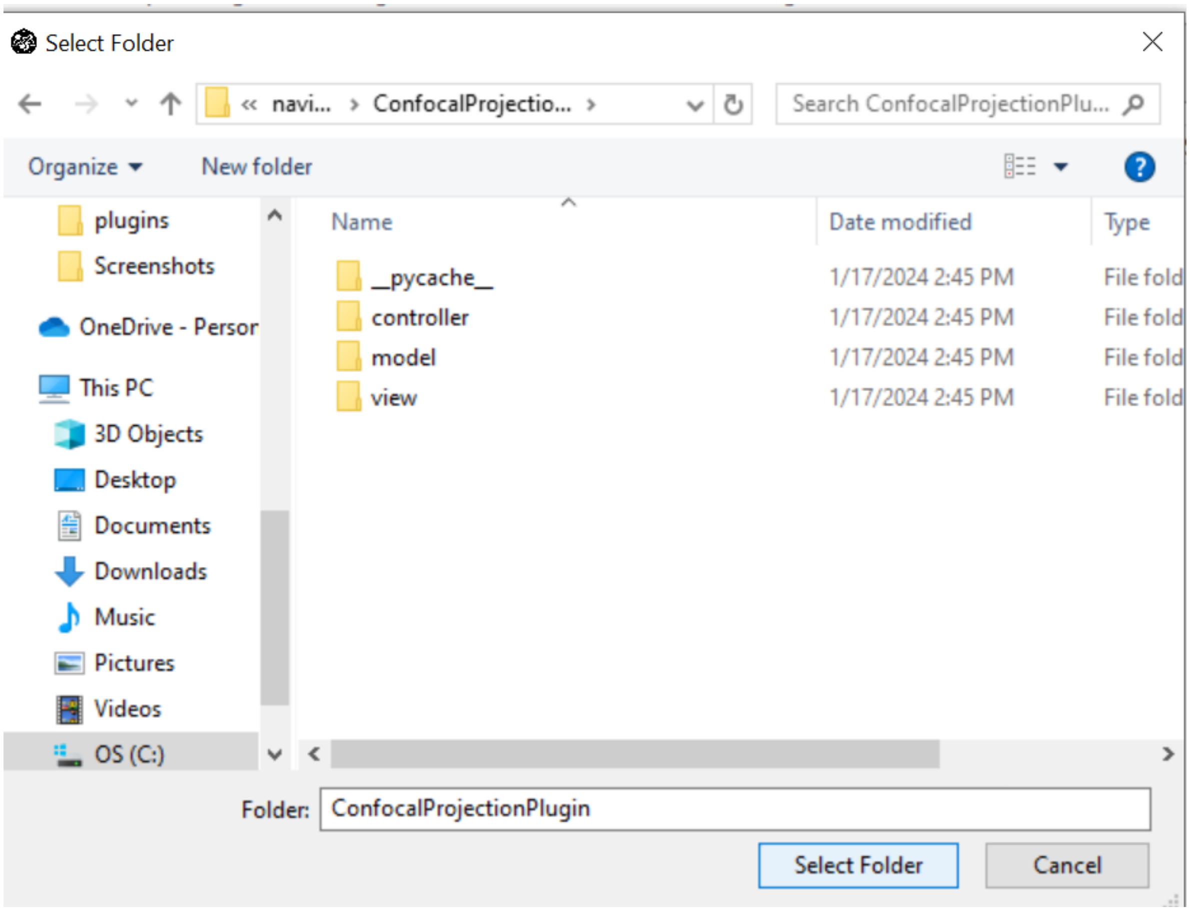

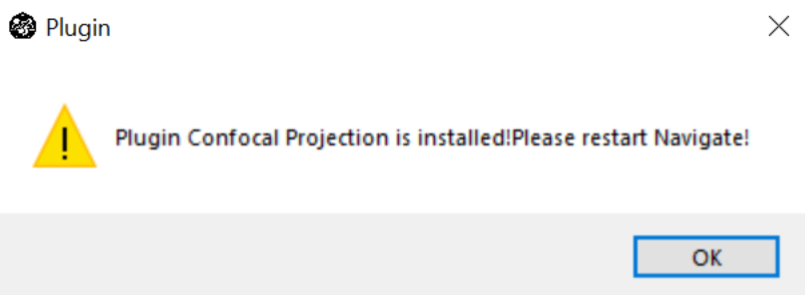

3. Restart **navigate** to use this installed plugin.

#### 10.3 Uninstalling a Plugin

Uninstalling a plugin is very easy.

1. Select *Plugins* → *Uninstall Plugins*. This will open a popup window where you can see all of the currently installed plugins.

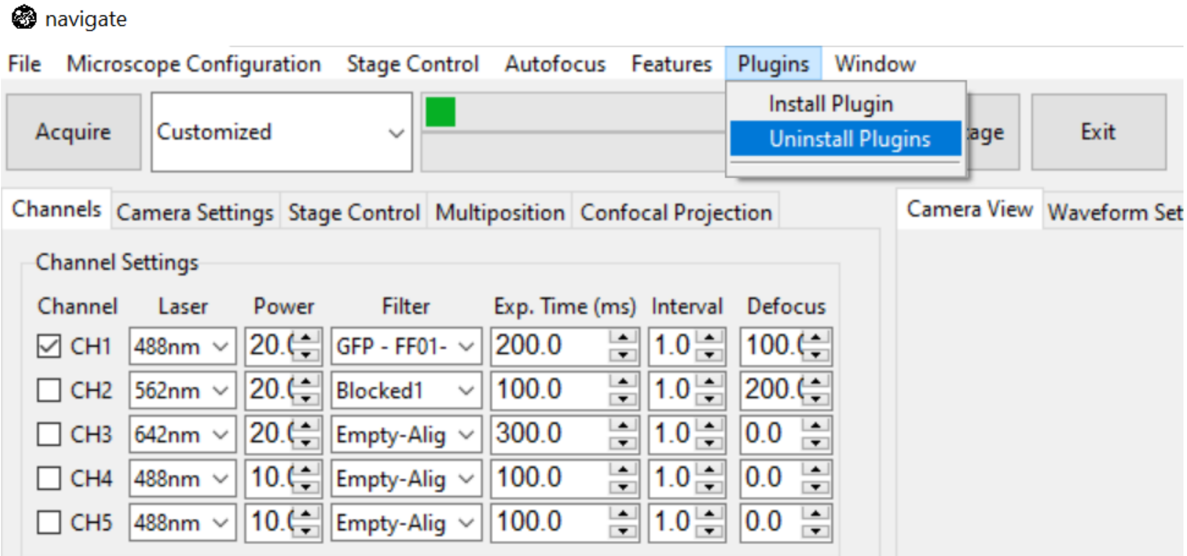

2. Select the plugin you want to uninstall.

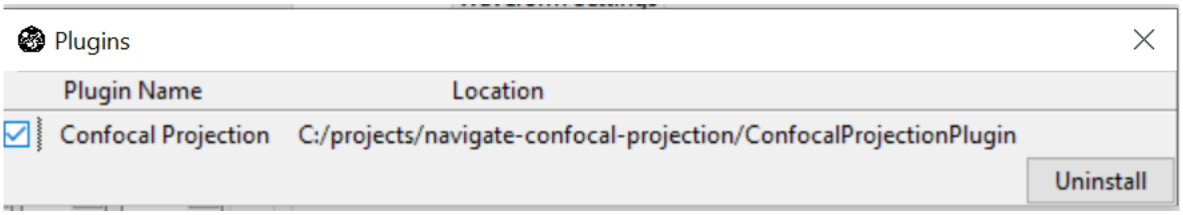

3. Click *Uninstall*.

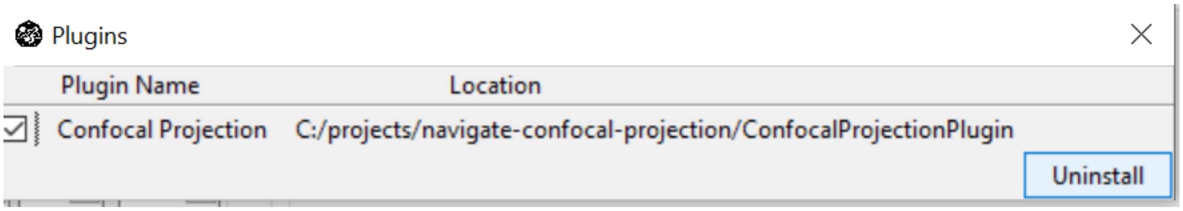

4. Restart **navigate** to fully remove the uninstalled plugin.

#### 10.4 Designing a Plugin

##### 10.4.1 Using a Plugin Template

A comprehensive **plugin template** is provided. Users could download the **plugin template** from github and build plugins on it.

Plugin Structure:

```
plugin_name/
     ├── controller/
     │    ├── plugin_name_controller.py
     |    ├── ...
     ├── model/
     |    ├── devices/
     │    │    └── plugin_device/
     │    │         ├── device_startup_functions.py
     │    │         ├── plugin_device.py
     │    │         └── synthetic_plugin_device.py
     │    └── features/
     │              ├── plugin_feature.py
     │              ├── ...
     │
     ├── view/
     |    ├── plugin_name_frame.py
     |    ├── ...
     │
     ├── feature_list.py
     ├── plugin_acquisition_mode.py
     └── plugin_config.yml
```

**Note:** The template shows a plugin with GUI, device, feature, feature_list and acquisition mode. If your plugin only incorporates some of these components, you should remove unused folders and files.

##### 10.4.2 Plugin Configuration

There should always have a `plugin_config.yml` file under the plugin folder, which tells **navigate** the plugin name, the GUI as a Tab or Popup and custom acquisition mode name. A typical plugin config is:

```
name: Plugin Name
view: Popup # or Tab
acquisition_modes:
  - name: Plugin Acquisition
    file_name: plugin_acquisition_mode.py
```

##### 10.4.3 Plugin GUI Elements

**navigate** supports plugins with their own GUIs. A custom plugin GUI can be integrated as a tab or a popup. Users should specify a view option in plugin_config.yml. If it is a popup, users can find the plugin under the *Plugins* menu in the **navigate** window. If it is a tab, it will appear next to the *Settings Notebooks*.

When creating a new plugin with a GUI, ensure that the plugin name is consistent with the naming conventions for the associated Python files (plugin_name_controller.py and plugin_name_frame.py). Both Python filenames should be in lowercase.

For example, if your plugin is named "My Plugin" (there is a space in between), the associated Python files should be named: my_plugin_frame.py and my_plugin_controller.py.

---

##### 10.4.4 Plugin Devices

The **navigate** plugin architecture allows you to integrate new hardware device. There can be more than one device inside a plugin. If they are different kinds of device, please put them into different folders. For each kind of device, there should be a device_startup_functions.py telling **navigate** how to start the device and indicating the reference name of the device to be used in configuration.yaml.

Device type name and reference name are given as following:

```
DEVICE_TYPE_NAME = "plugin_device"  # Same as in configuration.yaml, for example "stage",
↪ "filter_wheel", "remote_focus_device"...
DEVICE_REF_LIST = ["type", "serial_number"]
```

A function to load the device connection should be given,

```
def load_device(configuration, is_synthetic=False):
    # ...
    return device_connection
```

A function to start the device should be given,

```
def start_device(microscope_name, device_connection, configuration, is_synthetic=False):
    # ...
    return device_object
```

The template for device_startup_functions.py can be found in the plugin template.

---

##### 10.4.5 Plugin Features

**navigate** allows users to add new features. New feature objects and feature lists can each be a plugin or components of a plugin. Features and feature lists are automatically loaded into **navigate**.

Please visit here for details about how to build a new feature object and feature list.

---

##### 10.4.6 Custom Acquisition Modes

Navigate offers seamless support for custom acquisition modes, and registering a new mode is straightforward.

1. Download the template for plugin_acquisition_mode.py

2. Update the `feature_list`.

```
@AcquisitionMode
class PluginAcquisitionMode:
    def __init__(self, name):
        self.acquisition_mode = name

        self.feature_list = [
            # update here
        ]
```

3. Update the functions.

Users should tell **navigate** what to do before and after acquisition using the following functions.

```
def prepare_acquisition_controller(self, controller):
    # update here

def end_acquisition_controller(self, controller):
    # update here

def prepare_acquisition_model(self, model):
    # update here

def end_acquisition_model(self, model):
    # update here
```

4. Register the acquisition mode in `plugin_config.yml`.

```
acquisition_modes:
    - name: Custom Acquisition
      file_name: plugin_acquisition_mode.py
```

---

For more plugin examples, please visit navigate and Navigate Plugins.
